# Questioning Neanderthal admixture: on models, robustness and consensus in human evolution

**DOI:** 10.1101/2023.04.05.535686

**Authors:** Rémi Tournebize, Lounès Chikhi

**Author notes:** Laboratoire Évolution & Diversité Biologique (EDB UMR 5174), Université de Toulouse Midi-Pyrénées, CNRS, IRD, UPS, Toulouse Cedex 9, France.

## Abstract

Genomic and ancient DNA data have revolutionized palaeoanthropology and our vision of human evolution, with indisputable landmarks like the sequencing of Neanderthal and Denisovan genomes. Yet, using genetic data to identify, date and quantify evolutionary events—like ancient bottlenecks or admixture—is not straightforward, as inferences may depend on model assumptions. In the last two decades, the idea that Neanderthals and members of the *Homo sapiens* lineage interbred has gained momentum. From the status of unlikely theory, it has reached consensus among human evolutionary biologists. This theory is mainly supported by statistical approaches that depend on demographic models minimizing or ignoring population structure, despite its widespread occurrence and the fact that when ignored, population structure can lead to infer spurious demographic events. We simulated genomic data under a structured and admixture-free model of human evolution, and found that all the tested admixture approaches identified long Neanderthal fragments in our simulated genomes and an admixture event that never took place. We also observed that several published admixture models failed to predict important empirical diversity or admixture statistics, and that our model was best at predicting these statistics jointly. Our results suggest that models accounting for population structure are fundamental to improve our understanding of human evolution, and that admixture between Neanderthals and *Homo sapiens* needs to be re-evaluated in the light of structured models. Beyond the Neanderthal case, we argue that ancient hybridization events, which are increasingly documented in many species, including with other hominins, may also benefit from such reevaluation.

**Significance statement:** The idea that Neanderthals and some ancestral *Homo sapiens* populations interbred has gained momentum in the last two decades. Yet, this theory is mainly supported by statistical approaches that assume highly simplified models of hominin evolution. A major issue is that these methods have been poorly tested in the context of population structure, despite its widespread occurrence in many vertebrate species. We simulated data under a structured model and found that all tested methods identified spurious admixture events, suggesting a lack of robustness to population structure. Besides, our structured model was better at predicting several key genomic statistics than the tested admixture models. This suggests that admixture should be re-evaluated in the light of population structure, in hominins and beyond.

## Introduction

Population structure is increasingly recognized by palaeo-anthropologists, geneticists and palaeo-climatologists as a fundamental feature of *Homo sapiens* (*Hs*) evolution—not only in the recent, but also in the ancient past (Beaumont, 2004; Harding and McVean, 2004; Ramachandran et al., 2005; Mazet et al., 2016; Rodríguez et al., 2018; Arredondo et al., 2021; Ragsdale et al., 2022). Periods of geographical contraction or expansion of populations, of increased or decreased connectivity among regions of the African continent have likely been at the heart of this history, possibly in relation with climate and habitat change during the Pleistocene (Scerri et al., 2018). Yet, the integration of modern and ancient genomic data into one general framework to infer *Hs* demographic history is among the greatest challenges facing theoretical population geneticists and palaeo-anthropologists (Goldstein and Chikhi, 2002; Scerri et al., 2018, 2019).

Given that our closest relatives, the African great apes, are spatially and genetically structured over wide regions of Africa (Prado-Martinez et al., 2013; Lester et al., 2021), it seems reasonable to consider that *Hs* and other hominins have been structured across different but similarly wide regions of Africa during significant periods of their evolution. Consequently, it seems important to integrate intra-continental population structure in the models used for demographic inference if we wish to understand and properly represent the recent evolutionary history of hominins (Scerri et al., 2019). Currently, most inferential approaches applied to human genetic data tend to either ignore population structure altogether (Li and Durbin, 2011) or assume that it is mainly significant between continental populations (Gutenkunst et al., 2009), with a few exceptions (Currat and Excoffier, 2004).

The importance of model-based approaches to address these challenges has been stressed for several decades (Beaumont, 2004; Beaumont et al., 2002; Goldstein and Chikhi, 2002; Gerbault et al., 2014). Demographic models should be as simple as possible to avoid over-parameterization but should also capture important aspects of a necessarily complex history (Goldstein and Chikhi, 2002; Scerri et al., 2018, 2019; Chikhi, 2023). In addition, some features of the actual demographic dynamics can be confounding for inferential methods, whose robustness is another often overlooked issue. For instance, when ignoring population structure, researchers can identify and quantify with great precision population size changes even when the inferred changes never occurred (Wakeley, 1999; Beaumont, 2004; Chikhi et al., 2010; Mazet et al., 2016; Battey et al., 2020). Besides, inferential methods used to detect important events such as admixture or bottlenecks are more likely to be convincing if they can also predict realistic values for other well-established measures of human genetic diversity.

In the last two decades, ancient DNA (aDNA) studies have revolutionized palaeo-anthropology with the publication of thousands of ancient genomes, and notably, with the sequencing of ancient hominins like Neanderthals (*Hn*). Remarkably, researchers identified that a small but significant amount of DNA stretches sampled in some modern *Hs* populations were genetically closer to *Hn* alleles than to alleles found in other contemporaneous *Hs* populations (Green et al., 2010). This pattern has been interpreted as a result of admixture or interbreeding between *Hn* and some *Hs* populations around 50–60 thousand years ago (kya) (Sankararaman et al., 2012). While there is some debate regarding the tempo and mode of this archaic introgression, the current consensus is that some modern populations harbor Neanderthal DNA as a result of a recent admixture event. A few discordant voices have suggested that part of the admixture signal could be explained by spatial population structure leading to isolation by distance and incomplete lineage sorting (ILS) in some *Hs* populations (Eriksson and Manica, 2012, 2014). Others added that current admixture models failed to explain significant patterns of genomic diversity which can be explained without admixture when population structure is accounted for (Rodríguez et al., 2018). Under such admixture-free but structured models, the DNA tracts identified as introgressed should not be attributed to an actual introgression event: they are the result of shared ancestry and are inherited from a structured common ancestor to both *Hn* and *Hs*.

It is therefore important to ask whether admixture estimates are as robust to model departures (viz. intra-continental population structure) as it has been generally argued (Peter, 2016; Racimo et al., 2015). A related and open problem is whether current models of *Hn-Hs* admixture are compatible among themselves and whether they can reproduce well-established genome-wide patterns of human genetic diversity and differentiation.

Several statistics have been developed to detect and estimate genetic admixture or introgression (Sankararaman, 2020). We can, as a first approximation, classify these statistics as frequency- or LD-based. Frequency-based statistics compare allelic composition in *Hn* and *Hs* populations to identify patterns that are unlikely deriving from ILS but are more likely resulting from a recent admixture event. LD-based statistics aim at identifying stretches of DNA shared between *Hn* and specific *Hs* populations, that are too long to have been inherited independently from a common ancestor. Yet, for both frequency- or LD-based statistics, the likelihood that these shared SNPs or long DNA fragments derive from an admixture event rather than from ILS depends on the specificities of the assumed demographic, mutation (Amos, 2020) and recombination models. For instance, the *D* statistic, basis of the ABBA/BABA test, is expected to be zero if the contribution of Neanderthals to European and African *sapiens* is the same (Green et al., 2010). This expectation is deduced from a model where Europeans and Africans are modeled as continent-wide panmictic populations since their split, tens of kya, deriving from a panmictic (non-structured) ancestral population. An implicit assumption is that departures from this simple model should not influence the value and statistical significance of *D* in any notable way.

Eriksson and Manica (2012) demonstrated that this assumption was incorrect and that the *D* statistic was in fact not robust to spatial population structure. By simulating a one-dimensional (1D) stepping-stone model with African, Eurasian *Hs* and *Hn* demes, they estimated *D* values similar to those observed in real data even though they did not model admixture between *Hn* and *Hs* populations. In 2014, the same authors showed that the *DCFS* (doubly-conditioned frequency spectrum, Yang et al. (2012)) was not robust to population structure either. Thus, two statistics designed to detect admixture could be reproduced in the absence of admixture when population structure was modeled. Since then, new admixture statistics and models have been developed, that increasingly integrate patterns of LD and haplotype information, as these were thought to provide clearer support for introgression (Racimo et al., 2015; Sankararaman, 2020). In addition, the discovery and analysis of ancient *Hs* specimens, like Oase1, led to the identification of high *D* values and long Neanderthal-like DNA stretches that were suggested as further evidence of admixture (Fu et al., 2015). Nevertheless, little effort has been made to test the robustness to population structure of more recent admixture statistics, neither in present-day nor in ancient *Hs* samples.

The work presented here aims at clarifying the role of both admixture and population structure in human evolution and is inspired by two lines of research. The first is exemplified by the seminal theoretical work developed around the structured coalescent by Herbots (1994); Wilkinson-Herbots (1998) and others (Notohara, 1990; Takahata, 1991; Hudson, 1990; Tajima, 1990), who showed that gene trees sampled in structured populations have properties that no stationary panmictic model can replicate (Wakeley, 1999; Beaumont, 2004; Chikhi et al., 2010; Mazet et al., 2016; Rodríguez et al., 2018). The second line of research follows the critical work of Eriksson and Manica (2012, 2014). We extend their original model and approach to investigate the behaviour of additional frequency-based statistics, but also of LD-based admixture statistics. In particular, we show that, contrary to common claims, long fragments interpreted as introgressed from Neanderthal can be detected without any actual *Hn* admixture, once spatial structure is accounted for. We compare our results with those of eleven demographic models published in the literature and find that our model generally best predicts the statistics jointly. We acknowledge that admixture models could provide interesting avenues for research regarding past human evolution but predict that admixture estimates will significantly decrease as models increasingly integrate intra-continental population structure. We thus argue that admixture between *Hs* and *Hn* needs to be re-evaluated in the light of structured models.

## Results

### A model with intra-continental structure and without admixture: specificities and results

The demographic scenarios that we explored belong to a family of 1D stepping-stone models (Figure 1 for a simplified version, Figure S1 for the full model), inspired by the original model of Eriksson and Manica (2012, 2014) and by archaeological and palaeo-anthropological literature (Scerri et al., 2018, 2019). Still, our model differs from Eriksson and Manica’s (2012, 2014) in several ways (see Notes Sl.l for details). For instance, their model assumed a single African stepping-stone, whereas we found that an ancient bipartite metapopulation structure across the African continent significantly improved the prediction of several statistics (see Ragsdale et al. (2022) for a similar and independent suggestion of an ancient split in the hominin lineage). Specifically, our model assumed two sets of interconnected demes, where one ancient set of 10 demes (metapopulation M_A_) generated a second set of 10 demes (metapopulation M_B_) through a serial colonization process in a period that was allowed to vary from 500 kya to 9 Mya. Our simulations suggested that this event occurred prior to 2 Mya, and that this bipartite structure may have been maintained during most of the Pleistocene, with *F_ST_* values varying between 0.03 and 0.89 (Notes S6).

**Figure 1.**
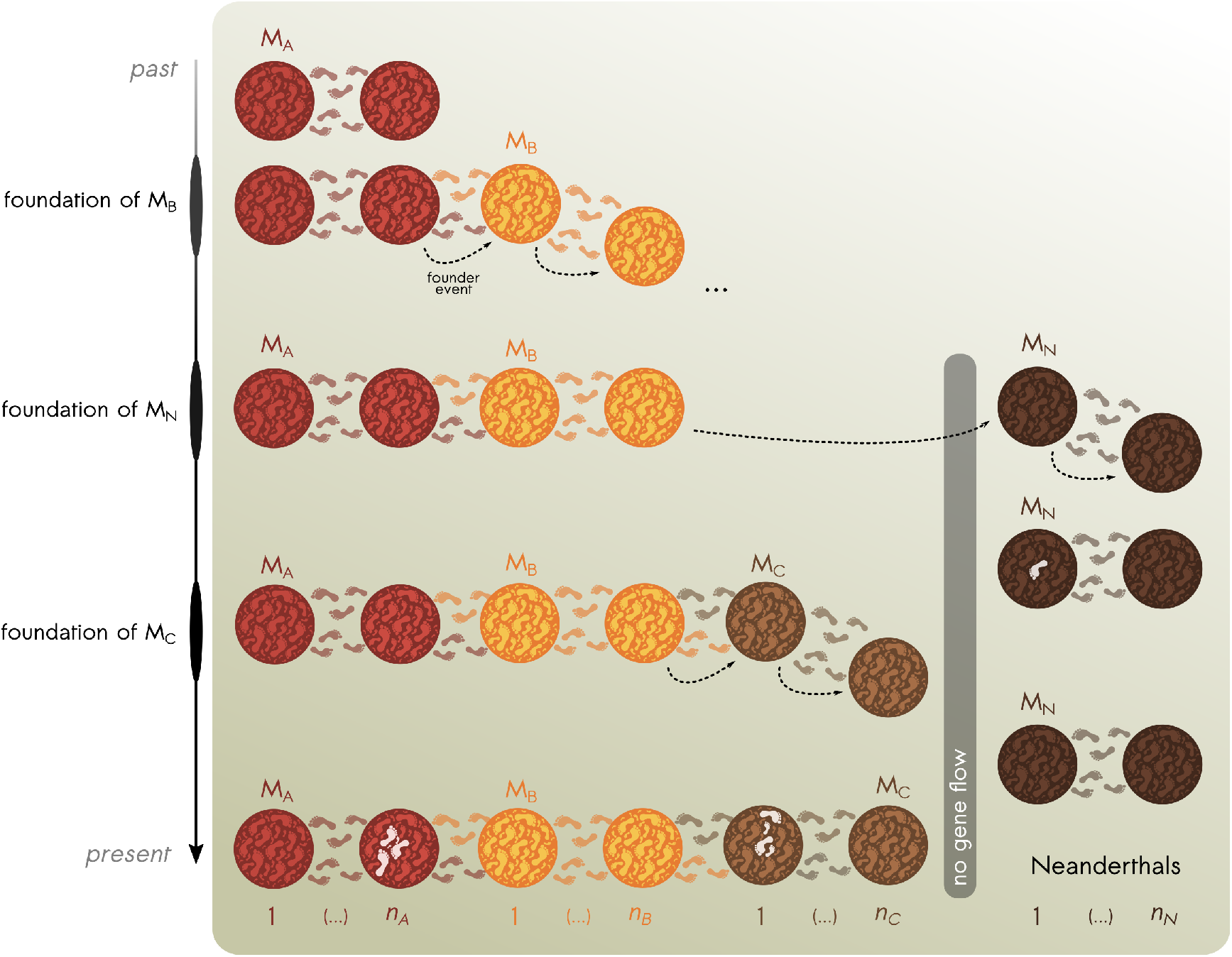
Simplified representation of the 1D structured model considered in this study. Time flows from top (past) to bottom (present), with an initial metapopulation M_A_ consisting of *n_A_* (=10) demes exchanging migrants with their neighbours. At some point in the past, the rightmost deme of M_A_ founds a new metapopulation M_B_ of *n_B_* (=10) demes, with which it will continue exchanging migrants till the present. Later, the rightmost deme of M_B_ founds the metapopulation M_N_ of *n_N_* (=10) demes which will become Neanderthals. The M_N_ metapopulation will never exchange migrants with any other deme from the other metapopulations. Closer towards the present, the rightmost deme of M_B_ founds M_C_ which corresponds to the expansion of *H. sapiens* towards Eurasia. White feet represent the sampled metapopulations (not the specific demes) for respective sampling times. The location of the sampled demes (within the corresponding metapopulations) is a random variable (see Notes S1.1). Fifty individuals are sampled in M_A_ and in M_C_ to represent modern-day YRI and CEU samples respectively. For the Neanderthals (M_N_), one individual is sampled at 50 kya. More details about the models can be found in the Materials and Methods section and in Notes S1.1.

Following the formation of this bipartite African structure among the species ancestral to *Hs* and *Hn*, we modeled the ancestors of *Hn* as a 1D stepping-stone metapopulation of 10 demes, created when individuals from the African M_B_ demes serially colonized Eurasia. Despite a wide prior for the split between the ancestors of *Hs* and *Hn* (from 400 kya to 1 Mya), the best scenarios tended to favor a split around 650 kya (see Notes S5). After this split, our model assumed that M_A_ and M_B_ would give rise to *Hs*, whereas the Eurasian metapopulation, M_N_, would become *Hn*. A later serial colonization of Eurasia by *Hs* was also simulated from the last M_B_ deme, with gene flow connecting the new *Hs* Eurasian metapopulation M_C_ with the African *Hs* metapopulation M_B_ (itself connected to the *Hs* metapopulation M_A_). Note however that none of the *Hs* metapopulations (M_A_, M_B_ and M_C_) ever exchanged gene flow with the *Hn* metapopulation (M_N_). In brief, our model does not allow admixture between *Hs* and *Hn*. The period providing the best support for *Hs* expansion in Eurasia was estimated around 40-65 kya, in agreement with current models and archaeological data (cf. published models in Notes S8).

We must stress here that our objective was not to find the best model explaining the data but rather, to identify a reasonable set of demographic parameters that could reproduce important summary statistics within a wide parameter space. To do so, we used a set of ten summary statistics (*AFS_CEU_*, *AFS_YRI_*, *F_ST_*, *D*, *DCFS*, *L_S_*, *π_YRI_*, *T_LD_*, *σ_T,LD_*, see Notes S1.4 for details) (Gutenkunst et al., 2009; Bhatia et al., 2013; Green et al., 2010; Yang et al., 2012; Browning et al., 2018; Sankararaman et al., 2012; Keinan et al., 2009) as a guide for the exploration of the parameter space (Table S2) and for the identification of the twenty most promising scenarios out of a million simulations. Some of these statistics (*D*, *DCFS*, *L_S_*, *T_LD_*) are generally used to detect or quantify admixture. We additionally computed three statistics (the *S’*-related match rate *m_S’_* (Browning et al., 2018) and two *PSMC* curves (Li and Durbin, 2011) for genomes sampled at present in M_A_ and M_C_ demes) to see if our model could predict observed values or curves from real present-day samples.

### Explaining admixture statistics without admixture

Despite important differences between our model and Eriksson and Manica (2012, 2014), our simulations confirmed their original findings (Figure 2F), viz. that large and realistic *D* values can be reproduced after accounting for population structure in the absence of *Hn* admixture. Our model generated *D* values that varied from −1.2% to 10.8% (IQR95), with an average at 4.7% similar to the observed value of 4.6% (Figure 2F). Using the mean relative absolute error (MRAE) as a sealed measure of discrepancy between model predictions and observed values (Notes S1.7), we estimated MRAE=52.4%±47.2% (mean ± SD). Similarly, we obtained non linear *DCFS* curves, which are typically suggested to be indicative of admixture (Yang et al., 2012) (Figure 2G).

**Figure 2.**
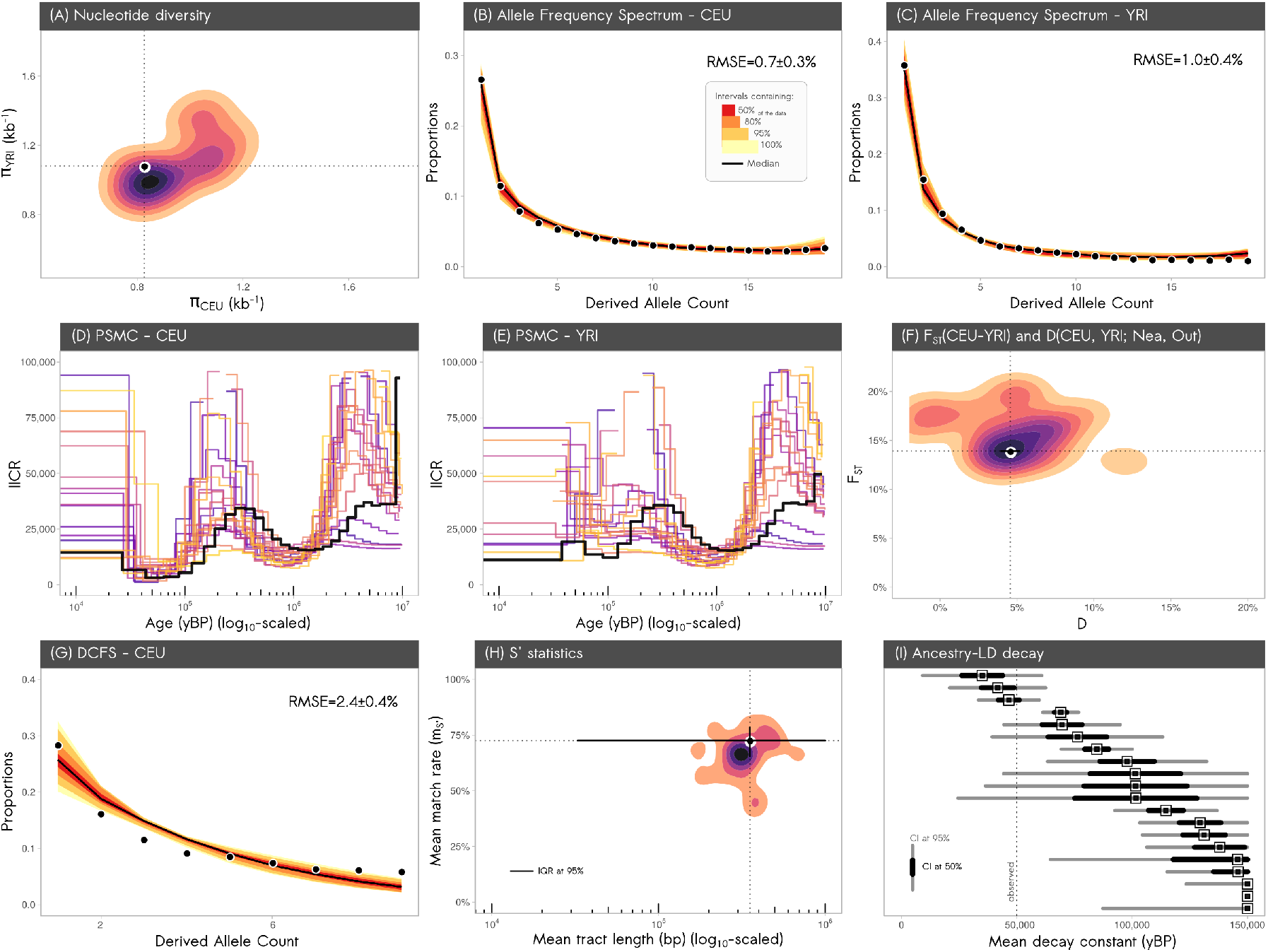
Performance of our admixture-free model compared to observed data (black) for the twelve statistics used in this study. The statistical distributions were obtained from the twenty accepted runs of our model, after simulating twenty chromosomes of 30 Mbp (uniform recombination). **Abbreviations.** CI: confidence interval; IQR: interquantile range; IICR: inverse instantaneous coalescence rate. (A) Nucleotide diversity in CEU and YRI. We represent the 2D density (Gaussian kernel) of the joint nucleotide diversities π estimated in simulated CEU and YRI samples. The black dot represents the observed values from real data. (B) Normalized CEU *AFS*. The black dots represent the observed *AFS* using ten real CEU individuals. The colors represent the 50%, 80%, 95% and 100% interquantile ranges of the normalized derived allele frequency spectrum calculated for ten simulated CEU samples across the twenty selected runs. The black line is the median across the simulated *AFSs*. RMSE is the root mean squared error and is given in percentage. **(C)** Normalized YRI *AFS*. Same as (B), for ten YRI samples. **(D)** CEU *PSMC* curves. The black curve is the observed CEU *PSMC* whereas the coloured PSMC curves were obtained from the twenty selected runs. The *x*-axis (log_10_-scaled) is expressed in years before present (with *μ*=1.2×10^-8^ and g=25 years). **(E)** YRI *PSMC* curves. Same as (D), for a single YRI sample per selected run. **(F)** Joint distribution of the *F_ST_* and *D*-statistic. The D-statistic (*x*-axis) was calculated as *D*(50 CEU, 50 YRI; 1 Neanderthal, Ancestral reference) and the *F_ST_* (*y*-axis) was computed between the CEU and YRI samples. Both statistics were computed for each of the twenty runs and plotted jointly using a 2D density (Gaussian kernel). **(G)** CEU *DCFS*. As in panel (B), an interquantile ribbon plot was obtained for the doubly conditioned site frequency spectra across the twenty selected runs. The *DCFS* was computed for five CEU individuals. The black line is the median *DCFS* across the twenty runs. The black dots represent the observed *DCFS*. **(H)** Joint distribution of mean Neanderthal fragments lengths and match rates. A 2D density (Gaussian kernel) of the distribution for the mean length (*x*-axis, in bp, log_10_-scaled) and the mean match rate *m_S’_* (*y*-axis) is represented for the putative Neanderthal segments estimated with *S’* in the 50 simulated CEU samples. The black dot represents the estimates from the observed data (with associated 95% IQR). (I) Ancestry-*LD* decayconstant. The *x*-axis represents the exponential decay constant of the ancestry-*LD* curve, and the twenty selected runs are represented on the *y*-axis. The point estimates (square dots) are classically-interpreted as the age of the admixture events. The whisks represent the 50% (black thickest segment) and the 95% confidence intervals (gray segment) around the jackknife mean (calculated using a weighted jackknife procedure). The point estimate of the decay constant computed from real data is -’--50 kya (vertical dashed line).

Our simulations further extend the studies of Eriksson and Manica (2012, 2014) by showing that empirical LD-based statistics can also be reproduced. For instance, using the *S’* approach (Browning et al., 2018), we identified large introgressed Neanderthal fragments in our simulated samples (Figure 2H). With an average length of 391 kb (IQR95=175—978 kb; MRAE=36.2%±55.3%) for the simulated genomes, which included the value computed from the observed data (354 kb), our model was thus able to explain empirical results and could generate large fragments (>500 kb). In addition, our model correctly predicted the mean empirical introgression rate (1.17%) estimated with a conditional random field (*CRF*). which fell within the IQR95 of our simulations (0.1%—1.7%; MRAE=51.9%±30.7%).

Another important admixture statistic, *m_S’_*, estimates the match rate between the *S’*-inferred ancestral alleles and the actual Neanderthal alleles within the *S’*-identified introgressed segments. This statistic is supposed to be specific to introgressed Neanderthal fragments since the tracts are first identified statistically, and only then compared to the Neanderthal sequences. In real data, the match rate was estimated as 72.6%, within the values estimated on our simulated data (average 66.4%; IQR95=44.3—83.7%; MRAE=13.3%±11.1%) (Figure 2H). We also cautiously applied the *CRF* approach to estimate the length of the Neanderthal fragments and estimated an average of 34 kb in the simulated data (IQR95=28—41 kb; MRAE=70.3%±3.5%), which was lower than the value obtained on real data (114 kb, Sankararaman et al. (2014)) (Figure S3). Using the ancestry-*LD* approach, we estimated that the decay constant was 109 kya on average (assuming a generation time of 25 years; IQR95=38—189 kya, MRAE= 129.8%±88.2%) (Figure 2I), which included the value of 48 kya inferred from observed data (Sankararaman et al., 2012). We note however that for the simulations under our structured model, *T_LD_* should reflect the simulated split time between *Hs* and *Hn* (> 415 kya for all scenarios, and ~650 kya on average for the retained models) since this is the time of the last genetic exchange between *Hs* and *Hn*. The estimated average of 109 kya is much more recent than that, questioning the classical interpretation of this statistic in the presence of population structure. In other words, these two statistics identified in our simulated *Hs* genomes: (i) a ~100 kya old admixture event that never took place, and (ii) Neanderthal fragments of ~34 kb (*CRF*), that cannot be the result of an introgression. Although our estimates based on simulated data do not perfectly match the values for the observed data, we note that when we allowed for a variable recombination rates along the genomes, the CRF-infcrrcd Neanderthal fragments in the simulated data increased in length (with an average of 75 kb; IQR95: 55—88 kb) and the ancestry-*LD* decay constant decreased to 88 kya (IQR95=23—190 kya; MRAE=97.0%±94.9%), values closer to the one inferred on real data.

Interestingly, moving away from admixture statistics, we computed the nucleotide diversity (*π*) of our simulated Eurasian and African individuals and found that our model was able to accurately predict the empirical autosomal *π* previously estimated by Keinan et al. (2009) in CEU (*π_CEU,obs_*=0.827 kb^-1^, *π_CEU,sim_*=0.917 kb^-1^, IQR95=0.77—1.16 kb^-1^, MRAE=16.5%±13.7%) and YRI (*π_YRI,obs_*=1.081 kb^-1^, *π_YRI,sim_*=1.092 kb^-1^, IQR95=0.90—1.40 kb^-1^, MRAE=11.2%±8.8%).

Lastly, *PSMC* curves inferred on empirical genomes from *Hs* individuals typically exhibit two humps (Li and Durbin, 2011; Mazet et al., 2016; Arredondo et al., 2021). Even though the *PSMC* was not used as a statistics for the exploration of the parameter space and for the selection of the twenty scenarios, our sub-models were able to reproduce the humps correctly for both simulated European and Yoruba individuals (Figure 2D,E). We note that the *PSMC* curves exhibited larger fluctuations for some of these scenarios and were not always perfectly timed with the observed ones. This mismatch however represents a secondary issue given the poorer fit of the *PSMC* curves computed under the other published models (Figure 3A,B).

**Figure 3.**
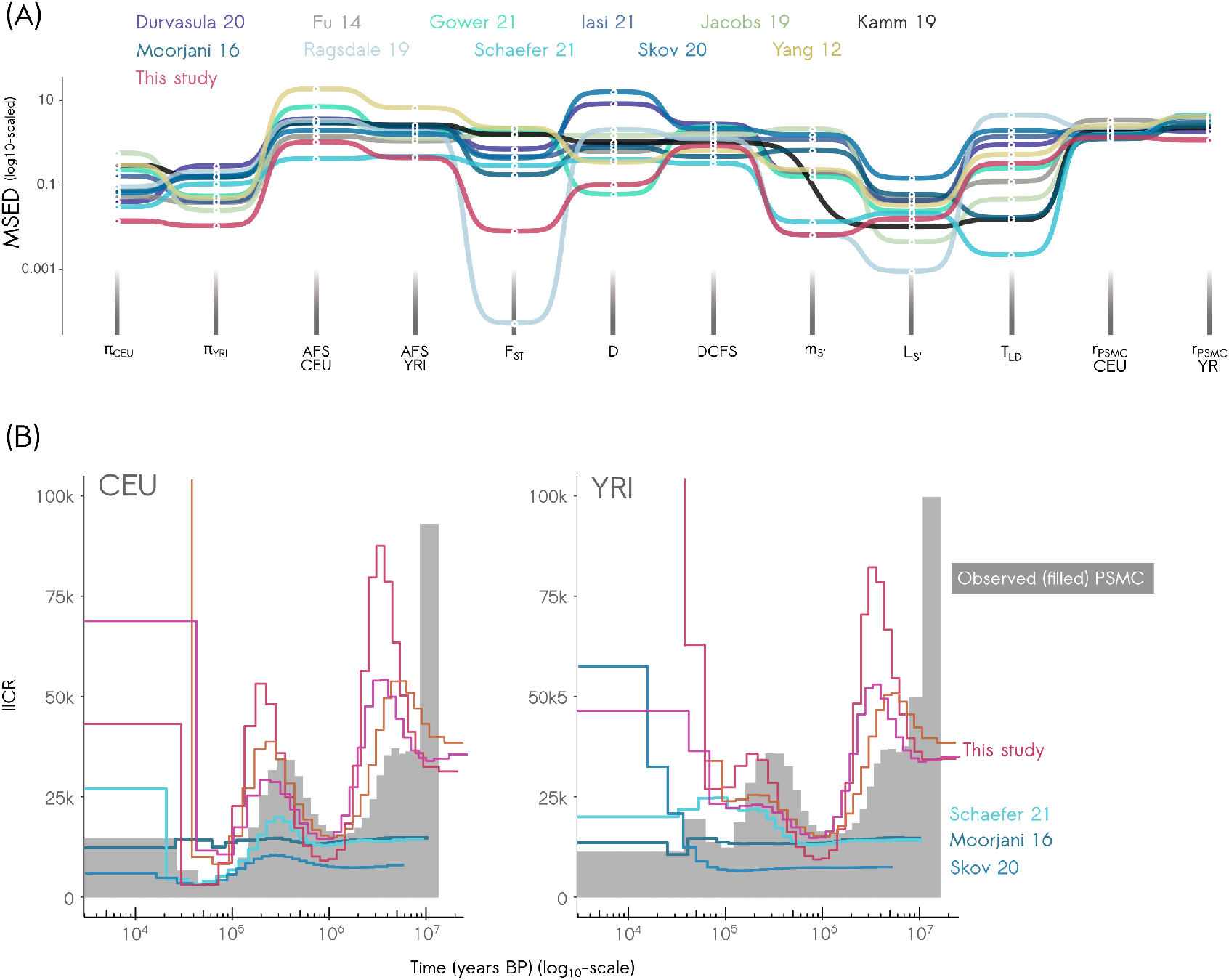
Model comparison. This figure compares the predictions of the eleven models for the twelve statistics (Figure 2) with those of our 1D structured model (red). All models except ours assume Neanderthal admixture into *Hs*. **(A)** Ranking of models for the twelve statistics. The statistics are represented in the *x*-axis, and each model is represented by a different color. The *y*-axis represents the minimum scaled Euclidean distance (MSED) to the observed statistics for twenty accepted runs from each model. The *y*-axis is log_10_-scaled. (B) Representation of the CEU (left) and YRI (right) *PSMC* curves from three among the twenty selected runs of our structured model (red shades) and from three published models. Two of these published models were chosen because they rank right after our model in the global model comparison (Schaefer et al., 2021; Moorjani et al., 2016). The *PSMC* curves for all published models are represented in Notes S8. The grey (filled) *PSMC* curves are the observed ones. Time (*x*-axis, log_10_-scaled) is given in years **BP.**

We performed additional robustness tests to account for the fact that modern datasets can typically use genomic data with different sampling strategies, coverage properties and sequencing quality. We filtered the simulated genetic data for the *S’* and *CRF* analyses so as to mimic the original analyses performed on the 1000 Genomes Project dataset with low coverage (Durbin et al., 2010) (cf. Notes S1.3.2). We found that our results were consistent irrespective of the shifts in allele frequency spectra induced by low coverage (Notes S10.2), to ancestral allele specification errors (Notes S10.3), to the sampling age of the simulated Neanderthal genomes (Notes S10.5) or to pseudodiploid genotypes (Notes S10.6).

### Comparing models with and without admixture

Using twelve statistics, we compared the performance of our structured model to that of eleven published models assuming Neanderthal admixture (Figure 3A). To this end, we simulated individual genomes of 0.6 Gbp (twenty 30-Mbp chromosomes) assuming a uniform recombination rate. Our model ranked first or second for nine statistics out of twelve, including the three that were predicted and not used for model selection (Table S4). Its worst ranking (seventh) was observed for the *T_LD_* statistics, which we saw above appears unreliable to date gene exchange under population structure. Several models performed poorly on statistics other than the ones originally used for model fitting (Notes S8). A total of six out of the eleven models failed to predict observed levels of genetic diversity in CEU or YRI (mean relative percentage error, MRPE, greater than 30% on either *π_CEU_* or *π_YRI_*), although we allowed mutation rate to vary—from 5×10^-9^ to 5×10^-8^ g^-1^ bp^-1^—to improve their ability to predict observed diversity levels. In addition to over and under-estimated nucleotide diversities, several models predicted biased *F_ST_* estimates (MRPE to true value from −91% for Yang et al. (2012) model up to 87% for Gower et al. (2021)). Also, ten models predicted biased *D* estimates (eight overestimations, two underestimations with |MRPE|>30%) with three models having |MRPE|>100%. For most models, the *T_LD_* statistic identified a recent admixture event (~35—53 kya), in agreement with the simulated admixture ages, suggesting that this statistic is accurate for dating admixture in models with no or very limited population structure. In one model, it estimated an interbreeding event happening less than 20 kya, a period when Neanderthals are currently thought to have been extinct for several thousands of years (Ragsdale and Gravel, 2019).

A set of seven models predicted match rates (*m_S’_*) lower than 60%, i.e. significantly less than the observed value of 72.6% or the mean value predicted by our model (66.4%). Five models predicted *CRF* introgression rates at odds with the observed value of 1.17% (|MRPE| > 50%). Finally, all the admixture models produce *PSMC* curves at odds with the empirical curves for CEU and YRI individuals (Li and Durbin, 2011), even when ignoring the most ancient past prior to a million years ago (Figure 3, Notes S8).

We further compared all models using the mean scaled Euclidean distance (MSED) across the twelve comparative statistics, as a scaled measure of discrepancy between simulations and observed data. For each model, we retained the minimum distance for each statistic across twenty simulations accepted using the same standard procedure (see Materials and Methods). For the published models, the only parameter which was let to vary across the 50 simulations was the mutation rate, to improve the model ability to predict diversity statistics. Then, we averaged the distances over the twelve statistics to obtain a single value per model (Notes S1.6). Our structured model had a MSED of 0.44 and ranked first (Table S9). The second-best model was Schaefer et al. (2021) with a distance of 0.68, followed by Moorjani et al. (2016) (0.97). We also compared the fit of our model against the admixture models assuming now a heterogeneous landscape of recombination (Hellenthal and Stephens, 2007). Here, our model and Schaefer’s et al. exhibited near-identical standardized distances (MSED=0.531 and 0.525 respectively). The third model was Moorjani et al. (2016), with a value of 0.853. When considering the average *rank* (instead of the continuous value) of the mean standardized Euclidean distances, our model always ranked first for both the uniform and the heterogeneous recombination rate simulations.

### Ancient DNA admixture estimates

The surprisingly high levels of empirical Neanderthal ancestry estimated in some ancient Eurasian *Hs* specimens—like Oase1 (Fu et al., 2015) or the recently sequenced Bacho Kiro samples (Hajdinjak et al., 2021)—, were suggested as strong evidence of *Hn* admixture into *Hs*. We wondered if our structured model could generate such patterns, too. To this end, We sampled single individuals (to mimic the rarity of aDNA specimens) in the *Hs* Eurasian metapopulation M¢ (from 0 to 45 kya) for the twenty accepted runs of our structured model, assuming uniform recombination. To mimic various types of aDNA genetic data, We considered three SNP sampling strategies: (i) “ *All*”: using all the SNPs simulated for 0.6 Gbp genomes per sample (this would mimic a high-coverage sequencing); (ii)“*1M*”: using a random subset of one million SNPs from the *All* data (to mimic the 1240K SNP array); (iii) “*Archaic*”: a subset of ascertained SNPs from *All* mimicking the Oase1 genotyping (Fu et al., 2015) (cf. Materials and Methods). When measuring the *D* statistic using diploid genotypes and the densest SNP sampling (*All*), we observed that spurious archaic admixture could be identified again (*D*=4.8% on average and *Z*=5.3, IQR95: −0.2—10.7), despite the fact that *Hs* and *Hn* never exchanged gene flow in the simulations. The *D* statistic was stable across the sampling ages (P-value=0.67), in agreement with the best admixture model We co-analyzed (P-value=0.50, Schaefer et al. (2021)). When We accounted for ascertainment biases (see Notes S1.8.4), the simulated *D* values included the empirical *D* estimated on several ancient specimens (Figure 4). Importantly, this included Oase1 when accounting for the *Archaic* ascertainment (Figure 4), a sample often presented as unambiguously admixed. The *D* values for ancient simulated samples and their stability across sampling ages were robust to pseudodiploidization typically used on real data (Skoglund et al., 2012) and SNP downsampling (ANOVA: P-value=0.99), but were on average four times larger using the *Archaic* pseudodiploid data compared to the *1M* diploid data. Our analyses thus suggest that the archaic admixture interpretation for statistics computed on aDNA samples might need to be re-assessed.

**Figure 4.**
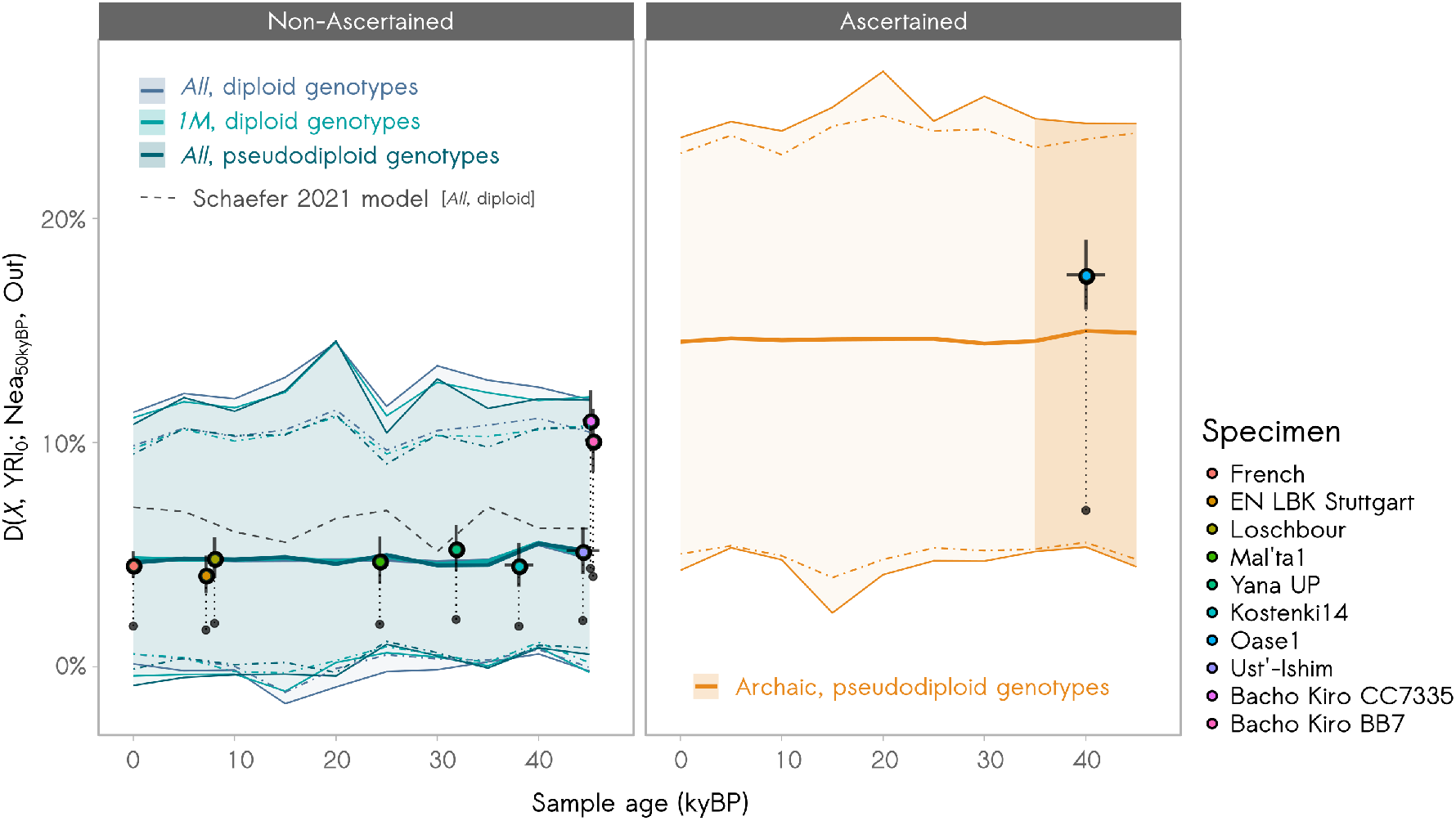
aDNA *D*-statistics. The figure represents the temporal trajectory of the D-statistic estimated on a time series of ancient Eurasian (M_C_ metapopulation) samples—simulated under our structured model together with empirical estimates (dots). The left panel represents the values computed from non-ascertained SNPs. The right panel represents the values computed using a SNP ascertainment mimicking the empirical data for Oase1 (Fu et ah, 2015). To obtain the different values and curves, we simulated single Eurasian samples (*X*) at increasing ages of the past (from present-day up to 45 kya) under the twenty accepted runs of our structured model. We then estimated *D* as *D*(*X*. 50 YRI; 1 Neanderthal, Ancestral reference). The shaded areas (contoured with solid fines) correspond to the interquantile ranges at 100% of the *D* values estimated for various (i) SNP sampling strategies (“*All*Γ: all simulated SNPs; “*1M*”: random subset of one million SNPs; “*Archaic*”: ascertained set of SNPs) and (ii) genotype quality (either diploid or pseudodiploid). The thick lines correspond to the median values across these ranges. The colored dot-dashed lines represent the interquantile ranges at 95%. For comparison, the dotted fine (left panel) is the trajectory of the *D*-statistics estimated under the Schaefer et al. (2021) model. We also represented the (i) *raw* (black points) values of *D* that we re-estimated for ten empirical ancient *Hs* samples from the A ADR v54.1 dataset—SNP array data—, and (ii) their *scaled* (colored points) estimates, obtained by multiplying all the raw *D* values by a constant factor so that the AADR-based raw estimate for the French present-day samples fits the published CEU genome-wide estimate (4.57%). The empirical sample ages are the calibrated ^14^C radiocarbon dates (AADR v54 annotation file). The error bars correspond to the 95% confidence intervals (in the case of the *D*-statistics, this is calculated using a weighted-block jackknife procedure).

Further, We calculated the decay constant of the single-sample ancestry-*LD* decay curve, *T_LD,1_* (Moorjani et al., 2016). We observed that the curves decayed more slowly for some ancient samples compared to present-day simulated samples, in line with empirical observations. For instance, the ~45 kya old Ust’-Ishim fossil exhibits ~1.8—4.2× longer segments (hence slower decay rate) than present-day genomes (Fu et al., 2014). When re-scaling the decay constant *{T_LDΛ_}* in kya after adding the age of the simulated samples, all values agreed across the time series (P-value=0.17 of a linear model) and identified an admixture event that was in most cases <100-200 kya, much more recent than the simulated *Hn-Hs* split times (Notes S7). Our results thus suggest that high levels of *Hn* admixture estimates and longer *Hn* segments can be inferred in aDNA samples under simple models of population structure without admixture.

## Discussion

The sequencing of ancient genomes like those of Neanderthal and Denisova has revolutionarized paleoanthropology. While these technological feats are remarkable, several lines of evidence suggest that caution should still be made regarding the interpretation of genomic and aDNA-based studies. Twenty years ago, when the general consensus was that Neanderthals could not have interbred with *Homo sapiens*, it was noted that the then available genetic data could not be used to rule out interbreeding between *Hn* and *Hs* (Nordborg, 1998; Goldstein and Chikhi, 2002). This statement was based on the work of Nordborg (1998) who had shown by simulations that Neanderthal mtDNA data only allowed to reject panmixia, between *Hs* and *Hn*. But because Nordborg’s conclusions were based on a model assuming a single panmictic population only, the need for considering structured models to study human evolution was already stressed (Goldstein and Chikhi, 2002). The simulation results presented here suggest that similar caution is still recommended today and that studies identifying admixture while ignoring intra-continental population structure should be taken with a grain of salt.

In the last two decades, population geneticists have learned much about (and from) structured models (Herbots, 1994; Hudson, 1990; Notohara, 1990; Wakeley, 1999; Beaumont, 2004; Edmonds et al., 2004; Currat and Excoffier, 2005, 2004; Chikhi et al., 2010, 2018; Mazet et al., 2016; Arredondo et al., 2021; Battey et al., 2020). In particular, it has been shown that when populations are structured, the interpretation of genetic data can be counter-intuitive and can lead to the spurious identification of selection (Currat et al., 2006; Battey et al., 2020) or of specific demographic events like population size changes (Wakeley, 1999; Chikhi et al., 2010; Mazet et al., 2016) and admixture (Eriksson and Manica, 2012, 2014). Worse, spurious signals are more likely to be detected and estimated precisely as genetic information increases (Chikhi et al., 2010). It was thus suggested that as genomic data would become more widely available, researchers might feel more convinced of the existence of events that perhaps never happened, as exemplified with the inference of population size changes in *PSMC* curves based on whole-genome sequences (Chikhi et al., 2018). Recently, several authors have proposed alternatives to panmictic models to interpret the *PSMC* of *Hs* and *Hn* (Mazet et al., 2016; Rodríguez et al., 2018; Arredondo et al., 2021), showing that structured models—with changes in connectivity that include *Hs, Hn* and their ancestor—could explain major patterns of the observed human genomic diversity, including patterns unexplained by other models (see also Scerri et al. (2018, 2019)). In a similar line of research, Eriksson and Manica (2012, 2014) showed that spurious admixture signals—as measured by two statistics (*D* and *DCFS*)—could be generated when intra-continental population structure was ignored. Thus, it should not come as a surprise if admixture parameters provide powerful adjustment variables when geneticists ignore population structure. This may be particularly problematic since admixture event(s) can compensate for the limitations of models that ignore or minimize other forms of connectivity at the intra- and inter-continental level.

In the present study, we confirmed that the *D* statistic and the *DCFS* computed from present-day *Hs* genomes can be reproduced by a demographic model with no admixture between *Hn* and *Hs*. IFc extended the work of Eriksson and Manica (2012, 2014) by showing that a structured model could also explain observed LD-based statistics. We showed that long DNA fragments that are identified as introgressed by LD-based methods like *S’* or *CRF* are expected in our admixture-free model. Thus, one important result and strength of our work is to show that structured models can reproduce a large set of genetic statistics *jointly*, despite the fact that complex relationships may exist between all these statistics (Notes S3). In addition, we showed that our structured model could also predict several statistics measured in aDNA samples. Our results are thus at odds with several reviews stating that admixture statistics are robust to population structure or that *Hn-Hs* admixture is now well established (Peter, 2016; Ahlquist et al., 2021; Theunert and Slatkin, 2017; Vernot and Akey, 2014).

The long Neanderthal tracts in non-African modern humans that are reported in the literature can thus also be generated from shared ancestry in the context of spatial structure. Consequently, these long tracts should not be seen as more “Neanderthal” than “ *sapiens*”, since they can be inherited from an ancestral structured population and maintained in both the *Hs* and *Hn* populations amidst a complex history of local changes in population sizes and connectivity. Thus, a parsimonious explanation could be that *Hn* fixed some ancestral alleles due to their historically smaller population sizes, whereas *Hs* maintained more standing ancestral variation, i.e. a larger proportion of the diversity present in the common ancestor of *Hs* and *Hn*.

We also evaluated how several published models of human evolution—which consider *Hn* admixture into *Hs—*could explain the aforementioned set of statistics. We found that the different models were not necessarily cross-compatible (e.g. admixture dating) and that several of them were not able to predict realistic values of several important measures of human genetic diversity or differentiation. When we compared the discrepancy between model predictions and observed data across twelve statistics, our model provided the highest fit (for nine out of twelve), including when we accounted for variations in recombination rates along the genome.

We note that our objective was not to infer the best scenario using a fully-fledged inferential approach (e.g. Beaumont et al. (2002)). Rather, we aimed at illustrating the principle that structured models represent a promising avenue of research to improve our understanding of human and hominin evolution. More specific structured models might be required in the future to assess the sensitivity and specificity of yet to come admixture statistics. In particular, we have introduced here a simple 1D stepping stone model in which we found that a bipartite ancient structure was required to account for the genomic diversity currently estimated in *Hs*. We interpret this surprising result as a hint that ancient population structure should be explicitly integrated in future models of hominin evolution. The parsimonious scenario mentioned above, which we find is currently the best in our comparative model set, does not necessarily mean that no admixture ever took place between *Hn* and *Hs*. What our results suggest is that, if admixture ever occurred, it is currently hard to identify using existing methods, due to their limited robustness to population structure.

Future work should explore these questions within more complex spatial models, or metapopulations integrating population extinctions (Scerri et al., 2018, 2019; Chikhi, 2023). As noted by Scerri et al. (2019), choosing meta-population over tree models requires to shift from the estimation of split times, for instance, towards the quantification of processes such as changes in connectivity, extinction or recolonization rates. Focusing on split times tends to stress differences between human groups, whereas metapopulation models might allow us to focus on the dynamics of ancient populations and may thus provide a less problematic approach to evolutionary anthropology (Chikhi, 2023).

The issues identified here should thus not be construed to the identification of a simple technical issue, since they question how narratives are constructed in our field and how consensus is built. While admixture between *Hn* and *Hs* was considered impossible by most researchers twenty years ago, it is now difficult to even question that it took place. The story that defines humans has changed, to the point that Neanderthals are now regularly and increasingly identified as being responsible for many adaptations and diseases observed in some present-day *Hs* populations.

We conclude by stressing that this work does not oppose the idea of ancient interbreeding between *Hs* and *Hn*. After all, they both are hominin species with long generation times who probably diverged relatively recently and likely had overlapping distributions for thousands of years (Talamo et al., 2023). This likely provided opportunity for them to interact and possibly interbreed. Yet, our work does suggest that current statistical approaches to identify these putative events remain subject to confounding factors, and notably, to population structure. Our results thus question the power of genetic data and current admixture statistics to provide clear and unambiguous evidence in favor of ancient interbreeding. Although admixture models provide interesting avenues for research, we wonder whether Neanderthal admixture estimates might not decrease (and admixture possibly vanish) when these models better integrated intra-continent population structure. This is however an open question given the infinitely large parameter space of structured models. We thus argue that admixture between *Hs* and *Hn*, and possibly other hominins, needs to be re-evaluated in the light of population structure. Ultimately, the questions asked and the modeling framework developed in this study are important beyond humans and hominins. Population structure may have affected the inference of hybridization in many living taxa, suggesting that some hybridization or introgression results might also benefit from a statistical re-evaluation under structured models.

## Materials and Methods

We provide here a brief outline of our approach but a detailed description can be found in the Supplementary Notes S1.

### The structured model

We assume a set of linear stepping stones—represented in Figure 1 (simplified version) and Figure S1 (detailed)—to which we will refer as “metapopulations” throughout the manuscript, because our model differs from classical stationary stepping stones which do not account for change in the number of demes. We consider that new demes are created as new regions are colonized, starting with ten demes in the most distant past (metapopulation M_A_) towards a full extent of 30 *Hs* demes (10 in metapopulation M_A_, 10 in M_B_, 10 in Mρ) and ten *Hn* demes in metapopulation M_N_ Details of the timings and demographic processes are provided in Notes S1.1. In short, the first metapopulation, M_A_, is comprised of ten demes where each deme is connected to its neighboring demes or deme. A second metapopulation, M_B_, is founded sequentially, one generation at a time, from the tenth M_A_ deme. Gene flow occurs between the tenth M_A_ and the first M_B_ deme, as well as between neighbouring demes within the M_B_ metapopulation. A new linear metapopulation, M_N_, is later founded from the tenth M_B_ deme, following the same process of stepping stone creation but with different deme sizes. M_N_ demes exchange genes among neighbours but do not exchange genes with M_A_ or M_B_ demes after the foundation of the first M_N_ deme. M_N_ will evolve into *Hn*, whereas M_A_ and M_B_ will evolve into *Hs*. As M_N_ is created, the deme sizes in M_A_ and M_B_ are allowed to change to a new value. Also, after the M_N_ foundation, gene flow between M_A_ and M_B_ is allowed to increase. At another later time period, a new metapopulation, M_C_, is founded from the tenth M_B_ deme to represent the colonization of Eurasia by *Hs*. Each creation of M_C_ demes starts with a multigenerational bottleneck. Gene flow occurs between the tenth M_B_ and the first M_C_ deme and between M_C_ neighboring demes. No gene flow is ever allowed between M_N_ and M_C_ (or M_A_ and M_B_) demes. When M_C_ is founded, M_A_ is allowed to have different deme sizes from M_B_ and gene flow is allowed to change between M_B_ demes. Altogether, this model accounts for three foundations of metapopulations, and for changes in connectivity and deme sizes. However, it does not allow gene flow between M_N_ with any of the other metapopulations, i.e. no gene flow is ever allowed between *Hn* and *Hs*.

The timing of these events, the deme sizes and migration rates between demes or metapopulations were allowed to vary within wide pre-defined ranges, consistent with the literature (Notes S1.1, Table S2). For instance, the split between M_N_ and M_B_ was allowed to occur between 400 kya and 1 Mya, to account for the dating uncertainty of the divergence between *Hs* and *Hn*. Details on the other parameters, including the 25 free parameters, can be found in Notes Sl.l. We note that We included three parameters which enabled the sampling of individuals in random demes within their respective metapopulation. This allowed introducing variability across runs, to avoid arbitrarily sampling individuals within specific demes of our structured model. We thus sampled 50 individuals at present in M_A_ (corresponding to present-day YRI) and 50 in M_C_ (present-day CEU). We also sampled one Neanderthal at 50 kya in M_N_ (Vindija33.19) (Prüfer et al., 2017).

A large number of simulations was produced using the coalescent simulator `msprime 1.1.1` (Hudson’s algorithm with recombination) (Kelleher et al., 2016; Baumdicker et al., 2022) and a two-step filtering strategy to minimize computation burden. For the initial exploration of the parameter space, each individual genome consisted of seven chromosomes of length 10 Mbp each. We screened the simulated data after simulating the first chromosome and excluded the runs where some statistics were clearly out of range compared to real data (see Notes S1 for details). We repeated this screening process after simulating the second chromosome. We completed the full (10 × 7 Mbp genetic data) simulations for the runs which successfully passed the two-step filtering process. Thus, from testing one million parameter combinations, we generated 10,361 genetic data sets from which the full set of statistics was computed (see below).

For these simulations, We considered a uniform recombination landscape with a rate of 10^-8^ events per base pair per lineage per generation (Halldorsson et al., 2019) and a binary mutation model with a rate equal to 1.2×10^-8^ mutations per base pair per lineage per generation (Jónsson et al., 2017). We did not account for gene conversion. All analyzes assumed a generation time of 25 years (Li and Durbin, 2011).

### Summary statistics

The simulated genetic data were summarized using a collection of statistics detailed in Notes S1.3, computed using original softwares, custom scripts and Python libraries (`scikit-allel 1.3.2`). Briefly, these include (i) within-population diversity statistics, in CEU and YRI populations: nucleotide diversity *π*, normalized derived allele frequency spectrum (*AFS*), *PSMC* (Li and Durbin, 2011); (ii) bctwccn-population differentiation statistics, between CEU and YRI: Hudson’s *F_ST_*, (iii) “admixture” statistics for Neanderthal signals into CEU compared to YRI: *D*-statistics (Green et al., 2010), doubly conditioned frequency spectrum (*DCFS*) (Yang et al., 2012), mean length (*L_S’_*) and mean match rate (*m_S’_*) of the putative Neanderthal-introgressed segments inferred by *S’* (Browning et al., 2018), introgression rate (*α_CRF_*) and mean length (*L_CRF_*) of the putative Neanderthal-introgressed segments inferred by a conditional random field (*CRF*) (Sankararaman et al., 2014), decay constant of the multisample ancestry-*LD* (*T_LD_*) (Sankararaman et al., 2012).

The corresponding empirical estimates for these summary statistics were extracted from published literature for sampling designs that matched our simulation framework. See Notes S2 for a detailed description of how these empirical estimates were obtained. Note that for some summary statistics (related to *S’, CRF* and ancestry-*LD*), we applied a filter to our simulated genetic data (minor allele count filtering), in order to mimic the type of empirical data analyzed in the original studies (see Notes S1.3.2).

### Run selection

Among the 10,361 filtered runs, we retained a set of twenty simulations which were inferred to be the closest to the empirical data according to ten summary statistics (Notes S1.4). We estimated the global distance of each simulation to the observed data by calculating *D*\* defined as:

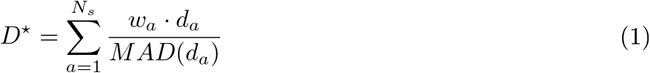

where *N_s_* is the total number of statistics considered (*N_s_* = 10), *d_a_* is the Euclidean distance between the observed and simulated values of the (vectorial or scalar) statistics *S_a_, w_a_* is the weight of the statistics *S_a_* and *MAD*(·) is the mean absolute deviation. Note that for *PSMC, d_a_* is defined as the cross-correlation distance (Notes S1.4.1).

Based on the twenty retained runs, we then simulated genetic data with larger genome sizes (600 Mbp) composed of twenty chromosomes of length 30 Mbp each. All subsequent analyses reported in this study are based on these larger datasets.

### Robustness analysis

We assessed the robustness of our analyses to variable sources of genomic uncertainty (low coverage data leading to SNP sampling biases, ancestral allele misspecification, pseudodiploid data), to the sampling ages of Neanderthal and to the nature of the recombination maps (uniform, heterogeneous vs. empirical). Details are provided in Notes S10.

### Comparison with other published models

We assessed the performance of our admixture-free structured model in comparison with a collection of eleven published models of human evolution which assume (i) admixture from at least one archaic species (always Neanderthals) and (ii) no or limited intra-continental population structure (Yang et al., 2012; Moorjani et al., 2016; Skov et al., 2020; Ragsdale and Gravel, 2019; Durvasula and Sankararaman, 2020; Gower et al., 2021; Iasi et al., 2021; Schaefer et al., 2021). A few models (Jacobs et al., 2019; Kamm et al., 2020) allowed for several *Hs* populations within a continent, but these were typically defined by a unique “ghost” population or by aDNA specimens, and corresponded to branches in an evolution tree where gene flow was often absent, or only represented by admixture events. Yet, given the great diversity of these published demographic models, a detailed description of each model (with visual representation) is provided in Notes S8 along with the simulation protocols and commands. For each published model (characterized by its parametric point estimates), we simulated 50 datasets by varying the mutation rates uniformly from 5×10^-9^ to 5×10^-8^ bp^-1^g^-1^ and selected the twenty runs closest to the observed data, following the same procedure as above. We then compared the performance of the twelve models by calculating, for each model *M_i_*, the minimum scaled Euclidean distance (MSED) *D*_M_i__*:

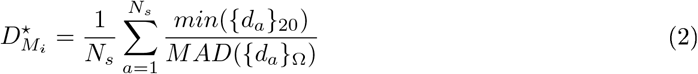

where *min*({*d_a_*}_20_) is the minimum of the Euclidean distances calculated for the (scalar or vectorial) statistics *S_a_* across the twenty accepted runs of the model *M_i_* and *MAD*({*d_a_}_Ω_*) is the mean absolute deviation of the distances *d_a_* across the twenty accepted runs of *all* compared models.

### Ancient DNA study

We investigated the values of two summary statistics (*D*-statistics (Green et al., 2010) and the single-sample ancestry-*LD* decay constant *T_LD-1_* (Moorjani et al., 2016)) in ancient samples that We simulated (i) under the twenty accepted runs of our structured model and (ii) Schaefer et al.’s (2021) model. Details on the simulations and analyses are provided in Notes S1.8.

## Code availability

All the scripts used in this study (including but not limited to: model simulation and plotting, statistics calculation, run selection, model comparison, figure plotting) as well as the demes YAML-formatted demographic histories of the twenty selected runs will be publicly available in a GitHub repository (freely available once the manuscript is accepted, can also be provided to the editor and referees if requested or required).

## Acknowledgements

We thank Olivier Mazet, Simon Boitard, Barbara Parreira, Armando Arredondo and members of the Population and Conservation Genetic group for their support and for useful discussions on this topic. We would like also to acknowledge the Bioinformatic Unit and the Informatics Team of the IGC, as well as CALMIP (project P23002) for their help and support with computational resources. We thank Olivier Mazet and Simon Boitard for their useful comments on the first version of this manuscript. LC and RT were funded by Fundação para a Ciência e Tecnologia (ref. PTDC-BIA-EVL/30815/2017). This work was also supported by the LABEX entitled TULIP (ANR-10-LABX-41 and ANR-11-IDEX-0002-02) as well as the IRP BEEG-B (International Research Project - Bioinformatics, Ecology, Evolution, Genomics and Behaviour). We acknowledge an Investissement d’Avenir grant of the Agence Nationale de la Recherche (CEBA: ANR-10-LABX-25-01).

