## Supplementary Materials for "Questioning Neanderthal admixture: on models, robustness and consensus in human evolution"

### Supporting Information for:
### Questioning Neanderthal admixture: on models, robustness and consensus in human evolution

Rémi Tournebize Lounès Chikhi

April 5, 2023

#### Summary

**S1 Detailed Material and Methods** 4

**S2 Observed statistics in present-day modern humans** 21

**S3 Collinearity between statistics** 26

**S4 Model performance with heterogeneous recombination** 27

**S5 Parameters of the 20 accepted runs from the structured model** 30

**S6 $F_{ST}$ trajectory between metapopulations** 31

**S7 Single-sample ancestry-*LD* decay constants for aDNA** **33**

**S8 Published models** **34**

### S1 Detailed Material and Methods

**General notes** If not stated otherwise, we consider by default that individuals (or alternatively: samples or specimens) as well as effective population sizes are **diploid**. Throughout the manuscript, we tried to avoid the term "archaic" which is neither clear nor very helpful (Scerri et al., 2019). However, in order to match original designations, we used it on some occasions where results or methods (usually from published studies) explicitly referred to "archaic" species, populations or admixture.

#### S1.1 Detailed description of the structured model

Our structured model starts with a chain (1D stepping-stone metapopulation) of ten equally sized demes (size $N_5$) connected by symmetric gene flow ($m_1$). This chain will be referred to as metapopulation $\mathrm{M_A}$. See Figure S1 for a visual representation of the model (produced with the Python library `demesdraw`). See Table S1.1 and S2 for details about the parameters.

At a time $T_4$, the rightmost deme of the metapopulation $\mathrm{M_A}$ founds a new chain, $\mathrm{M_B}$, following a sequence of stepping-stone expansion events happening every generation. This new metapopulation $\mathrm{M_B}$ reaches a full size of ten equally-sized demes (same size as $\mathrm{M_A}$: $N_5$). Within $\mathrm{M_B}$, demes are connected by asymmetric gene flow (with rates $m_7$ and $m_8$). The two demes connecting $\mathrm{M_A}$ to $\mathrm{M_B}$ also have asymmetric gene flow (with rates $m_9$ and $m_{10}$). The two connected metapopulations model an hominid species ancestral to both *Hs* and *Hn*.

At a time $T_3$ between 400 kya and 1 Mya (after $T_4$), the rightmost deme of the metapopulation $\mathrm{M_B}$ founds a new chain ($\mathrm{M_N}$) of ten equally-sized demes (size $N_4$), following a similar sequence of stepping-stone expansion events happening every generation. Demes within $\mathrm{M_N}$ are connected by symmetric gene flow (same rate as in $\mathrm{M_A}$: $m_1$). This metapopulation $\mathrm{M_N}$ will ultimately evolve into **Neanderthals**. By contrast, the metapopulations $\mathrm{M_A}$ and $\mathrm{M_B}$ will evolve into ***Hs***. When $\mathrm{M_N}$ is founded, we allow the size of the demes within $\mathrm{M_A}$ and $\mathrm{M_B}$ to change (new size: $N_2$). We note that the metapopulation $\mathrm{M_N}$ never sends nor receives any migrant to nor from any other metapopulation. It is therefore totally isolated since its foundation. This is in clear contrast with the majority of human demographic models which assume some gene flow between humans and Neanderthals thousands of years after their divergence.

At a time $T_2$ between 40 kya and 1 Mya (note: $T_2$ chronologically follows $T_3$), we allow gene flow between the metapopulations $\mathrm{M_A}$ and $\mathrm{M_B}$ to increase, respectively: from $m_9$ to $m_4$ and from $m_{10}$ to $m_5$.

At time $T_1$ (between 40 kya and 70 kya, always after $T_2$), a third *Hs* metapopulation ($\mathrm{M_C}$) is founded by the stepping-stone expansion of the last deme of $\mathrm{M_B}$. This period corresponds to the *Hs* expansion into Eurasia. At its full extent, $\mathrm{M_C}$ is comprised of ten equally-sized demes (size $N_3$), connected by asymmetric gene flow (with rates $m_2$ and $m_3$). Gene flow between the $\mathrm{M_B}$ and the $\mathrm{M_C}$ chains is symmetric (rate $m_6$). Each founder event is modelled every $dT1$ years as a multi-generation bottleneck, with founder size $n_3$ and duration $dT_2$. At the same time as $\mathrm{M_C}$ is formed, we allow (i) the symmetric migration within $\mathrm{M_B}$ to change to the same rate as within $\mathrm{M_A}$ (i.e. $m_1$); (ii) the sizes of the demes within $\mathrm{M_A}$ to change to $N_1$.

Also, we include three sampling parameters $p_i$ corresponding to the relative positions (from 0 to 1) of the demes in which we sample the YRI (within $\mathrm{M_A}$), the CEU (within $\mathrm{M_C}$) and the Vindija Neanderthal (within $\mathrm{M_N}$). To obtain the index $i$ (0-based indexing) of the deme in which samples are to be drawn, we compute $i = \lfloor p(\#M_x - 1)\rceil$ where $\lfloor \cdot \rceil$ is the rounding function, $\#M_x$ is the number of demes within metapopulation $M_x$ where sampling should be done, and $p$ is the relative sampling position. This formulation leads to downsample (by 2-fold in the prior distribution) the demes at the two extreme ends of the chains, which is an intended feature given the low probability for the analyzed populations to originate from any extreme of the metapopulations.

Note that some asymmetries (e.g. in gene flow) implemented in this model naturally arose during preliminary exploration of the parameter space. However these patterns should not be overinterpreted because we consider this model as a first step towards building a general inferential framework for structured models.

Note that all parameters described above assume a forward-time perspective (including migration rates). They are converted to coalescent parameters in the backward perspective for the simulations under `msprime`. The parametric input files to our demographic simulations (provided in the GitHub repository) are the coalescent backward-time parameters.

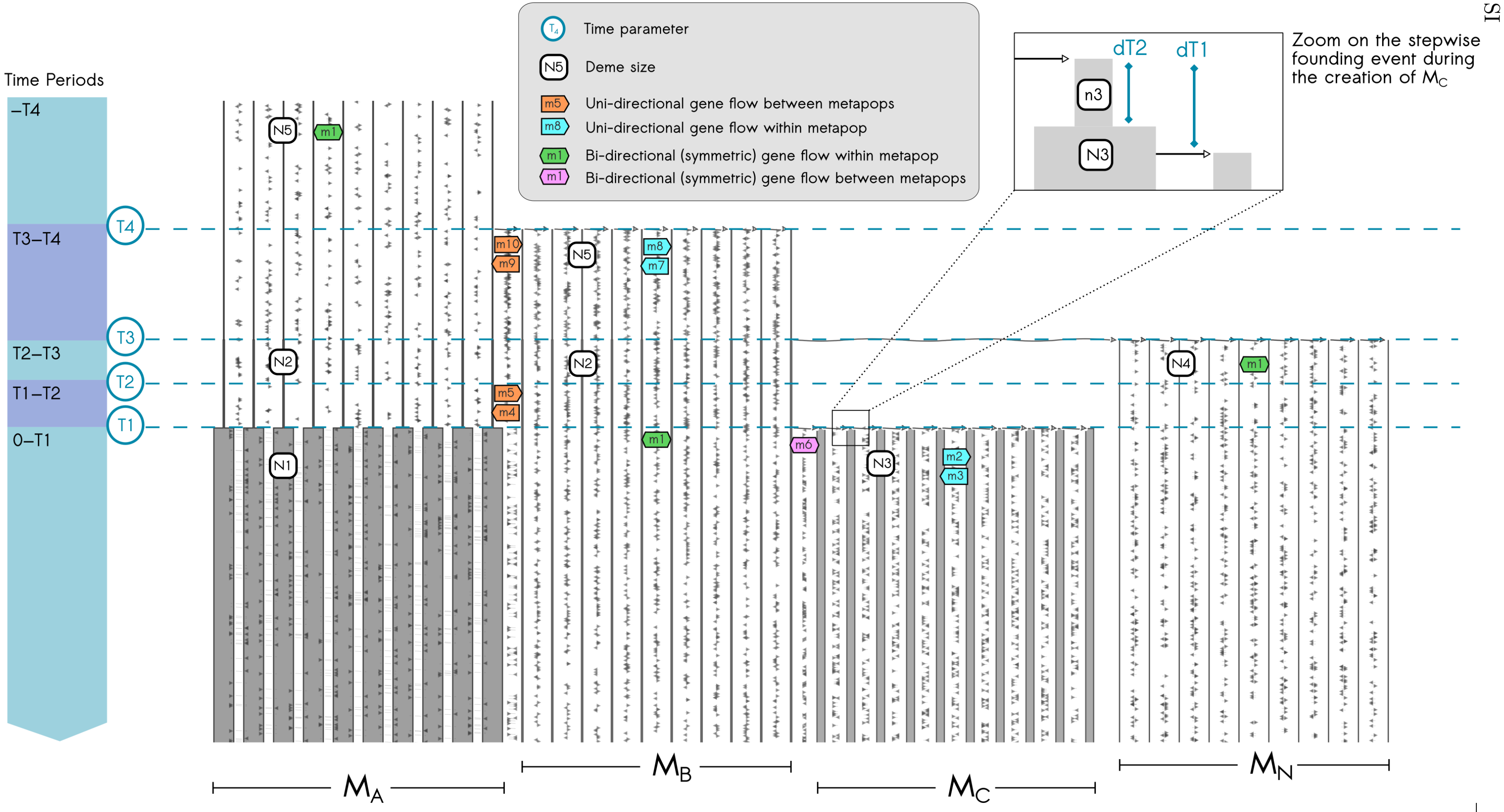

Figure S1: Visual representation of our structured model. Note that time flows from top to bottom. The migration arrows are in the forward-time perspective. The model representation was produced using the Python library `demesdraw`.

| Period | Parameter type | Codename+ | Parameter | Description |
|---|---|---|---|---|
| —T4 | Deme size | **n.rift.AfW** | **N5** | Effective size of each deme in $M_A$ |
| | Within-metapop gene flow | **m.rift.AfW>** | **m1** | Intrinsic migration rate from one deme to the deme on its right, within $M_A$ |
| | Within-metapop gene flow | **m.rift.AfW<** | **m1** | Instrinsic migration rate from one deme to the deme on its left, within $M_A$ |
| T3—T4 | Time | **T.rift** | **T4** | Time point |
| | Deme size | n.nea.AfW | N5 | Effective size of each deme in $M_A$ |
| | Deme size | **n.nea.AfE** | **N5** | Effective size of each deme in $M_B$ |
| | Within-metapop gene flow | m.nea.AfW> | m1 | Intrinsic migration rate from one deme to the deme on its right, within $M_A$ |
| | Within-metapop gene flow | **m.nea.AfE>** | **m8** | Instrinsic migration rate from one deme to the deme on its right, within $M_B$ |
| | Within-metapop gene flow | m.nea.AfW< | m1 | Instrinsic migration rate from one deme to the deme on its left, within $M_A$ |
| | Within-metapop gene flow | **m.nea.AfE<** | **m7** | Instrinsic migration rate from one deme to the deme on its left, within $M_B$ |
| | Between-metapopulation gene flow | **m.nea.AfW_from_AfE** | **m9** | Intrinsic migration rate from the first deme of $M_B$ into the last deme of $M_A$ |
| | Between-metapopulation gene flow | **m.nea.AfE_from_AfW** | **m10** | Intrinsic migration rate from the last deme of $M_A$ into the first deme of $M_B$ |
| T2—T3 | Time | **T.nea** | **T3** | Time point |
| | Deme size | **n.mis5.AfW** | **N2** | Effective size of each deme in $M_A$ |
| | Deme size | **n.mis5.AfE** | **N2** | Effective size of each deme in $M_B$ |
| | Deme size | **n.mis5.Nea** | **N4** | Effective size of each deme in $M_N$ |
| | Within-metapop gene flow | m.mis5.AfW> | m1 | Intrinsic migration rate from one deme to the deme on its right, within $M_A$ |
| | Within-metapop gene flow | m.mis5.AfE> | m8 | Instrinsic migration rate from one deme to the deme on its right, within $M_B$ |
| | Within-metapop gene flow | **m.mis5.Nea>** | **m1** | Instrinsic migration rate from one deme to the deme on its right, within $M_N$ |
| | Within-metapop gene flow | m.mis5.AfW< | m1 | Instrinsic migration rate from one deme to the deme on its left, within $M_A$ |
| | Within-metapop gene flow | m.mis5.AfE< | m7 | Instrinsic migration rate from one deme to the deme on its left, within $M_B$ |
| | Within-metapop gene flow | **m.mis5.Nea<** | **m1** | Instrinsic migration rate from one deme to the deme on its left, within $M_N$ |
| | Between-metapopulation gene flow | m.mis5.AfW_from_AfE | m9 | Intrinsic migration rate from the first deme of $M_B$ into the last deme of $M_A$ |
| | Between-metapopulation gene flow | m.mis5.AfE_from_AfW | m10 | Intrinsic migration rate from the last deme of $M_A$ into the first deme of $M_B$ |
| T1—T2 | Time | **T.mis5** | **T2** | Time point |
| | Deme size | n.ooa.AfW | N2 | Effective size of each deme in $M_A$ |
| | Deme size | n.ooa.AfE | N2 | Effective size of each deme in $M_B$ |
| | Deme size | n.ooa.Nea | N4 | Effective size of each deme in $M_N$ |
| | Within-metapop gene flow | m.ooa.AfW> | m1 | Intrinsic migration rate from one deme to the deme on its right, within $M_A$ |
| | Within-metapop gene flow | m.ooa.AfE> | m8 | Instrinsic migration rate from one deme to the deme on its right, within $M_B$ |
| | Within-metapop gene flow | m.ooa.Nea> | m1 | Instrinsic migration rate from one deme to the deme on its right, within $M_N$ |
| | Within-metapop gene flow | m.ooa.AfW< | m1 | Instrinsic migration rate from one deme to the deme on its left, within $M_A$ |
| | Within-metapop gene flow | m.ooa.AfE< | m7 | Instrinsic migration rate from one deme to the deme on its left, within $M_B$ |
| | Within-metapop gene flow | m.ooa.Nea< | m1 | Instrinsic migration rate from one deme to the deme on its left, within $M_N$ |
| | Between-metapopulation gene flow | **m.ooa.AfW_from_AfE** | **m4** | Intrinsic migration rate from the first deme of $M_B$ into the last deme of $M_A$ |
| | Between-metapopulation gene flow | **m.ooa.AfE_from_AfW** | **m5** | Intrinsic migration rate from the last deme of $M_A$ into the first deme of $M_B$ |
| 0—T1 | Time | **T.ooa** | **T1** | Time point |
| | Duration | **j.ooa.EuA_into_AfE** | **dT1** | Duration between successive founding events of $M_C$ demes |
| | Duration | **jd.ooa.EuA_into_AfE** | **dT2** | Duration of the bottleneck for the founding of each $M_C$ deme |
| | Deme size | **jn.ooa.EuA_into_AfE** | **n3** | Effective size of each deme $M_C$ when it is founded |
| | Deme size | **n.0.AfW** | **N1** | Effective size of each deme in $M_A$ |
| | Deme size | n.0.AfE | N2 | Effective size of each deme in $M_B$ |
| | Deme size | **n.0.EuA** | **N3** | Effective size of each deme in $M_C$ (after instantaneous expansion, following the founding bottleneck) |
| | Deme size | n.0.Nea | N4 | Effective size of each deme in $M_N$ |
| | Within-metapop gene flow | m.0.AfW> | m1 | Intrinsic migration rate from one deme to the deme on its right, within $M_A$ |
| | Within-metapop gene flow | **m.0.AfE>** | **m1** | Instrinsic migration rate from one deme to the deme on its right, within $M_B$ |
| | Within-metapop gene flow | **m.0.EuA>** | **m2** | Instrinsic migration rate from one deme to the deme on its right, within $M_C$ |
| | Within-metapop gene flow | m.0.Nea> | m1 | Instrinsic migration rate from one deme to the deme on its right, within $M_N$ |
| | Within-metapop gene flow | m.0.AfW< | m1 | Instrinsic migration rate from one deme to the deme on its left, within $M_A$ |
| | Within-metapop gene flow | **m.0.AfE<** | **m1** | Instrinsic migration rate from one deme to the deme on its left, within $M_B$ |
| | Within-metapop gene flow | **m.0.EuA<** | **m3** | Instrinsic migration rate from one deme to the deme on its left, within $M_C$ |
| | Within-metapop gene flow | m.0.Nea< | m1 | Instrinsic migration rate from one deme to the deme on its left, within $M_N$ |
| | Between-metapopulation gene flow | m.0.AfW_from_AfE | m4 | Intrinsic migration rate from the first deme of $M_B$ into the last deme of $M_A$ |
| | Between-metapopulation gene flow | m.0.AfE_from_AfW | m5 | Intrinsic migration rate from the last deme of $M_A$ into the first deme of $M_B$ |
| | Between-metapopulation gene flow | **m.0.AfE_from_EuA** | **m6** | Intrinsic migration rate from the first deme of $M_C$ into the last deme of $M_B$ |
| | Between-metapopulation gene flow | **m.0.EuA_from_AfE** | **m6** | Intrinsic migration rate from the last deme of $M_B$ into the first deme of $M_C$ |

Table S1: Description of the parameters of our structured models. Note that the descriptions here (including migration rates) are given in a forward-time perspective. They are converted to backward-time for the coalescent simulations (see GitHub repository). The parameters in **bold** are the ones which are new or whose values change compared to a previous time period (in the forward direction). For prior distributions, see Table S2.

Notes. + Codenames are the parameter names in the input parameter file of our simulations, see our Github repository.

Table S2: Prior distribution of the 25 free parameters used in our structured model. For details about these parameters, see Table S1.1.
Rules. Times parameters (in kya) must check: T4 > T3 > T2 > T1.
Units. Deme sizes (= parameter names starting with **N**) are diploid. Time parameters (= parameter names containing the character **T**) are in years before present, later converted to generations BP assuming $g = 25$ years. Migration rates (= parameter names starting with **m**) are the probabilities that any lineage in a deme migrate to another deme per generation.

| **Parameter** | **Distribution** | **Lower bound** | **Upper bound** |
|---|---|---|---|
| N1 | $\log_{10}$-uniform | 1,000 | 60,000 |
| N2 | $\log_{10}$-uniform | 1,000 | 10,000 |
| N4 | $\log_{10}$-uniform | 10 | 2,500 |
| N3 | $\log_{10}$-uniform | 2,000 | 60,000 |
| m1 | $\log_{10}$-uniform | $10^{-3}$ | 0.1 |
| m2 | $\log_{10}$-uniform | $10^{-6}$ | 0.1 |
| m3 | $\log_{10}$-uniform | $10^{-6}$ | 0.1 |
| m4 | $\log_{10}$-uniform | $10^{-8}$ | $10^{-3}$ |
| m5 | $\log_{10}$-uniform | $10^{-3}$ | 0.5 |
| m6 | $\log_{10}$-uniform | $10^{-7}$ | $10^{-4}$ |
| T1 | uniform | 40,000 | 70,000 |
| dT1 | $\log_{10}$-uniform | 25 | 5,000 |
| n3 | $\log_{10}$-uniform | 2 | 5,000 |
| dT2 | $\log_{10}$-uniform | 1 | 2,000 |
| m7 | $\log_{10}$-uniform | $10^{-5}$ | 0.1 |
| m8 | $\log_{10}$-uniform | $10^{-3}$ | 0.1 |
| T2 | uniform | 40,000 | 1,000,000 |
| m9 | uniform | $10^{-5}$ | $3{\times}10^{-5}$ |
| m10 | uniform | $3.5{\times}10^{-7}$ | $8{\times}10^{-7}$ |
| T3 | uniform | 400,000 | 1,000,000 |
| N5 | uniform | 500 | 1,500 |
| T4 | uniform | 500,000 | 9,000,000 |
| *Relative position of the sampled demes:* | | | |
| $p_{YRI}$ | uniform | 0 | 1 |
| $p_{CEU}$ | uniform | 0 | 1 |
| $p_{Vindija}$ | uniform | 0 | 1 |

#### S1.2 Simulations

We simulated one million genomic datasets under a custom 1D stepping-stone model without Neanderthal admixture in *Hs*. We used the coalescent simulator `msprime` 1.1.1 with Hudson's standard time-continuous algorithm with recombination (Kelleher et al., 2016; Baumdicker et al., 2022). For each simulation, we randomly drew values for the 25 free parameters considering predefined search ranges that are detailed in Table S2 For each dataset, we sampled in random demes of each respective metapopulation:

- 50 diploid individuals sampled at present in the African metapopulation $M_A$: corresponding to **YRI**.
- 50 diploid individuals sampled at present in the Eurasian metapopulation $M_C$: **CEU**.
- one Neanderthal diploid individual sampled 2,000 generations BP (50 kya assuming a generation time of $g = 25$ years) in $M_N$: **Vindija33.19 Neanderthal** (Prüfer et al., 2017).

Note that the index of the deme sampled in each metapopulation is a free parameter of the model, cf. Notes S1.1 for how this random sampling is performed.

For the initial exploration of the parameter space, each individual genome consisted of 7 chromosomes of length 10 Mbp each. We first considered a uniform recombination landscape with a rate of $10^{-8}$ events per base pair per lineage per generation (Halldorsson et al., 2019). We used a binary mutation model (`BinaryMutationModel` in `msprime`), i.e. two alleles only with between-allele transitions of 1.0 and 1.0, where a new mutation at an already mutated site generates a state which depends on the previous one (`state_independent=False`). The mutation rate was fixed at $1.2\times10^{-8}$ mutations per base pair per lineage per generation following (Jónsson et al., 2017) (estimate from parent-offspring data). Note that we also checked that our results were consistent with alternative mutation models (Jukes-Cantor 1969). Also, we considered no gene conversion. All analyzes assumed a generation time of $g = 25$ years (Li and Durbin, 2011). Since `msprime` simulates haploid chromosomes, we randomly drew two haploid chromosomes without replacement to generate diploid data.

To limit computation burden, we screened the datasets after simulating the first two chromosomes in order to avoid simulating the rest of the genome if some genetic summary statistics showed preliminary evidence of unrealistic values. Specifically, after simulating the first chromosome, we calculated the genetic diversity in both CEU and YRI and stopped the simulation if $\pi_{\mathrm{CEU}}$ was outside of the range 0.5–1.5 kb$^{-1}$ (Table S6) and $\pi_{\mathrm{YRI}}$ was outside of the range 0.5–1.8 kb$^{-1}$. We then checked the values of eight statistics after simulating the second chromosome. Specifically, we assessed if: $\pi_{\mathrm{CEU}}$ was within the range 0.5–1.5 kb$^{-1}$; $\pi_{\mathrm{YRI}}$ within 0.5–1.8 kb$^{-1}$; $F_{ST}$ between CEU and YRI within 0.05–0.35; the frequency of singletons in CEU within 0.18–0.35; the frequency of singletons in YRI within 0.28–0.42; the frequency of singletons of the *DCFS* in CEU within 0.2–0.4; the mean length of the *S'*-identified Neanderthal segments in CEU was within 100kb—600kb and the $m_{S'}$ match rate of the *S'* segments were within 0.65–0.90 (Table S6). If any statistics lay outside of the specified range, the simulation was stopped and the dataset deleted. This procedure led to simulate full genetic data ($7 \times 10$ Mbp) for 10,361 runs (i.e. 0.1% of the 1M parameter combinations), with a significant speed-up of the parameter exploration process. Note that we further excluded duplicate genetic datasets, i.e. those which happened to have identical parameter combinations due to the random sampling within the prior distributions.

In all our simulation studies, we assumed ideal genotypes with no missing data. Also, if not stated otherwise (e.g. for aDNA studies, Notes S1.8), all the genotypes were considered diploid (not pseudodiploid).

**Simulation of heterogeneous recombination landscape.** For situations which required to model a more realistic recombination landscape with varying local recombination rates, we followed an approach described in Hellenthal and Stephens (2007) which accounts for recombination hotspots. Specifically, for each chromosome, we note $H$ the total number of points of recombination rate changes. $H$ was randomly sampled in a Poisson distribution with $\lambda = \frac{L}{W}$ where $L$ is the size of the chromosome (in bp) and $W$ the expected spacing (in bp) between two successive rate change points ($W$=40 kbp, Hellenthal and Stephens (2007)). Each local recombination region $h$ ($h$=*1,...,H*) was characterized by (i) its left boundary (sampled in a uniform distribution from 1 to $L$ bp); (ii) its length (sampled in a uniform distribution from 1,000 to 2,000 bp) and (iii) its local recombination rate, defined as $\rho_{bg} \cdot \eta_h$with $\rho_{\mathrm{bg}}$ the background recombination rate (in bp$^{-1}$g$^{-1}$) and $\eta_{\mathrm{h}}$ the intensity of the rate change, sampled in a $\log_{10}$-uniform distribution from 10 to 316 (Hellenthal and Stephens, 2007). Note that if the length between two rate change points fell below 5 kbp, the second rate change was not implemented, as suggested by Hellenthal and Stephens (2007). Outside of the $H$ regions, the recombination rate $\rho_{\mathrm{bg}}$ was set as $\rho_{\mathrm{bg}}=2.325\times10^{-9}$ bp$^{-1}$g$^{-1}$ resulting (in composition with the other parameters) in a standard genome-wide average recombination rate of $1.0\times10^{-8}$ bp$^{-1}$g$^{-1}$.

#### S1.3 Computation of summary statistics

##### S1.3.1 Non-admixture statistics

Simulated genetic data were summarized using a collection of genetic statistics informative about within-population diversity and between-population differentiation.

**Nucleotide diversity ($\pi$)** We estimated the genetic diversity $\pi$ on the 50 CEU and the 50 YRI diploids separately as the mean number of pairwise differences per kilobase between individuals from each population, using `scikit-allel 1.3.5` (Miles et al., 2021).

***AFS*** The derived allele frequency spectra (*AFS*) in present-day CEU and YRI were calculated by considering a subset of 10 random individuals in each population, to match the published *AFS* procedure (Gutenkunst et al., 2009). After excluding the monomorphic entries, each *AFS* was scaled by the sum of the remaining entries (from DAC=1–9), so that each value of the spectrum represents the probability of observing a SNP at a particular derived allele frequency in the target population.

**$F_{ST}$** The $F_{ST}$ between the 50 CEU and 50 YRI diploids was calculated with `scikit-allel 1.3.5` using Hudson's estimator (Hudson et al., 1992):

$$\hat{F}_{ST} = 1 - \frac{\bar{H}_w}{H_b} \tag{S1}$$

where $\bar{H}_w$ is the average of the within-population expected heterozygosities across SNPs over the two populations and $H_b$ is the between-population expected heterozygosity (Bhatia et al., 2013). All SNPs were used, without ascertainment.

***PSMC*** We applied the pairwise sequential Markovian coalescent method (`PSMC v. 0.6.5-r67`) to the genetic data simulated for one random sample from the CEU population and one from the YRI population. We generated the *PSMC* input file assuming a bin size of 100 bp as in Li and Durbin (2011), and ran the inference with: `-p 4+25*2+4+6` for time patterning, `-t 15` (number of $T_{\mathrm{MRCA}}$ bins), `-N 20` (number of iterations). In accordance with the mutation and recombination rates used for the simulations, the ratio of the mutation rate over the recombination rate (`-r`) was set as $r := \mu/\rho$ (=1.2 in the uniform recombination landscape case). We retrieved the last iteration (`-N 20`) for subsequent plotting and analyses. We note that the maximum number of iterations is lower than the one used in several empirical studies (`-N 25`) but (i) we did not observe any significant fit gain by running more iterations above 20 for several test scenarios and (ii) we wanted to speed-up the simulation process given the large number of parameter combinations that we had to test. As for all other temporal results reported in this study, we plotted *PSMC* curves in absolute years using a generation time of 25 years (Li and Durbin, 2011). Note that the scaling of times and population size assumed the mutation rate used in the simulation.

##### S1.3.2 Statistics sensitive to admixture

The simulated genetic datasets were also analyzed using statistics designed to detect and/or quantify *Hn* admixture from Neanderthal into Eurasian *Hs*: the *D*-statistics, the doubly conditioned site frequency spectrum (*DCFS*), the decay constant of the Neanderthal-ancestry *LD* curve ( $T_{LD}$), and summaries of the purportedly archaic introgression segments identified using *S'* and a conditional random field (*CRF*) approach.

***D*-statistic** We calculated the value of the *D*-statistic as *D(CEU, YRI; Vindija33.19, Outgroup)* considering the simulated 50 present-day CEU, 50 present-day YRI, the 50 kya Vindija33.19 Neanderthal genome and an outgroup individual (in our simulation framework, a virtual individual having the diploid ancestral alleles on every sites, but see Notes S10.3 for simulations with ancestral polarization errors). We used the allele frequency formula described in Appendix A of Patterson et al. (2012) and implemented in the Python library `scikit-allel 1.3.5`. The standard errors were estimated following Fu et al. (2015) (SI 16), using a weighted-block jackknife procedure discarding 5 cM blocks at each jackknife step. Using the jackknife-mean $D_j$ and the jackknife-SE ($SE_j$), we calculated the $Z$-score as $Z = D_j/SE_j$.

***DCFS*** The doubly conditioned site frequency spectrum (*DCFS*) was introduced by Yang et al. (2012) and its shape was suggested as a test to disentangle scenarios of admixture from ancient population structure. The *DCFS* is mathematically analogue to the derived *AFS* in CEU but is calculated for specifically ascertained SNPs supposedly enriched in signals of Neanderthal admixture, i.e. SNPs for which: (i) there is at least one ancestral allele in a randomly sampled YRI chromosome and (ii) one derived allele in a randomly sampled Neanderthal chromosome. Following the original article, we calculated the *DCFS* for five diploid CEU individuals and considered the simulated Vindija33.19 Neanderthal sample and one simulated YRI sample for ascertainment. In contrast with the procedure described in Yang et al. (2012), we averaged the *DCFS* over all 50 simulated YRI individuals while randomly sampling five CEU individuals at each iteration. We considered this averaging strategy to minimize the uncertainty associated with sampling a single chromosome in the YRI population. To mimic the haplotype-picking procedure in the Neanderthal and YRI genomes (as done in Yang et al. (2012)), if the simulated diploid genomes were homozygous, we took one of the duplicated alleles and if they were heterozygous, we sampled one of the two alleles with equal probability. We considered the *DCFS* bounds with derived allele counts from one to nine, and subsequently scaled the *DCFS* by the sum of all entries, so that each entry $i$ corresponded to the average probability of observing an ascertained SNP with derived allele count $i$ in the CEU sample.

**Note** We applied two other statistical approaches designed to identify segments of archaic ancestry in modern human genomes: *S'* (Browning et al., 2018) and *CRF* (Sankararaman et al., 2014). *S'* aims at identifying introgressed segments using a target population (here, CEU) and a purportedly non-admixed population (here, YRI) without requiring genomes of the admixing population (here, Neanderthals). Unlike *S'*, the conditional random field (*CRF*) approach uses an archaic genome to assess ascertainment and haplotype divergence patterns between the archaic genome and the CEU and YRI modern humans.

**Note on MAC filtering**

For modern humans, Sankararaman et al. (2014) and Browning et al. (2018) used the the 1000 Genomes Project dataset (Durbin et al., 2010), which is low coverage (on average 7.4×) with a significant under-representation of low-frequency variants (Sudmant et al., 2015). To mimic the features of the original empirical data that were used to call putative introgressed segments, we post-processed our simulated data to exclude some low-frequency variants, as a function of their original minor allele counts (MAC) in CEU and YRI. We followed the procedure described in Sankararaman et al. (2012), referred to as "MAC-filtering" henceforth. The acceptance rate $p_a$ of a SNP was set logarithmically proportional to its MAC in CEU and YRI:

$p_a$ = 0, 0.25, 0.50, 0.75, 0.80, 0.90, 0.95 0.96, 0.97, 0.98, 0.99 for minor allele counts equal to 0, 1, 2, 3, 4, 5, 6, 7, 8, 9, ≥10, respectively.

***S'*** For the *S'* analyses, we used the MAC-filtered dataset with the simulated 50 CEU and 50 YRI, considering only SNPs that were (i) polymorphic across the two populations and (ii) for which the frequency of the purportedly introgressed alleles in YRI was ≤ 1% (`-maxfreq` option). Based on the mutation rate used to generate the simulated data, we retained the segments with a score greater than the default threshold of 150,000 (Browning et al., 2018). As done in Browning et al. (2018), we estimated the match rate $m_{S'}$ between the Neanderthal genome and the CEU genomes at the $S'$-inferred segments. Unlike Browning et al. (2018) used the Altai Neanderthal, we used the 50 kya simulated Neanderthal (corresponding to Vindija33.19). We checked that the results were consistent between the two sampling options (Supplementary Notes S10.5). A SNP was considered matching the Neanderthal genome if the putative introgressed allele inferred by $S'$ at this location was found at least once in the Neanderthal genome. The match rate was then calculated as the total number of SNP matches within the retained segments over the total number of analyzed SNPs within the retained segments. Finally, we summarized the distribution of the physical lengths of all segments (whatever their match rate) using the mean and the 2.5% and 97.5% percentiles of the length distribution.

***CRF*** For the *CRF* analysis, we used the MAC-filtered dataset with the simulated 50 CEU and 50 YRI and the 50 kya Neanderthal genome—corresponding to Vindija33.19, as done in Sankararaman et al. (2014)—, considering only the SNPs that were polymorphic in CEU. The *CRF* method requires haplotype information, therefore we used the phased simulated genetic data for this analysis. For each SNP and each CEU haplotype, the emission probabilities (for the two states: introgressed or non-introgressed) were calculated using the fixed parameters published in the original study, as also done in, e.g. Choin et al. (2021). For each CEU haplotype, we identified a purportedly "introgressed segment" as a contiguous tract of SNPs with marginal posterior probabilities of Neanderthal ancestry $p_N$ greater than 90% and with a genetic length above 0.02 cM, following Sankararaman et al. (2012). We then calculated the introgression rate $\alpha_{\text{CRF}}$ as the proportion of SNPs with $p_N \geq 0.90$ over all analyzed SNPs for each haplotype, averaged over all CEU haplotypes. We summarized the distribution of the segment physical lengths using the mean and the 2.5% and 97.5% percentiles of the length distribution. We stress that this approach relies on a demographic model which assumes that Neanderthal admixture happened 1,900 generations ago (47,500 years ago assuming $g = 25$ years) with rate 3%, to calibrate the expected length of the recombining introgressed blocks. We thus used it as a summary statistics whose meaning is unclear (outside of the original model) but which has been computed in other studies (Choin et al., 2021).

**Ancestry-*LD* decay constant ($T_{LD}$, population-level)** In addition to the $S'$ and *CRF* approach which identify putatively introgressed segments, we also analyzed our simulated data with an approach based on the distribution of ancestry-informative linkage disequilibrium (ancestry-*LD*) (Moorjani et al., 2011; Sankararaman et al., 2012). Specifically, Sankararaman et al. (2012) implemented an approach (`computed`) estimating the LD distribution in a target population for a set of ascertained SNPs that are assumed informative about archaic (Neanderthal, here) admixture in a panmictic population model with single admixture pulse. They suggest that the exponential decay constant of the LD curve should then be proportional to the admixture age, as it provides a proxy of the distribution of the blocks of archaic ancestry in the genomes of the target population. For this analysis, we also used the MAC-filtered dataset with the 50 simulated CEU and performed the ascertainment with the 50 simulated YRI and the 50 kya-simulated Neanderthal genome (corresponding to Vindija33.19) to follow Sankararaman et al. (2012). We ascertained the SNPs following the authors' "Scheme 0", i.e. retaining only SNPs: (i) having at least one derived allele in the Neanderthal genome, (ii) a derived allele frequency in CEU ranging from 0 (excluded) to 10% (excluded).

We implemented our own version of the program to speed-up the calculation over thousands of simulations (because we can import the dataset just once to calculate several summary statistics, instead of repeating I/O operations by using `computed` separately) and also because we could not compile nor use the binary executable of `computed` on our computational resources. We checked that our implementation led to the same results as the original `computed` software on a local computer (cf. Notes S10.1).

Briefly, genetic distances were binned by bins of size $10^{-3}$ cM, from 0.02 to 1 cM, as in Sankararaman et al. (2012). Note however that in `computed`, the way the index $b$ of the bin (in 0-based indexing) to which any genetic distance $d$ is assigned is equivalent to:

$$b = \lfloor \frac{d - (m - w)}{w} \rfloor \tag{S2}$$

where $m$ is the minimum genetic distance considered (here, $m = 0.02$ cM) and $w$ is the binwidth ($= 10^{-3}$ cM). The intervals are thus defined by their right-bound. This implementation leads to collect genetic distance values which, in fact, start at $m - w = 0.019$ cM. However, in our implementation, we favoured defining intervals by the left-bound, leading to calculate $b$ as:

$$b = \lfloor \frac{d - m}{w} \rfloor \tag{S3}$$

We favoured this notation as the first bin (0.02 cM) corresponds to the distance values contained in the interval 0.02—0.021 (i.e. we collect distances which really start at 0.02 cM) instead of the interval 0.019—0.02 (which instead start at 0.019 cM), for consistency with the definition of the LD decay curve boundaries (i.e. "from 0.02 to 1.0 cM").

For each bin (with index $b$), we calculated $\bar{C}_b$ the average allelic covariance in CEU across the set $S$ of all pairs of SNPs $i$, $j$ separated by genetic distances contained in the bin $b$. Formally:

$$\bar{C}_b = \frac{1}{\#S_b} \sum_{S_b} \frac{\sum_{\#G_i} (G_i - \bar{G}_i)(G_j - \bar{G}_j)}{\#G_i - 1} \tag{S4}$$

where $\#S_b$ is the size of the set $S_b$ (i.e. the number of SNP pairs separated by genetic distances contained in the bin $b$); $G_i$ is the vector of genotypes of individuals in CEU at SNP $i$ and $\#G_i$ is the number of genotypes. Note that we calculated the unbiased covariance estimate instead of the biased one, as done in `computed` (where we would normalize by $\#G_i$).

To avoid fitting unrealistic exponential models to noisy LD decay curves, we set all covariance values $\bar{C}_b$ (for all $b$) as missing if more than half of the distance bins did not have a covariance estimate because of the absence of SNP pairs within. A last difference between the `computed` implementation and ours is that by default, we assume that covariance in any bin $b$ (when $S_b = \emptyset$) is NaN, whereas `computed` sets these missing values as 0.

Using a non-linear least squares regression, we fitted an exponential function ($y$=$Ae^{-td}$+$c$ with $y$ the allelic covariance and $d$ the genetic distance in Morgans) to the decay curve and retained the estimated coefficient $t$ as a measure of $T_{LD}$ (in generations), the purported time of Neanderthal admixture. To estimate the standard error for $T_{LD}$, we performed a weighted block-jackknife procedure following Moorjani et al. (2011) and Sankararaman et al. (2012), considering each chromosome as an independent block and setting the weights proportional to their respective SNP counts (Busing et al., 1999). We then calculated the confidence interval at 95% as $t_j \pm 1.96 \times SE_t$ where $t_j$ is the jackknife-mean decay constant and $SE_t$ the jackknife-SE. Note that in contrast to the analyses done in Sankararaman et al. (2012) for empirical data, we did not correct for errors in the genetic map since we used the ideal simulation-based recombination maps.

#### S1.4 Run selection

Among the one million simulated datasets, we retained a set of 20 runs which were considered the closest to the empirical data according to ten summary statistics, detailed in Table S3. Note that several of these statistics (e.g. *AFS*, *DCFS*...) are vectorial.

Table S3: Description of the summary statistics used for run selection.

| **Statistics** $S_a$ | **Description** | **Type of diversity analyzed** | **Weight** $w_a$ |
|---|---|---|---|
| *AFS* in YRI | | within-population diversity | 1 |
| *AFS* in CEU | | within-population diversity | 2 |
| *D*-statistic | | admixture | 2 |
| $T_{LD}$ | Mean* of the ancestry-*LD* decay constant | admixture | 2 |
| *SE(*$T_{LD}$*)* | SE* of the ancestry-*LD* decay constant | admixture | 1 |
| $L_{S'}$ | Average length of the *S'*-identified segments | admixture | 2 |
| $F_{ST}$ | (CEU; YRI) | between-population diversity | 1 |
| *DCFS* | | admixture | 2 |
| $\pi_{CEU}$ | | within-population diversity | 1 |
| $\pi_{YRI}$ | | within-population diversity | 1 |

* Jackknife-based values.

Thus, to estimate the global distance $D^\star$ of each simulation to the observed data, we first calculated the Euclidean distance $d_a$ between the simulated and the observed (scalar or vectorial) statistics, for each statistic $S_a$ (except for the *PSMC* curves, see below Notes S1.4.1):

$$d_a = \sqrt{\sum_{i=1}^{\#S_a} (S_{a,sim,i} - S_{a,obs,i})^2} \tag{S5}$$

with $S_{a,sim,i}$ the $i^{th}$ entry of the statistic $S_a$ (from the simulated data); $S_{a,obs,i}$ the $i^{th}$ entry of the statistic $S_a$ (from the observed data) and $\#S_a$ the size of $S_a$ ($\#S_a > 1$ if $S_a$ is vectorial, $\#S_a = 1$ if $S_a$ is a scalar). Naturally, for scalar statistics, $d_a$ is simply the absolute difference between $S_{a,sim}$ and $S_{a,obs}$. For each statistic $S_a$, we then scaled the distances $d_a$ across the simulations so that the distribution of distances for each statistic $S_a$ across all simulations has a mean absolute deviation (MAD) of one:

$$d_a^\star = \frac{d_a}{MAD(d_a)} \tag{S6}$$

with $MAD(d_a)$ the mean absolute deviation of the distances associated to the statistic $S_a$. Lastly, we calculated a single global distance index $D^\star$ for each simulation as:

$$D^\star = \sum_{a=1}^{\#S_a} w_a \cdot d_a^\star = \sum_{a=1}^{\#S_a} \frac{w_a \cdot d_a}{MAD(d_a)} \tag{S7}$$

where $w_a$ is the weight of the statistic $S_a$ (Table S3), so that statistics can contribute with varying importance to the global distance. We then ordered the simulations in increasing order of $D^\star$ and selected the twenty first simulations.

##### S1.4.1 Special case of *PSMC* distances

To compare the simulated *PSMC* curves ($\phi$) with the observed *PSMC* ($\phi_o$), we used the cross-correlation distance ($D_c$) as this distance allows to better represent the similarity between two curves in terms of inverse instantaneous coalescent rate (IICR) fluctuations, even if the two compared curves have temporal lags (i.e. shifts along the temporal axis). The distance was calculated as:

$$D_c = 1 - max(\{ACC(\phi, \phi_o, k)\}_{k \in [0; n_\phi - 1]}) \tag{S8}$$

with $ACC(\phi, \phi_o, k)$ the absolute value of the cross-correlation coefficient between $\phi$ and $\phi_o$ for lag $k$ and $n_\phi$ the number of step points in $\phi$. Note that prior to $D_c$ calculation, the *PSMC* curves were pre-processed: (i) they were truncated to the interval 10,000—8,000,000 years BP (since *PSMC* inference has low performance for the recent past and can generate spurious population size changes at the oldest time period) and (ii) aligned to the same step points to facilitate distance calculation. To do so, we interpolated the $y$-values of $\phi$ at the step points given in $\phi_o$ using the constant method of the R function `approx()` (i.e. for any point $p$, we took the value of the first more recent point, $p-1$, since the inference is done under the coalescent and therefore, backward in time) to match the step-wise visual nature of the *PSMC*.

#### S1.5 Observed statistics

For all studied genetic summary statistics, we retrieved the values estimated on the empirical data from previously published works. Details on how these statistics were estimated, the samples used and the corresponding values are described in Notes S2. Briefly, we retrieved:

- the genetic diversities $\pi$ in CEU and YRI from Keinan et al. (2009);
- the $F_{ST}$ between CEU and YRI from Bhatia et al. (2013);
- the *AFS* in CEU and YRI from Gutenkunst et al. (2009);
- the *PSMC* in a French and a Yoruba individual from Prado-Martinez et al. (2013);
- the $D$-statistic from Green et al. (2010);
- the *DCFS* from Yang et al. (2012);
- the $S$'-inferred putative Neanderthal introgressed segments from Browning et al. (2018);
- the *CRF*-inferred putative Neanderthal introgressed segments from Sankararaman et al. (2014);
- the $T_{LD}$ (putative age of Neanderthal admixture using the population-level ancestry-$LD$) from Sankararaman et al. (2012).

#### S1.6 Comparison with other published models

We assessed the performance of our structured model in comparison with a collection of eleven other published demographic models of human evolution which assume (i) archaic admixture from at least one archaic species (Neanderthal), (ii) a tree-like model of human evolution with (iii) simple panmictic populations or a very simple population structure (Yang et al., 2012; Fu et al., 2014; Skov et al., 2020; Jacobs et al., 2019; Ragsdale and Gravel, 2019; Durvasula and Sankararaman, 2020; Kamm et al., 2020; Gower et al., 2021; Iasi et al., 2021; Moorjani et al., 2016; Schaefer et al., 2021). Details on these demographic models (with visual representation) and on the simulation protocol and commands are provided in Notes S8.

The statistical approach that we used for model comparison was based on a **minimum scaled Euclidean distance** (MSED) for each model, noted here $D^{\star}_{M_i}$. Practically, for each published model, we retrieved the point estimates and simulated 50 datasets by varying the mutation rates uniformly from $5 \times 10^{-9}$ to $5 \times 10^{-8}$ bp$^{-1}$g$^{-1}$. We employed this varying mutation rate as a mean to introduce variance in the simulated genetic diversity and also because the original models were estimated with various mutation rate assumptions (usually $1.2 \times 10^{-8}$, $1.25 \times 10^{-8}$ or $2.5 \times 10^{-8}$) or did not rely on the mutation clock (Ragsdale and Gravel, 2019). For each published model, we then selected the 20 runs closest to the observed data, following the same procedure as described in Notes S1.4. We thus retained a total of 240 simulations: 20 for each of the 11 published models + 20 simulations for our structured model. Then, using a similar framework as described in Notes

S1.4, we calculated for each summary statistic $S_a$, the **per-statistic scaled Euclidean distance** $d^{\star}_{a,M_i}$, as the minimum scaled distance across the $n_i$ accepted simulations of the model $M_i$:

$$d^{\star}_{a,M_i} = \frac{min(\{d_a\}_{n_i})}{MAD(\{d_a\}_{\Omega})} \quad \text{(S9)}$$

where $\{d_a\}_{n_i}$ is the vector (of size $n_i = 20$) of the Euclidean distances (except for *PSMC*, see Notes S1.4.1) for the summary statistics $S_a$ across the $n_i$ accepted runs for model $M_i$; $MAD(\{d_a\}_{\Omega})$ is the mean absolute deviation of the distances $d_a$ across the $n_i$ accepted runs for *all* models (i.e. across 240 values).

To provide a single measure of model performance, we then calculated the minimum scaled Euclidean distance (MSED) $D^{\star}_{M_i}$ for each model $M_i$, as the average of the per-statistic scaled Euclidean distances across the 12 summary statistics (Table S4).

$$D^{\star}_{M_i} = \frac{1}{N_{stats}} \sum_{a=1}^{N_{stats}} d^{\star}_{a,M_i} = \frac{1}{N_{stats}} \sum_{a=1}^{N_{stats}} \frac{min(\{d_a\}_{n_i})}{MAD(\{d_a\}_{\Omega})} \quad \text{(S10)}$$

We note that the per-statistic distances are not weighted for model comparison, so that all summary statistics $S_a$ contribute equally to the final index. This means that our model is expected to be more disadvantaged since runs were selected on a different weighing scheme, but the uniform weighing of the model comparison did not appear to significantly impact our model performance.

Table S4: Description of the summary statistics used to calculate the MSED.

| **Statistics $S_a$** | **Description** | **Type of diversity analyzed** |
|---|---|---|
| *AFS* in YRI | | within-population diversity |
| *AFS* in CEU | | within-population diversity |
| *D*-statistic | | admixture |
| $T_{LD}$ | Mean* of the ancestry-*LD* decay constant | admixture |
| $L_{S'}$ | Average length of the *S'*-identified segments | admixture |
| $m_{S'}$ | Match rate of the *S'*-identified segments | admixture |
| $F_{ST}$ | (CEU; YRI) | between-population diversity |
| *DCFS* | | admixture |
| $\pi_{CEU}$ | | within-population diversity |
| $\pi_{YRI}$ | | within-population diversity |
| *PSMC* in YRI | | within-population diversity |
| *PSMC* in CEU | | within-population diversity |

* Jackknife-based values.

#### S1.7 Statistical analyses

To assess the accuracy of model predictions for particular statistics, we used the Root-Mean-Square Deviation (RMSD) (which is in the unit of the statistics) or the Mean Relative Absolute Error (MRAE) (which is dimensionless, thus allowing magnitude comparison across statistics).

For any summary statistics $S_a$ defined as a vector of size $\#S_a$ (if $S_a$ is scalar, $\#S_a = 1$):

$$RMSD = \sqrt{\frac{1}{\#S_a} \sum_{i=1}^{\#S_a} (S_{a,sim,i} - S_{a,obs,i})^2} \quad \text{(S11)}$$

$$MRAE = \frac{1}{\#S_a} \sum_{i=1}^{\#S_a} |\frac{S_{a,sim,i} - S_{a,obs,i}}{S_{a,obs,i}}| \quad \text{(S12)}$$

We also implemented the Relative Percentage Error (RPE) for scalar summary statistics as a more intuitive, dimensionless measure of discrepancy between the simulated and observed values:

$$RPE = 100 \cdot \frac{S_{a,sim} - S_{a,obs,i}}{S_{a,obs}} \tag{S13}$$

Unlike the RMSD or MRAE, that RPE values can be negative (when the model underestimates the observed value) or positive (when the model overestimates).

#### S1.8 Ancient DNA study

We investigated the values of two summary statistics ($D$, $T_{LD,1}$) in ancient samples that were simulated (i) under the 20 accepted runs of our structured model and (ii) Schaefer et al. (2021)'s model as this model generally ranked best among the published models that we compared. We chose these two statistics as they are easily and widely applied to aDNA samples.

##### S1.8.1 aDNA simulations under the structured model

Ancient Eurasian samples were simulated in the third deme (0-based indexing) of the $M_C$ metapopulation ; and at 10 different time points (from present up to 45 kya by steps of 5 kya). For each sampled deme and sampling time point, we simulated a single diploid individual to mimic the rarity of empirical aDNA data. All other samples (50 present-day YRI + Vindija33.19 Neanderthal) were the same as in the other simulations (Notes S1.2). We simulated individual genomes with twenty chromosomes of 30 Mbp, with uniform recombination rate ($1.0{\times}10^{-8}$) and a mutation rate of $1.2{\times}10^{-8}$.

##### S1.8.2 aDNA simulations under the Schaefer et al. (2021) model

We simulated single ancient samples in the CEU population at 10 different time points (from present up to 45 kya by steps of 5 kya), 50 samples in YRI at present at 1 Vindija Neanderthal sample at 50 kya in the deme named "Vindija" of the original model (Notes S8.11). As previously, we simulated genomes of size 20×30 Mbp, with uniform recombination rate ($1.0{\times}10^{-8}$) and a mutation rate of $1.2{\times}10^{-8}$. The generation time used for simulations as well as for the scaling of temporal statistics was the same as in our structured model: $g = 25$ years.

##### S1.8.3 Statistics

***D*-statistic** We calculated the $D$-statistic as: *D(1 ancient Eurasian, 50 present-day YRI; 1 Vindija Neanderthal, 1 Outgroup)*. In the simulations, the outgroup is a virtual individual having the diploid ancestral allele on every sites. Genotypes were not MAC-filtered. We used the frequency formula described in Appendix A of Patterson et al. (2012) and implemented in the Python library `scikit-allel` 1.3.5. The standard errors were estimated following Fu et al. (2015) (SI 16), using a weighted-block jackknife procedure discarding 5 cM blocks at each jackknife step. Using the jackknife-mean $D_j$ and the jackknife-SE ($SE_j$), we calculated the $Z$-score as $Z = D_j/SE_j$.

**Mean decay constant of the single-sample ancestry-*LD* curve ($T_{LD,1}$)** The previously described ancestry-$LD$ approach ($T_{LD}$) requires population-data in the target population and is therefore not suitable for cases where we have only a single sample, as witn aDNA. Fu et al. (2014) and Moorjani et al. (2016) developed a comparable approach but with a different implementation allowing to calculate the allelic covariance across pairs of alleles of putative Neanderthal ancestry within a single diploid sample. Thus, for our aDNA analyses, we used the single-sample ancestry-$LD$ approach with the ascertainment scheme 0 described in Moorjani et al. (2016), i.e.: (i) at least one derived allele in Neanderthal and (ii) only ancestral alleles in YRI. This ascertainement was originally suggested in order to minimize background correlation (i.e. pre-admixture) and highlight the Neanderthal ancestry signature. To match the pipeline of the original study, we did not use the MAC-filtered dataset. The ancestry-*LD* was then estimated using the authors' publicly available software (*github.com/priyamoorjani/Neanderthal_ dating*), based on the same parameters as for the multi-sample approach (genetic distances from 0.02 to 1 cM by steps of $10^{-3}$ cM). For each simulation, we retained the jackknife-mean and the standard error for $T_{LD,1}$ as estimated with the weighted jackknife procedure (Busing et al., 1999).

###### S1.8.4 Simulated genetic data quality

Given that empirical aDNA specimens have been genotyped and analyzed with different technologies and with varying data quality, we calculated the two statistics ($D$, $T_{LD,1}$) using different genetic datasets that we artificially transformed to best mimic empirical studies:

- **SNP sampling strategy**
  - ***All***: we used all the SNPs simulated over the 20×30 Mbp genomes—this SNP sampling would mimic empirical data obtained from high-coverage sequencing (e.g. Ust'-Ishim data with a coverage of 42×).
  - ***1M***: we used a random subset of one million SNPs from the *All* data—this sampling strategy would mimic the specimens genotyped on the 1240K enrichment capture (Fu et al., 2015).
  - ***Archaic***: we used a subset of ascertained SNPs from *All*–this strategy would mimic the specimens (like Oase1) genotyped on the Archaic SNP array. The ascertainment scheme that we applied followed Fu et al. (2015): to be retained, a SNP must have at least one Neanderthal allele which differs from the majority allele in a panel of 24 YRI diploids (sampled at random in our simulations).
- **Genotype properties**
  - **Diploid** genotypes—this mimics data for which were retrieved large quantities of DNA from the ancient specimens (e.g. Ust'-Ishim). This is the default genotype type for present-day humans.
  - **Pseudodiploidized** genotypes—this mimics lower-quality aDNA specimens (e.g. Oase1). This genotype type is common for ancient humans. In order to pseudodiploidize genotypes, we sampled at random one of the two alleles of the heterozygous genotypes and set it into two copies. Thus, a 0/1 genotype would randomly be assigned 0/0 with a probability of 50% or 1/1 with probability 50% (Skoglund et al., 2012).

###### S1.8.5 Empirical analysis

To compare the $D$ values estimated in our simulated ancient samples with the ones estimated on real ancient specimens, we could either retrieve the empirical statistics from the literature or re-calculate them ourselves. We discarded the first option as ancient specimens were described in independent studies which would use variable metrics of archaic ancestry. To improve the consistency and the reliability of the comparison, we thus re-calculated the *D(X, YRI; Vindija Neanderthal, Outgroup)* statistics separately for ten ancient specimens and for one present-day European population (28 individuals with MasterID "*French.DG*" from AADR v54.1). These empirical samples are described in Table S5. We used the Allen Ancient DNA Resource v54.1 dataset publicly available on the Reich's Lab website[1], which consists in 10,067 ancient samples genotyped at 1,233,013 positions (1240K panel).

[1] *https://reich.hms.harvard.edu/allen-ancient-dna-resource-aadr-downloadable-genotypes-present-day-and-ancient-dna-data*, retrieved on Dec. 30th 2022.

For each ancient specimen separately and for the modern-day "*French.DG*", the $D$-statistic was calculated for autosomal SNPs only, after excluding SNPs with at least one missing genotype over all analyzed individuals. The samples were:

- $X$: the ancient specimen or the 28 "*French.DG*" individuals,
- *YRI*: 24 present-day individuals with population label "*Yoruba.DG*",
- *Neanderthal*: the Shotgun diploid "*Vindija_ snpAD.DG*" (Vindija33.19) Neanderthal sample,
- *Outgroup*: ancestral reference with MasterID "*Ancestor.REF*".

Given that the $D$ estimates calculated by this mean are conditioned on the ascertainment of the SNP array, their values will necessarily differ from empirical estimates calculated on whole-genome data, and notably, from our simulated estimates where no such ascertainment bias exists. To correct for the ascertainment bias, we considered that the ascertained $D$ value estimated here on the "*French.DG*" population (1.82%) should be equal to the unbiased whole-genome $D$ value estimated on CEU in Green et al. (2010) (4.57%). We thus assumed conservative to rescale all $D$ values by a common multiplicative factor of $4.57/1.82 = 2.5$ (see Fu et al. (2015) p. 13 for a comparable strategy). The raw and scaled values of the $D$-statistics are provided in Table S5.

Table S5: Description of the ten ancient specimens and the modern French population from the AADR v54.1 dataset that we re-analyzed.

| Specimen | MasterID | GeneticID | Year* | Country | $^{14}$C** | $^{14}$C** (SE) | $D$ | SE | $D$ scaled |
|---|---|---|---|---|---|---|---|---|---|
| Bulgaria_BachoKiro_LatePleistocene_BB7 | BB7-240 | BB7-240_noUDG | 2021 | Bulgaria | 45,371 | 406 | 0.040 | 0.007 | 0.101 |
| Bulgaria_BachoKiro_LatePleistocene_CC7335 | CC7-335 | CC7-335_noUDG | 2021 | Bulgaria | 45,117 | 396 | 0.044 | 0.007 | 0.110 |
| Germany_EN_LBK_Stuttgart.DG | I0018 | Stuttgart.DG | 2014 | Germany | 7,168 | 70 | 0.016 | 0.004 | 0.041 |
| Luxembourg_Loschbour.DG | I0001 | Loschbour.DG | 2014 | Luxembourg | 8,025 | 64 | 0.019 | 0.005 | 0.049 |
| Romania_Oase_UP_enhanced | Oase1 | Oase1_noUDG | 2015 | Romania | 40,022 | 956 | 0.070 | 0.008 | 0.175 |
| Russia_Kostenki14 | Kostenki14 | Kostenki14 | 2014 | Russia | 38,052 | 725 | 0.018 | 0.005 | 0.045 |
| Russia_MA1_HG.SG | MA1 | MA1_noUDG.SG | 2013 | Russia | 24,320 | 120 | 0.019 | 0.005 | 0.048 |
| Russia_Ust_Ishim.DG | Ust_Ishim | UstIshim_snpAD.DG | 2014 | Russia | 44,366 | 816 | 0.021 | 0.005 | 0.052 |
| Russia_Yana_old2_UP.SG | Yana2 | Yana_old2_noUDG.SG | 2019 | Russia | 31,850 | 202 | 0.021 | 0.005 | 0.053 |
| French.DG | | | 2020 | France | 0 | 0 | 0.018 | 0.003 | 0.046 |

* Year of first publication.
** $^{14}$C radiocarbon age (years BP).

#### S2 Observed statistics in present-day modern humans

The values of the considered summary statistics estimated on observed genetic data ("observed statistics" henceforth) were retrieved from published studies (Table S6). Below, we detail the data and methods used in the original articles to produce these empirical estimates.

**Nucleotide diversity ($\pi$)** We extracted the nucleotide diversity measured in observed West Africans and North Europeans from Keinan et al. (2009). The authors used DNA sequences from five unrelated West Africans (four YRI + one African American with > 95% confidence in an African origin for both chromosomes) and five unrelated North European (three CEU + two European Americans) individuals from a public genomic database. Reads were aligned to the hg17 assembly using `ssahaSNP` (Ning et al., 2001). The mean number of aligned autosomal bases in West Africans was 475 Mb and 563 Mb in North Europeans. Then, a single mosaic of non-overlapping reads was generated for each individual, resulting in a haploid autosomal genome with limited strand bias. Nucleotide diversity was calculated by counting the number of differences at each position of the genome between two randomly sampled haploid genomes from a given population. Standard deviations were calculated using a jackknife with blocks of size 100 kb. The analysis resulted in estimates of $\pi = 0.827 \pm 0.004$ (mean $\pm$ SE) per kb in North Europeans and $\pi = 1.081 \pm 0.005$ per kb in West Africans.

***$F_{ST}$*** We retrieved the observed value of the $F_{ST}$ between CEU and YRI from Bhatia et al. (2013), since $F_{ST}$ was not computed in Keinan et al. (2009). The authors used the 1000 Genomes Project Phase 1 database, with 59 unrelated YRI individuals and 60 unrelated CEU individuals. Considering only autosomes, they computed $F_{ST}$ using different estimators and found that the Hudson's estimator (calculated as a ratio of averages across multiple SNPs) was less biased than the average of ratios and more robust to unbalanced sample sizes. Specifically, for any given SNP, the ratio of averages for the Hudson's estimator $\hat{F}_{ST}$ is (Hudson et al., 1992):

$$\hat{F}_{ST} = 1 - \frac{H_w}{H_b} \tag{S14}$$

with $H_w$ the mean number of differences within-CEU and within-YRI averaged across SNPs, and $H_b$ the mean number of differences between CEU and YRI averaged across SNPs. This can be decomposed in terms of allele frequencies as:

$$\hat{F}_{ST} = \frac{(p_1 - p_2)^2 - \frac{p_1(1-p_1)}{n_1-1} - \frac{p_2(1-p_2)}{n_2-1}}{p_1(1-p_2) + p_2(1-p_1)} \tag{S15}$$

with $p_i$ the mean allele frequencies in population $i$; $n_i$ the haploid sample size in population $i$. Using this method, they estimated $\hat{F}_{ST} = 0.139 \pm 5 \times 10^{-4}$ (mean $\pm$ SE) (for SNPs polymorphic in YRI), $\hat{F}_{ST} = 0.141 \pm 6 \times 10^{-4}$ (for SNPs polymorphic in CEU), $\hat{F}_{ST} = 0.142 \pm 7 \times 10^{-4}$ (for SNPs polymorphic in both CEU and YRI). We considered the $\hat{F}_{ST}$ value calculated for SNPs polymorphic in YRI but given the marginal difference between the three reported $\hat{F}_{ST}$ values, any one of these would lead to very comparable results in our study.

***AFS*** We retrieved the observed derived 2D-*AFS* from Gutenkunst et al. (2009) which was calculated using the NIEHS Environmental Genome Project SNP database (Sanger resequencing), with 22 unrelated YRI and 12 unrelated CEU individuals. The cumulated genomic segments considered spanned 5.01 Mb, located in noncoding regions of 219 autosomal genes. Ancestral alleles were identified using the panTro2 chimpanzee reference, with further correction of ancestral allele misidentification using a context-dependent substitution model (e.g. taking into account CpG effects). Due to the high quality of the original data, no correction was applied for possible sequencing errors. To account for missing data, they projected the *AFS* down to 10 diploids in each population using a hypergeometric distribution. The downscaling resulted in a total of 15,726 SNPs for CEU and YRI. For ease of comparison with our simulations, we then scaled the 2D-*AFS* by dividing each entry with the sum across all entries: each entry thus represented the probability that a SNP was found at any joint frequency $f_{CEU}$ in CEU and $f_{YRI}$ in YRI. We also calculated each population-specific 1D-*AFS* by summing over the 2D-*AFS* row-wise or column-wise. Each population-specific 1D-*AFS* was then scaled by dividing each entry with the sum across all entries. The derived 2D-*AFS* was downloaded from GADMA GitHub[2].

***PSMC*** We used the observed *PSMC* trajectories originally estimated in Prado-Martinez et al. (2013) (kindly provided by Arredondo et al. (2021)) from the diploid whole-genome resequencing data of a French and a Yoruba individual (HGDP) aligned on the hg18 human assembly. The authors ran `PSMC v. 0.6.5-r67` with the options `-N25 -t15 -r5 -p 4+25*2+4+6` (Notes S12.4 of the original study). When comparing and plotting the empirical *PSMC* ($\phi_o$) trajectory along with the simulated *PSMC*(s), we used the 20$^{\text{th}}$ iteration of $\phi_o$ and converted it into an IICR trajectory assuming $g = 25$ years and the same mutation rate as the one used to produce the compared simulated *PSMC*(s).

***D*-statistic** We retrieved the observed value of the *D*-statistic *D(CEU, YRI; Vindija Neanderthal, Chimpanzee)* calculated in Green et al. (2010) (original Supplementary Materials SOM15, Table S44). The authors calculated *D* using sequencing reads produced either with the ABI3730 or the Illumina $GA_{II}$ technology, but we used the values estimated for the ABI3730 data as these were found to be more precise (SOM 15). The authors generated sequences for four YRI and two CEU using the ABI3730 sequencer producing 752 bp-long reads on average. Variants were called with `ssahaSNP` (Ning et al., 2001) and subsequently filtered to retain those which (i) mapped to the panTro2 sequence, (ii) had a Phred-scaled quality score $\geq 40$, (iii) had a Phred-scaled quality score on the five flanking bases $\geq 15$, (iv) had a good mappability and copy number stability relatively to panTro2. The Neanderthal genome used was the pooled Vindija introduced in the same article (coverage $\sim$30$\times$). *D* was then calculated using allele frequencies in each population (formula S15.2 from the original study), where each frequency at a given site was estimated by pooling the reads of all individuals from a given population, weighted by the read coverage in each individual at the site (Green et al., 2010). The standard error was originally computed using a weighted block jackknife with 800 contiguous blocks, as this roughly corresponds to a block length of $\sim$4 cM, but the authors showed that the results are stable as soon as the blocks are $> 2$ cM (Table S44 from the original study). The resulting *D* is equal to $4.57\% \pm 0.33\%$ (mean $\pm$ SE) ($Z = 13.8$) (Table S44 from the original study). We note that the value of *D* was comparable when the authors used other sequencing technologies.

***DCFS*** The observed doubly conditioned site frequency spectrum (*DCFS*) was retrieved from Yang et al. (2012). The authors used 7 YRI individuals and 5 CEU individuals from the CGDP database (Drmanac et al., 2010) sequenced at an average depth of 45$\times$. For Neanderthal, they used the pooled Vindija sequence, filtered using the Green et al. (2010) protocol. The ancestral alleles were extracted from the 1000 Genomes Project database (estimated using a common alignment of humans, chimpanzees, orangutans and rhesus macaque genomes). Only biallelic transversion SNPs were considered, to minimize the impact of substitutions due to DNA damage (Yang et al., 2012). The doubly conditioned ascertainment of SNPs was performed by (i) randomly sampling a single read in the Neanderthal BAM file at a given position and (ii) randomly sampling a single allele in the YRI population at this position. A SNP was taken into account if the allele was (i) derived in Neanderthal and (ii) ancestral in YRI. The *DCFS* was calculated by counting the number of derived alleles in CEU at each of these ascertained SNPs. We used the scaled *DCFS*, i.e. where each entry was divided by the total sum of SNPs in the *DCFS*.

[2] https://github.com/noscode/GADMA/blob/master/examples/optimize_ga_for_YRI_CEU/YRI_CEU.fs

***S'*** We retrieved the results of the empirical *S'* analysis performed by Browning et al. (2018) from a public repository[3]. The authors used the 1000 Genomes Phase 3 dataset and the HapMap genetic map[4], considering 90 CEU individuals as the target population and 108 YRI individuals at the outgroup population. For the *S'* analysis, they set the minimum MAF cutoff in YRI at 1%. For any given SNP, the *S'*-inferred ancestral allele was compared to the allelic content in Neanderthal using the Altai Neanderthal genome (52×) as reference. The Altai variants were masked if coverage depth was < 10×, Phred-scaled mapping quality score < 25 or if they were near indels/tandem repeats or regions with poor mappability. We downloaded the original results from the *CEU_sprime_results.tar.gz* archive. We retrieved each identified segment with a score greater than 150,000, following Browning et al. (2018). Following the original study, the *S'*-inferred ancestral allele was declared to match Neanderthal if it was present in at least one copy in Neanderthal, else it was declared a mismatch (or was set as missing if the site was masked in the filtered Neanderthal). For each segment, we counted the total number of SNPs with allelic match ($n_1$) and the total number of SNPs with allelic mismatch ($n_0$). The match rate $m_{S'}$ was estimated as $n_1/(n_0 + n_1)$ (following (Browning et al., 2018)) and the segment length as the difference between the physical positions of the first and last SNP within. If the segment comprised less or as many as 3 SNPs, it was discarded. We then calculated the mean, standard deviation and 2.5%- and 97.5%-percentiles of the distribution of match rates and segment lengths across all retained segments. These analyses resulted in a mean match rate of 72.6% ± 3.7% (mean ± SD) and a mean length of 353.7 kb ± 163.7 kb (mean ± SD) in CEU.

***CRF*** We retrieved the results of the empirical *CRF* analysis performed by Sankararaman et al. (2014) from their article (Table 1) and Supplementary Data[5]. The authors used the 1000 Genomes Phase 1 dataset, considering 176 phased diploid YRI invididuals and 85 phased CEU individuals. The genetic map was derived from the combined Oxford LD map (Myers et al., 2005). The Neanderthal sample used was the Altai Neanderthal (52×). Alleles were polarized using the panTro2 sequence. The Neanderthal variants were masked if the Phred-scaled genotype quality score was < 30, Phred-scale mapping quality score < 30, or if they were near tandem repeats or regions with poor mappability. The analysis was run on biallelic SNPs which were polymorphic in the target population (i.e. CEU, here), with the parameters $\lambda = 1900$, $\theta = 0.97$ and `onlypolymorphic = 1`. Considering sites with a minimum marginal probability of Neanderthal ancestry of 90%, they estimated a genome-wide proportion of Neanderthal ancestry on autosomes averaged across all CEU individuals as 1.17% ± 0.08% (mean ± SD) (Table 1 of (Sankararaman et al., 2014)). We calculated the confidence interval at 95% as 1.17% ± 1.96 × 0.08%. To estimate the distribution of the high-confidence Neanderthal tiling paths ("segments" henceforth) in CEU, we retrieved the results from the original Supplementary Data, considering the files *chr-*.thresh-90.length-0.00.gz* which provide the list of analyzed SNPs with the marginal probability of Neanderthal ancestry averaged across all CEU haplotypes. To estimate the distribution of segment lengths, we considered all the segments with an average marginal probability greater than 90% and a size greater than 0.02 cM, following Sankararaman et al. (2014). Then, for each CEU haplotype, we stored the length of all detected segments. We then calculated the mean, standard deviation and 2.5%- and 97.5%-percentiles of the segment lengths across all the CEU haplotypes taken together, resulting in a mean length of 114.1 kb ± 153.7 kb (mean ± SD). We stress here that this method assumes a demographic model with Neanderthal admixture happening 1,900 generations ago.

**Ancestry-*LD* decay constant ($T_{LD}$)** We considered the putative age of Neanderthal admixture estimated by Sankararaman et al. (2012) with the population-level ancestry-*LD* approach (software `computed`). The authors used the 1000 Genomes Pilot 1 data on the CEU (target) and YRI (outgroup) populations, with the the pooled Vindija (Green et al., 2010) as the Neanderthal sample. SNPs were polarized using the chimpanzee reference panTro2. We considered the results obtained by

[3] https://data.mendeley.com/datasets/y7hyt83vxr/1
[4] http://bochet.gcc.biostat.washington.edu/beagle/genetic_maps/
[5] https://reich.hms.harvard.edu/datasets/landscape-neandertal-ancestry-present-day-humans (Part 4)

the authors with their default ascertainment "Scheme 0", i.e. for SNPs which: (i) are polymorphic in CEU; (ii) have a derived allele frequency below 10% in CEU and (iii) have at least one derived allele in Neanderthal. They estimated the ancestry-$LD$ curve as the covariance between genotypes for each pair of ascertained SNPs separated by genetic distances ranging from 0.02 cM to 1.0 cM (by steps of $10^{-3}$ cM) (Notes S10.1). They estimated the exponential decay constant (after fitting a single-term exponential model with ordinary least squares regression) and corrected its value to account for genetic map uncertainty at short distances. As a point estimate for the putative age of admixture, we averaged the corrected ages ($t_{GF}$) estimated by the authors using the Decode map (Kong et al., 2010) and the CEU Oxford LD map (Myers et al., 2005) ($1,900$ and $1,961$ generations ago, respectively) (Table S6 in the original study).

| Statistics | Reference | African | European | Neanderthal | Outgroup | Genome | Method | Values*** |
|---|---|---|---|---|---|---|---|---|
| Genetic diversity $\pi$ | Keinan et al. 09 | 5 unrelated West-Africans (incl. 4 YRI) | 5 unrelated North Europeans (incl. 3 CEU) | | | WGS, autosomes | Average number of divergent sites per base pair | CEU: 0.827 $\pm^{SE}$ 0.004 kb$^{-1}$, YRI: 1.081 $\pm^{SE}$ 0.005 kb$^{-1}$ |
| $F_{ST}$ | Bhatia et al. 13 | 59 unrelated YRI (1KG) | 60 unrelated CEU (1KG) | | | WGS, autosomes | Hudson's $F_{ST}$, ratio of averages | 0.139 $\pm$ $5 \cdot 10^{-4}$ (polymorphic in YRI) |
| 2D-*AFS* 1D-*AFS* | Gutenkunst et al. 19 | 22 unrelated YRI | 12 unrelated CEU | | panTro2 | NIEHS SNP database, non-coding regions | Projected down to 10 diploids per population, ancestral allele misidentification correction | cf. GitHub |
| *PSMC* | Prado-Martinez et al. 13 | 1 Yoruba (HGDP) | 1 French (HGDP) | | | WGS | | cf. GitHub |
| *D*-statistic | Green et al. 10, Table S44 | 4 YRI | 2 CEU | Vindija (pooled*, not pseudodiploidized) | panTro2 | Whole-genome, autosomes, ABI3730 sequencing | Weighted block jackknife over 4 Mb blocks | 4.57% $\pm^{SE}$ 0.33% |
| *DCFS* | Yang et al. 12 | 7 YRI (CGDP) | 5 CEU (CGDP) | Vindija (pooled*, BAM file) | 1KG ancestral | WGS (45×), autosomes | | cf. GitHub |
| *S'* | Browning et al. 18 | 108 YRI (1KG3) | 99 CEU (1KG3) | Altai (52×) | | WGS, autosomes; HapMap map | MAF$_{YRI}$=1%, score >150k | Mean match rate**: 72.6% $\pm^{SD}$ 3.7%; mean length**: 353.7 kb $\pm^{SD}$ 163.7 kb |
| *CRF* | Sankararaman et al. 14 | 176 phased YRI (1KG1) | 85 phased CEU (1KG1) | Altai (52×) | panTro2 | WGS, autosomes; Oxford combined LD map | Marginal probability > 0.90, min length 0.02 cM | Introgression rate: 1.17% $\pm^{SD}$ 0.08% (Table 1); mean length**: 114.1 kb $\pm^{SD}$ 153.7 kb |
| Ancestry-*LD* | Sankararaman et al. 12 | YRI (1KG1) | CEU (1KGI) | Vindija (pooled*) | panTro2 | WGS, autosomes | Empirical map correction | 1,900—1,961 generations BP |

Table S6: Summary of samples, methods and data used to estimate observed statistics. We also report their estimated values.
* Pooled Vindija refers to the Neanderthal sequence obtained by pooling Vi33.16, Vi33.25 and Vi33.26 (Green et al., 2010).
** Re-estimated from the Supplementary Data provided by the authors.
*** We note $\pm^{SE}$ for standard errors and $\pm^{SD}$ for standard deviations.

### S3 Collinearity between statistics

Figure S2 represents the matrix of correlation between the distances to the observed for the summary statistics estimated in the twenty accepted runs of our structured model. It shows blocks of statistics positively inter-correlated, as well as statistics negatively correlated. This suggests that whenever the fit of certain statistics is improved (by changing the parameters of the model, for instance), the fit of other (anticorrelated) statistics worsens, highlighting optimization trade-offs.

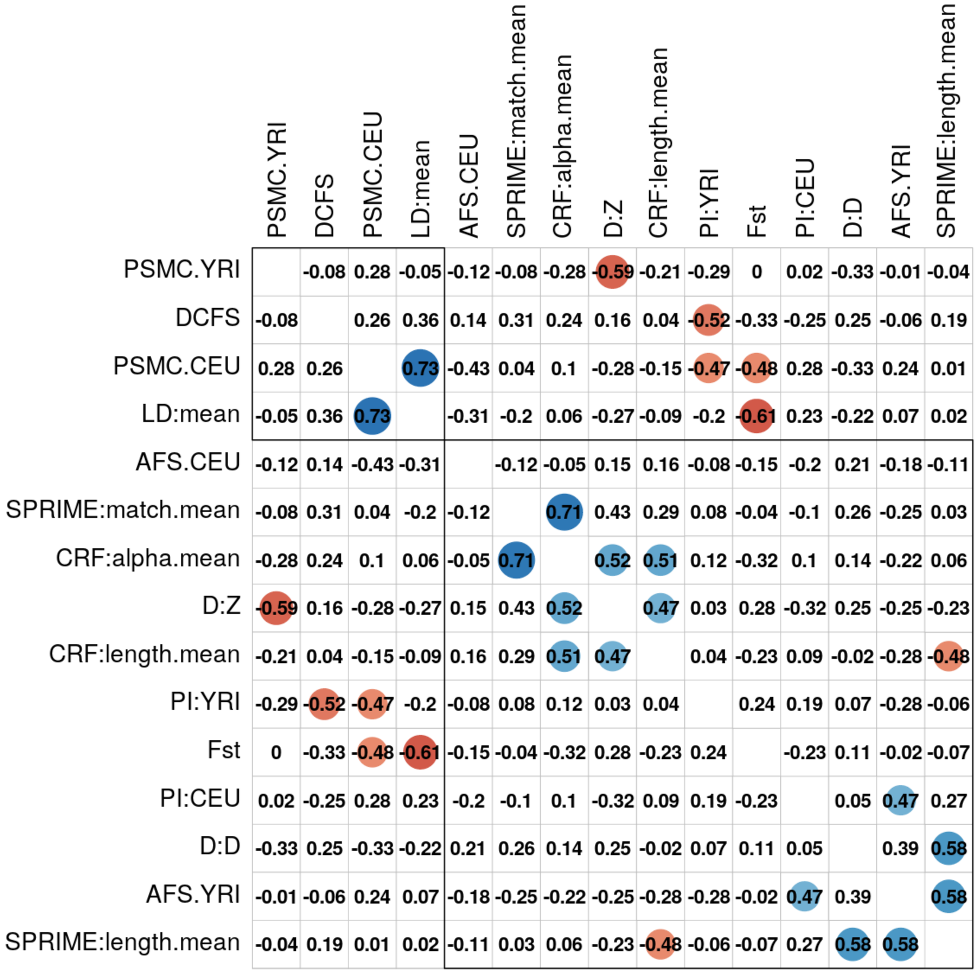

Figure S2: Pearson's pairwise correlation coefficients of the distances to the observed between fifteen genetic statistics estimated on the twenty selected runs of our structured model. Note that the correlated values are not the raw values of the statistics, but their Euclidean distance to the empirical statistics, except for *PSMC* for which we used the cross-correlation distance. When a correlation coefficient was significant at a level of 5%, we added a circle whose size and color were proportional to the correlation coefficient (blue if positive, red if negative). Statistics were ordered and clustered into two groups according to a hierarchical clustering algorithm (R package `corrplot`).
**Notes.** "LD:mean" is $T_{LD}$, "CRF:alpha.mean" is $\alpha_{CRF}$, "D:D" is $D$, "D:Z" is the $Z$-score for $D$, "SPRIME:match.mean" is $m_{S'}$.

### S4 Model performance with heterogeneous recombination

Figures in this section S4 were obtained assuming a **heterogeneous landscape of recombination** (cf. Notes S1.2).

**Caption of Figure S3:** Performance of our admixture-free model compared to observed data (black) for the twelve statistics used in this study, assuming a heterogeneous landscape of recombination with hotspots. The statistical distributions were obtained from the twenty accepted runs of our model, after simulating twenty chromosomes of 30 Mbp. **Abbreviations.** CI: confidence interval; IQR: interquantile range; IICR: inverse instantaneous coalescence rate. **(A)** Nucleotide diversity in CEU and YRI. We represent the 2D density (Gaussian kernel) of the joint nucleotide diversities $\pi$ estimated in simulated CEU and YRI samples. The black dot represents the values estimated from real data. **(B)** Normalized CEU *AFS*. The black dots represent the observed *AFS* using ten real CEU individuals. The colors represent the 50%, 80%, 95% and 100% interquantile ranges of the normalized derived allele frequency spectrum calculated for ten simulated CEU samples across the twenty selected runs. The black line is the median across the simulated *AFS*s. RMSE is the root mean squared error and is given in percentage. **(C)** Normalized YRI *AFS*. Same as (B), for ten YRI samples. **(D)** CEU *PSMC* curves. The black curve is the observed CEU *PSMC* whereas the coloured *PSMC* curves were obtained from the twenty selected runs. The $x$-axis ($\log_{10}$-scaled) is expressed in years before present (with $\mu = 1.2 \times 10^{-8}$ and g=25 years). **(E)** YRI *PSMC* curves. Same as (D), for a single YRI sample per selected run. **(F)** Joint distribution of the $F_{ST}$ and $D$-statistic. The $D$-statistic ($x$-axis) was calculated as $D$(50 CEU, 50 YRI; 1 Neanderthal, Ancestral reference) and the $F_{ST}$ ($y$-axis) was computed between the CEU and YRI samples. Both statistics were computed for each of the twenty runs and plotted jointly using a 2D density (Gaussian kernel). **(G)** CEU *DCFS*. As in panel (B), an interquantile ribbon plot was obtained for the doubly conditioned site frequency spectra across the twenty selected runs. The *DCFS* was computed for 5 CEU individuals. The black line is the median *DCFS* across the twenty runs. The black dots represent the observed *DCFS*. **(H)** Joint distribution of mean Neanderthal fragments lengths and match rates. A 2D density (Gaussian kernel) of the distribution for the mean length ($x$-axis, in bp, $\log_{10}$-scaled) and the mean match rate $m_{S'}$ ($y$-axis) is represented for the putative Neanderthal segments estimated with *S'* in the 50 simulated CEU samples. The black dot represents the estimates from the observed data (with associated 95% IQR). **(I)** Ancestry-*LD* decay constant. The $x$-axis represents the exponential decay constant of the ancestry-*LD* curve, and the twenty selected runs are represented on the $y$-axis. The point estimates (square dots) are classically interpreted as the age of the admixture events. The whisks represent the 50% (darkest thickest segment) and the 95% confidence intervals (colored segment with transparency) around the jackknife mean (calculated using a weighted jackknife procedure). The point estimate of the decay constant computed from real data is ~50 kya (vertical dashed line).

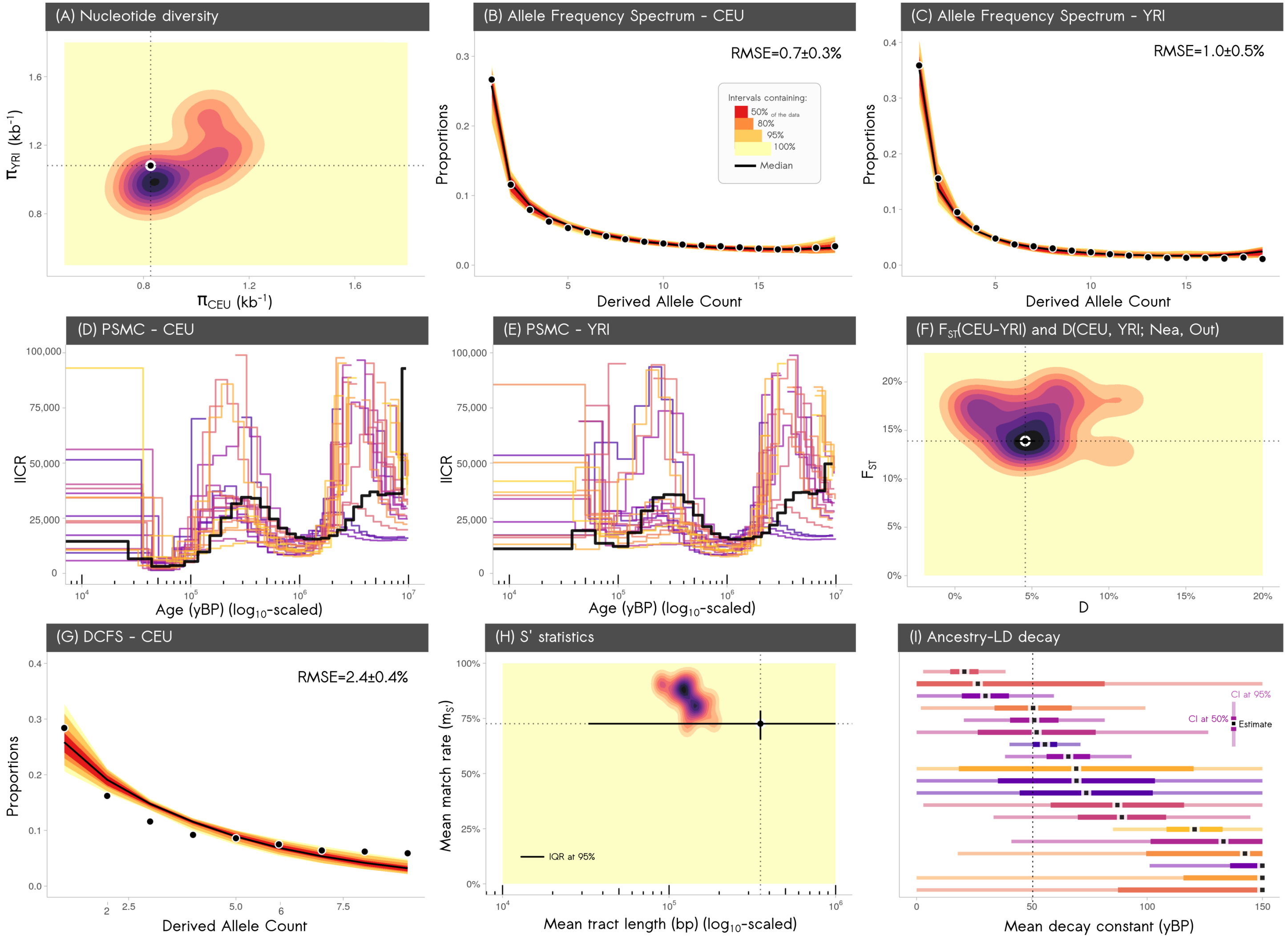

Figure S3: See the figure caption above.

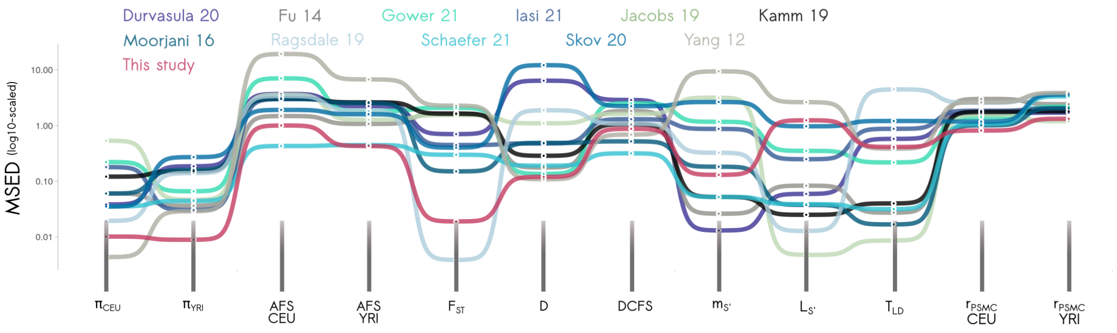

Figure S4: **Model comparison assuming a heterogeneous landscape of recombination with hotspots.** This figure compares the predictions of the eleven published models for twelve statistics, with those of our 1D structured model (red). All models except ours assume Neanderthal admixture into *Hs*. The statistics are represented in the $x$-axis, and each model is represented by a different color. The $y$-axis represents the minimum scaled Euclidean distance (MSED) to the observed statistics for twenty accepted runs from each model. The $y$-axis is $\log_{10}$-scaled.

### S5 Parameters of the 20 accepted runs from the structured model

Table S7: Values of the 25 free parameters for the first ten runs among the twenty accepted. For details about the parameters, see Table S1.1.

| RunID: | 1012447 | 1014230 | 1050670 | 1161505 | 152019 | 200752 | 234400 | 344388 | 35483 | 483636 |
|---|---|---|---|---|---|---|---|---|---|---|
| N1 | $2.12\times10^{4}$ | $3.40\times10^{4}$ | $2.87\times10^{4}$ | $1.46\times10^{4}$ | $2.40\times10^{4}$ | $6.53\times10^{3}$ | $3.96\times10^{3}$ | $5.09\times10^{3}$ | $3.80\times10^{4}$ | $2.39\times10^{4}$ |
| N2 | $1.39\times10^{3}$ | $2.30\times10^{3}$ | $3.17\times10^{3}$ | $3.03\times10^{3}$ | $2.59\times10^{3}$ | $1.38\times10^{3}$ | $7.96\times10^{3}$ | $2.08\times10^{3}$ | $2.27\times10^{3}$ | $2.61\times10^{3}$ |
| N4 | $5.10\times10^{2}$ | $2.02\times10^{1}$ | $2.90\times10^{2}$ | $3.77\times10^{2}$ | $1.29\times10^{1}$ | $1.23\times10^{2}$ | $1.20\times10^{3}$ | $3.07\times10^{2}$ | $1.97\times10^{2}$ | $1.43\times10^{1}$ |
| N3 | $5.30\times10^{4}$ | $1.35\times10^{4}$ | $2.34\times10^{4}$ | $2.51\times10^{4}$ | $1.10\times10^{4}$ | $4.28\times10^{4}$ | $1.39\times10^{4}$ | $1.79\times10^{4}$ | $1.43\times10^{4}$ | $1.33\times10^{4}$ |
| m1 | $4.11\times10^{-2}$ | $4.31\times10^{-2}$ | $3.11\times10^{-2}$ | $2.06\times10^{-3}$ | $6.51\times10^{-2}$ | $5.09\times10^{-3}$ | $1.64\times10^{-3}$ | $9.59\times10^{-3}$ | $1.36\times10^{-2}$ | $3.28\times10^{-2}$ |
| m2 | $2.31\times10^{-6}$ | $9.05\times10^{-5}$ | $1.28\times10^{-4}$ | $1.50\times10^{-4}$ | $3.35\times10^{-2}$ | $7.42\times10^{-4}$ | $4.59\times10^{-4}$ | $6.42\times10^{-4}$ | $6.06\times10^{-3}$ | $1.07\times10^{-6}$ |
| m3 | $4.68\times10^{-2}$ | $1.23\times10^{-2}$ | $2.64\times10^{-4}$ | $6.77\times10^{-3}$ | $1.12\times10^{-2}$ | $8.05\times10^{-4}$ | $1.13\times10^{-5}$ | $8.41\times10^{-4}$ | $1.23\times10^{-3}$ | $4.29\times10^{-2}$ |
| m4 | $4.85\times10^{-4}$ | $8.92\times10^{-5}$ | $1.16\times10^{-5}$ | $1.19\times10^{-5}$ | $1.11\times10^{-7}$ | $2.56\times10^{-4}$ | $1.15\times10^{-8}$ | $1.25\times10^{-8}$ | $2.03\times10^{-7}$ | $7.03\times10^{-4}$ |
| m5 | $1.46\times10^{-2}$ | $3.39\times10^{-3}$ | $8.98\times10^{-3}$ | $1.29\times10^{-3}$ | $1.53\times10^{-3}$ | $3.21\times10^{-1}$ | $6.95\times10^{-3}$ | $2.07\times10^{-3}$ | $1.51\times10^{-1}$ | $6.47\times10^{-2}$ |
| m6 | $5.38\times10^{-5}$ | $3.81\times10^{-6}$ | $8.00\times10^{-5}$ | $5.34\times10^{-6}$ | $3.03\times10^{-6}$ | $1.35\times10^{-5}$ | $5.85\times10^{-7}$ | $6.29\times10^{-7}$ | $3.11\times10^{-7}$ | $2.31\times10^{-5}$ |
| T1 | $4.91\times10^{4}$ | $5.39\times10^{4}$ | $5.58\times10^{4}$ | $4.15\times10^{4}$ | $4.84\times10^{4}$ | $6.12\times10^{4}$ | $5.27\times10^{4}$ | $6.40\times10^{4}$ | $6.52\times10^{4}$ | $4.57\times10^{4}$ |
| dT1 | $7.97\times10^{2}$ | $2.77\times10^{1}$ | $9.57\times10^{1}$ | $5.36\times10^{2}$ | $8.98\times10^{2}$ | $1.14\times10^{3}$ | $3.19\times10^{2}$ | $1.08\times10^{3}$ | $5.97\times10^{2}$ | $5.83\times10^{1}$ |
| n3 | $1.82\times10^{3}$ | $3.09\times10^{2}$ | $5.44\times10^{1}$ | $1.84\times10^{3}$ | $3.15\times10^{0}$ | $1.91\times10^{3}$ | $1.62\times10^{2}$ | $2.24\times10^{3}$ | $3.10\times10^{0}$ | $2.95\times10^{1}$ |
| dT2 | $3.64\times10^{0}$ | $5.07\times10^{2}$ | $1.45\times10^{2}$ | $1.86\times10^{2}$ | $3.68\times10^{1}$ | $1.63\times10^{3}$ | $7.08\times10^{2}$ | $1.85\times10^{3}$ | $5.34\times10^{0}$ | $2.04\times10^{1}$ |
| m7 | $4.64\times10^{-4}$ | $8.26\times10^{-4}$ | $3.14\times10^{-4}$ | $1.30\times10^{-5}$ | $1.68\times10^{-2}$ | $1.97\times10^{-3}$ | $1.12\times10^{-4}$ | $3.13\times10^{-3}$ | $3.13\times10^{-3}$ | $2.15\times10^{-2}$ |
| m8 | $2.54\times10^{-3}$ | $2.47\times10^{-3}$ | $2.49\times10^{-3}$ | $2.10\times10^{-3}$ | $2.21\times10^{-2}$ | $5.58\times10^{-3}$ | $8.56\times10^{-2}$ | $5.74\times10^{-3}$ | $4.02\times10^{-3}$ | $1.95\times10^{-2}$ |
| T2 | $2.88\times10^{5}$ | $4.53\times10^{5}$ | $2.82\times10^{5}$ | $3.70\times10^{5}$ | $4.41\times10^{5}$ | $2.04\times10^{5}$ | $4.24\times10^{5}$ | $3.88\times10^{5}$ | $8.41\times10^{5}$ | $4.06\times10^{5}$ |
| m9 | $3.00\times10^{-5}$ | $2.36\times10^{-5}$ | $2.98\times10^{-5}$ | $1.85\times10^{-5}$ | $2.47\times10^{-5}$ | $2.68\times10^{-5}$ | $1.29\times10^{-5}$ | $2.85\times10^{-5}$ | $1.70\times10^{-5}$ | $1.93\times10^{-5}$ |
| m10 | $4.88\times10^{-7}$ | $7.50\times10^{-7}$ | $7.81\times10^{-7}$ | $5.49\times10^{-7}$ | $3.81\times10^{-7}$ | $6.69\times10^{-7}$ | $4.15\times10^{-7}$ | $5.59\times10^{-7}$ | $5.27\times10^{-7}$ | $5.83\times10^{-7}$ |
| T3 | $9.61\times10^{5}$ | $4.63\times10^{5}$ | $4.35\times10^{5}$ | $5.48\times10^{5}$ | $6.97\times10^{5}$ | $5.63\times10^{5}$ | $4.88\times10^{5}$ | $9.95\times10^{5}$ | $8.47\times10^{5}$ | $5.06\times10^{5}$ |
| N5 | $1.34\times10^{3}$ | $1.06\times10^{3}$ | $7.95\times10^{2}$ | $1.06\times10^{3}$ | $7.46\times10^{2}$ | $8.45\times10^{2}$ | $7.97\times10^{2}$ | $1.03\times10^{3}$ | $1.41\times10^{3}$ | $1.01\times10^{3}$ |
| T4 | $6.18\times10^{6}$ | $6.93\times10^{6}$ | $6.94\times10^{6}$ | $8.29\times10^{6}$ | $5.97\times10^{6}$ | $8.98\times10^{6}$ | $3.80\times10^{6}$ | $5.29\times10^{6}$ | $6.26\times10^{6}$ | $7.10\times10^{6}$ |
| $p_{YRI}$ | $4.38\times10^{-1}$ | $7.09\times10^{-1}$ | $7.23\times10^{-1}$ | $5.09\times10^{-1}$ | $3.42\times10^{-1}$ | $3.67\times10^{-1}$ | $3.98\times10^{-1}$ | $4.68\times10^{-1}$ | $5.05\times10^{-1}$ | $8.95\times10^{-3}$ |
| $p_{CEU}$ | $5.65\times10^{-1}$ | $4.39\times10^{-2}$ | $3.96\times10^{-1}$ | $1.46\times10^{-1}$ | $4.61\times10^{-1}$ | $2.83\times10^{-2}$ | $5.85\times10^{-1}$ | $4.68\times10^{-1}$ | $8.38\times10^{-1}$ | $4.23\times10^{-1}$ |
| $p_{Vindija}$ | $7.61\times10^{-1}$ | $9.02\times10^{-1}$ | $9.87\times10^{-1}$ | $8.48\times10^{-1}$ | $7.36\times10^{-1}$ | $5.40\times10^{-1}$ | $4.34\times10^{-1}$ | $5.30\times10^{-1}$ | $6.77\times10^{-1}$ | $7.41\times10^{-1}$ |

Table S8: Values of the 25 free parameters for the last ten runs among the twenty accepted. For details about the parameters, see Table S1.1.

| RunID: | 522039 | 544094 | 56031 | 566619 | 573942 | 575635 | 581743 | 761711 | 785396 | 83213 |
|---|---|---|---|---|---|---|---|---|---|---|
| N1 | $1.78\times10^{4}$ | $1.96\times10^{3}$ | $1.63\times10^{3}$ | $6.41\times10^{3}$ | $5.79\times10^{3}$ | $6.98\times10^{3}$ | $3.91\times10^{4}$ | $4.75\times10^{4}$ | $5.85\times10^{3}$ | $3.40\times10^{3}$ |
| N2 | $1.80\times10^{3}$ | $5.98\times10^{3}$ | $5.71\times10^{3}$ | $2.34\times10^{3}$ | $7.36\times10^{3}$ | $3.20\times10^{3}$ | $5.73\times10^{3}$ | $2.07\times10^{3}$ | $2.16\times10^{3}$ | $2.92\times10^{3}$ |
| N4 | $5.54\times10^{1}$ | $1.65\times10^{2}$ | $4.03\times10^{1}$ | $3.57\times10^{2}$ | $4.83\times10^{1}$ | $3.99\times10^{1}$ | $1.14\times10^{1}$ | $3.03\times10^{2}$ | $6.26\times10^{1}$ | $1.11\times10^{1}$ |
| N3 | $2.76\times10^{4}$ | $5.03\times10^{4}$ | $8.19\times10^{3}$ | $1.38\times10^{4}$ | $2.01\times10^{4}$ | $3.44\times10^{4}$ | $7.42\times10^{3}$ | $2.16\times10^{4}$ | $1.75\times10^{4}$ | $2.03\times10^{4}$ |
| m1 | $5.42\times10^{-2}$ | $1.10\times10^{-2}$ | $4.36\times10^{-2}$ | $3.54\times10^{-2}$ | $3.07\times10^{-3}$ | $8.91\times10^{-2}$ | $3.69\times10^{-3}$ | $2.78\times10^{-2}$ | $4.44\times10^{-2}$ | $4.25\times10^{-2}$ |
| m2 | $9.91\times10^{-6}$ | $9.51\times10^{-4}$ | $1.57\times10^{-5}$ | $7.37\times10^{-4}$ | $4.53\times10^{-5}$ | $3.55\times10^{-3}$ | $1.32\times10^{-2}$ | $6.51\times10^{-6}$ | $2.57\times10^{-3}$ | $9.38\times10^{-4}$ |
| m3 | $6.44\times10^{-4}$ | $1.50\times10^{-2}$ | $7.87\times10^{-2}$ | $3.04\times10^{-6}$ | $1.13\times10^{-6}$ | $2.99\times10^{-5}$ | $1.29\times10^{-2}$ | $8.25\times10^{-5}$ | $2.48\times10^{-4}$ | $1.94\times10^{-4}$ |
| m4 | $1.69\times10^{-6}$ | $6.93\times10^{-4}$ | $2.19\times10^{-7}$ | $1.34\times10^{-5}$ | $1.48\times10^{-5}$ | $4.75\times10^{-5}$ | $2.38\times10^{-5}$ | $5.86\times10^{-8}$ | $4.12\times10^{-8}$ | $2.30\times10^{-8}$ |
| m5 | $7.45\times10^{-2}$ | $3.98\times10^{-3}$ | $6.46\times10^{-2}$ | $2.28\times10^{-1}$ | $2.81\times10^{-3}$ | $1.11\times10^{-2}$ | $3.00\times10^{-2}$ | $1.18\times10^{-2}$ | $2.14\times10^{-2}$ | $2.92\times10^{-3}$ |
| m6 | $2.37\times10^{-6}$ | $1.12\times10^{-7}$ | $7.48\times10^{-5}$ | $1.43\times10^{-6}$ | $2.46\times10^{-5}$ | $1.88\times10^{-5}$ | $3.53\times10^{-5}$ | $1.51\times10^{-5}$ | $2.85\times10^{-6}$ | $3.80\times10^{-5}$ |
| T1 | $4.97\times10^{4}$ | $4.32\times10^{4}$ | $6.01\times10^{4}$ | $5.49\times10^{4}$ | $5.93\times10^{4}$ | $4.03\times10^{4}$ | $4.33\times10^{4}$ | $4.09\times10^{4}$ | $4.56\times10^{4}$ | $4.94\times10^{4}$ |
| dT1 | $3.38\times10^{3}$ | $3.00\times10^{1}$ | $1.28\times10^{2}$ | $1.77\times10^{3}$ | $4.49\times10^{1}$ | $1.51\times10^{3}$ | $3.45\times10^{2}$ | $3.49\times10^{1}$ | $2.94\times10^{3}$ | $8.25\times10^{1}$ |
| n3 | $8.88\times10^{0}$ | $6.17\times10^{1}$ | $1.18\times10^{1}$ | $9.62\times10^{0}$ | $7.55\times10^{1}$ | $1.63\times10^{1}$ | $2.07\times10^{0}$ | $4.21\times10^{1}$ | $1.26\times10^{2}$ | $1.67\times10^{2}$ |
| dT2 | $1.83\times10^{1}$ | $1.06\times10^{3}$ | $9.07\times10^{0}$ | $1.79\times10^{0}$ | $1.12\times10^{3}$ | $1.22\times10^{2}$ | $1.29\times10^{1}$ | $1.71\times10^{0}$ | $6.43\times10^{0}$ | $3.44\times10^{2}$ |
| m7 | $2.03\times10^{-5}$ | $1.26\times10^{-2}$ | $6.71\times10^{-5}$ | $1.27\times10^{-5}$ | $1.04\times10^{-2}$ | $8.15\times10^{-4}$ | $2.22\times10^{-5}$ | $1.90\times10^{-5}$ | $2.15\times10^{-4}$ | $5.73\times10^{-5}$ |
| m8 | $2.06\times10^{-3}$ | $5.24\times10^{-2}$ | $1.55\times10^{-2}$ | $1.87\times10^{-3}$ | $1.66\times10^{-2}$ | $3.07\times10^{-3}$ | $2.37\times10^{-3}$ | $1.56\times10^{-3}$ | $1.44\times10^{-3}$ | $2.48\times10^{-3}$ |
| T2 | $2.82\times10^{5}$ | $1.14\times10^{5}$ | $2.91\times10^{5}$ | $2.98\times10^{5}$ | $4.05\times10^{5}$ | $3.05\times10^{5}$ | $2.35\times10^{5}$ | $3.39\times10^{5}$ | $6.21\times10^{5}$ | $2.68\times10^{5}$ |
| m9 | $1.83\times10^{-5}$ | $1.27\times10^{-5}$ | $1.65\times10^{-5}$ | $2.42\times10^{-5}$ | $1.65\times10^{-5}$ | $2.23\times10^{-5}$ | $1.46\times10^{-5}$ | $2.48\times10^{-5}$ | $1.87\times10^{-5}$ | $2.12\times10^{-5}$ |
| m10 | $5.88\times10^{-7}$ | $3.82\times10^{-7}$ | $7.49\times10^{-7}$ | $6.58\times10^{-7}$ | $6.90\times10^{-7}$ | $5.94\times10^{-7}$ | $7.66\times10^{-7}$ | $5.31\times10^{-7}$ | $5.73\times10^{-7}$ | $5.20\times10^{-7}$ |
| T3 | $8.78\times10^{5}$ | $5.88\times10^{5}$ | $5.20\times10^{5}$ | $6.65\times10^{5}$ | $5.74\times10^{5}$ | $5.66\times10^{5}$ | $6.97\times10^{5}$ | $6.60\times10^{5}$ | $6.80\times10^{5}$ | $4.15\times10^{5}$ |
| N5 | $6.74\times10^{2}$ | $9.72\times10^{2}$ | $7.39\times10^{2}$ | $1.24\times10^{3}$ | $1.13\times10^{3}$ | $1.21\times10^{3}$ | $1.24\times10^{3}$ | $1.13\times10^{3}$ | $9.71\times10^{2}$ | $8.56\times10^{2}$ |
| T4 | $5.44\times10^{6}$ | $2.82\times10^{6}$ | $8.90\times10^{6}$ | $2.43\times10^{6}$ | $6.30\times10^{6}$ | $8.36\times10^{6}$ | $5.51\times10^{6}$ | $2.25\times10^{6}$ | $7.48\times10^{6}$ | $5.49\times10^{6}$ |
| $p_{YRI}$ | $5.91\times10^{-1}$ | $7.86\times10^{-1}$ | $6.51\times10^{-1}$ | $1.46\times10^{-1}$ | $9.33\times10^{-1}$ | $2.37\times10^{-1}$ | $8.68\times10^{-1}$ | $1.94\times10^{-3}$ | $4.28\times10^{-1}$ | $2.13\times10^{-1}$ |
| $p_{CEU}$ | $2.53\times10^{-1}$ | $8.70\times10^{-1}$ | $1.40\times10^{-2}$ | $2.99\times10^{-1}$ | $6.80\times10^{-1}$ | $7.70\times10^{-1}$ | $3.44\times10^{-1}$ | $1.48\times10^{-1}$ | $9.95\times10^{-1}$ | $9.47\times10^{-1}$ |
| $p_{Vindija}$ | $1.90\times10^{-1}$ | $3.74\times10^{-1}$ | $9.85\times10^{-1}$ | $9.72\times10^{-1}$ | $9.10\times10^{-1}$ | $2.42\times10^{-1}$ | $4.22\times10^{-1}$ | $2.44\times10^{-1}$ | $7.74\times10^{-1}$ | $6.62\times10^{-2}$ |

### S6 $F_{ST}$ trajectory between metapopulations

We calculated the trajectory of genetic differentiation between and within *Hs* metapopulations $M_A$, $M_B$ and $M_C$ in the run 1014230 of our structured model. To this end, we simulated 10 diploid individuals in each metapopulation at 30 time points (when applicable) from present to 6.5 Mya, spaced on a base-10 logarithmic scale. For computation speed and given that $F_{ST}$ estimates remain stable even for a limited number of markers, we simulated 10 chromosomes of 7 Mbp for each individual, using otherwise the same parameters as for the other general simulations presented in this study. We calculated four independent trajectories:

- **between metapopulations $\mathbf{M_A}$ and $\mathbf{M_B}$** where we calculated the $F_{ST}$ between 10 individuals in the fifth deme (1-based indexing) of $M_A$ and 10 in the fifth deme (1-based indexing) of $M_B$ (i.e. close to the center of the metapopulations).
- **between metapopulations $\mathbf{M_B}$ and $\mathbf{M_C}$** where we calculated the $F_{ST}$ between 10 individuals in the fifth deme (1-based indexing) of $M_B$ and 10 in the fifth deme (1-based indexing) of $M_C$ (i.e. close to the center of the metapopulations).
- **within metapopulation $\mathbf{M_A}$**: we calculated the $F_{ST}$ between the first and the last demes of the $M_A$ metapopulation.
- **within metapopulation $\mathbf{M_B}$**: same as above, but within metapopulation $M_B$ instead.

**Results** The $F_{ST}$ trajectories can be divided into three phases (Figure S6):

1. After the onset of the colonization of the African metapopulation $M_B$ (∼6.5 Mya), genetic differentiation significantly builds up between the two metapopulations $M_A$ and $M_B$. However, $F_{ST}$ remains stable between the extreme demes of each metapopulation. At all times during this period, the differentiation between metapopulations is higher than within, which is expected given that the connectivity between metapopulations is lower than the one within each metapopulation.
2. After the onset of the colonization of the Eurasian metapopulation $M_N$ (which will become Neanderthals), genetic differentiation decreases between $M_A$ and $M_B$ as well as within each metapopulation. There is an increase in connectivity between as well as within metapopulations. At some point during this period, the genetic differentiation within $M_A$ becomes greater than the one between $M_A$ and $M_B$, i.e. the two extreme demes within $M_A$ are genetically more differentiated than the central demes from $M_A$ and $M_B$ are between each other. Connectivity increases between metapopulations, although it decreases within $M_A$.
3. After the onset of the Eurasian sapiens metapopulation $M_C$ (corresponding to the expansion of *Hs* into Eurasia), the $F_{ST}$ values stabilize between as well as within metapopulations. Genetic differentiation between metapopulations is greater than the differentiation within metapopulations, by a factor of 100 compared to $M_A$ and 2 compared to $M_B$. The differentiation between $M_C$ and $M_B$ is ∼6-times greater than in-between $M_A$ and $M_B$.

Across the whole considered period (from present 6.5 Mya), $F_{ST}$ between $M_A$ and $M_B$ ranges between 0.03—0.89. It ranges between 0—0.03 within $M_A$ and between 0.01—0.37 within $M_B$.

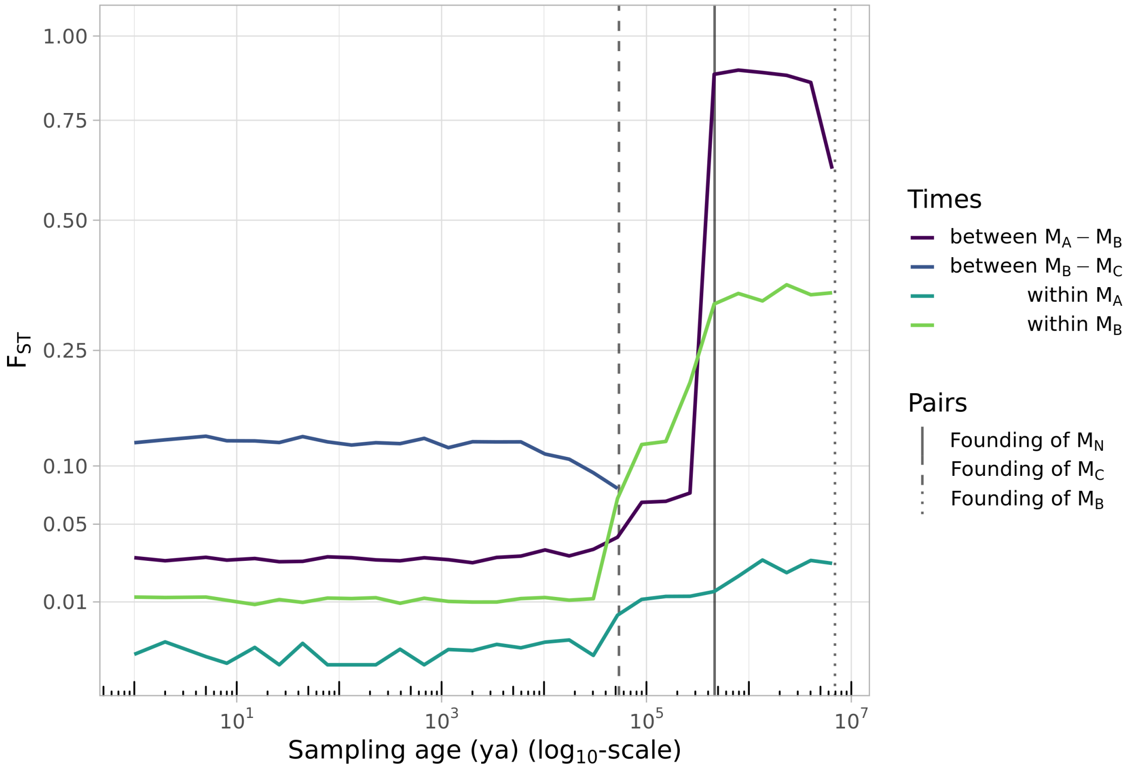

Figure S5: Evolution of the $F_{ST}$ for three sampling strategies, from present up to 6.5 Mya. Note that the $x$-axis is in $\log_{10}$-scale and the $y$-axis is in square-root scale.

789 # S7 Single-sample ancestry-*LD* decay constants for aDNA

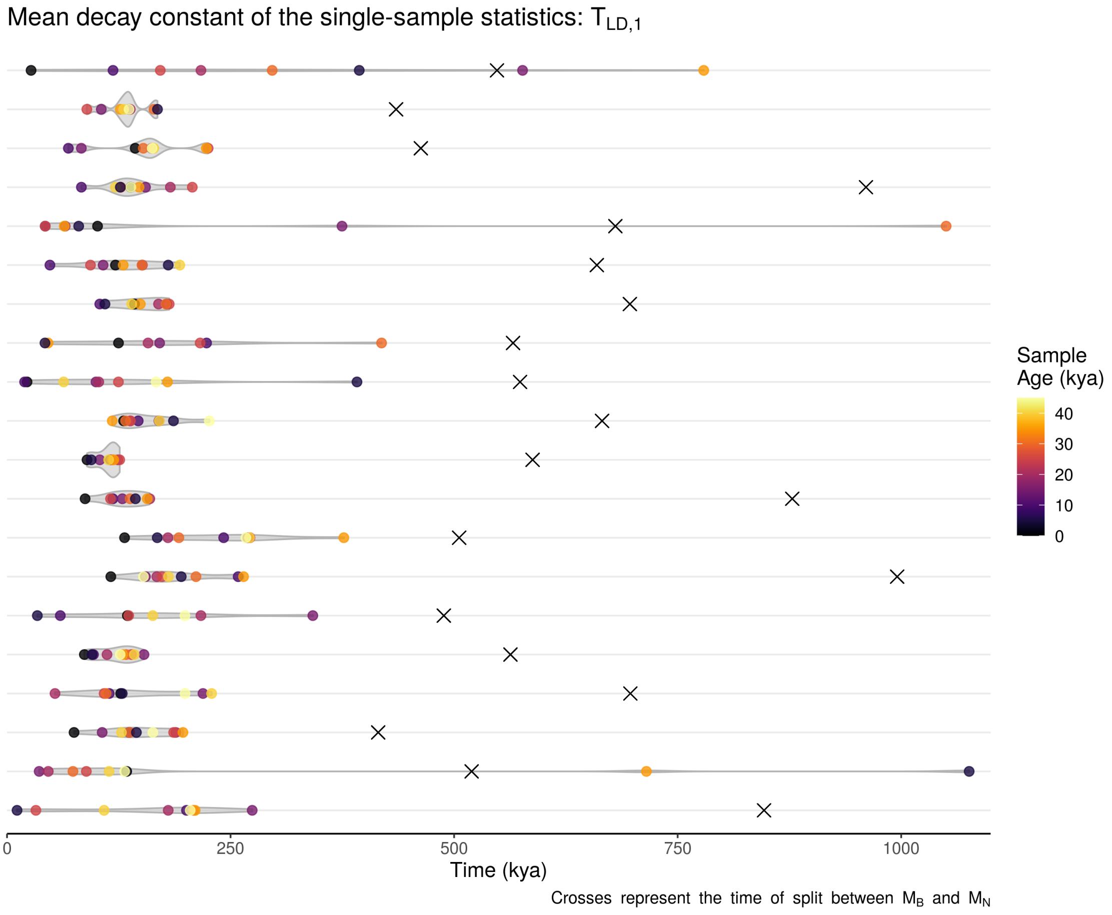

Figure S6: Distribution of the $T_{LD,1}$ estimates (in kya, colored dots, with coloring proportional to the age of the simulated samples) obtained for a time series of ancient Eurasian samples simulated under the twenty selected runs (one per row) of our structured model. For details on the simulations and statistical analyses, see Notes S1.8. The simulated *Hn-Hs* split ages (i.e. founding of $M_N$ from $M_B$) are represented, for each run, with a cross. Note that the $T_{LD,1}$ estimates assume a generation time of 25 years and include the simulated sample ages.

### S8 Published models

We considered eleven demographic models from the human evolution literature, which assume (i) introgression from Neanderthal into *Hs* and (ii) a tree-like model of human evolution with no or very limited intra-continental population structure. For all these models, we simulated genetic data for 50 CEU (or a proxy of CEU for models fitted on other European populations) (sampled at present), 50 YRI (or a proxy) (sampled at present) and one Neanderthal (sampled 50,000 years BP, corresponding to Vindija33.19), using `msprime 1.1.1` with a binary mutation model (Baumdicker et al., 2022). We simulated genomes of twenty chromosomes (30 Mb each), i.e. 600 Mb per individual in total (~20% of the human genome size), varying the mutation rates between $5.0 \cdot 10^{-9}$ and $5.0 \cdot 10^{-8}$ per bp per generation, to improve the ability of models to explain the empirical genetic diversity. For instance, Ragsdale and Gravel (2019) model relied on the recombination map rather than the mutation clock. If we had to use the currently accepted mutation rate, their model would predict $\pi_{CEU}$ on the order of 0.26 $kb^{-1}$, which is three times lower than the observed.

For all simulations, the recombination rate was fixed as $\rho = 10^{-8}$ $b^{-1}g^{-1}$ (uniform along the genome). We simulated each model using the generation time specified in the original study, to preserve the specific scaling of all the time parameters. Consequently, given that the sampling times of ancient genomes are provided in our simulations in years BP, the scaling of the sampling in generations BP could vary slightly across the models. Note however that for model comparison and result plotting, the timing of the *PSMC* curves and the ancestry-*LD* decay constants ($T_{LD}$) were scaled in absolute years BP assuming a fixed generation time of 25 years. This common calibration was chosen to allow the comparison of common temporal statistics across all models. Also, the time and population size scaling of the *PSMC* curves assumed the mutation rate used in the simulation, so that the scaling of $\theta$ remained consistent with the genetic diversity that was simulated for each model and mutation rate combination.

Note than in the following plots—where we represent the statistical estimates over multiple simulations for the same model—, we selected the twenty runs that were the closest to the observed data, following the same procedure as described in Notes S1.4.

#### S8.1 Ragsdale et al. (2019)

Ragsdale et al. (2019) observed that several widely used models in human evolution failed to reproduce LD patterns in present-day genomes. Using a two-locus statistic with a likelihood-based inference, their best model fit involved archaic admixture events in ancestral African, Asian and European populations, with an unknown lineage admixing with African populations before and after the split between Africans and Eurasians.

Specifically, their model generalizes the standard model published in Gutenkunst et al. (2009) by introducing two deeply divergent hominid branches (Fig. S7). One branch (Neanderthal) admixed with the Eurasian ancestors of CEU and CHB and another one (unidentified archaic African) admixed with the ancestors of YRI before as well as after the YRI-CEU split. The admixture between archaic and modern human branches were modelled using continuous and symmetric gene flow. We used the model represented in Fig. 4A and Table 1 of Ragsdale and Gravel (2019) in which the two deeply divergent branches split independently from the one leading to modern humans, ∼500 kya. The putative Neanderthal branch was estimated to contribute 1.2% ± 0.6% ancestry in CEU and CHB. The authors assumed a generation time of 29 years.

Practically, we retrieved the Ragsdale et al. (2019) model from `stdpopsim` 0.1.3b1, model `"OutOfAfricaArchaicAdmixture_5R19"`.

**Simulation results** The original study did not use nor specify a mutation rate, since the authors calibrated their absolute event ages and population sizes using the recombination clock solely (African-American genetic map from Hinch et al. (2011)). When simulating their model, we found that a currently accepted mutation rate in humans ($1.2 \times 10^{-8}$) led to a large underestimation of the nucleotide diversities in both CEU and YRI (Fig. S9-A). This observation remained true for any mutation rate below $2.5 \times 10^{-8}$ (Fig. S8-A). A better fit of $\pi_{CEU}$ and $\pi_{YRI}$ could only be obtained for $\mu$ ranging from $3 \times 10^{-8}$ to $4 \times 10^{-8}$, which are significantly higher than currently accepted rates. The Ragsdale et al. model predicts fairly well the scaled $AFS$ in CEU and YRI but overestimates the proportion of singleton SNPs. The model accurately predicts the observed $F_{ST}$ between CEU and YRI (Fig. S8-F, $y$-axis). The $PSMC_{CEU}$ trajectory departs significantly from the empirical curve $\phi_o$, with a single peak instead of the two bumps observed in $\phi_o$. The model peak occurs ∼100 kya (∼300 kya earlier than $\phi_o$), i.e. between the simulated YRI-CEU split and the start of the archaic-African admixture. In periods older than the peak, the estimated $PSMC$ stabilizes at a much lower value than estimated on the observed data (Fig. S8-D).

Regarding the admixture-sensitive statistics, the $DCFS$ has a L-shape but strongly overestimates the proportion of singleton ascertained SNPs by ∼30%. The $D$ statistic is also generally overestimated (Fig. S8-F, $x$-axis). The $S'$-inferred putatively archaic tracts in the model explain well the observed match rate and average length. The mean lengths of the tracts inferred with the $CRF$ methods are in good agreement with the observed but the introgression rate $\alpha_{CRF}$ is significantly underestimated by ∼2.2-fold on average. Lastly, the ancestry-$LD$ curves suggest an admixture event ∼19 kya (assuming $g = 25$), significantly earlier than reported in previous works. This very recent decay constant is however in line with the end of the period of migration from Neanderthal into CEU that is simulated in the original scenario.

Overall, if we exclude the introgression rates predicted with $CRF$, the Ragsdale et al. (2019) model tends to overestimate the signatures of genetic similarity between CEU and Neanderthals compared to YRI and Neanderthals. Besides, it does not accurately explain the patterns of within-population genetic diversity ($PSMC_{CEU}$, $\pi$) at the whole-genome scale for mutation rates in line with the ones commonly accepted.

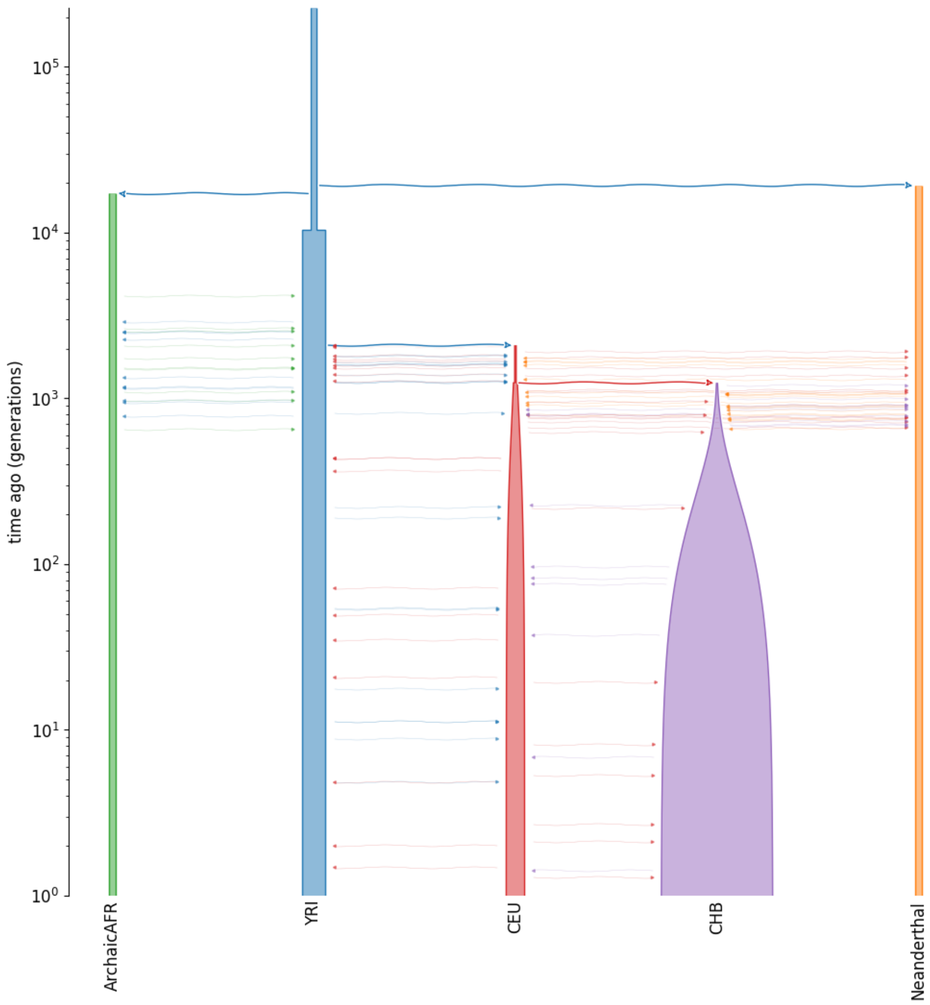

Figure S7: Visual representation of the scenario from Ragsdale et al. (2019) obtained using `demesdraw`. The $y$-axis is in generations BP and $\log_{10}$-scaled.

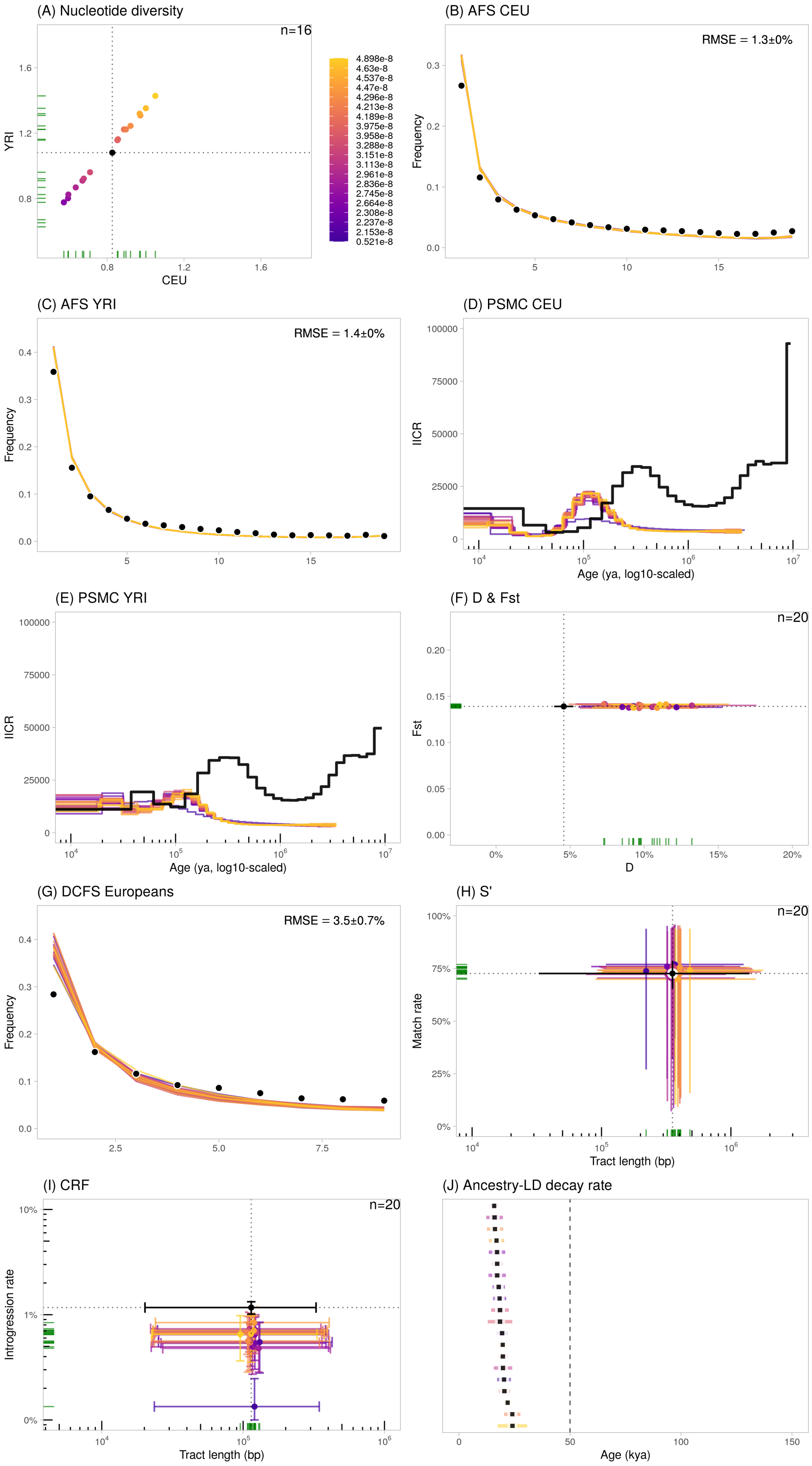

Figure S8: Statistics calculated using the Ragsdale et al. (2019) scenario, using varying mutation rates ($5 \times 10^{-9}$ to $5 \times 10^{-8}$ b$^{-1}$g$^{-1}$). We plotted the 20 runs closest to the observed data. The error bars represent the confidence intervals at 95% ($= 1.96 \cdot SE$) except for $S'$ (H) and *CRF* (I) where they represent the 2.5- and 97.5%-percentiles of the distributions.

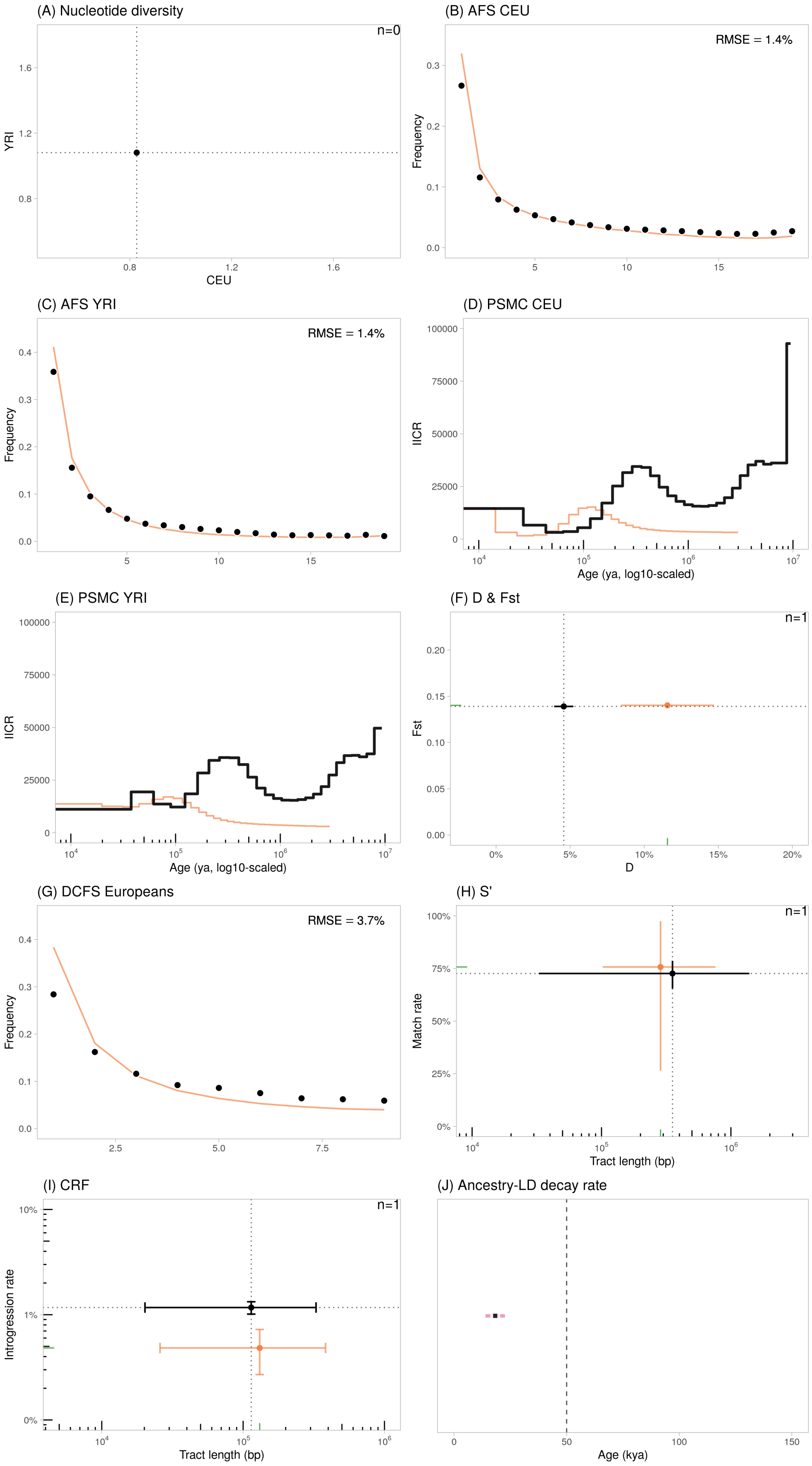

Figure S9: Statistics calculated using the Ragsdale et al. (2019) scenario with the standard mutation rate of $1.2 \times 10^{-8}$ b$^{-1}$g$^{-1}$. The error bars represent the confidence intervals at 95% ($= 1.96 \cdot SE$) except for $S'$ (H) and $CRF$ (I) where they represent the 2.5- and 97.5%-percentiles of the distributions. In (A), the observed $\pi$ values are outside of the figure limits.

#### S8.2 Durvasula et al. (2020)

Durvasula et al. (2020) observed that current models of human evolution failed to explain genetic statistics like conditional site frequency spectra (CSFS). They showed that accounting for a contribution from archaic hominins into the genomes of present-day Africans increased the fit of the model to the observed CSFS. Using the CSFS statistics—which do not require reference archaic genomes—and an ABC-based inference framework, they compared six major demographic models. These models are generalization of a base model published in Prüfer et al. (2017) which included an admixture event from modern humans into Neanderthals (5%) and admixture from Neanderthals into the ancestors of non-Africans (3%). However, these new models differ among themselves in the topology of the introgression event from an unknown archaic lineage into the ancestors of modern humans and the split time of the archaic lineage. We simulated data under Model C.1 which had the best fit to the observed CSFS (Table S4, Table S8 in Durvasula and Sankararaman (2020)). This model with four populations includes an admixture event (with rate $\alpha$=5%) from an unknown archaic lineage into the ancestors of modern humans soon before the YRI-CEU split. The unknown archaic lineage split was estimated to occur before the Neanderthal-Sapiens split (Fig. S10). The authors used a generation time of 29 years.

Practically, we used the `ms` command provided by the authors in their Supplementary Material p. 16, where populations are ordered as: CEU, YRI, Neanderthal, Denisovan:

```
ms 202 1 -t 480 -r 519.99948 1000000 -I 4 100 100 1 1 -en 0 3 0.1 -en 0 4 0.1
-es 0.04310345 1 0.97 -ej 0.05387931 5 3 -ej 0.05387931 2 1
-es 0.055 1 alpha -es 0.2155172 4 0.94 -en 0.2155172 7 0.2
-es 0.2836207 3 0.95 -ej 0.2836207 8 1 -ej 0.3577586 4 3
-ej 0.4741379 3 1 -ej 0.6 6 1 -ej 0.7844828 7 1
```

Listing 1: Command `ms` of model C.1 from Durvasula et al. SM, p. 16.

We modified the sampling to have 50 CEU diploids, 50 YRI diploids, one Neanderthal diploid sampled at 50 kya (corresponding to Vindija33.19). The `ms` command was converted into a `demes` graph object using `demes::from_ms()` with $N_o = 10^4$ (as specified by the authors) and later into a `msprime` demography object using `msprime::from_deme()`.

**Simulation results** Mutation rates below $1.2 \times 10^{-8}$ lead to underestimate the nucleotide diversities in both CEU and YRI (< 0.6 kb$^{-1}$) (Fig. S11-A). The best fit for $\pi_{YRI}$ would be obtained for $\mu \sim 2.5 \times 10^{-8}$. The model slightly overestimates low-frequency variants in CEU while it strongly underestimates the proportion of singleton SNPs in YRI (Fig. S11-B,C). As a consequence, the model also underestimates the $F_{ST}$ between CEU and YRI by 30% on average (Fig. S11-F, $y$-axis). The $PSMC_{CEU}$ trajectory departs significantly from the empirical curve, with the absence of any visible oscillation. The estimated effective population size is constant for all the times considered, with $IICR \sim 10k$, in line with the simulated population size (10k) and lack of population structure.

Regarding the admixture-sensitive statistics, the $DCFS$ has a L-shape but strongly overestimates the proportion of singleton ascertained SNPs by more than ~45% (Fig. S11-G). The $D$ statistic is also strongly overestimated, ranging from 15% to 20% depending on the mutation rate (Fig. S11-F, $x$-axis). The $S'$-inferred putatively archaic tracts have match rates to the simulated Neanderthal genome lower than empirical data (Fig. S11-H, $y$-axis), with a bias positively correlated to the mutation rate. Unlike Ragsdale and Gravel (2019), the Durvasula et al. (2020) model slightly underestimates the mean length of the observed $CRF$ tracts and slightly overestimates the introgression rate (Fig. S11-I). The ancestry-$LD$ curves have decay constants at ~41 kya (assuming $g = 25$), overall concordant with the Neanderthal admixture age that was simulated (43 kya assuming $g = 25$).

The Durvasula et al. (2020) model thus tends to overestimate the signatures of genetic similarity between CEU and Neanderthals compared to YRI and Neanderthals. Also, it poorly predicts the patterns of within-population genetic diversity ($PSMC_{CEU}$, $AFS$ YRI) at the whole-genome scale.

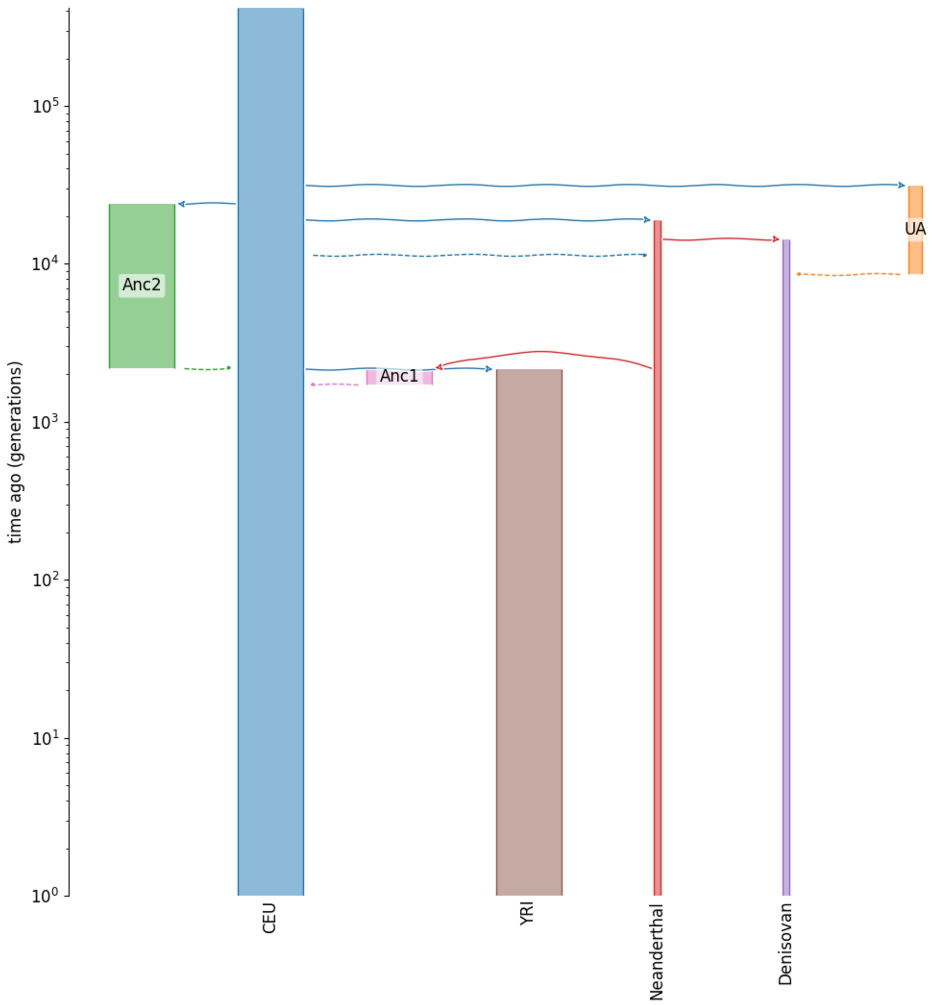

Figure S10: Visual representation of the scenario from Durvasula et al. (2020) . The $y$-axis is in generations BP and $\log_{10}$-scaled.

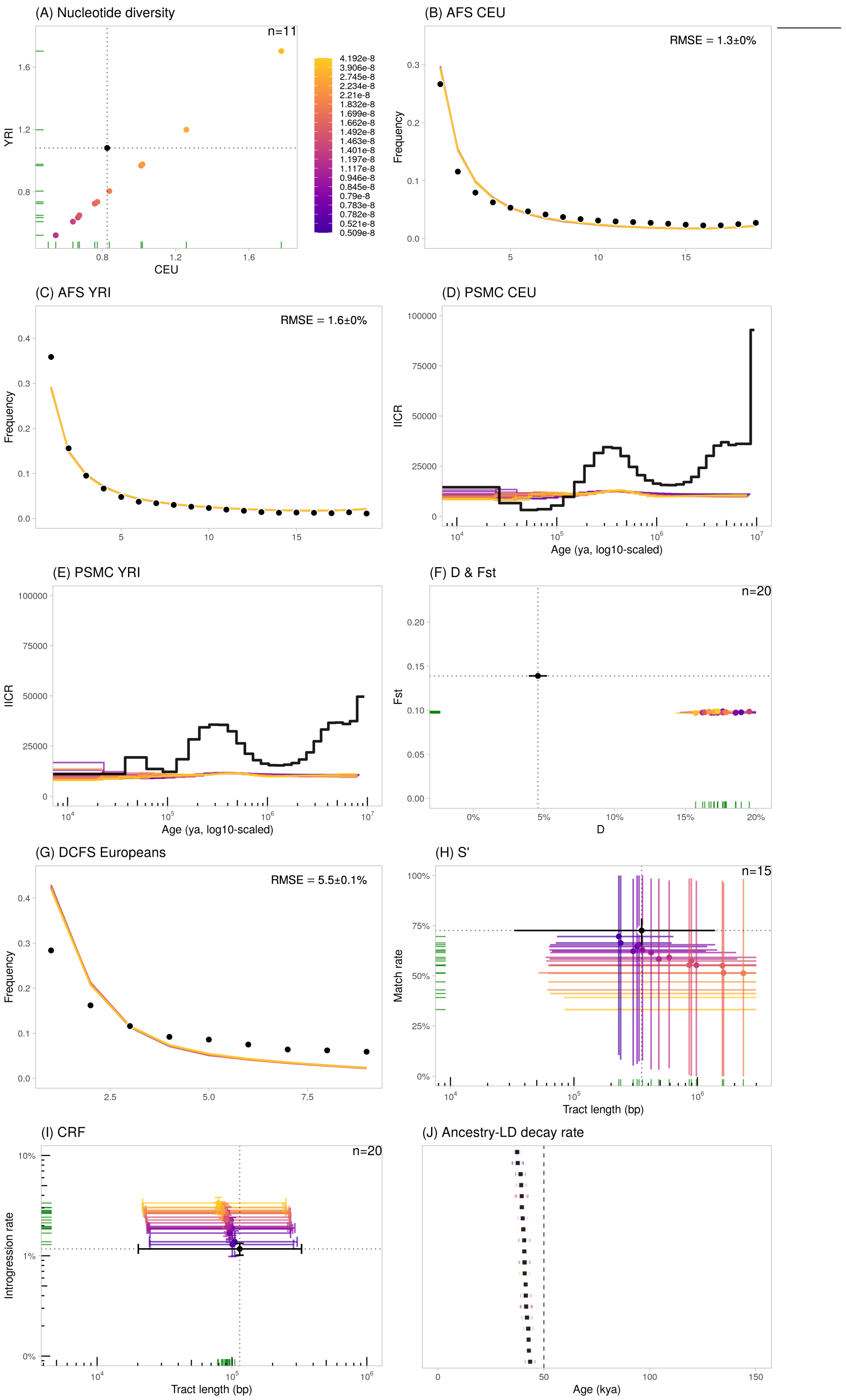

Figure S11: Statistics calculated using the Durvasula et al. (2020) scenario, using varying mutation rates ($5 \times 10^{-9}$ to $5 \times 10^{-8}$ b$^{-1}$g$^{-1}$). We plotted the 20 runs closest to the observed data. The error bars represent the confidence intervals at 95% ($= 1.96 \cdot SE$) except for $S'$ (H) and $CRF$ (I) where they represent the 2.5- and 97.5%-percentiles of the distributions.

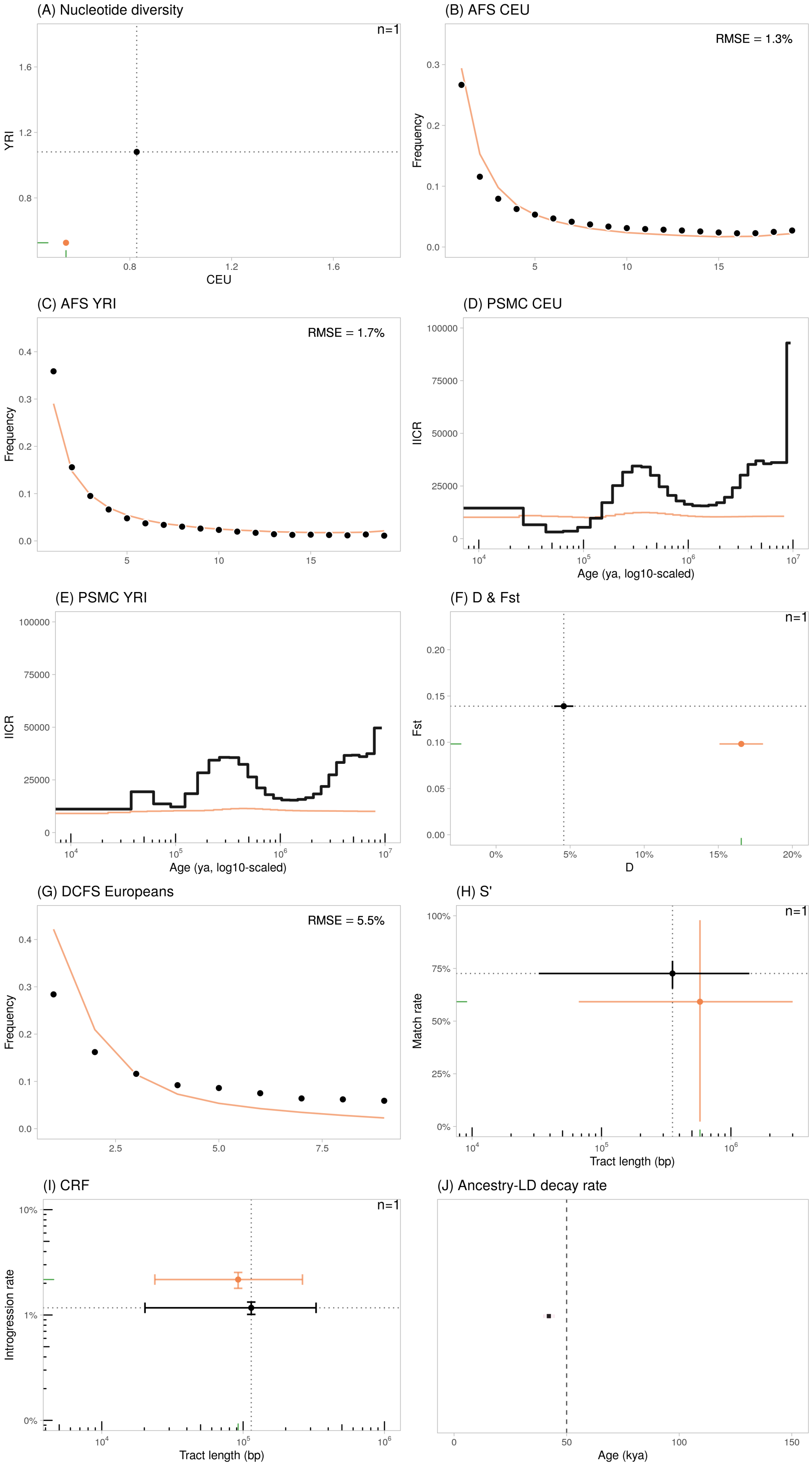

Figure S12: Statistics calculated using the Durvasula et al. (2020) scenario with the standard mutation rate of $1.2 \times 10^{-8}$ b$^{-1}$g$^{-1}$. The error bars represent the confidence intervals at 95% ($= 1.96 \cdot SE$) except for $S'$ (H) and $CRF$ (I) where they represent the 2.5- and 97.5%-percentiles of the distributions.

#### S8.3 Jacobs et al. (2019)

Jacobs et al. (2019) investigated the evolutionary history of archaic and modern humans with a particular focus on several modern Papuan populations and archaic Denisovan populations. Using mismatch distributions on the ancestry blocks in Papuans as a summary statistics, they extended an archaic demographic model published in Malaspinas et al. (2016) to assess the divergence time between the Denisovan-related components of ancestry in Papuan genomes.

Their model includes an ancient *Hs* population splitting into a branch that ultimately leads to present-day Africans (YRI) and to an unidentified non-African ancestral lineage (Fig. S13). The latter lineage founds a Papuan lineage, later followed by a Eurasian lineage. West Eurasians (CEU) and East Asians (CHB) split 1,293 generations ago. Multiple Neanderthal pulses are modelled, into (i) the unidentified non-African ancestral lineage; (ii) the ancient Eurasians before the CEU-CHB split; (iii) the Papuan lineage; (iv) the East Asian lineage. Additionally, two pulses from divergent Denisovans are modelled into ancient Papuans. Compared to the Malaspinas et al. (2016) model, this one thus implements two pulses of Denisova introgression. The authors used a generation time of 29 years.

Practically, we retrieved the Jacobs et al. (2019) model from `stdpopsim` 0.1.3b1, model: `"PapuansOutOfAfrica_10J19"`.

**Simulation results** Estimates for genetic diversity in CEU and YRI tend to be significantly larger than the observed ($\pi > 1.5$ kb$^{-1}$) for most mutation rates, including for $\mu = 1.2 \times 10^{-8}$ ($\pi_{CEU} = 1.5$ kb$^{-1}$, i.e. 1.8-times larger) (Fig. S14-A). The model explains relatively well the scaled $AFS$ in both CEU and YRI although it tends to underestimate the singleton SNP proportion in both populations (Fig. S14-B,C). The model also overestimates other low-frequency variants in CEU (Fig. S14-B). The model strongly underestimates the $F_{ST}$ between CEU and YRI by 66% on average (Fig. S14-F, $y$-axis). The $PSMC_{CEU}$ curve reproduces fairly well the empirical one for the time period between 10 and 500 kya, although it is shifted towards the recent past with a lag of ∼250 ky. Indeed, the first bump reaches its maximum at ∼150 kya and ends (forward in time) at ∼60 kya (corresponding roughly to the formation of the CEU branch, 51 kya). Earlier than 500 kya, the $PSMC_{CEU}$ curve is flat, with $IICR \sim 35k$. The $PSMC_{YRI}$ lacks clear oscillations (Fig. S14-E).

Regarding the admixture-sensitive statistics, the $DCFS$ has a near-linear shape which would be interpreted as a signal of ancestral population structure without archaic admixture. The $D$-statistic is underestimated by ∼50% on average (Fig. S14-F, $x$-axis). The $S'$-inferred putatively archaic tracts have a match rate nearly half that of the observed value, with mean lengths showing a variable bias depending on the mutation rate. The putative archaic tracts identified with $CRF$ strongly underestimate both the mean lengths of the tracts as well as the introgression rate (with rates $\alpha_{CRF}$ ranging from 0.03% to 0.5%). The bias is negatively correlated with the mutation rate. The ancestry-$LD$ curves have decay constants at the furthest 49 kya (assuming $g = 25$).

Overall, when considering altogether the $DCFS$, $D$-statistic, $S'$ and $CRF$ tracts, the Jacobs et al. (2019) model tends to underestimate the signatures of genetic similarity between (CEU, Neanderthals) compared to (YRI, Neanderthals) and does not explain several observed patterns of genetic diversity. For instance, it underestimates by ∼3-fold the genetic differentiation between present-day CEU and YRI.

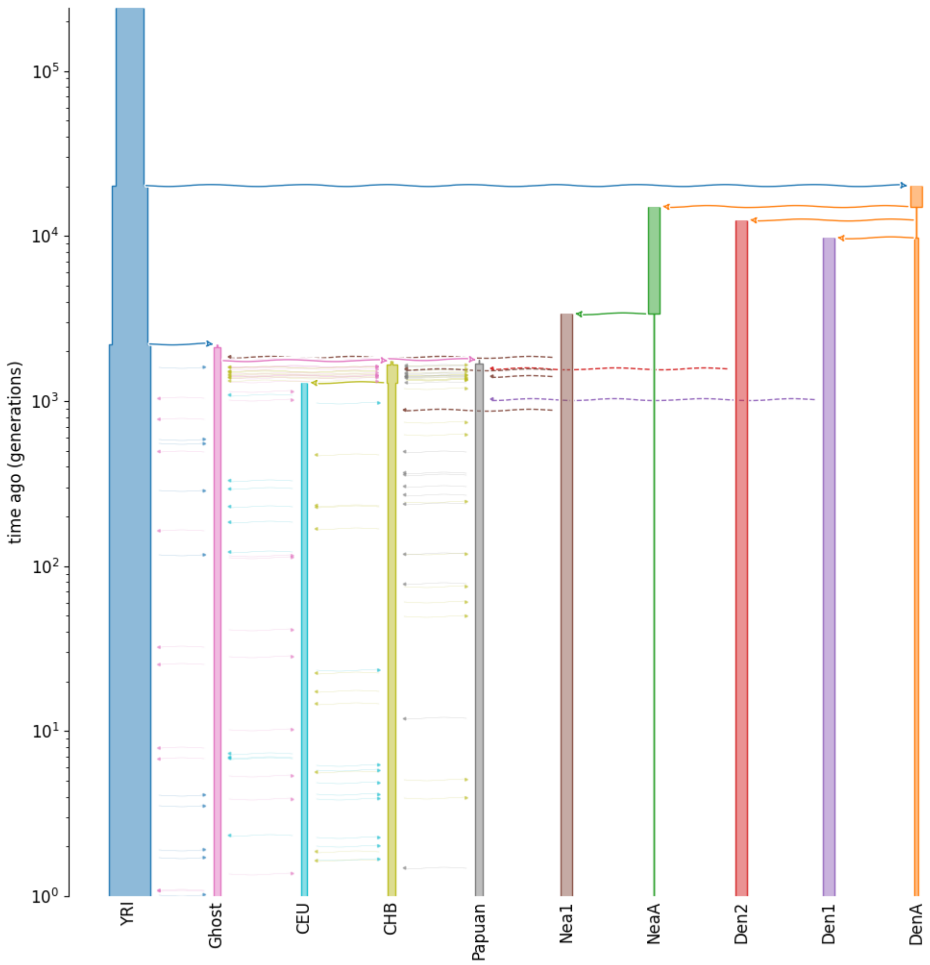

Figure S13: Visual representation of the scenario from Jacobs et al. (2019) . The $y$-axis is in generations BP and $\log_{10}$-scaled.

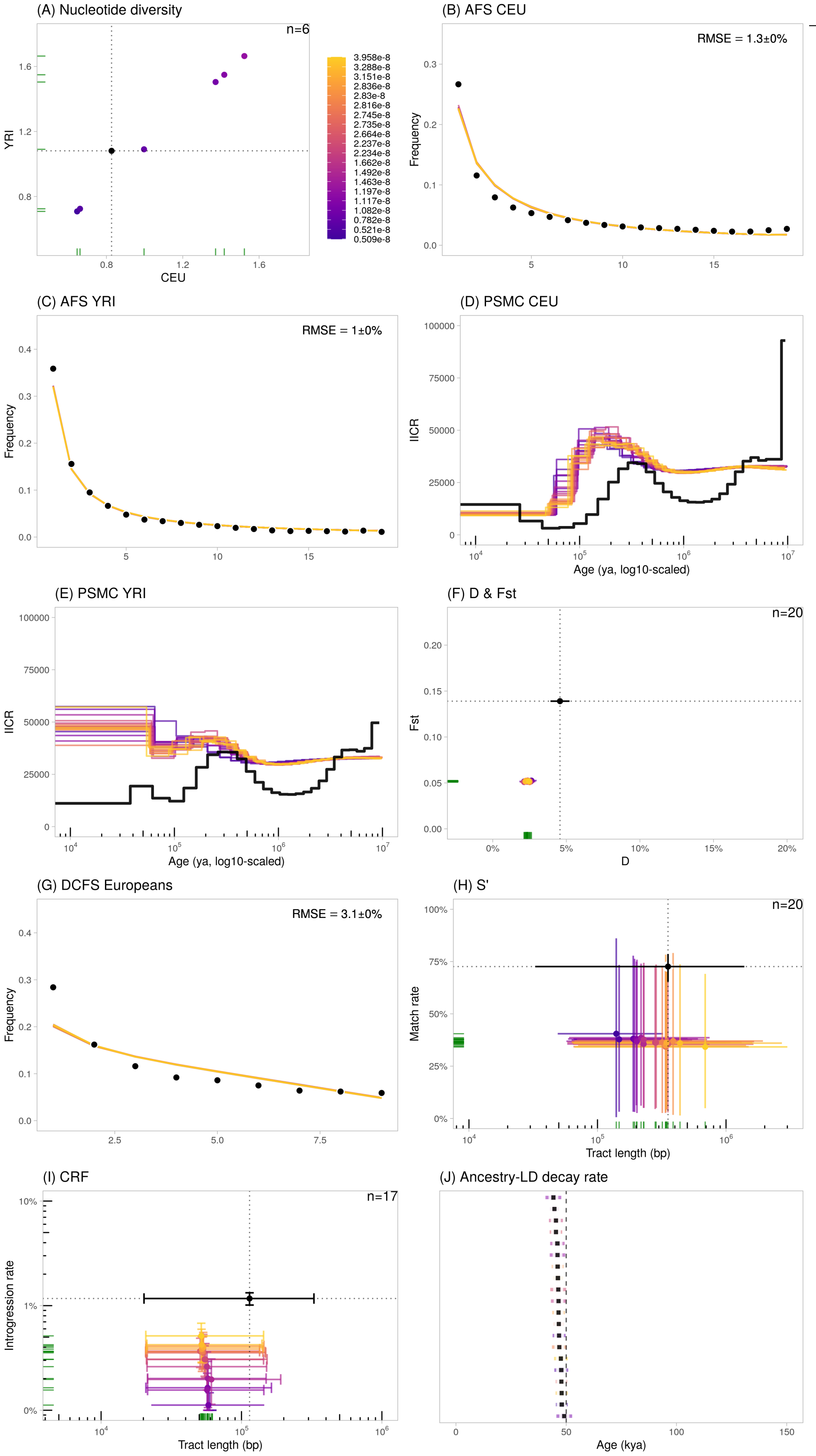

Figure S14: Statistics calculated using the Jacobs et al. (2019) scenario, using varying mutation rates ($5 \times 10^{-9}$ to $5 \times 10^{-8}$ $b^{-1}g^{-1}$). We plotted the 20 runs closest to the observed data. The error bars represent the confidence intervals at 95% ($= 1.96 \cdot SE$) except for $S'$ (H) and $CRF$ (I) where they represent the 2.5- and 97.5%-percentiles of the distributions.

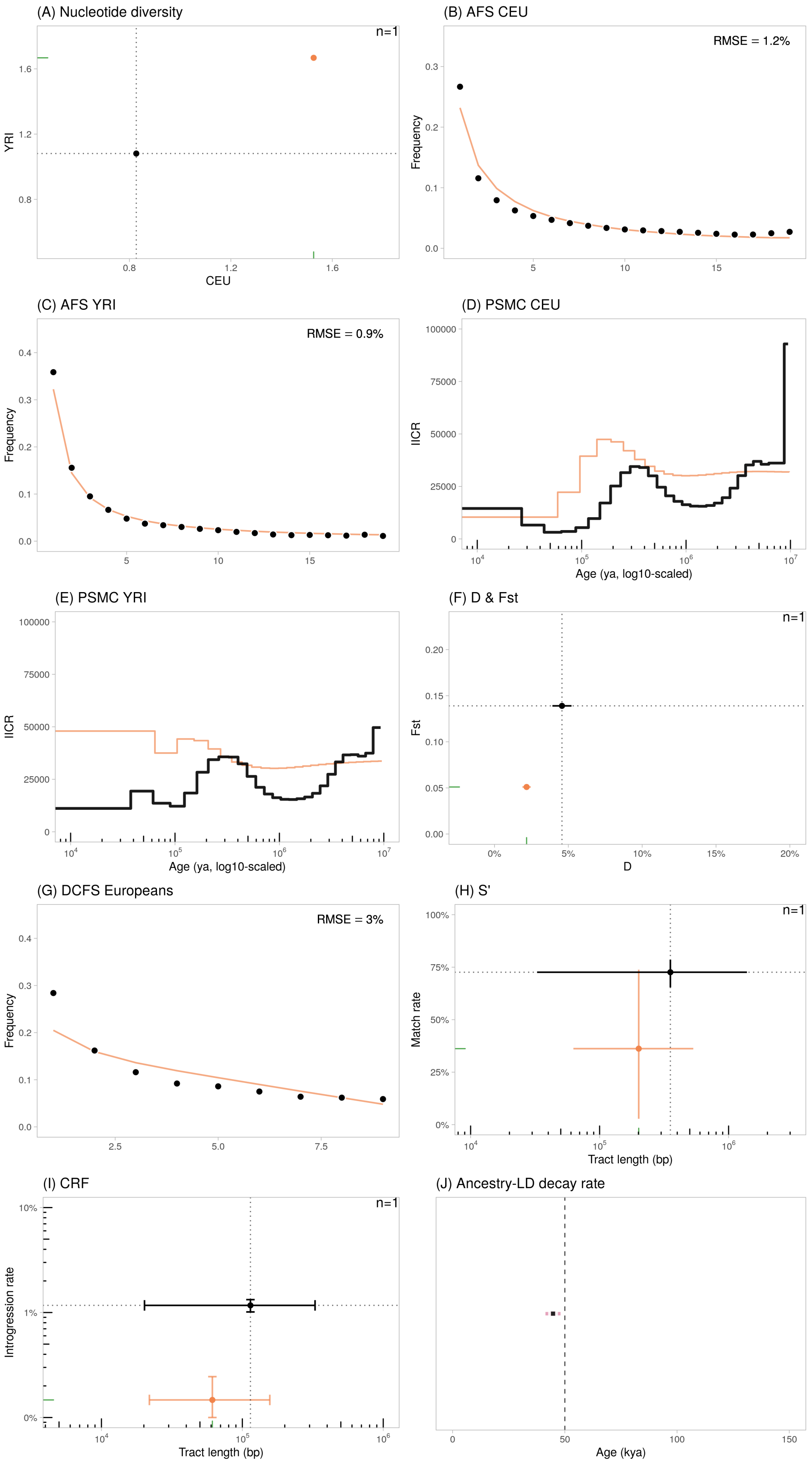

Figure S15: Statistics calculated using the Jacobs et al. (2019) scenario with the standard mutation rate of $1.2 \times 10^{-8}$ b$^{-1}$g$^{-1}$. The error bars represent the confidence intervals at 95% ($= 1.96 \cdot SE$) except for $S'$ (H) and $CRF$ (I) where they represent the 2.5- and 97.5%-percentiles of the distributions.

#### S8.4 Kamm et al. (2019)

Kamm et al. (2019) introduced `momi2`, an inferential method which uses the $AFS$ over multiple populations to infer the demographic history while accounting for multiple pulses of gene flow between populations. Using this approach with eight ancient and modern sampled human populations, they tested the presence of "Basal Eurasian" ancestry in present-day Europeans. This hypothesis stems from the inference of an ancestry component in early European farmers that was identified as outgroup to all non-African populations, while retaining similar intensity of drift. In several populations, the Basal Eurasian ancestry was found to be negatively correlated with Neanderthal ancestry, suggesting that Basal Eurasians were a branch that off-shooted before any admixture from Neanderthal. Later on, by admixing with Neanderthal-admixed non-African populations, Basal Eurasians would dilute the signal of Neanderthal ancestry in later non-Africans.

Specifically, their best likelihood model (Fig. S16) integrates some form of population structure, although with limited gene flow between demes. In this scenario, modern humans split with Neanderthals ~700 kya. Then, within modern humans, Africans (here, Mbuti) split with non-Africans around 100 kya. The earliest split with non-Africans occurred soon after, between Basal Eurasians and a lineage that would later lead to (i) four ancient lineages: Ust-'Ishim, with no known ancestry in present-day humans; Loschbour, representative of Mesolithic Western Hunter-Gatherers; LBK, representative of Neolithic early European farmers; MA1, Paleolithic humans from Mal'ta (Siberia) and (ii) two modern populations: Han and Sardinians. We used here Mbuti and Sardinians, respectively, to represent an African and a European population. In this model, Sardinians have a signature of double admixture from Loschbour (3.2%) and earlier from Basal Eurasians (9.4%). The single pulse of Neanderthal admixture was estimated to occur at 56.8 kya, with rate 3.0%. The authors used a generation time of 25 years.

Practically, we retrieved the Kamm et al. (2019) model from `stdpopsim 0.1.3b1`, model `"AncientEurasia_9K19"`.

**Simulation results** The best fit for the nucleotide diversities in CEU and YRI is obtained using a mutation rate of $\sim 1.2 \times 10^{-8}$, in line with the authors estimating a best-fit mutation rate of $1.22 \times 10^{-8}$ in their study. The model explains well the scaled $AFS$ in both CEU and YRI although it underestimates the singleton SNP proportion in both populations (Fig. S17-B,C). This could be explained by the absence of a recent expansion in their model. The $F_{ST}$ between CEU and YRI in the simulated data is significantly higher than the observed by 67% on average (Fig. S17-F, $y$-axis). However, note that the authors inferred their model using Sardinian and Mbuti samples whereas our observed estimate of $F_{ST}$ were obtained for CEU and YRI, respectively. Therefore, inconsistency between numbers can be driven by the variable differentiation across populations. The $PSMC_{CEU}$ curve reproduces very well the empirical one for the time period between 10 kya and 500 kya, and is among the best fit observed across all compared published models. However, earlier than 500 kya, the model predicts a flat trajectory, with $IICR \sim 20k$. We note also that $PSMC_{YRI}$ shows one bump, which is driven by changes in population sizes (cf. Figure S16) and not by population structure or changes in connectivity.

Regarding the admixture-sensitive statistics, the $DCFS$ shows a very weak L-shape and the proportion of singleton ascertained SNPs is underestimated by ~20%. The $D$-statistic is overestimated by 62% on average (Fig. S17-F, $x$-axis). The $S'$-inferred putatively archaic tracts have a match rate which decreases as the mutation rate increases. The best estimate is obtained for $\mu \sim 0.7 \times 10^{-8}$, which is much lower than most commonly reported mutation rates in humans (Fig. S17-H, $x$-axis). Regarding the $CRF$-inferred tracts, the introgression rates tend to be overestimated with a bias correlated to the mutation rate (Fig. S17-I, $y$-axis. The mean lengths are slightly underestimating the empirical value by 27% on average. The ancestry-$LD$ curves have decay constants ranging at the furthest 57 kya (assuming $g = 25$), concordant with the estimated Neanderthal pulse of admixture 56.8 kya.

Overall, the Kamm et al. (2019) model tends to explain several admixture statistics well, except the $DCFS$. Yet, the good fit obtained for $S'$ and $CRF$ parameters are obtained for mutation rates that are much lower than currently accepted rates in humans (Fig. S18). We stress again that the comparison here is limited by the fact that we consider Sardinians and Mbuti (the populations used in the original study) instead of the CEU and YRI populations from which the empirical estimates we consider were obtained.

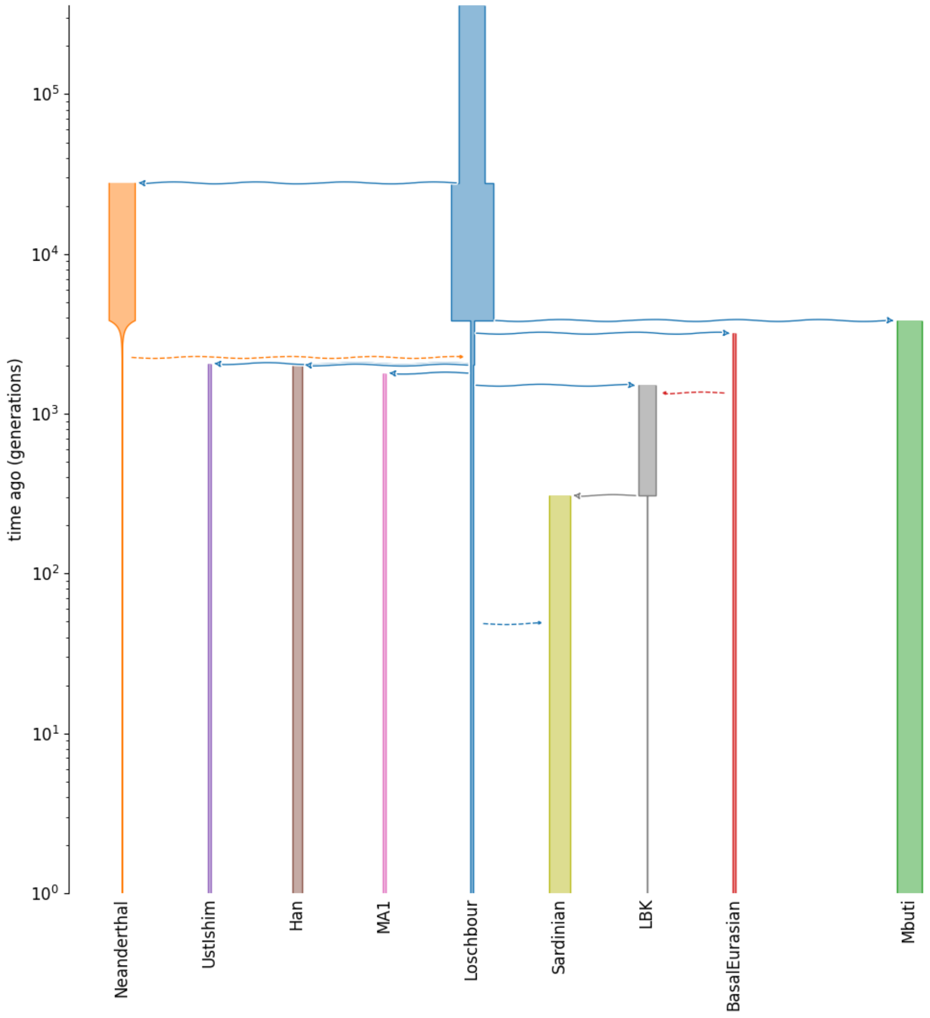

Figure S16: Visual representation of the scenario from Kamm et al. (2019) . The $y$-axis is in generations BP and $\log_{10}$-scaled.

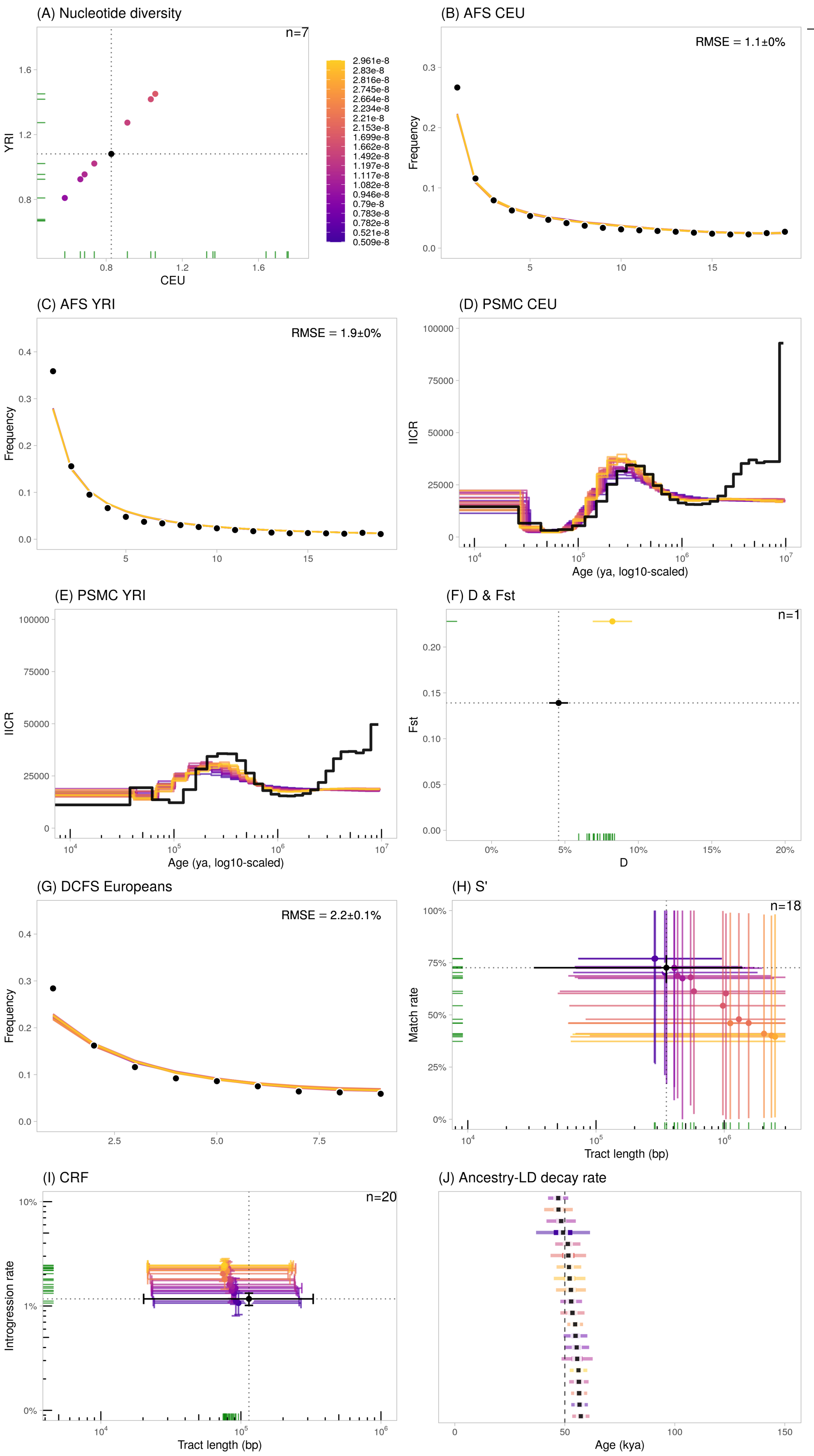

Figure S17: Statistics calculated using the Kamm et al. (2019) scenario, using varying mutation rates ($5 \times 10^{-9}$ to $5 \times 10^{-8}$ b$^{-1}$g$^{-1}$). We plotted the 20 runs closest to the observed data. The error bars represent the confidence intervals at 95% ($= 1.96 \cdot SE$) except for $S'$ (H) and $CRF$ (I) where they represent the 2.5- and 97.5%-percentiles of the distributions.

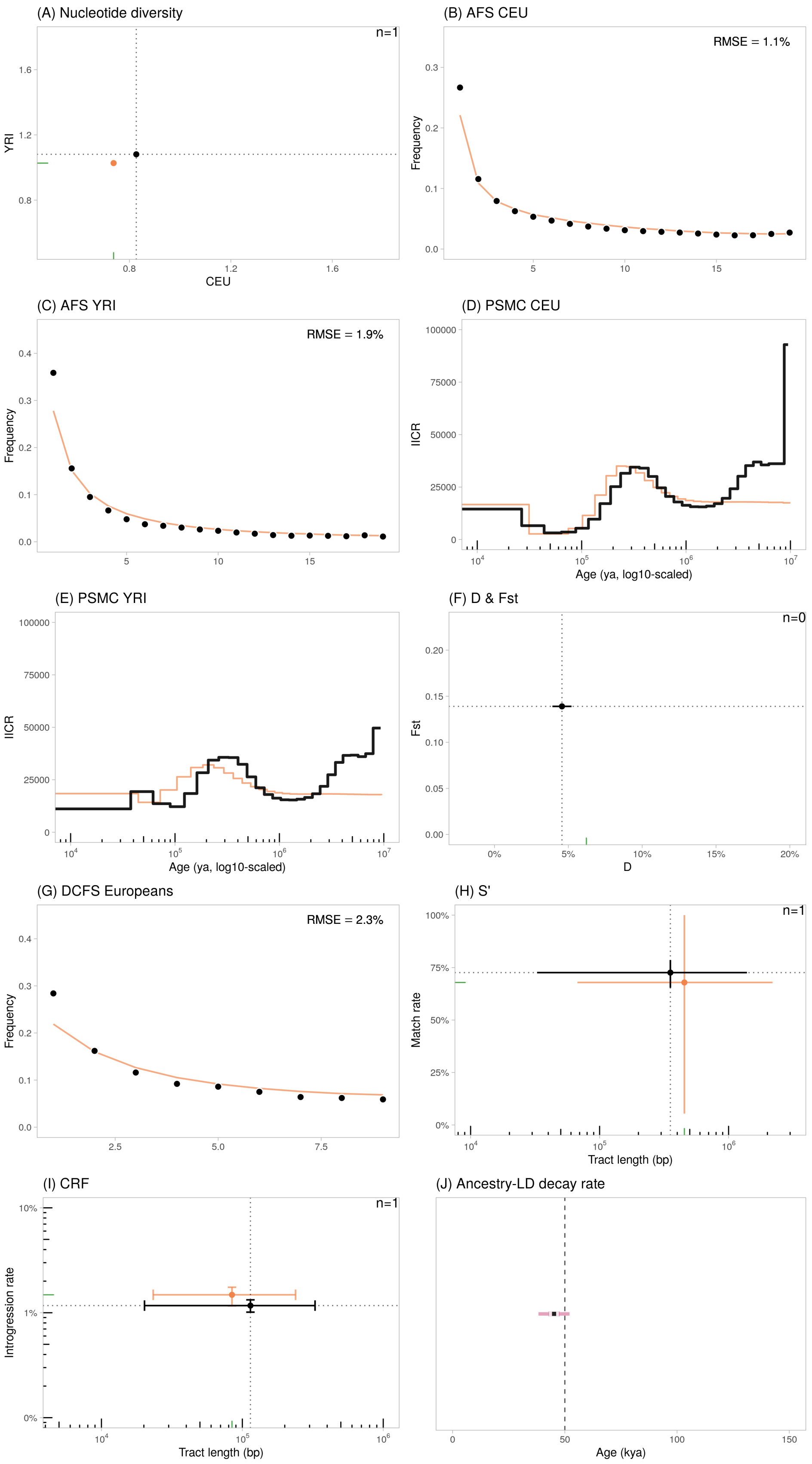

Figure S18: Statistics calculated using the Kamm et al. (2019) scenario with the standard mutation rate of $1.2 \times 10^{-8}$ $b^{-1}g^{-1}$. The error bars represent the confidence intervals at 95% ($= 1.96 \cdot SE$) except for $S'$ (H) and $CRF$ (I) where they represent the 2.5- and 97.5%-percentiles of the distributions. In the (F) panel, the dot is outside of the figure limits (towards the top).

#### S8.5 Iasi et al. (2021)

Iasi et al. (2021) investigated the dynamics of Neanderthal admixture and developed an "extended admixture pulse model" allowing the estimation of the timing and duration of the gene flow between Neanderthal and modern humans, using patterns of ancestry LD. Their inference suggested that gene flow from Neanderthal could have extended over thousands of years. We retrieved the simple demographic model that the authors used for performance evaluation, which assumes constant population sizes.

In this model, after its split with the (ancestral) "YRI" population, the (ancestral) "CEU" population receives continuous (unidirectional) gene flow from Neanderthal for a period stretching over 800 generations (i.e. from 50 kya to 30 kya), for a cumulated migration rate over the whole period equal to 0.029. The authors assume a generation time of 25 years.

Practically, we retrieved the Iasi et al. (2021) model from `stdpopsim 0.1.3b1`, model: `"OutOfAfricaExtendedNeandertalAdmixturePulse_3I21"`.

**Simulation results** Realistic prediction of the nucleotide diversities in CEU and YRI is obtained for mutation rates significantly larger ($> 1.8 \times 10^{-8}$) than the currently accepted rate. Using $\mu = 1.2 \times 10^{-8}$, diversities are significantly underestimated (~0.49 kb$^{-1}$ in both populations). The model tends to slightly overestimate the frequency of singletons in CEU (with overestimation of other low-frequency variants) and more significantly underestimate the frequency of singletons in YRI (Fig. S20-B,C). The $F_{ST}$ between CEU and YRI in the simulated data slightly underestimates the observed value by 20% on average (Fig. S20-F, $y$-axis). The $PSMC_{CEU}$ curve is flat ($IICR \sim 20k$) and strongly departs from the empirical trajectory.

Regarding the admixture-sensitive statistics, the $DCFS$ shows a very weak L-shape and the proportion of singleton ascertained SNPs is underestimated by ~20%. The $D$-statistic is rather well estimated (Fig. S20-F, $x$-axis). The $S'$-inferred putatively archaic tracts have a match rate about half the empirical value. Some mutation rates provide realistic tract lengths (Fig. S20-H, $x$-axis), including the standard rate of $1.2 \times 10^{-8}$. Regarding the $CRF$-inferred tracts, except for a few cases, the introgression rates tend to be significantly lower than the observed (Fig. S20-I, $y$-axis) (with a bias negatively correlated with $\mu$). Same is observed for the mean tract lengths (Fig. S20-I, $x$-axis). The ancestry-$LD$ decay constants range from 35 kya to 41 kya (assuming $g = 25$).

Overall, the Iasi et al. (2021) model tends to underestimate the values of several admixture statistics like the $DCFS$ and the $S'$ statistics. We note that the model fails to reproduce the $PSMC$ trajectory in CEU and YRI, which is expected since this model is constant-size with no population structure nor changes in connectivity among populations.

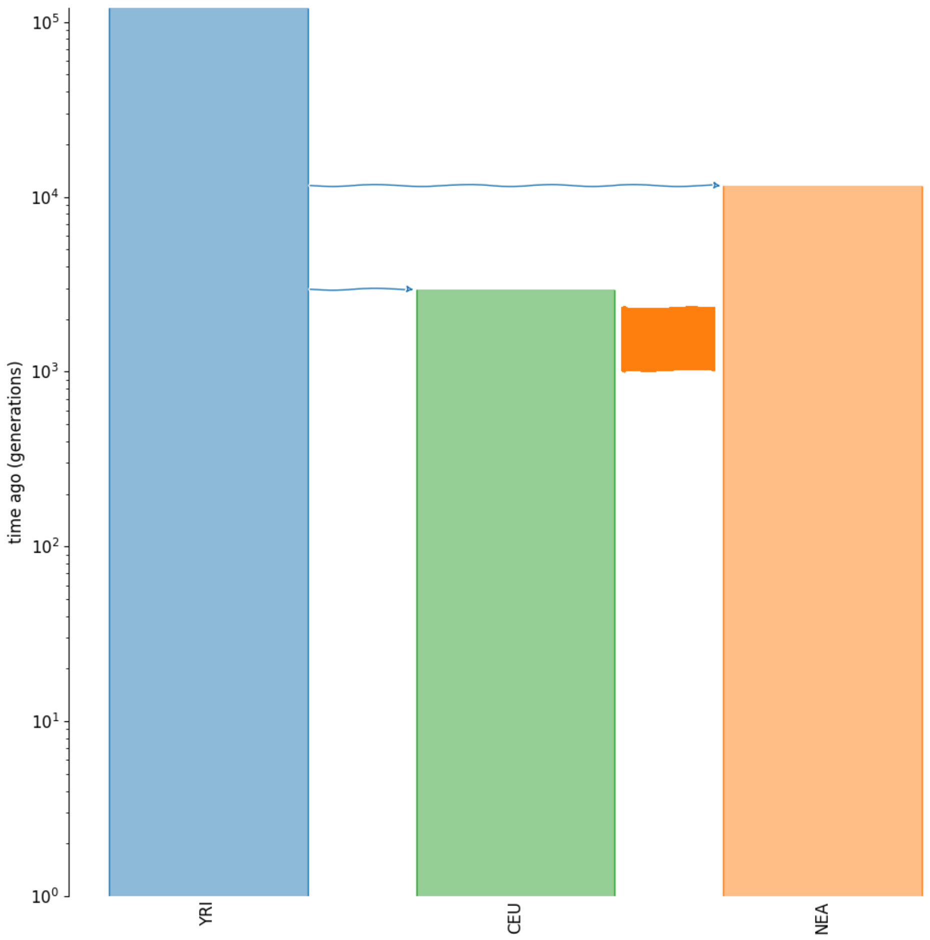

Figure S19: Visual representation of the scenario from Iasi et al. (2021) . The $y$-axis is in generations BP and $\log_{10}$-scaled.

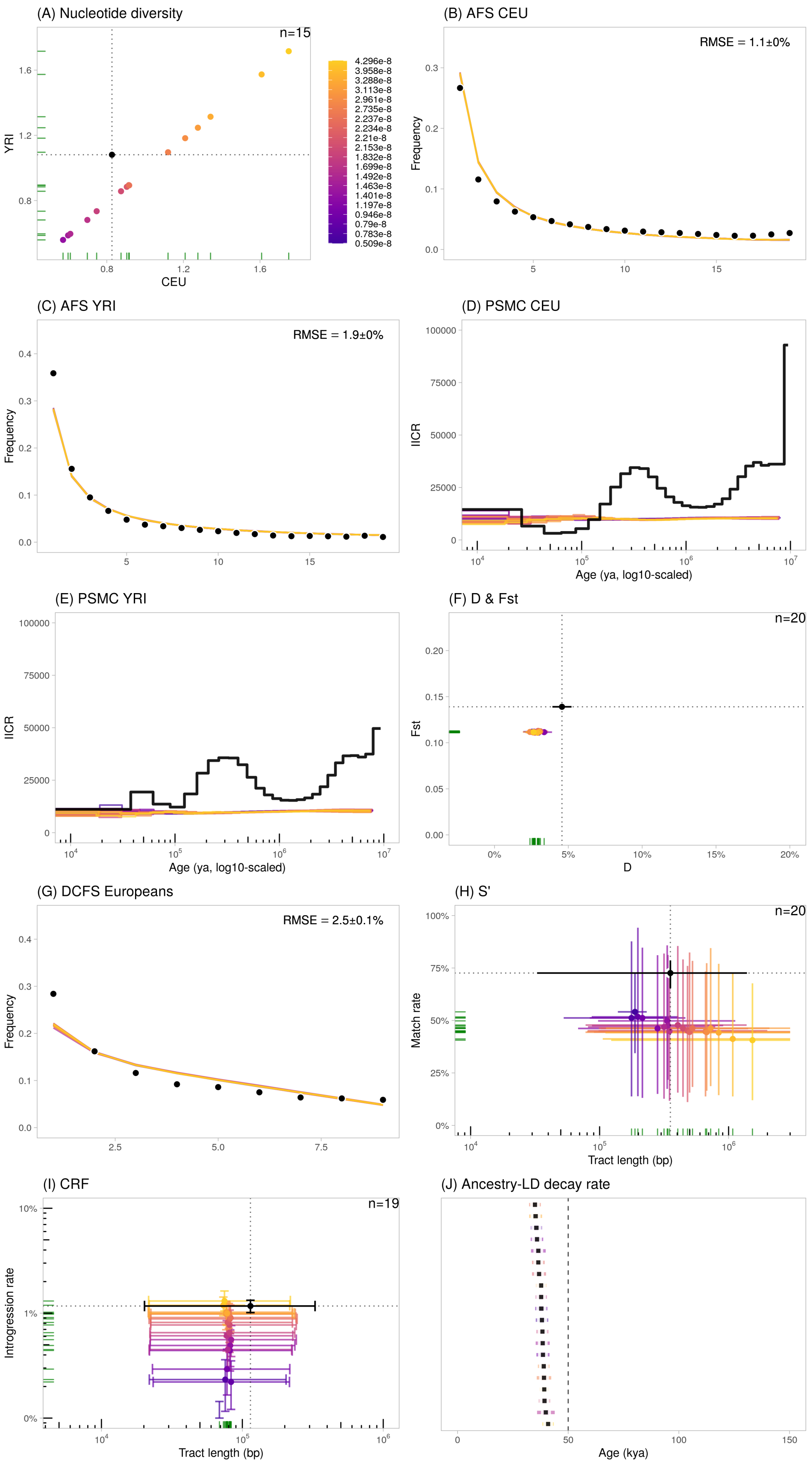

Figure S20: Statistics calculated using the Iasi et al. (2021) scenario, using varying mutation rates ($5 \times 10^{-9}$ to $5 \times 10^{-8}$ b$^{-1}$g$^{-1}$). We plotted the 20 runs closest to the observed data. The error bars represent the confidence intervals at 95% ($= 1.96 \cdot SE$) except for $S'$ (H) and $CRF$ (I) where they represent the 2.5- and 97.5%-percentiles of the distributions.

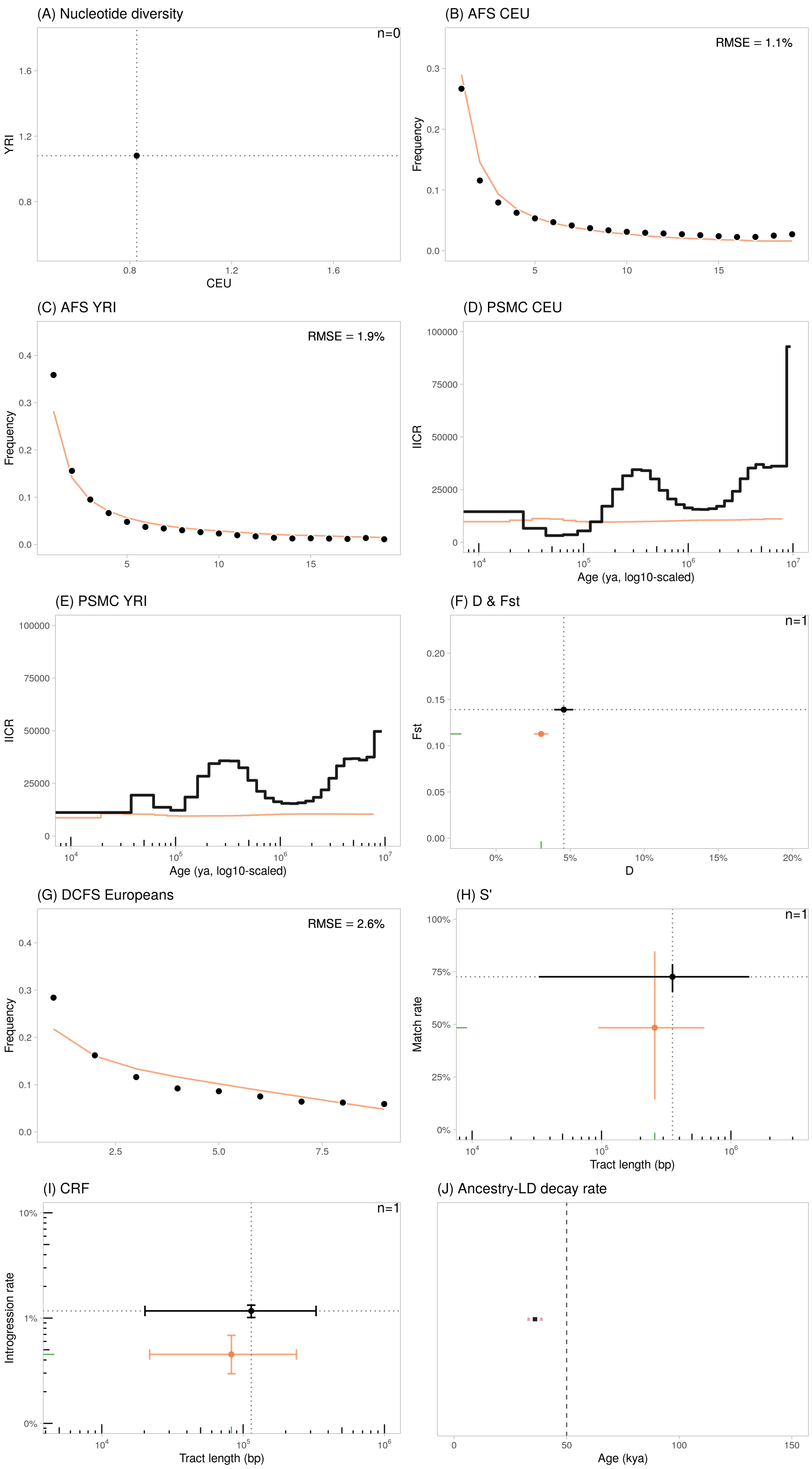

Figure S21: Statistics calculated using the Iasi et al. (2021) scenario with the standard mutation rate of $1.2 \times 10^{-8}$ b$^{-1}$g$^{-1}$. The error bars represent the confidence intervals at 95% ($= 1.96 \cdot SE$) except for $S'$ (H) and $CRF$ (I) where they represent the 2.5- and 97.5%-percentiles of the distributions.

#### S8.6 Gower et al. (2021)

Gower et al. (2021) implemented convolutional neural networks (CNN) to detect signatures of adaptive introgression. Their machine learning approach requires training data, that they simulated under a demographic model with archaic admixture, under neutral as well as selective sweep models. The authors built a composite demographic model ("Model A1") with three populations (CEU, YRI and Neanderthals), using topology and parameter values drawn from three publications (Appendix 3, Table 1 from Gower et al. (2021)):

- Kuhlwilm et al. (2016) for the effective size of Neanderthals, YRI and the population ancestral to CEU and YRI ("Anc" in Fig. S22),
- Ragsdale and Gravel (2019) for parameters related to the recent expansion history of CEU,
- Prüfer et al. (2017) for the parameters related to the split time of Neanderthal and all parameters related to the gene flow from Neanderthal into CEU.

Specifically, the model represents an ancestral population that founds two isolated populations: CEU and YRI, with CEU experiencing an exponential growth following its founding. At 1,896 generations BP, CEU receives a 2.25% migration pulse from the Neanderthal population. The authors used a generation time of 29 years. Practically, we simulated the Model A1 using the YAML file provided by the authors in a GitHub repository[6].

**Simulation results** For most mutation rates, genetic diversity in CEU tends to be underestimated, which is notable at $\mu = 1.2 \cdot 10^{-8}$ with $\pi_{CEU} \sim 0.50$ kb$^{-1}$, although such bias is not observed in YRI (Fig. S23-A). Although the $AFS$ in YRI is rather well predicted (despite an underestimation of the frequency of singletons), the model poorly predicts the $AFS$ in CEU, with a very significant underestimation of singleton SNPs and low-frequency variants (Fig. S23-B,C). This poor fit on the European diversity further impacts the differentiation index, with all $F_{ST}$ significantly larger than the observed value by 1.8-fold on average (Fig. S23-F, $y$-axis). The $PSMC_{CEU}$ curve shows a single bump, ∼350 ky earlier than the corresponding bump on the empirical trajectory. Earlier, it stabilizes with $IICR \sim 20k$ without further oscillations.

Regarding the admixture-sensitive statistics, the $DCFS$ shows a linear shape, and the proportion of singleton ascertained SNPs is very significantly underestimated (Fig. S23-G). The $D$-statistic is rather well estimated (Fig. S23-F, $x$-axis, cf. green ticks). The $S'$-inferred archaic tracts tend to be on average as long as the empirical tracts (Fig. S23-H, $x$-axis), but the match rates are slightly overestimated by 11% on average (Fig. S23-H, $y$-axis). Regarding the $CRF$-inferred tracts, the mean length is accurately predicted although the introgression rates are slightly overestimated (Fig. S23-I). The ancestry-$LD$ curves have decay constants around 44 kya (assuming $g = 25$).

Overall, the Gower et al. (2021) model strongly underestimates the "admixture" signal as quantified through the $DCFS$ shape, and fails to accurately predict the distribution of genome-wide diversity in the CEU ($AFS$), leading to a biased estimation of the differentiation between CEU and YRI.

[6] https://github.com/grahamgower/genomatnn/blob/main/demographic_models/HomininComposite_4G20.yaml

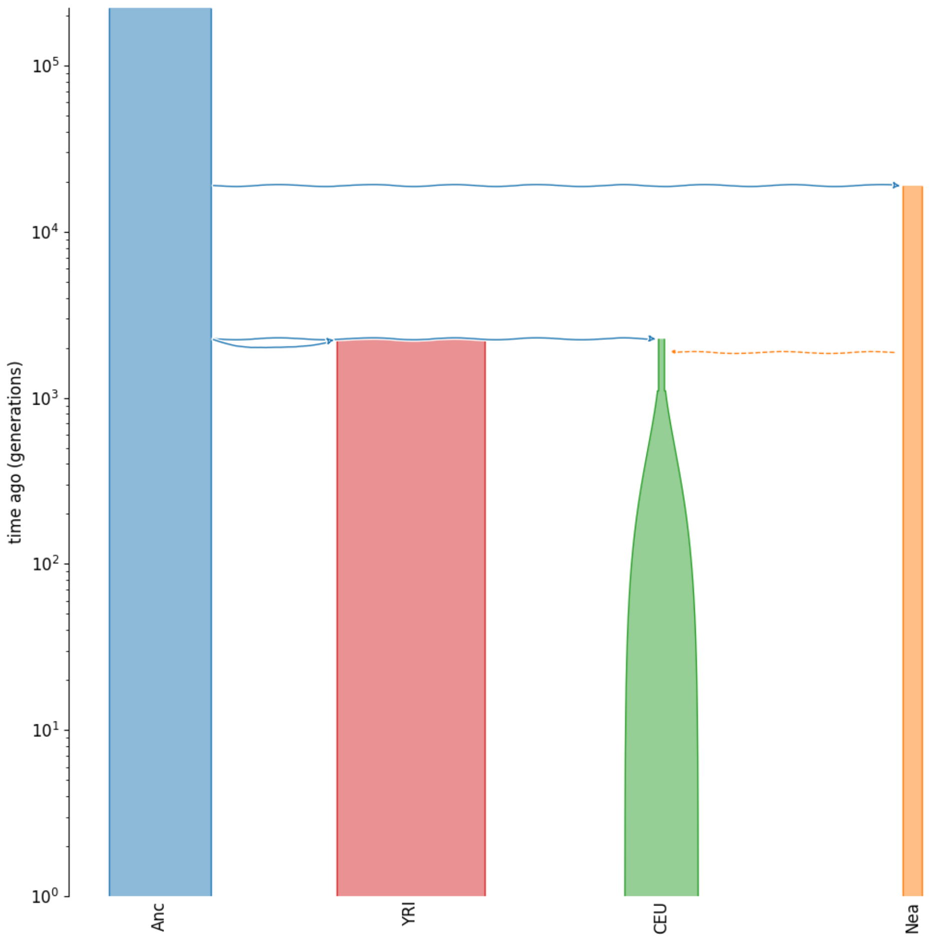

Figure S22: Visual representation of the scenario from Gower et al. (2021) . The $y$-axis is in generations BP and $\log_{10}$-scaled.

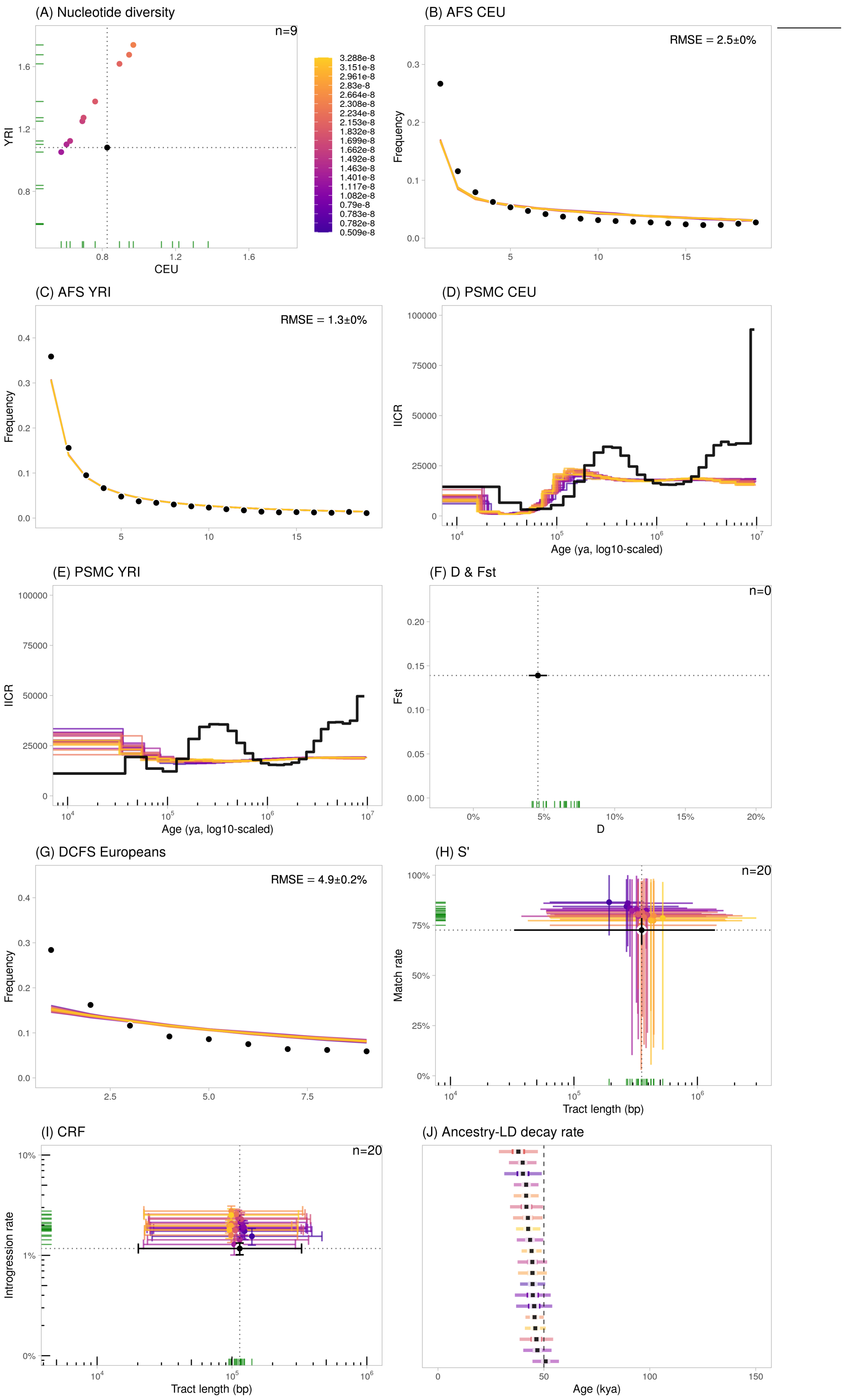

Figure S23: Statistics calculated using the Gower et al. (2021) scenario, using varying mutation rates ($5 \times 10^{-9}$ to $5 \times 10^{-8}$ b$^{-1}$g$^{-1}$). We plotted the 20 runs closest to the observed data. The error bars represent the confidence intervals at 95% ($= 1.96 \cdot SE$) except for $S'$ (H) and $CRF$ (I) where they represent the 2.5- and 97.5%-percentiles of the distributions. Note that in (F), the dots are outside of the figure limits, towards the top.

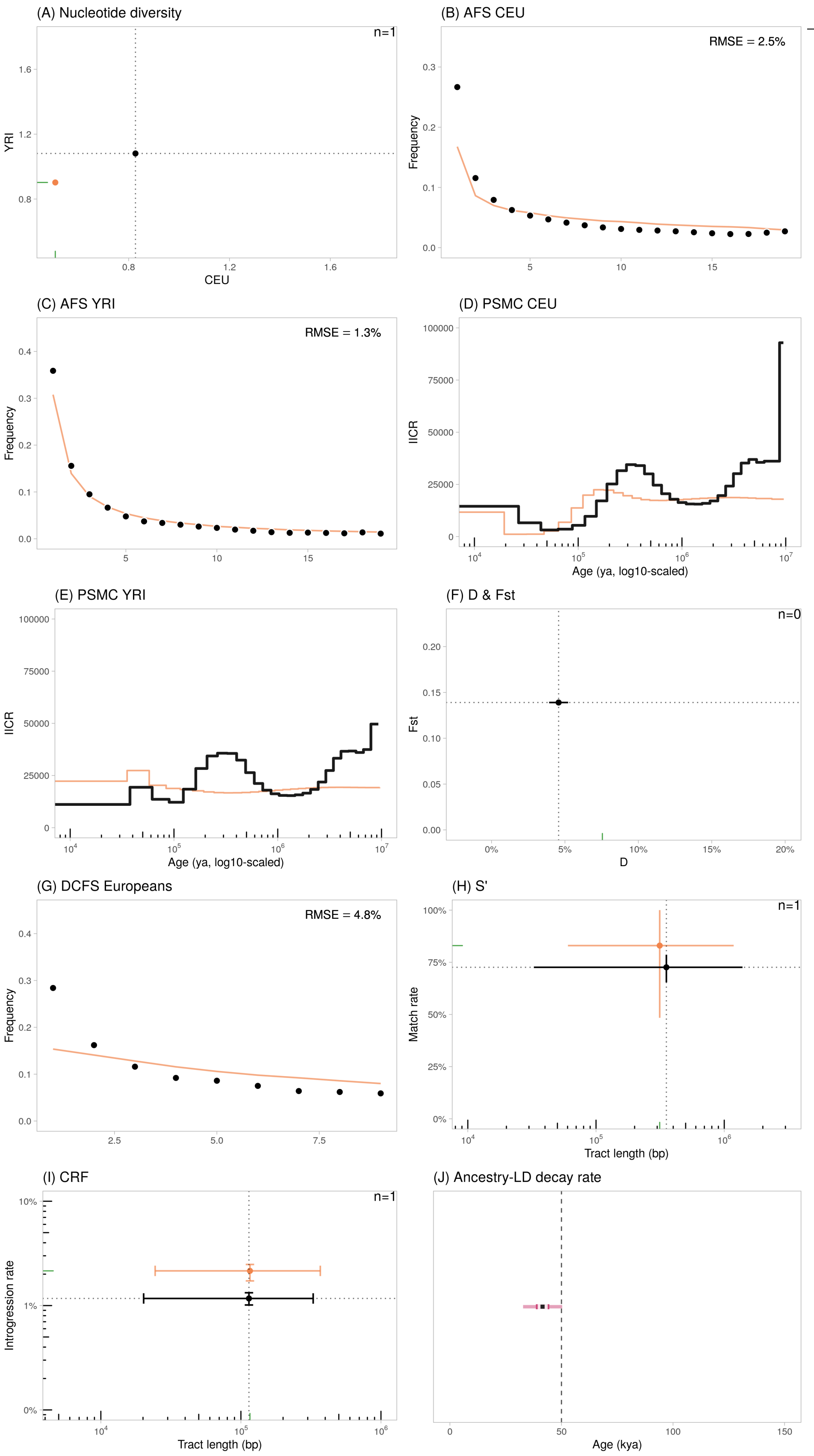

Figure S24: Statistics calculated using the Gower et al. (2021) scenario with the standard mutation rate of $1.2 \times 10^{-8}$ b$^{-1}$g$^{-1}$. The error bars represent the confidence intervals at 95% ($= 1.96 \cdot SE$) except for $S'$ (H) and $CRF$ (I) where they represent the 2.5- and 97.5%-percentiles of the distributions. Note that in (F), the dots are outside of the figure limits, towards the top.

#### S8.7 Yang et al. (2012)

Yang et al. (2012) implemented an archaic admixture model connecting YRI, non-African *Hs* (CEU) and Neanderthal, to investigate the performance of their *DCFS* statistics. They assumed a single episode of admixture from Neanderthal into non-Africans (at time $t_{GF}$ set at 50 kya) and panmixia within all three populations. The authors also included a bottleneck in the non-African population, reducing the population size by a factor 100 for 100 generations. We considered the scenario in which the bottleneck is older than the admixture as it provided the best fit to the *DCFS* (Yang et al., 2012), along with an admixture rate of 5% and no migration between CEU and YRI (Fig. 2c of Yang et al. (2012)). The authors used a generation time of 25 years.

Practically, we retrieved the `ms` command provided by the authors in their Appendix B (p. 8) using parameters of their Table 1, where populations are ordered as: Neanderthal, YRI and CEU:

```
-I 3 1 1 10 -n 2 100 -n 3 100 -m 3 2 0 -m 2 3 0
-es 0.05 3 0.95 -ej 0.05 4 1
-en 0.1 3 1 -ej 0.1125 3 2
-en 0.1150 2 1 -en 0.125 3 100
-ej 0.3 2 1
```

Listing 2: Command `ms` from Yang et al. (2012) Appendix B, bottleneck older than admixture.

We modified the sampling to have 50 CEU diploids, 50 YRI diploids, 1 diploid Neanderthal sampled at 50 kya (corresponding to Vindija33.19). The `ms` command was converted into a `demes` graph object using `demes::from_ms()` with $N_o = 10^4$ (as specified by the authors, p. 4) and later into a `msprime` demography object using `msprime::from_deme()`.

**Simulation results** Realistic prediction of the nucleotide diversities in CEU and YRI are obtained for mutation rates significantly greater ($> 1.5 \times 10^{-8}$) than the currently accepted rate. Using $\mu = 1.2 \times 10^{-8}$, diversities are significantly underestimated (<0.6 kb$^{-1}$). The model significantly overestimates the frequency of singletons in CEU along with an underestimation of all other variants. This pattern also affects the *AFS* of YRI (Fig. S26-B,C). The $F_{ST}$ between CEU and YRI in the simulated data is also significantly underestimated, by more than 11-fold (Fig. S26-F, $y$-axis). The $PSMC_{CEU}$ and $PSMC_{YRI}$ curves are flat before 100 kya, and later reach population sizes more than 5-times higher than the empirical curve.

Regarding the admixture-sensitive statistics, the *DCFS* is rather well estimated. Yet, it does not perfectly fit the empirical curve despite the fact that this model was used for performance testing. The $D$-statistic is well estimated (Fig. S26-F, $y$-axis). The $S'$-inferred putatively archaic tracts have wide-ranging distribution and most points suggest a significant underestimation of the match rate (Fig. S26-H, $y$-axis), including for the standard rate of $1.2 \times 10^{-8}$. Regarding the *CRF*-inferred tracts, the introgression rates are realistic but the mean tract lengths are underestimated by 32% on average (Fig. S26-I, $x$-axis). The ancestry-*LD* decay constants ($\sim$52 kya assuming $g = 25$) are in accordance with the age implemented in the model.

Overall, the Yang et al. (2012) model fails to reproduce the *PSMC* trajectory in CEU and YRI as well as several genome-wide statistics (e.g. *AFS* or $F_{ST}$). Still, it provides realistic measures for several admixture-sensitive statistics like the *DCFS*, $D$-statistic or the *CRF*-inferred introgression rate.

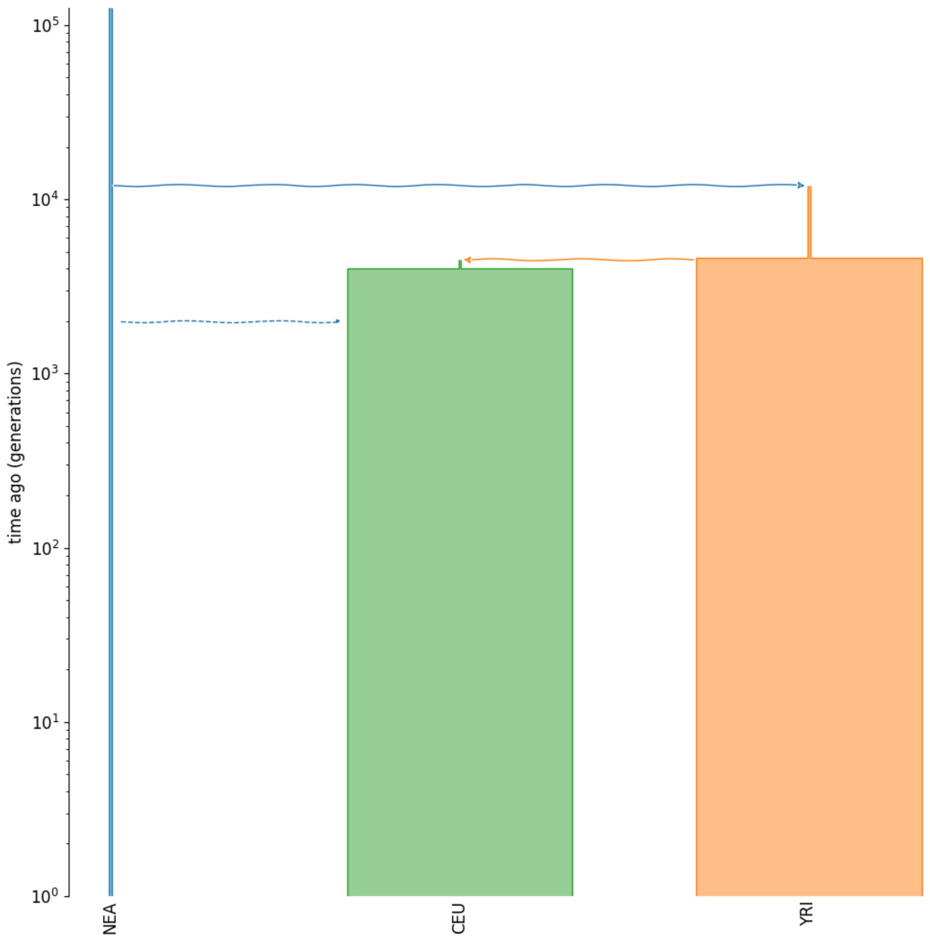

Figure S25: Visual representation of the scenario from Yang et al. (2012) . The $y$-axis is in generations BP and $\log_{10}$-scaled.

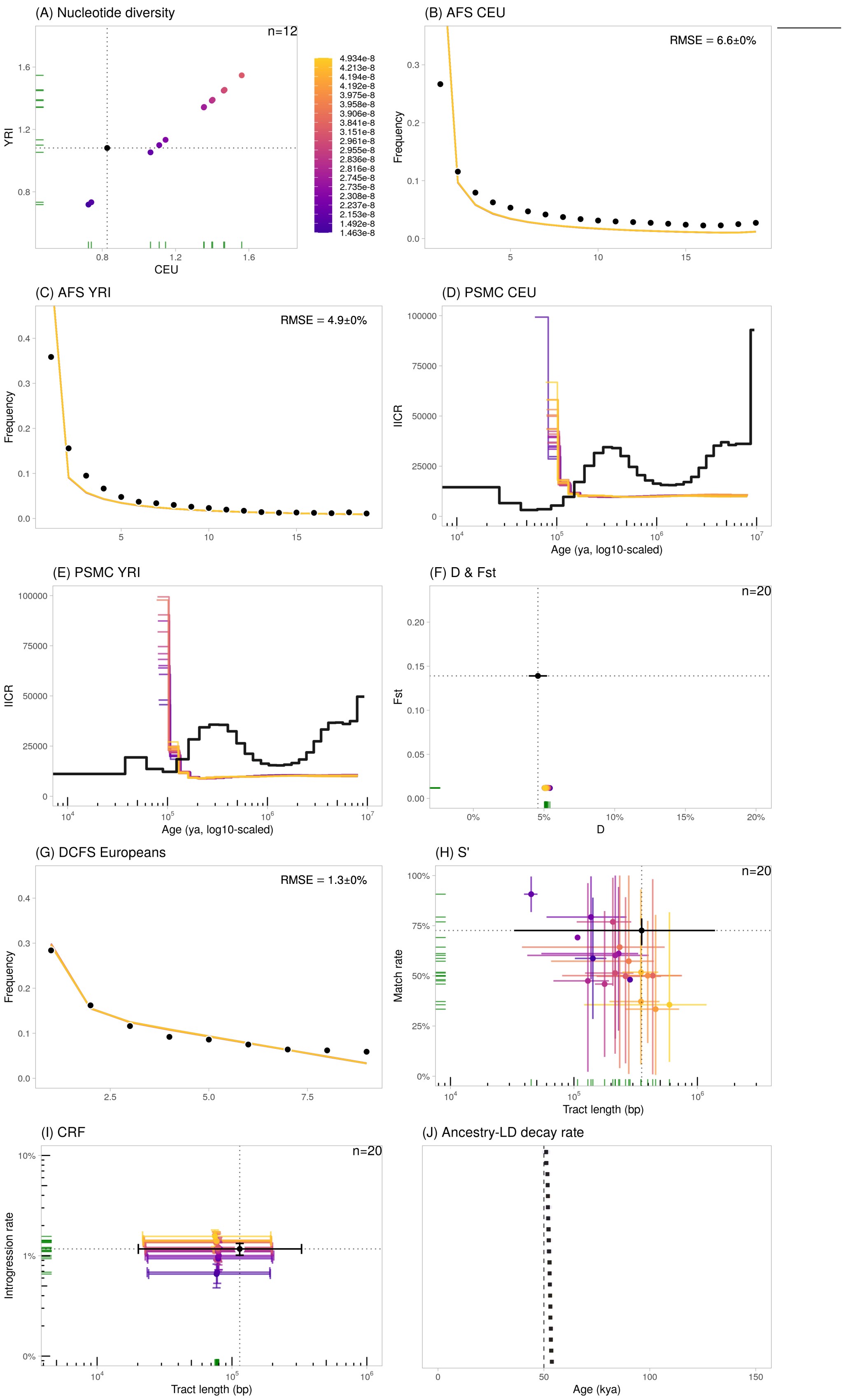

Figure S26: Statistics calculated using the Yang et al. (2012) scenario, using varying mutation rates ($5 \times 10^{-9}$ to $5 \times 10^{-8}$ b$^{-1}$g$^{-1}$). We plotted the 20 runs closest to the observed data. The error bars represent the confidence intervals at 95% ($= 1.96 \cdot SE$) except for $S'$ (H) and $CRF$ (I) where they represent the 2.5- and 97.5%-percentiles of the distributions.

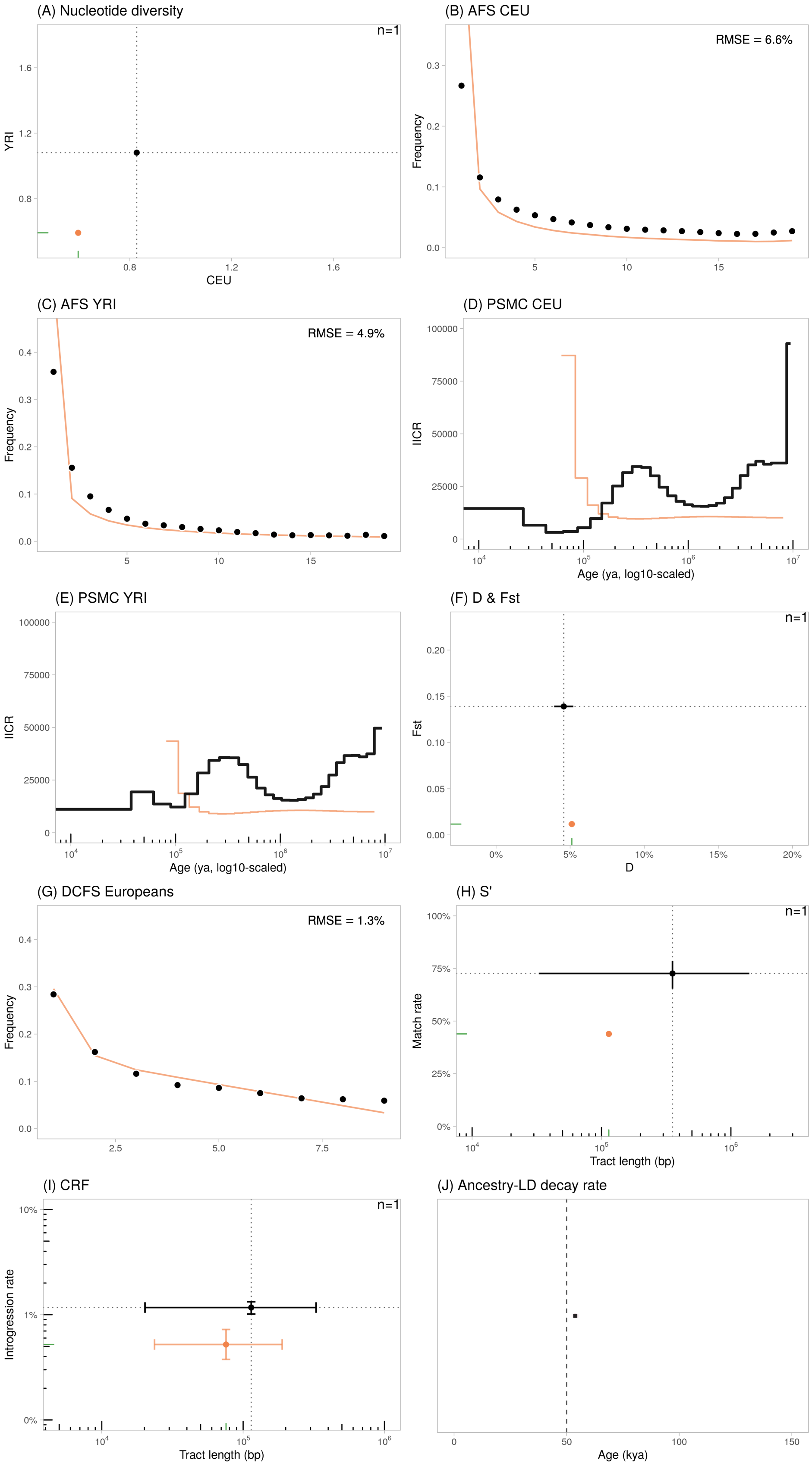

Figure S27: Statistics calculated using the Yang et al. (2012) scenario with the standard mutation rate of $1.2 \times 10^{-8}$ b$^{-1}$g$^{-1}$. The error bars represent the confidence intervals at 95% ($= 1.96 \cdot SE$) except for $S'$ (H) and $CRF$ (I) where they represent the 2.5- and 97.5%-percentiles of the distributions. In (H), the colored dot lacks bars because only one fragment was detected with $S'$ in the simulated dataset.

## S8.8 Fu et al. (2014)

Fu et al. (2014) investigated the performance of their Neanderthal admixture dating using a series of demographic models with increasing complexity. We considered the more complex model (Simulation 4, Supplementary Information 18, p. 113), derived from the model fit by Gravel et al. (2011) with modifications to account for the genetic diversity and timing of ancient DNA specimens (especially the Tianyuan individual dated c. 40 kya). In this model, the Neanderthal sample (Altai) split from a Neanderthal population which is the source of introgression into non-African *Hs* (2,200 generations ago, with rate 3%). The model implements an instantaneous population expansion for the common ancestor of all present-day humans, as well as a recent exponential population growth in the European and East Asian populations following their split-with-bottleneck from the African population. The authors used a generation time of 29 years.

Practically, we used the `ms` command provided by the authors in their Supplementary Material p. 113, where populations are ordered as: pop1=African, pop2=European, pop3=Asian, pop4-7=Ust'-Ishim, pop8-11=Neanderthal:

```
ms 68 1 -I 11 20 20 20 1 1 1 1 1 1 1 1 -en 0 1 1 -en 0 2 2.41428571428571
-en 0 3 3.23571428571429 -eg 0 2 97.6909397920288 -eg 0 3 123.512032930849
-en 0 4 7.14285714285714e-11 -en 0 5 7.14285714285714e-11
-en 0 6 7.14285714285714e-11 -en 0 7 7.14285714285714e-11
-en 0 8 7.14285714285714e-11 -en 0 97.14285714285714e-11
-en 0 10 7.14285714285714e-11 -en 0 11 7.14285714285714e-11
-ej 0.0321428571428571 5 4 -ej 0.0321428571428571 7 6
-en 0.0321430357142857 4 0.714285714285714
-en 0.0321430357142857 6 0.714285714285714 -ej 0.0321446428571429 6 4
-en 0.0321464285714286 4 0.714285714285714 -ej 0.0357142857142857 2 3
-ej 0.0357142857142857 4 3 -en 0.0357160714285714 3 0.132857142857143
-en 0.0357160714285714 3 0.132857142857143 -es 0.0392857142857143 3 0.97
-en 0.0392875 12 0.178571428571429 -en 0.0392875 3 0.132857142857143
-ej 0.0428571428571429 9 8 -ej 0.0428571428571429 11 10
-en 0.0428573214285714 8 0.178571428571429 -en 0.0428573214285714 10
    0.178571428571429
-ej 0.0428589285714286 10 8 -en 0.0428607142857143 8 0.178571428571429
-ej 0.0535714285714286 3 1 -en 0.0535732142857143 1 1 -ej 0.0714285714285714 12 8
-en 0.0714303571428571 8 0.178571428571429 -en 0.107142857142857 1 0.521428571428571
-ej 0.214285714285714 8 1 -en 0.2142875 1 0.714285714285714
-r 28000 50000000 -t 42000
```

Listing 3: Command `ms` from Fu et al. (2014) SM p. 113.

We modified the sampling to have 50 CEU diploids, 50 YRI diploids, 1 diploid Neanderthal sampled at 50 kya (corresponding to Vindija33.19). The `ms` command was converted into a `demes` graph object using `demes::from_ms()` with $N_o = 14 \cdot 10^3$ and later into a `msprime` demography object using `msprime::from_deme()`. $N_o$ was deduced given the mutation rate set by the authors ($\mu = 1.5 \cdot 10^{-8}$), the sequence length ($L = 50 \cdot 10^6$ bp, from the `-r` switch), then $N_o = \theta/(4 \cdot \mu \cdot L)$ with $\theta = 42000$ (cf. command).

**Simulation results** Realistic prediction of the nucleotide diversities in CEU and YRI are obtained for mutation rates significantly greater ($> 2.5 \times 10^{-8}$) than the currently accepted rate. Using $\mu = 1.2 \times 10^{-8}$, diversities are significantly underestimated (0.34 kb$^{-1}$ for CEU for instance, and 0.48 kb$^{-1}$ for YRI). The model seems to predict realistic derived *AFS* for both CEU and YRI (Fig. S29-B,C), however the $F_{ST}$ between CEU and YRI is significantly overestimated by ∼70%. The $PSMC_{CEU}$ and $PSMC_{YRI}$ curves are flat before 100 kya ($IICR \sim 15k$), with a very small bump ∼100 kya for $PSMC_{CEU}$ (Fig. S29-D).

Regarding the admixture-sensitive statistics, the *DCFS* is nearly linear, with a significant underestimation of the singleton category (Fig. S29-G). The *D*-statistic is overestimated by 50% on average (Fig. S29-F, $x$-axis). The $S'$-inferred putatively archaic tracts show realistic match rate (albeit slightly underestimated) and average length distribution (Fig. S29-H). Regarding the *CRF*-inferred tracts, the introgression rates are realistic but the mean tract lengths are slightly underestimated by 24% on average (Fig. S29-I, $x$-axis). The ancestry-$LD$ decay constants (∼52 kya assuming $g = 25$) are in line with the admixture age simulated in the model (55 kya assuming $g = 25$).

Overall, the Fu et al. (2014) model fails to reproduce the $PSMC$ trajectory in CEU and YRI and the level of genetic differentiation between CEU and YRI. The inaccurate estimation of the $F_{ST}$ was already pointed out in Moorjani et al. (2016) (p. 9 of the Supplementary Materials). Although this model provides realistic measures for the haplotype-related statistics, it does not provide a realistic modeling of the $DCFS$ in CEU.

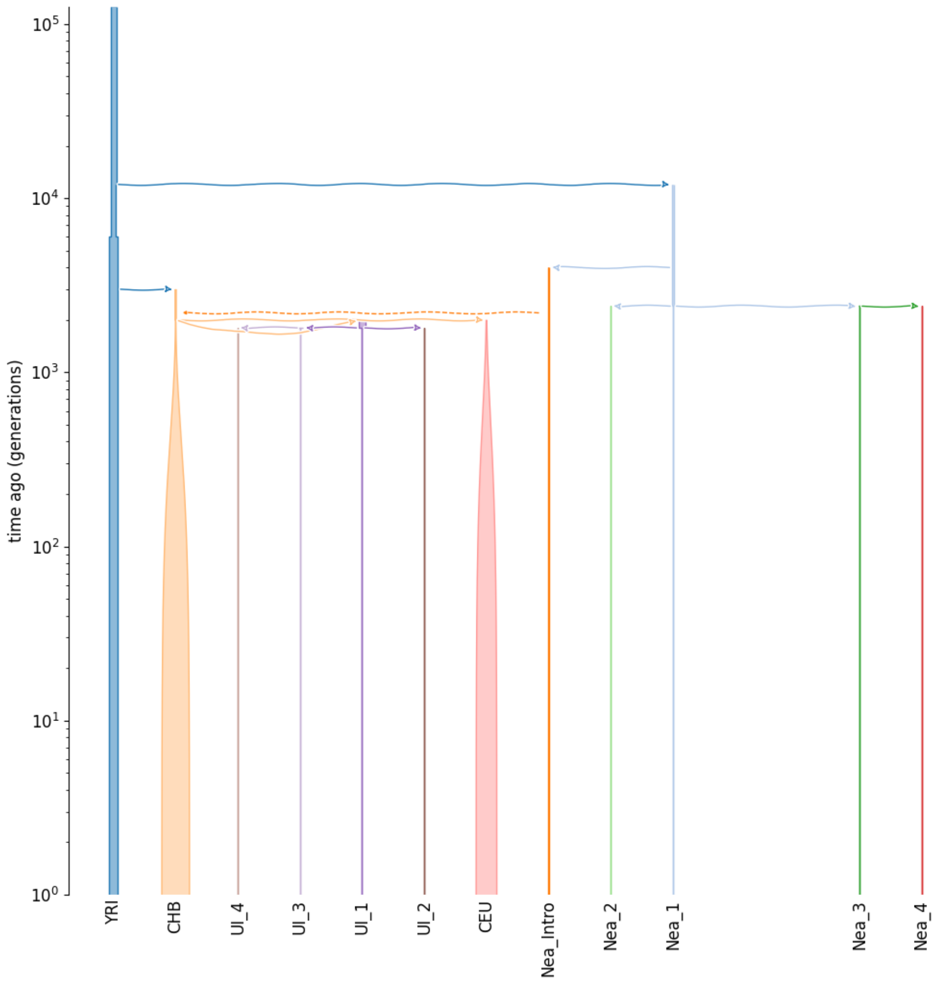

Figure S28: Visual representation of the scenario from Fu et al. (2014) . The $y$-axis is in generations BP and $\log_{10}$-scaled.

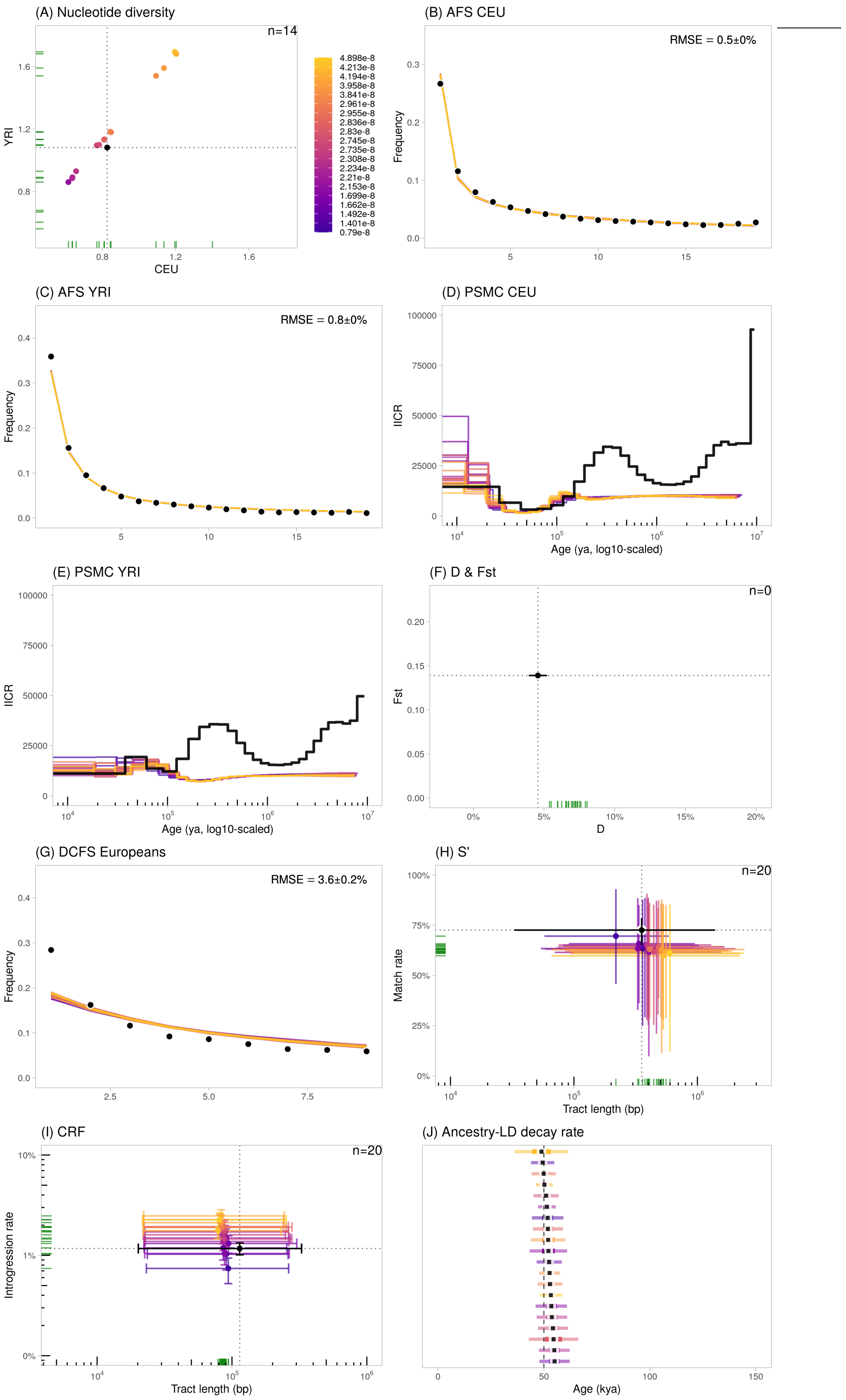

Figure S29: Statistics calculated using the Fu et al. (2014) scenario, using varying mutation rates ($5 \times 10^{-9}$ to $5 \times 10^{-8}$ b$^{-1}$g$^{-1}$). We plotted the 20 runs closest to the observed data. The error bars represent the confidence intervals at 95% ($= 1.96 \cdot SE$) except for $S'$ (H) and $CRF$ (I) where they represent the 2.5- and 97.5%-percentiles of the distributions. In (F), the dots are outside of the figure limits, towards the top.

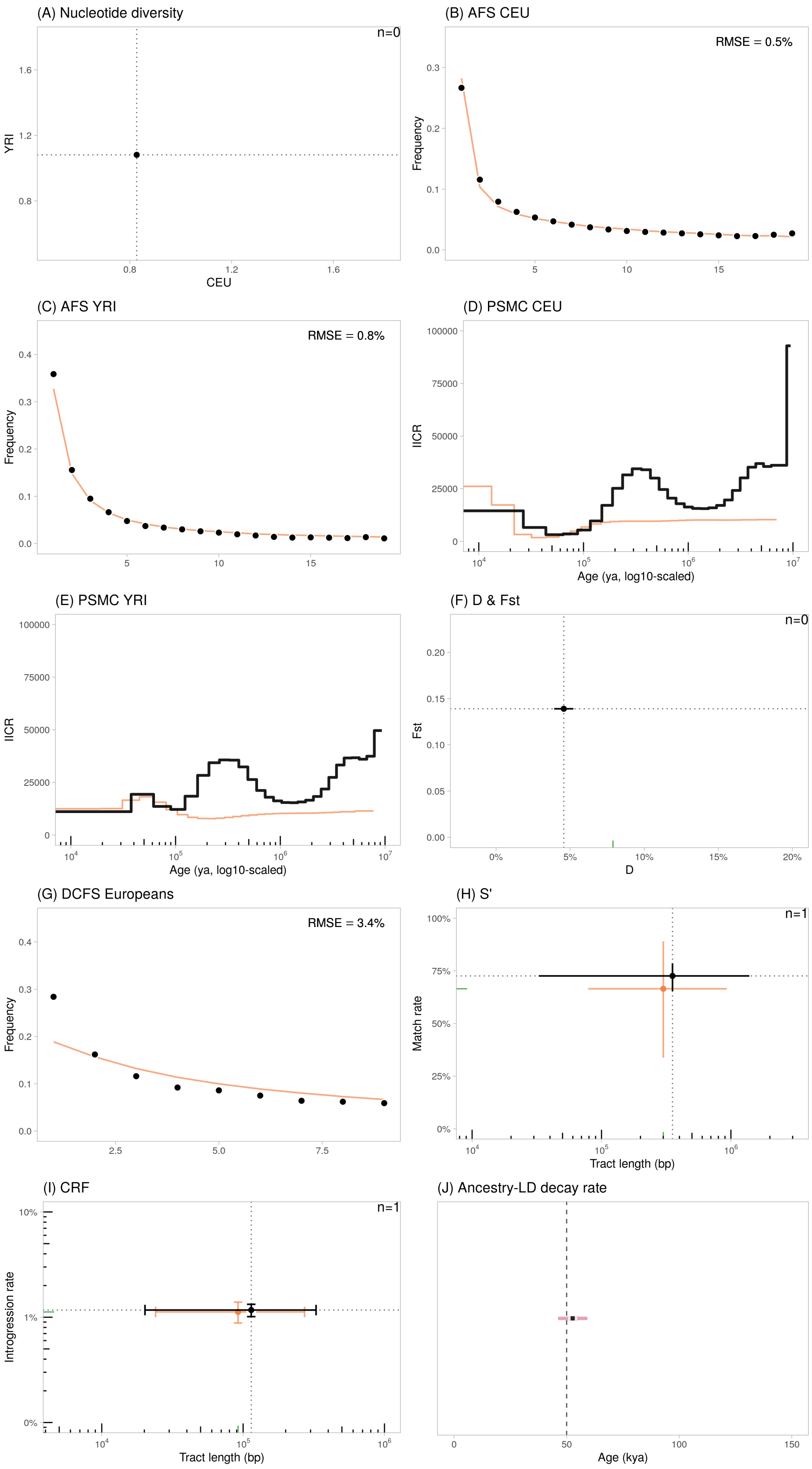

Figure S30: Statistics calculated using the Fu et al. (2014) scenario with the standard mutation rate of $1.2 \times 10^{-8}$ b$^{-1}$g$^{-1}$. The error bars represent the confidence intervals at 95% ($= 1.96 \cdot SE$) except for $S'$ (H) and $CRF$ (I) where they represent the 2.5- and 97.5%-percentiles of the distributions. Note that in (A) and (F), the dots are outside of the figure limits (towards the bottom-left and towards the top, respectively).

#### S8.9 Moorjani et al. (2016)

Moorjani et al. (2016) assessed the performance of their single-sample ancestry-*LD* decay approach to estimate the timing of archaic admixture, using a demographic model with parameters chosen to match the empirical $F_{ST}$ between YRI and CEU ($F_{ST}$=0.15) and the *D*-statistic with Neanderthals ($D$=5%). This model S2a assumes that ancient Europeans diverged from West Africans and received a single pulse of gene flow from Neanderthals with rate 3% at age $t_n$. We used $t_n = 56$ kya (under the authors' $g = 28$, i.e. 2,000 generations ago) as the date of the shared Neanderthal gene flow, following Notes S2c of the original study. The authors used a generation time of 28 years.

Practically, we used the `ms` command provided by the authors in their Supplementary Material p. 7, where populations are ordered as: pop1=west African, pop2=European, pop3-6=Neanderthal:

```
ms 44 1 -r 20000 50000000 -t 30000 -I 6 20 20 1 1 1 1 -en 0 1 1
-en 0 2 1 -en 0 3 1e-10 -en 0 4 1e-10 -en 0 5 1e-10 -en 0 6 1e-10
-es tn 2 0.97 -en 0.02500025 7 0.25 -en 0.02500025 2 1
-ej 0.05 4 3 -ej 0.05 6 5 -en 0.05000025 3 0.25 -en 0.05000025 5 0.25
-ej 0.0500025 5 3 -en 0.050005 3 0.25 -ej 0.075 2 1
-en 0.0750025 1 1 -ej 0.1 7 3 -en 0.1000025 3 0.25 -ej 0.3 3 1
-en 0.3000025 1 1
```

Listing 4: Command `ms` from Moorjani et al. (2016) SM p. 7.

We modified the sampling to have 50 CEU diploids, 50 YRI diploids, 1 diploid Neanderthal sampled at 50 kya (corresponding to Vindija33.19). The `ms` command was converted into a `demes` graph object using `demes::from_ms()`. Contrary to the model S2a which assumes $N_o = 1 \cdot 10^4$, we used the value from the more complex model S2b ($N_o = 1.4 \cdot 10^4$) because it led to an improved fit for nearly all summary statistics under $\mu = 1.2 \times 10^{-8}$.

**Simulation results** The best fit for the nucleotide diversities in CEU and YRI are obtained for realistic mutation rates which include the standard value of $\sim$1.2$\times 10^{-8}$. We observed an overestimation of low-frequency variants on the *AFS* CEU and a reverse underestimation by 21% of the singleton variants on the *AFS* YRI (Fig. S32-B,C). The model properly predicts the observed $F_{ST}$ (Fig. S32-E, $y$-axis). The $PSMC_{CEU}$ curve is flat, with $IICR \sim 20k$, which can be expected given that the model does not include population structure nor changes in population sizes or connectivity.

Regarding the admixture-sensitive statistics, the *DCFS* is well estimated, with a significant L-shape in agreement with the empirical curve (S32-G). The *D*-statistic is also well estimated (Fig. S32-F, $x$-axis). The $S'$-inferred putatively archaic tracts have a match rate which tend to underestimate the empirical value ($< 70\%$) and a wide distribution of mean tract lengths, some of which encompass the empirical value for certain mutation rates (including the standard one) (Fig. S32-H, $x$-axis). Regarding the *CRF*-inferred tracts, the introgression rates vary as a function of the mutation rate (Fig. S32-I, $y$-axis) while the mean lengths are slightly underestimating the empirical value, by 33% on average. The ancestry-*LD* decay constants are centred around 50 kya (assuming $g = 25$), concordant with the simulated Neanderthal pulse of admixture from the model.

Overall, the Moorjani et al. (2016) model provides realistic values for the admixture-sensitive statistics but poorly predicts a few genome-wide statistics like the derived *AFS* and the *PSMC*.

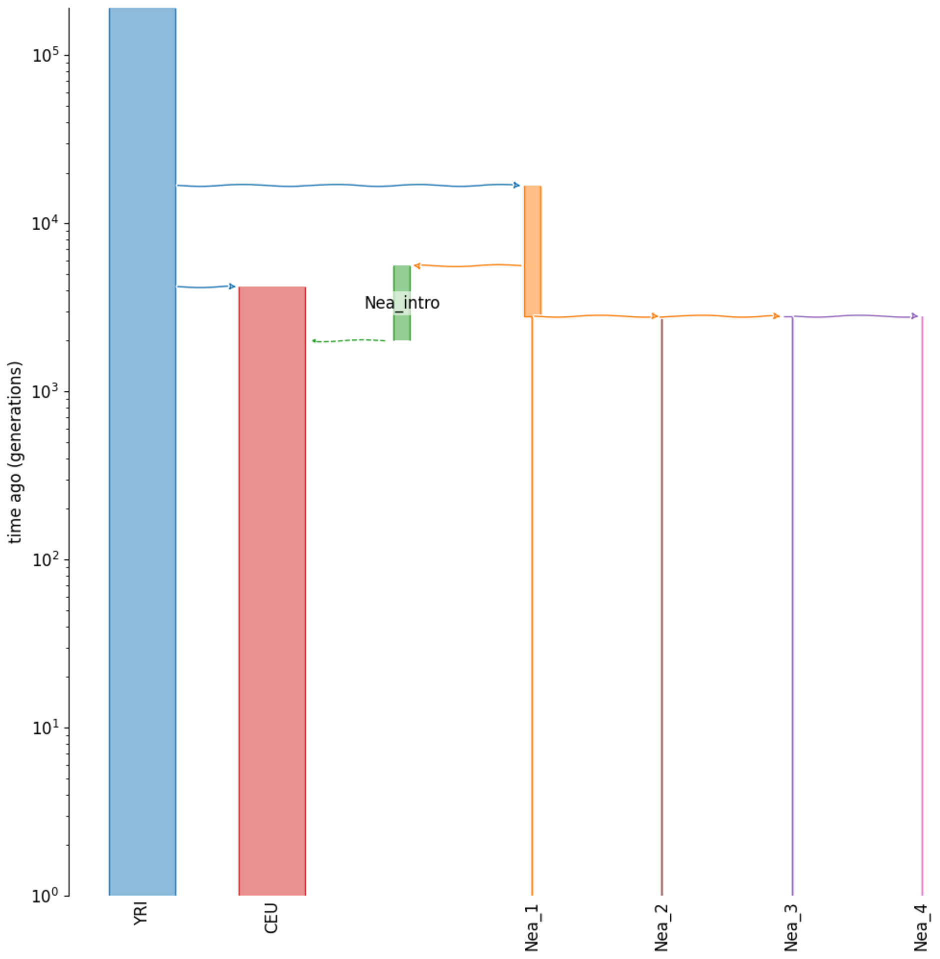

Figure S31: Visual representation of the scenario from Moorjani et al. (2016) . The $y$-axis is in generations BP and $\log_{10}$-scaled.

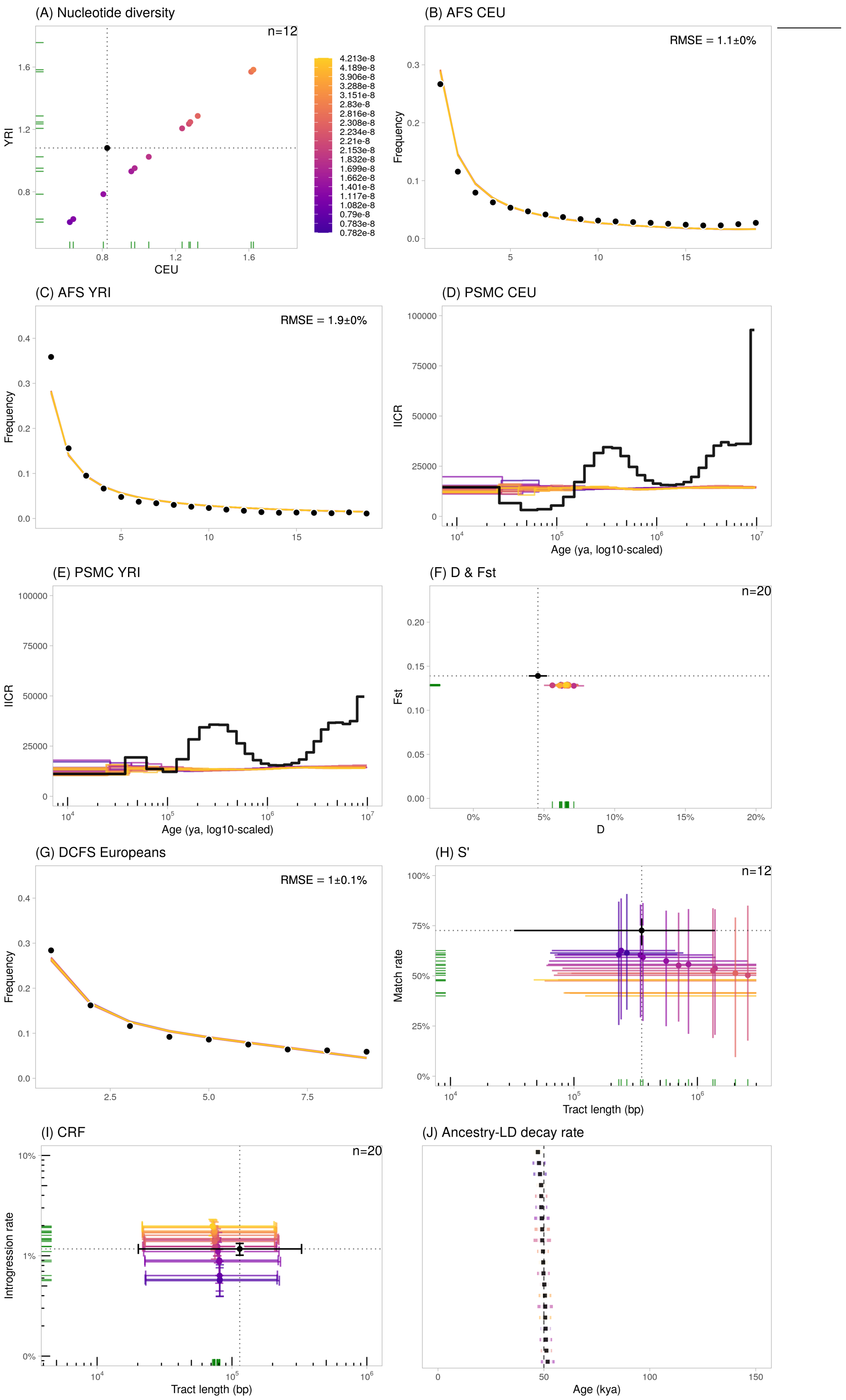

Figure S32: Statistics calculated using the Moorjani et al. (2016) scenario, using varying mutation rates ($5 \times 10^{-9}$ to $5 \times 10^{-8}$ b$^{-1}$g$^{-1}$). We plotted the 20 runs closest to the observed data. The error bars represent the confidence intervals at 95% ($= 1.96 \cdot SE$) except for $S'$ (H) and $CRF$ (I) where they represent the 2.5- and 97.5%-percentiles of the distributions.

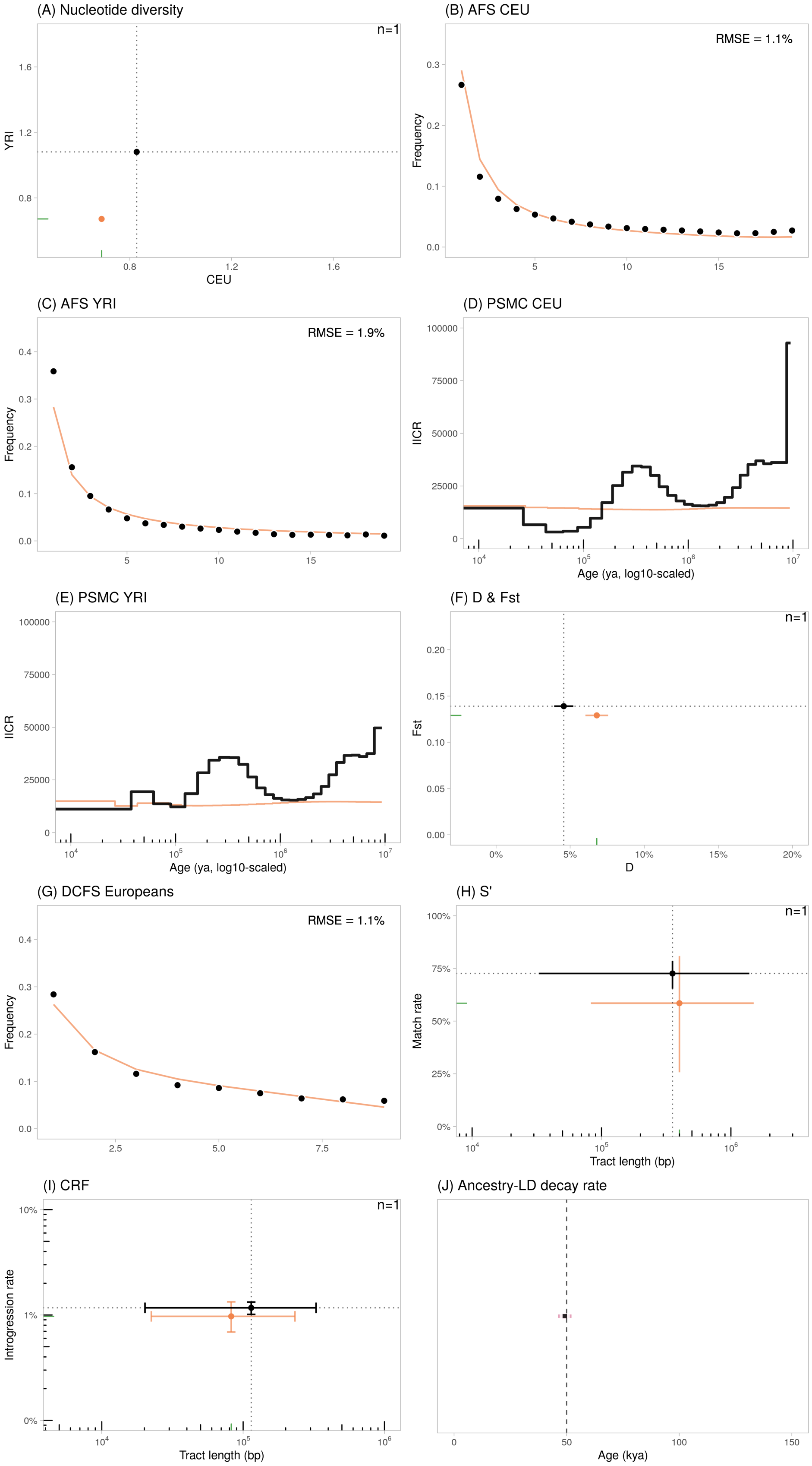

Figure S33: Statistics calculated using the Moorjani et al. (2016) scenario with the standard mutation rate of $1.2 \times 10^{-8}$ b$^{-1}$g$^{-1}$. The error bars represent the confidence intervals at 95% ($= 1.96 \cdot SE$) except for $S'$ (H) and $CRF$ (I) where they represent the 2.5- and 97.5%-percentiles of the distributions.

#### S8.10 Skov et al. (2020)

Skov et al. (2020) implemented various demographic models to assess the performance of their hidden Markov model to detect introgressed archaic segments. Among these, they tested a complex demographic scenario with multiple bottlenecks in the European population (Figure 3.1.1 of their Supplementary Material, p. 48). This model includes a single pulse of gene flow from the introgressing Neanderthal population into the ancestors of European populations (here modeled as Icelandic) with rate 2%, and varying population sizes in the archaic hominin lineages. The authors assumed a generation time of 29 years.

Practically, we retrieved the Skov et al. (2020) model formatted in the `msprime 0.7` syntax from the authors' GitHub[7]. We then converted it to the `msprime 1.1.1` syntax using `msprime::from_old_style()`.

**Simulation results** Realistic prediction of the nucleotide diversities in CEU and YRI were obtained for mutation rates significantly greater ($> 2.5 \times 10^{-8}$) that the currently accepted rate. Using $\mu = 1.2 \times 10^{-8}$, diversities are significantly underestimated (0.34—0.37 kb$^{-1}$ for CEU and YRI respectively). The model seems to predict realistic derived *AFS* for CEU (Fig. S35-B), but significantly overestimates the singleton variants in YRI (Fig. S35-C). The $F_{ST}$ between CEU and YRI is realistic, although slighly overestimated by 20% (Fig. S35-F, $y$-axis). The $PSMC_{CEU}$ curve is nearly flat ($IICR \sim 15k$), with a very discrete bump ~200-300 kya, at about the same period as the empirical trajectory. The overall trajectory still departs significantly from the empirical one (S35-D), as does $PSMC_{YRI}$ (S35-E).

Regarding the admixture-sensitive statistics, the *DCFS* is excessively L-shaped compared to the empirical decay, with a significant overestimation of low-frequency variants and an underestimation of high-frequency ones (Fig. S35-G). The $D$-statistic is extremely overestimated, by more than 6-fold (Fig. S35-F, $x$-axis). The $S'$-inferred putatively archaic tracts have match rates which underestimate the empirical value by ~2-fold (Fig. S35-H, $y$-axis). For some mutation rates, the mean tract length encompasses the empirical value (Fig. S35-H, $x$-axis). Regarding the *CRF*-inferred tracts, the mean lengths are realistic (Fig. S35-I, $x$-axis). The ancestry-*LD* decay constants are ~35 kya (assuming $g = 25$).

Overall, the Skov et al. (2020) model fails to reproduce the *PSMC* trajectory in CEU and YRI and the genome-wide derived *AFS* in YRI. Also, it fails to provide realistic *DCFS* and *S'*-inferred match rates despite satisfying performance on other admixture-sensitive statistics. We note however that for a standard, realistic mutation rate of $1.2 \times 10^{-8}$, the model underestimates very significantly the actual genetic diversity in the genomes of CEU and YRI samples (S36-A) along with too high values for the $D$-statistics.

[7] https://github.com/LauritsSkov/ArchaicSimulations/blob/master/SI%203%20Dataset%20-%20Simulation%20script.py

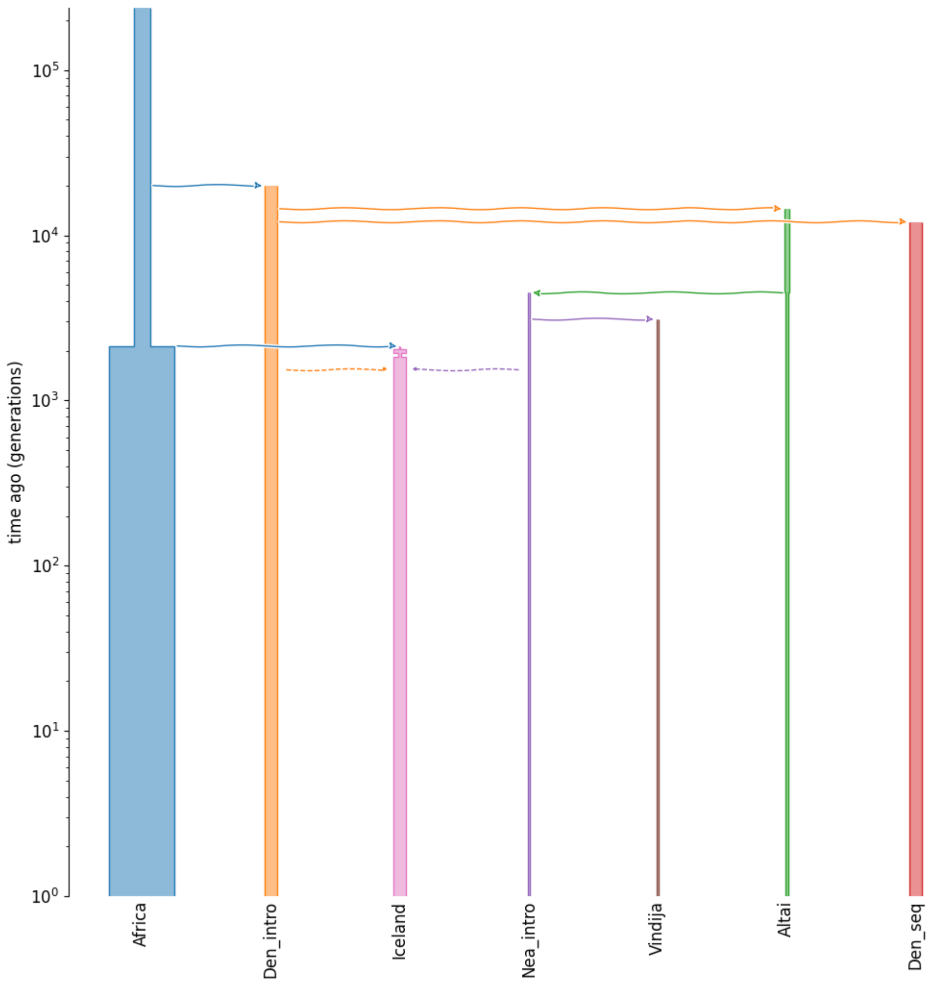

Figure S34: Visual representation of the scenario from Skov et al. (2020) . The $y$-axis is in generations BP and $\log_{10}$-scaled.

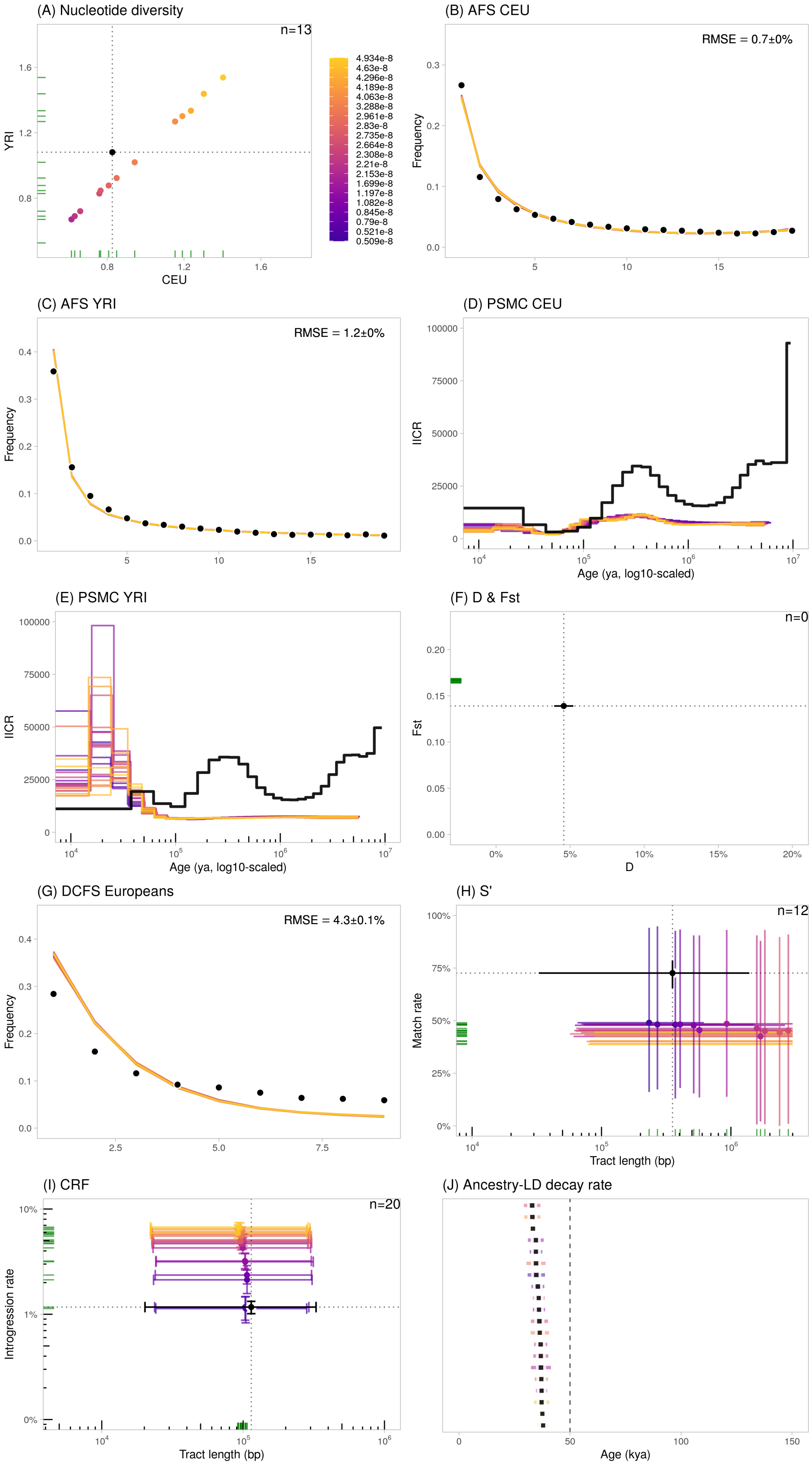

Figure S35: Statistics calculated using the Skov et al. (2020) scenario, using varying mutation rates ($5 \times 10^{-9}$ to $5 \times 10^{-8}$ b$^{-1}$g$^{-1}$). We plotted the 20 runs closest to the observed data. The error bars represent the confidence intervals at 95% ($= 1.96 \cdot SE$) except for $S'$ (H) and $CRF$ (I) where they represent the 2.5- and 97.5%-percentiles of the distributions. Note that in (F), the dots are outside of the figure limits, towards the right.

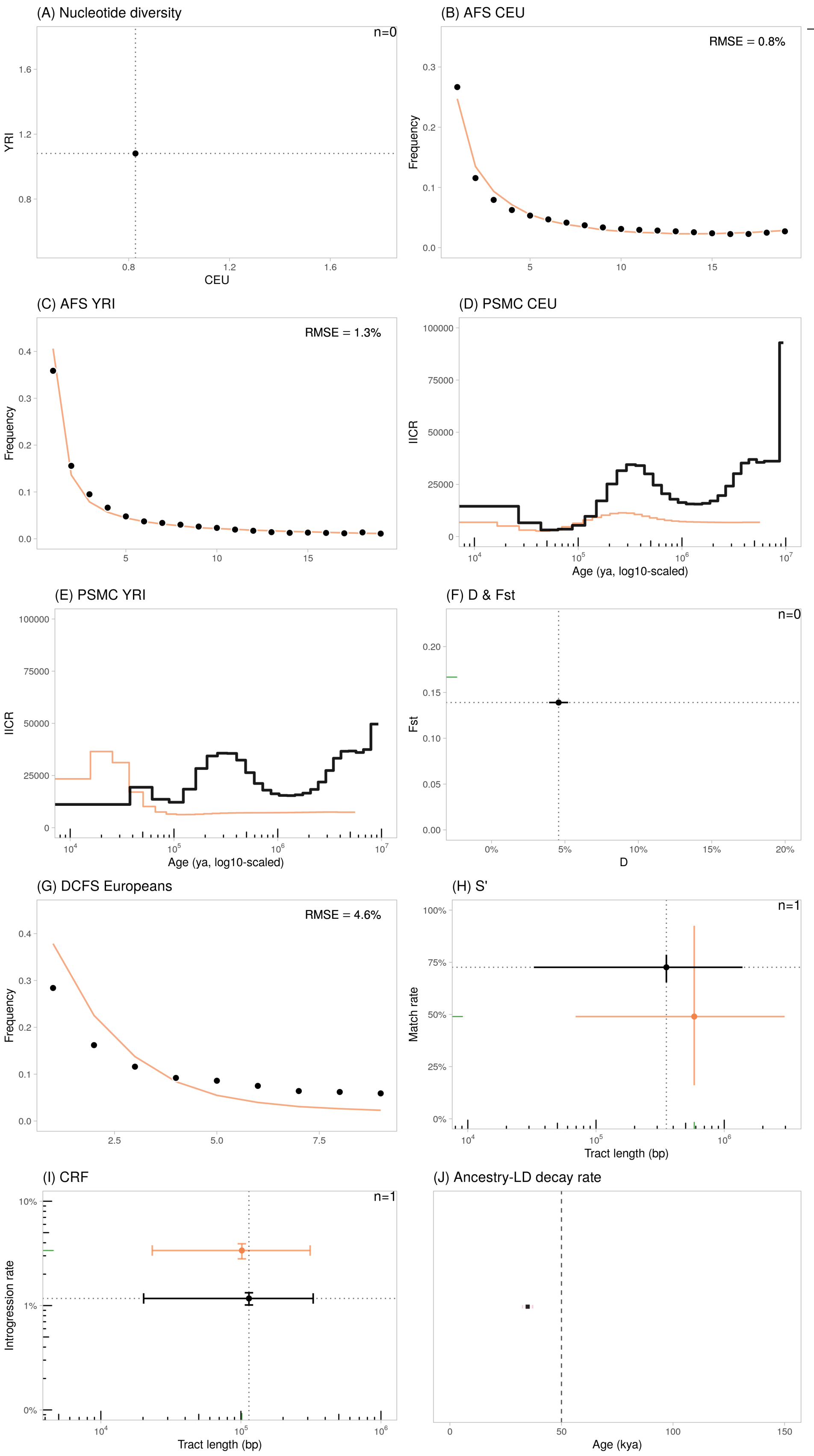

Figure S36: Statistics calculated using the Skov et al. (2020) scenario with the standard mutation rate of $1.2 \times 10^{-8}$ b$^{-1}$g$^{-1}$. The error bars represent the confidence intervals at 95% ($= 1.96 \cdot SE$) except for $S'$ (H) and $CRF$ (I) where they represent the 2.5- and 97.5%-percentiles of the distributions. Note that in (A) and (F), the dot is outside of the figure limits (towards the bottom-left for (A) and towards the right for (F)).

#### S8.11 Schaefer et al. (2021)

Schaefer et al. (2021) implemented a method, SARGE, to detect archaic ancestry along target genomes, using ancestral recombination graphs (ARGs). This supervised learning approach requires training data, that the authors simulated using a demographic model derived from the Gutenkunst et al. (2009) "OOA model". They however modified this model to account for the Altai Neanderthal, Vindija Neanderthal and Altai Denisovan demography. They assumed a single pulse of gene flow from introgressing Neanderthals into the ancestors of non-Africans 2,000 generations ago, with varying admixture rates $x$. We fixed $x$ at 2%, in line with most studies cited here, to best match the observed $D$-statistics. The authors used a generation time of 25 years.

Practically, we used the `scrm` command provided by the authors in their main text p. 12, where $[1-x]$ means one minus the admixture proportion. We excluded the switches `-T` and `-eI` which do not apply here. The populations are ordered as YRI (pop1), CEU (pop2), CHB (pop3), Altai Neanderthal (pop4), Neanderthal introgressing lineage (pop5), Vindija Neanderthal (pop6), Denisova (pop7):

```
scrm 456 1 -t 17253.7128713 -r 13802.970297 25000000 -I 7 150 150 150 0 0 0 0
-n 1 1.68 -n 2 3.74
-n 3 7.29 -n 4 0.231834158238 -n 5 0.231834158238 -n 6 0.231834158238
-n 7 0.0260813428018 -eg 0 2 116.010723 -eg 0 3 160.246047
-m 2 3 2.797460 -m 3 2 2.797460 -ej 0.028985 3 2 -en 0.028985 2 0.287184
-em 0.028985 1 2 7.293140 -em 0.028985 2 1 7.293140
-es 0.0144896348899 3 [1-x] -ej 0.0144896348899 8 7
-es 0.0362240872247 2 [1-x] -ej 0.0362240872247 9 5
-ej 0.0724481744495 6 5 -ej 0.197963 2 1 -en 0.303501 1 1
-ej 0.099616239868 5 4 -ej 0.304282332688 7 4 -ej 0.416577003084 4 1
```

Listing 5: Command `ms` from Schaefer et al. (2021) p. 12.

We modified the sampling to have 50 CEU diploids, 50 YRI diploids, 1 diploid Neanderthal sampled at 50 kya (corresponding to Vindija33.19). The `ms` command was converted into a `demes` graph object using `demes::from_ms()` with $N_o = 13,803$ and later into a `msprime` demography object using `msprime::from_deme()`. $N_o$ was deduced given the mutation rate set by the authors ($\mu = 1.25 \cdot 10^{-8}$), the sequence length ($L = 25 \cdot 10^6$ bp, from the `-r` switch), then $N_o = \theta/(4 \cdot \mu \cdot L)$ with $\theta = 17253.7128713$ (cf. command).

**Simulation results** Realistic prediction of the nucleotide diversities in CEU and YRI were obtained for mutation rates in the order of magnitude of ~$1.6\times10^{-8}$. Using $\mu = 1.2 \times 10^{-8}$, diversities are underestimated by ~25-30% (0.57—0.79 kb$^{-1}$ for CEU and YRI respectively). The model predicts realistic derived $AFS$ for both CEU and YRI (Fig. S35-B), and realistic $F_{ST}$ values between CEU and YRI (Fig. S35-F, $y$-axis). The $PSMC_{CEU}$ curve has one bump ~300—400 kya, concordant with the one visible on the empirical trajectory, but the simulated trajectory remains flat earlier into the past, in contrast with the empirical curve (S35-D). The $PSMC_{YRI}$ departs significantly from the empirical trajectory (S35-E).

Regarding the admixture-sensitive statistics, the $DCFS$ is L-shape and fits well the observed curve (Fig. S35-G). The observed $D$-statistic is also well estimated (Fig. S35-F, $x$-axis). The $S'$-inferred putatively archaic tracts have realistic match rates and mean lengths (Fig. S35-H), regardless of the mutation rate that was used. Regarding the $CRF$-inferred tracts, the introgression rates are realistic (Fig. S35-I, $y$-axis), but the mean tract lengths tend to underestimate the empirical value by 26% on average (Fig. S35-I, $x$-axis). The ancestry-$LD$ curves have decay constants ~45 kya (assuming $g = 25$).

Overall, the Schaefer et al. (2021) model provided the best prediction for all analyzed statistics among the published models that we tested, despite its poor ability to predict the empirical $PSMC$ in CEU and YRI. The fact that it does not reproduce the second ancient bump of the $PSMC$ can be explained by the fact that the model does not include changes in population size nor connectivity prior to 1 Mya. We note that although the concordance of the first *PSMC* bump with the empirical trajectory is good for CEU (but not for YRI), the amplitude of this bump still does not match the empirical one.

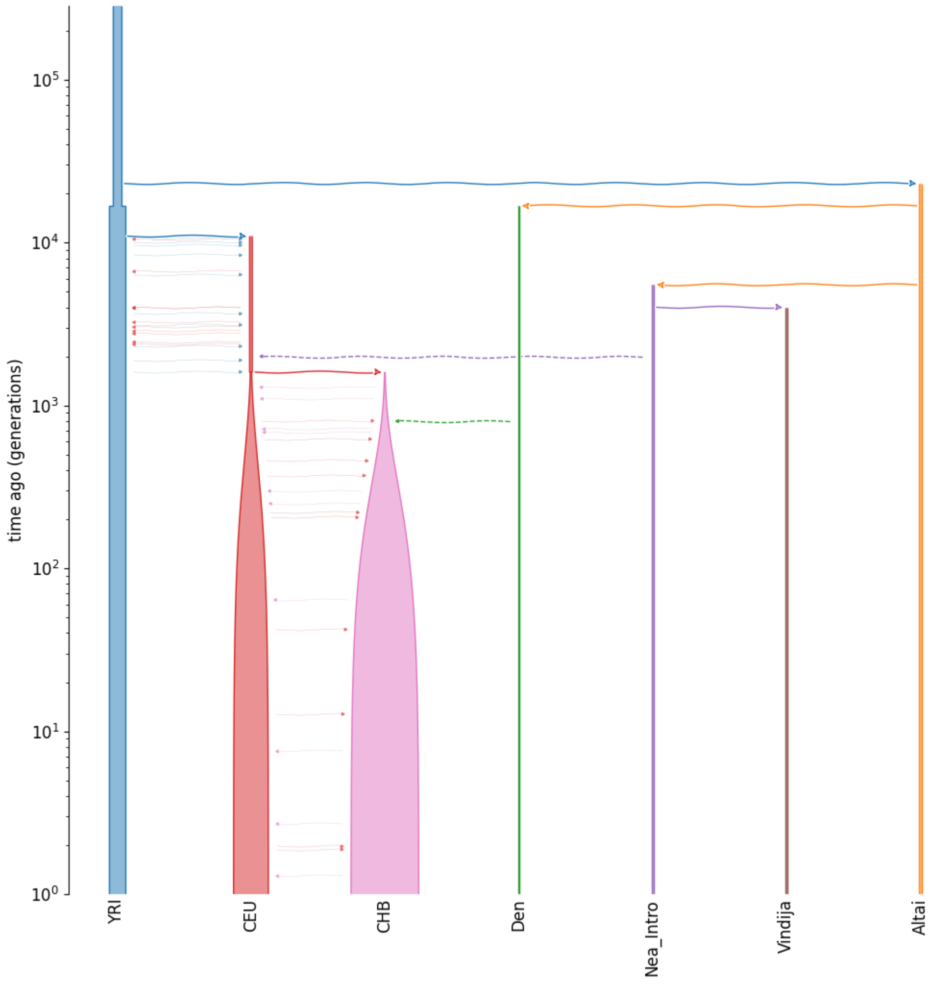

Figure S37: Visual representation of the scenario from Schaefer et al. (2021) . The $y$-axis is in generations BP and $\log_{10}$-scaled.

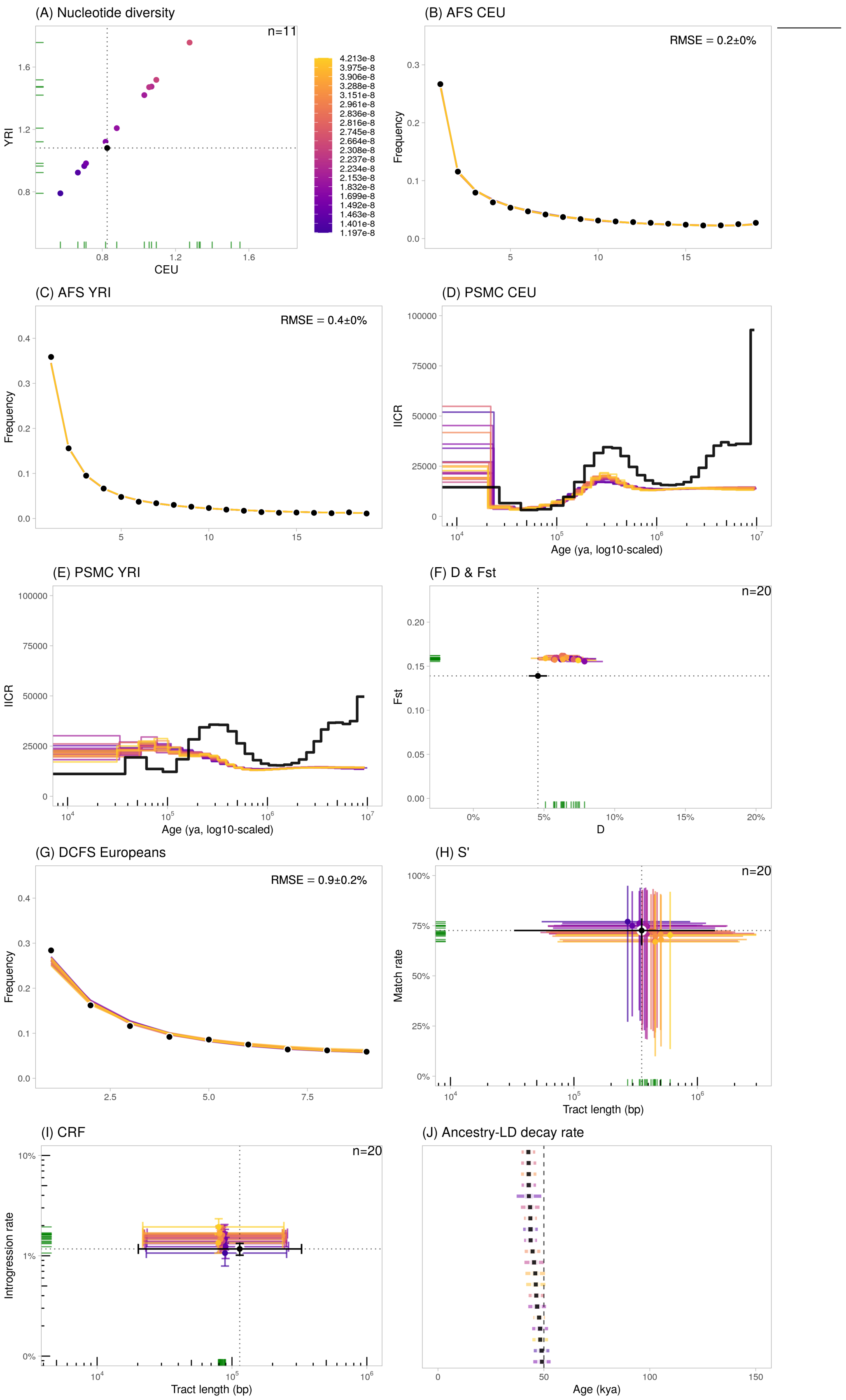

Figure S38: Statistics calculated using the Schaefer et al. (2021) scenario, using varying mutation rates ($5 \times 10^{-9}$ to $5 \times 10^{-8}$ b$^{-1}$g$^{-1}$). We plotted the 20 runs closest to the observed data. The error bars represent the confidence intervals at 95% ($= 1.96 \cdot SE$) except for $S'$ (H) and $CRF$ (I) where they represent the 2.5- and 97.5%-percentiles of the distributions.

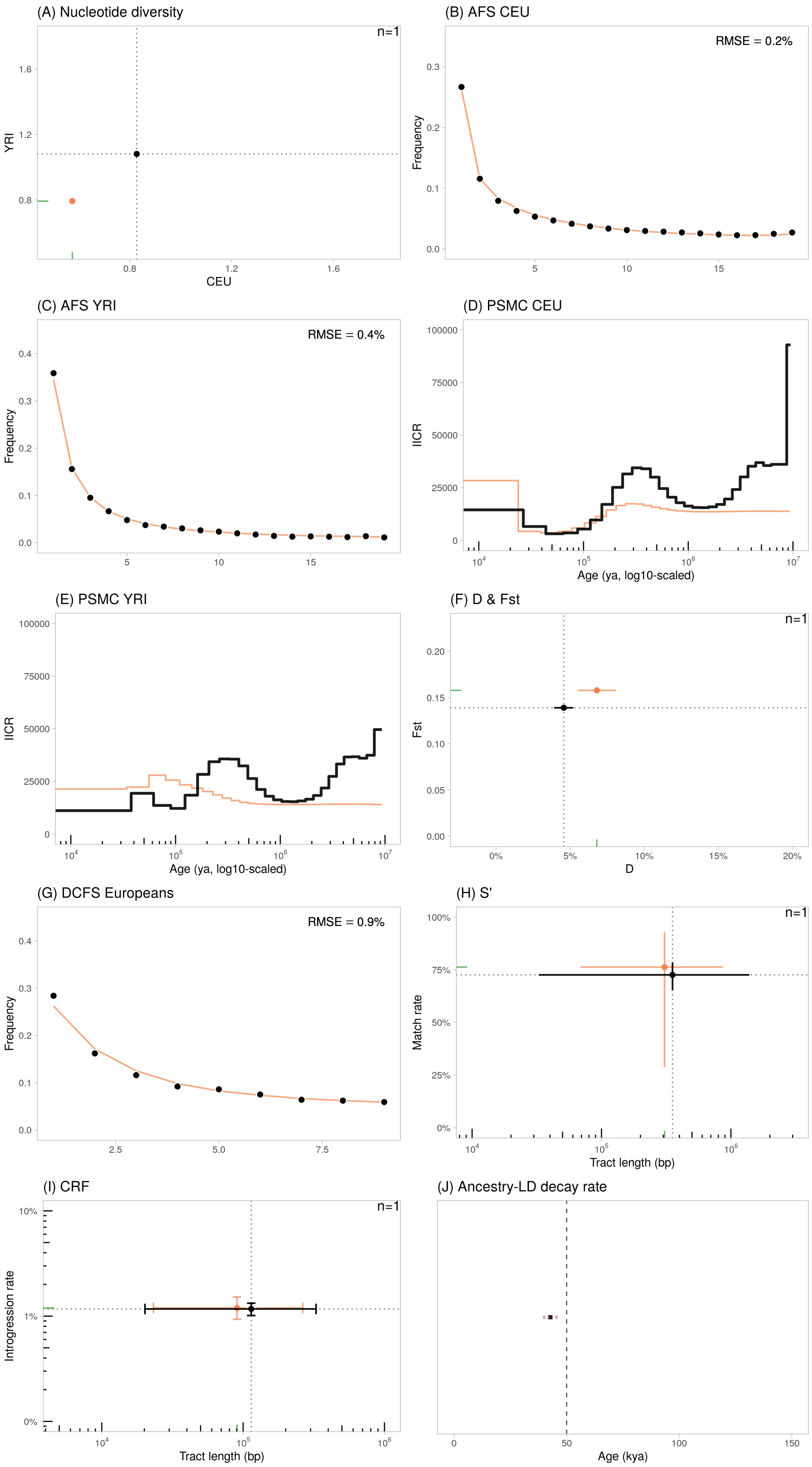

Figure S39: Statistics calculated using the Schaefer et al. (2021) scenario with the standard mutation rate of $1.2 \times 10^{-8}$ b$^{-1}$g$^{-1}$. The error bars represent the confidence intervals at 95% ($= 1.96 \cdot SE$) except for $S'$ (H) and $CRF$ (I) where they represent the 2.5- and 97.5%-percentiles of the distributions.

### S9 Comparison of all models

Table S9: Minimum scaled Euclidean distance (MSED) of the twelve compared models (including ours), under uniform and heterogeneous recombination maps (cf. Notes S1.6). Models are displayed by alphabetical order.

| **Model** | **MSED**, uniform | **MSED**, heterogeneous |
|---|---|---|
| Durvasula et al. (2020) | 1.932 | 1.677 |
| Fu et al. (2014) | 1.091 | 0.989 |
| Gower et al. (2021) | 1.57 | 1.601 |
| Iasi et al. (2021) | 1.373 | 1.152 |
| Jacobs et al. (2019) | 1.469 | 1.303 |
| Kamm et al. (2019) | 1.158 | 1.05 |
| Moorjani et al. (2016) | 0.973 | 0.853 |
| Ragsdale et al. (2019) | 1.631 | 1.615 |
| Schaefer et al. (2021) | 0.684 | 0.525 |
| Skov et al. (2020) | 2.63 | 2.352 |
| Yang et al. (2012) | 3.049 | 4.008 |
| This study | 0.44 | 0.531 |

In the Figure S40 below, we represented the values of the summary statistics estimated on genetic data simulated with $\mu = 1.2 \times 10^{-8}$ bp$^{-1}$g$^{-1}$ for the twelve compared models: the 11 published ones assuming Neanderthal admixture, and our structured model (run ID 1014230). All models were simulated with genomes of twenty chromosomes of 30 Mbp each and a uniform recombination rate ($1.0 \times 10^{-8}$ bp$^{-1}$g$^{-1}$).

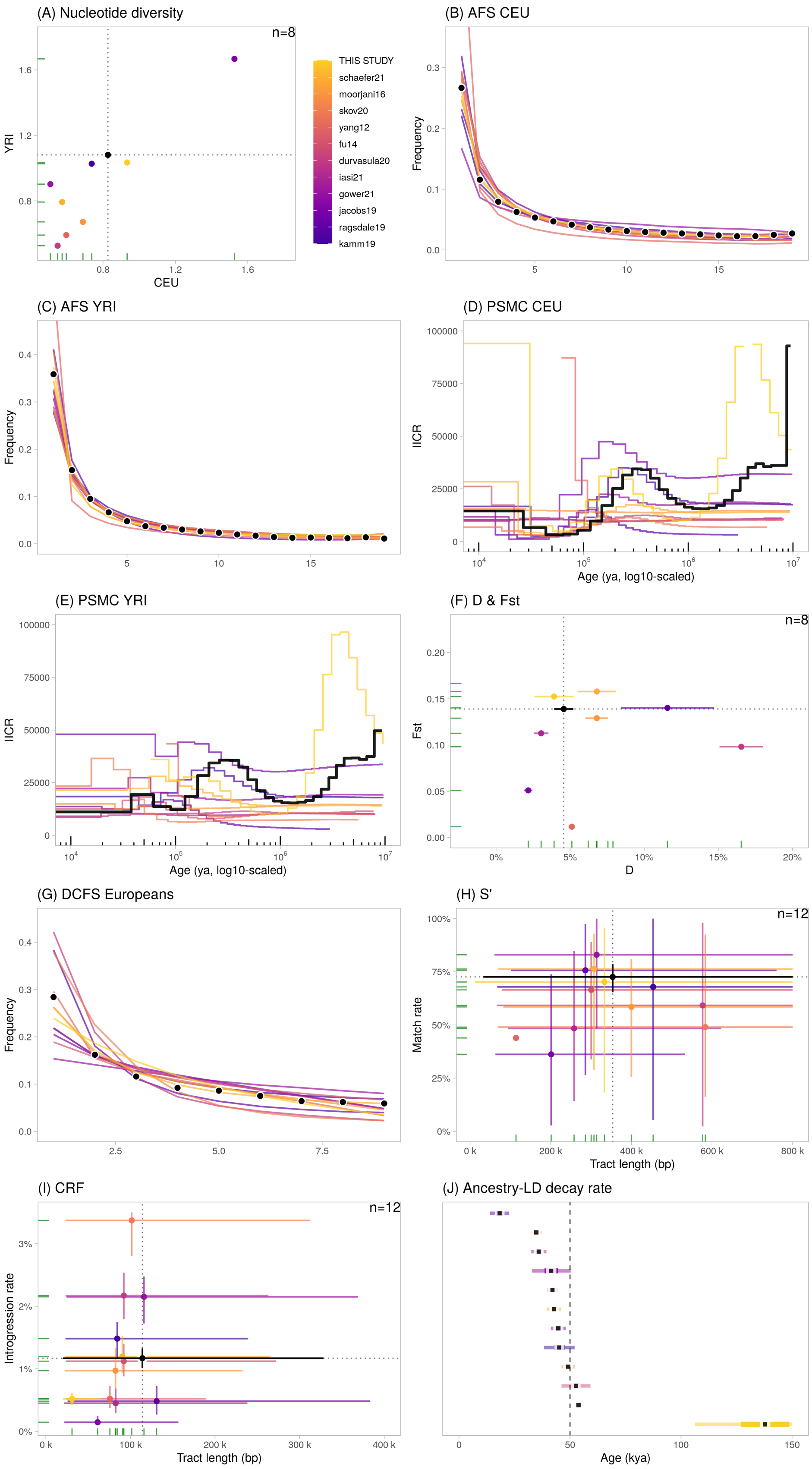

Figure S40: Genetic statistics estimated on the 12 models, including the 11 published models assuming Neanderthal admixture and our structured model (yellow color). The error bars represent the confidence intervals at 95% ($= 1.96 \cdot SE$) except for $S'$ (H) and $CRF$ (I) where they represent the 2.5- and 97.5%-percentiles of the distributions. In (H), the orange dot without bars corresponds to the value estimated under the Yang et al. (2012) model, for which a single segment was detected with $S'$.

### S10 Robustness to statistical parameters and algorithms

Note that in the following subsections, the plots were generated using a single run (ID 1014230) out of the twenty accepted runs from our structured model. We chose to represent the results from a single demographic scenario in order to simplify the comparison but the conclusions hold with the other runs.

#### S10.1 Algorithm to compute population-level ancestry-*LD*

For speed gain and compatibility issues with the installation and use of the `computed` C software (Sankararaman et al., 2012), we re-implemented the method in Python. In order to check the consistency of the estimates obtained with both algorithms, we installed `computed` on a local computer and performed the inference using simulated data.

**Simulations** We simulated 30 genetic datasets under the NGFI model described in Sankararaman et al. (2012), with a modification described hereafter. The original NGFI model assumes three isolated populations (CEU, YRI and Neanderthal) with same size $N_e = 10,000$; CEU diverged from YRI at 2,500 generations BP and Neanderthal diverged from YRI at 10,000 generations BP. There is no population size change in CEU. The modification we made to NGFI is the possibility of Neanderthal admixture into CEU. Admixture ages could vary uniformly between 400 and 2,400 generations ago (i.e. 10 to 60 kya), with rates equal to 0% (no-admixture), 2.5% or 5%. We simulated genomes of size 10×10 Mbp (matching the overall genome size simulated in Sankararaman et al. (2012), i.e. 100 Mbp) with the same mutation and recombination rates as in our other simulations (i.e. $\mu = 1.2 \times 10^{-8}$ and $\rho = 1.0 \times 10^{-8}$). We sampled 50 YRI diploids, 50 CEU diploids (at present) and 1 Vindija Neanderthal diploid (at 2,000 generations BP). The simulated SNPs were then ascertained using the original ascertainment "Scheme 0" (cf. Notes S1.3.2).

**Statistical analysis using lans** We analyzed the ascertained genetic data, restricting the genotypes to the CEU population only, using `computed`. The bounds of the genetic distances were set as default, i.e. from 0.02 cM to 1.0 cM by steps of $10^{-3}$ cM. We estimated the decay constant from the global *LD* decay curve (i.e. summed over all simulated chromosomes).

**Statistical analysis using our implementation** We also analyzed the ascertained genetic data using our Python implementation. The bounds of the genetic distances were the same as for the `computed` analysis. However here, we performed two sub-analyses: (i) "*Original*": this inference framework is based on statistical parameters that best match the `computed` implementation; (ii) "*This study*": this set-up is the one we performed by default for all analyses, and differs from "*Original*" in four ways: (i) we use the unbiased covariance estimate instead of the biased; (ii) the distance bins are defined by their left boundary instead of the right; (iii) the default value for missing covariance estimates in a bin is NaN instead of 0; (iv) we exclude a decay constant if more than half of the bins have missing covariance estimates (to avoid poor fit on noisy curves) (cf. Notes S1.3.2).

**Results** In the case of non-null admixture ($\alpha > 0\%$), `computed` and our Python implementations (using both the "*Original*" or "*This study*" parameter sets) give nearly identical results (Figure S41), in agreement with the simulated admixture ages under unstructured model. The RMSE is 116 years using `computed`, 116 using our Python implementation with "*Original*" set-up and 112 using our Python implementation with "*This study*" set-up. The correlation between the `computed`-inferred ages and ours ("*This study*") is $R^2 > 0.999$ ($P < 10^{-16}$).

When no admixture is simulated ($\alpha = 0$), our Python implementation with "*Original*" set-up gives nearly identical results to `computed` ($R^2 > 0.999$; $P < 10^{-16}$) (Figure S41). The inferred decay constants are highly correlated between `computed` and our Python implementation with the

"*This study*" set-up ($R^2 > 0.966$; $P < 10^{-7}$), but we note that values differed for three specific runs (Figure S41-B). In the absence of admixture, our simulated model is identical to the NGFI model described in Sankararaman et al. (2012) study, where they estimated a decay constant of 8,847 ± 126 generations BP. We were not able to reproduce this value exactly, since some parameters of their simulation studies are missing (mutation and recombination rates, type of recombination landscape). In this study, using the original `computed` software, we estimated decay constants ranging from 2,139 to 6,161 generations BP (53-154 kya with $g = 25$ years) for the 13 runs without admixture, with mean 4,104 (∼103 kya) and standard deviation 1,233 (∼31 ky). There is therefore a high variance for a same model, due to the stochasticity of coalescent simulations and the noisiness of the LD decay curve, which can lead to estimate a wide range of admixture ages even in the absence of any such admixture events.

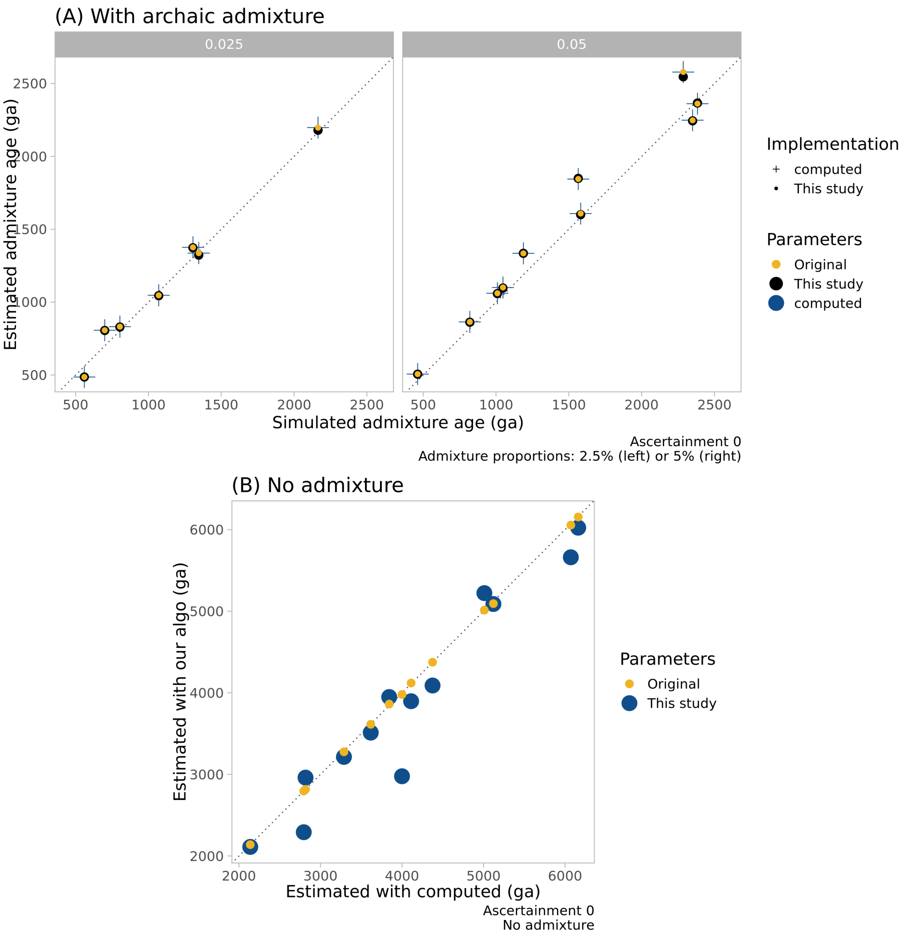

Figure S41: Performance of the estimation of the ancestry-*LD* decay constant (equivalent to the "admixture" age under a panmictic model with single-pulse admixture) between the original software `computed` (Sankararaman et al. (2012), cross-shaped point) and our Python implementation ("*This study*", point shapes). In our Python implementation, we either analyzed the data using the original parameters of `computed` ("*Original*", black color) or the ones we used as default in all our analyses ("*This study*", blue). All reported decay constants are in generations BP. **(A)** Estimated admixture ages compared to the actual simulated values ($x$-axis), for admixture rates equal to 2.5% (left graph) or 5% (right). **(B)** Decay constants estimated on simulations without admixture, using our Python implementation ($y$-axis) vs. `computed` ($x$-axis). Note that in our implementation, when using the same parameter set-up as `computed`, our results are nearly identical to those obtained with `computed` directly, confirming that our implementation performs equivalently to the original software.

#### S10.2 MAC-filtering

In order to match the low-coverage properties of the 1000 Genomes Project data that was used to estimate several genetic statistics in published studies (cf. Notes S6), we artificially impoverished our simulated data in low-frequency SNPs. To do so, we applied a probabilistic filter whereby each original SNP was retained with a probability positively and logarithmically proportional to the original minor allele count (MAC) in the CEU and YRI populations (cf. Notes S1.3.2), as done in Sankararaman et al. (2012). To investigate the robustness of our results to this data filtering, we calculated the same set of summary statistics using a (i) genetic dataset that was processed with the MAC-filtering and (ii) a genetic dataset that was not filtered. The two datasets were obtained under the run ID 1014230 for a genome of size $20 \times 30$ Mb with a uniform recombination map, with the same sampling strategy as described in Methods.

**Results** As illustrated in Fig. S42, the estimates between the MAC-filtered and non-filtered datasets were identical for nearly all statistics, except for the $S'$ analyses for which the mean tract length is $\sim$50% smaller on the non-filtered than the MAC-filtered dataset. Despite this difference, the non-filtered estimate actually provided a better fit to the empirical value compared to the MAC-filtered data (Fig. S42-H, $x$-axis). This comparison suggests that the main conclusions of our work are maintained, irrespective of the biased or non-biased representation of low-frequency SNPs due to low-coverage.

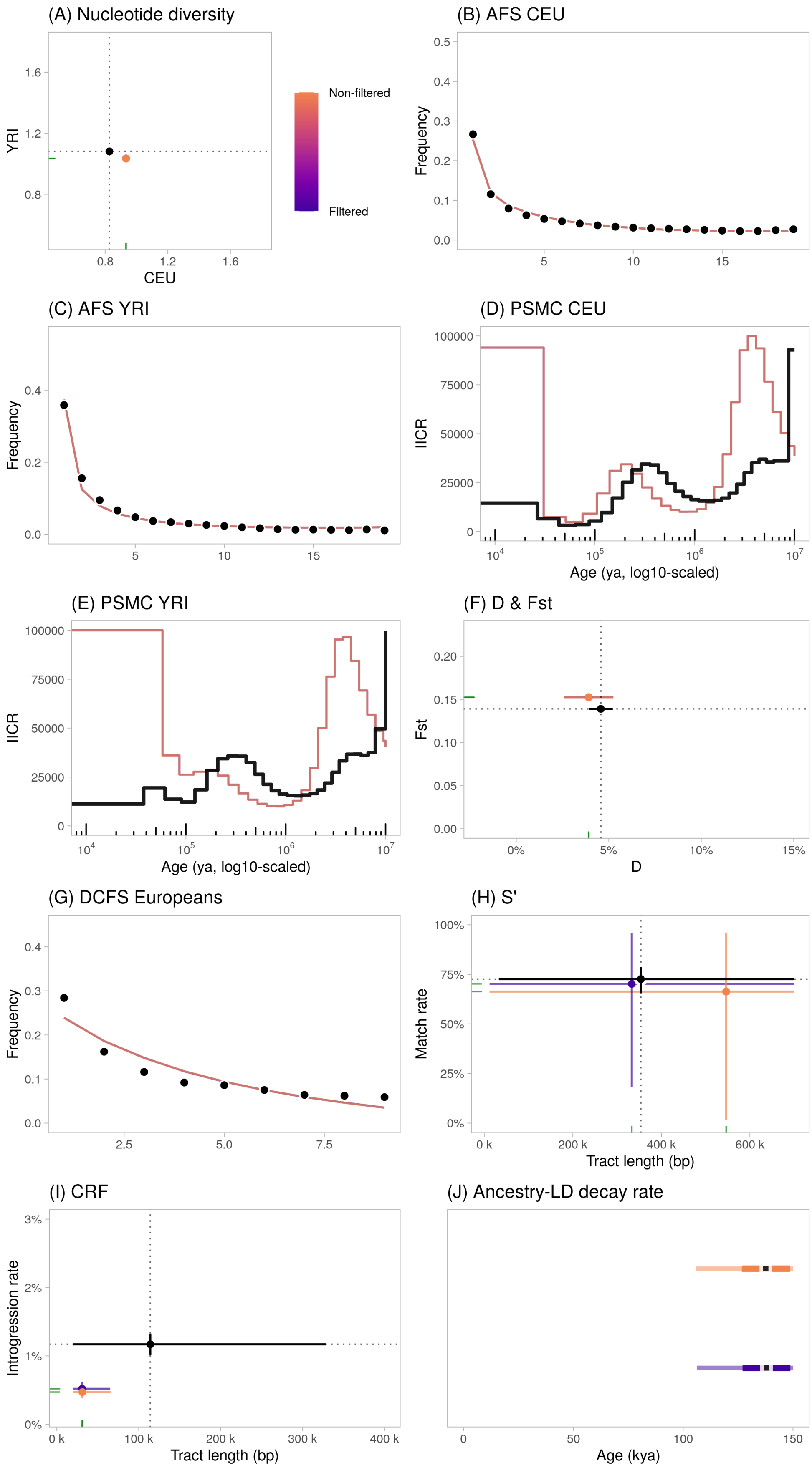

Figure S42: Summary statistics calculated on genetic data ($20 \times 30$ Mbp) simulated under the run 1014230 of our structured model after filtering out the SNPs using a MAC-sensitive thresholding (orange), or not (purple). The error bars represent the confidence intervals at 95% ($= 1.96 \cdot SE$) except for $S'$ (H) and $CRF$ (I) where they represent the 2.5- and 97.5%-percentiles of the distributions.

#### S10.3 Ancestral allele state misspecification

In several observed genetic data, including the 1000 Genomes Project database, the identification of the ancestral allele ("allele polarization") is conditioned on the reliability of genome alignments and the depth of the phylogenetic divergence with the outgroup species (Keightley and Jackson, 2018; Zeng et al., 2019). Previous studies have observed for instance that polarization error are particularly strong at hypermutable CpG sites. This ancestral allele misspecification can bias the estimation of summary statistics relying on allele polarization, like the derived allele frequency spectra. For instance, when ancestral states are misspecified, the $AFS$ can be artificially enriched in very high-frequency variants since in some cases, a (truly) singleton derived variant will appear as a singleton (falsely) ancestral variant, thus as a derived allele at high frequency in the target population (Williamson et al., 2005). To investigate how robust were our results to ancestral allele misspecification, we simulated genetic data ($20 \times 30$ Mbp) under the run 1014230 of our structured model. Then, we generated an additional dataset, after applying polarization errors with a rate of 3% to the previous data. Previous studies estimated polarization error rate on the order of ∼1%—4% in the 1000 Genomes data (Glémin et al., 2015).

Practically, to model ancestral allele misspecification, noting $\epsilon$ the ancestral polarization error rate (i.e. the probability that the ancestral allele is misspecified), we reversed alleles in the non-outgroup genotypes for each SNP with probability $\epsilon$. In the modified `EIGENSTRAT` format we used, diploid genotypes are encoded as 0 for homozygous ancestral, 1 for heterozygous and 2 for homozygous derived. Thus, we calculated the reverse of any genotype $G$ as $G' = 2 - G$, while leaving the ougroup reference genotype $G_{ref}$ as $G_{ref} = 0$.

**Results** As illustrated in Fig. S43, the estimates are identical for nearly all statistics between the original dataset and the one with 3% ancestral misspecification. We note that there is a slight difference in the $AFS$ for the maximum-frequency variants, with slight increase of this category for the 3%-misspecified data (as expected) (Fig. S43-C). This comparison suggests that the main conclusions of our work are maintained, irrespective of the level of ancestral allele misspecification, in the limits of a reasonable polarization error rate (Glémin et al., 2015).

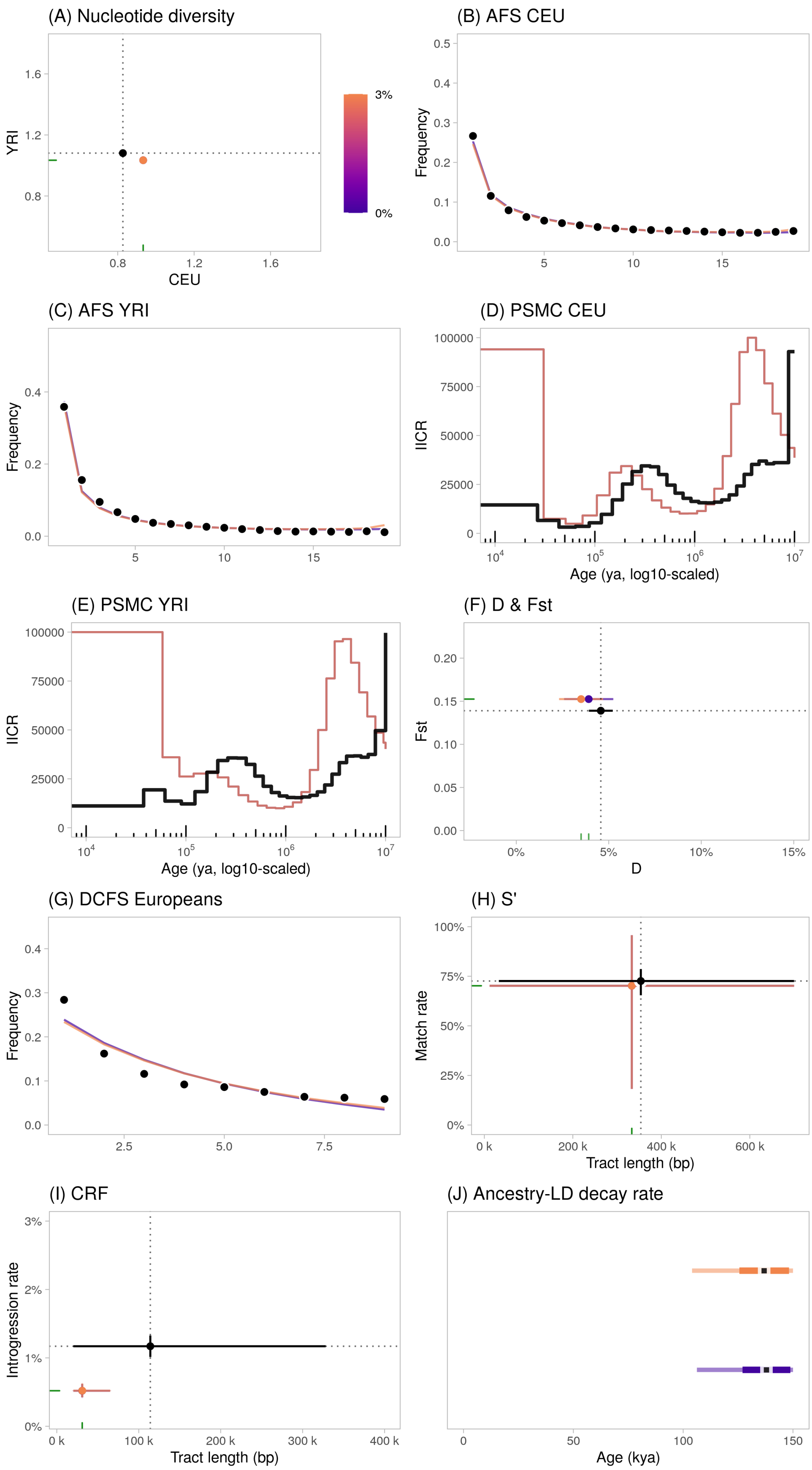

Figure S43: Summary statistics calculated on genetic data ($20 \times 30$ Mbp) simulated assuming 3% of ancestral allele misspecification (orange), or perfect ancestral polarization (purple). The error bars represent the confidence intervals at 95% ($= 1.96 \cdot SE$) except for $S'$ (H) and $CRF$ (I) where they represent the 2.5- and 97.5%-percentiles of the distributions.

#### S10.4 Empirical recombination maps

The landscape of human recombinations (the average density of crossing-overs along chromosomes across time) is heterogeneous (Halldorsson et al., 2019). Several statistics used for demographic inference and archaic introgression analysis are function of LD and therefore can depend on the type of recombination landscape used in the simulations. To investigate the sensitivity of our results to the properties of the recombination map, we used the run 1014230 of our structured model and simulated genetic data using empirical human recombination maps. We considered twenty human autosomes with equal size (50 Mbp). To this end, we trimmed the segments at both ends of each chromosome for which recombination rates were either missing or non-varying (because likely extrapolated). We then truncated the right-end of the trimmed sequences to a maximum size per chromosome of 50 Mb (Table S10). We estimated the local recombination rates along the chromosomes by calculating the mean variation in genetic positions per physical base pairs between two successive positions of the reference maps. If we note $p_i$ and $p_j$ the physical positions (in base pairs) of sites $i$ and $j$ and $g_i$ and $g_j$ their respective genetic positions (in cM), and if we assume that all four variables are defined, the local recombination rate was calculated as $\rho_{ij} = 10^6 \cdot \frac{g_j - g_i}{b_p - b_i}$ cM/Mb. We considered two genetic maps lifted over the GRCh37 assembly. The two are population-wise LD-based genetic maps where recombination rates were inferred from the observed breakdown of LD between markers. The LD-based approaches, when compared to directly observed patterns of crossover in pedigree data, were found to provide a robust estimate of the historical proportion of meioses over thousands of generations of evolution (Kong et al., 2010).

- "HapMap", the GRCh37-lifted map of HapMap II (build 35), inferred using the `LDhat` method using genotypes from CEU, YRI and JPT+CHB unrelated individuals (Frazer et al., 2007) (downloaded[8] on Jan 12th 2022).
- "Spence19": the GRCh37-map inferred from the 26 populations of the 1000 Genomes Project, inferred using the demography-aware `pyrho` method (Spence and Song, 2019) (downloaded[9] on Jan 12th 2022).

[8] ftp://ftp-trace.ncbi.nih.gov/1000genomes/ftp/technical/working/20110106_recombination_hotspots
[9] https://drive.google.com/drive/folders/1Tgt_7GsDO0-o02vcYSfwqHFd3JNF6R06

Table S10: Location of the chromosome sequences that were used for the simulated data with empirical recombination maps. All positions are in Mbp on the original GRCh37 assembly.
! End position when retaining only 50 Mbp.
* Positions obtained after trimming the starts and ends with null or missing recombination rates.

| | HapMap | | | Spence19 | | |
|---|---|---|---|---|---|---|
| Chromosome | Start* | End* | Trunc. End*! | Start* | End* | Trunc. End*! |
| 1 | 0.06 | 249.22 | 50.06 | 0.01 | 249.24 | 50.01 |
| 2 | 0.01 | 243.09 | 50.01 | 0.01 | 243.19 | 50.01 |
| 3 | 0.06 | 197.87 | 50.06 | 0.06 | 197.96 | 50.06 |
| 4 | 0.01 | 191.03 | 50.01 | 0.01 | 191.04 | 50.01 |
| 5 | 0.02 | 180.72 | 50.02 | 0.01 | 180.9 | 50.01 |
| 6 | 0.09 | 171.05 | 50.09 | 0.06 | 171.05 | 50.06 |
| 7 | 0.04 | 159.13 | 50.04 | 0.01 | 159.13 | 50.01 |
| 8 | 0.16 | 146.3 | 50.16 | 0.01 | 146.3 | 50.01 |
| 9 | 0.04 | 141.11 | 50.04 | 0.01 | 141.15 | 50.01 |
| 10 | 0.07 | 135.5 | 50.07 | 0.06 | 135.52 | 50.06 |
| 11 | 0.2 | 134.95 | 50.2 | 0.09 | 134.95 | 50.09 |
| 12 | 0.15 | 133.78 | 50.15 | 0.06 | 133.84 | 50.06 |
| 13 | 19.02 | 115.11 | 69.02 | 19.02 | 115.11 | 69.02 |
| 14 | 19.62 | 107.29 | 69.62 | 19 | 107.29 | 69 |
| 15 | 20.01 | 102.51 | 70.01 | 20 | 102.52 | 70 |
| 16 | 0.08 | 90.16 | 50.08 | 0.06 | 90.29 | 50.06 |
| 17 | 0.01 | 81.05 | 50.01 | 0 | 81.19 | 50 |
| 18 | 0.01 | 78.02 | 50.01 | 0.01 | 78.02 | 50.01 |
| 19 | 0.25 | 59.1 | 50.25 | 0.06 | 59.12 | 50.06 |
| 20 | 0.06 | 62.95 | 50.06 | 0.06 | 62.97 | 50.06 |

The mean recombination rates calculated across the retained maps are: $1.41\times10^{-8}$ $bp^{-1}g^{-1}$ for "HapMap" and $1.04\times10^{-8}$ $bp^{-1}g^{-1}$ for "Spence19".

**Results** The simulations performed using the "HapMap" or the "Spence19" genetic maps lead to nearly identical results on most statistics, except for the $D$-statistic and, as expected, the LD-based statistics: *S'*, *CRF* and $T_{LD}$. The $D$-statistic is slightly (and not significantly) lower under the empirical maps compared to the uniform map or even the hotspot-simulated map (cf. Notes S1.2). The difference could be due to a stochastic variation of the statistical values due to limited genome size rather than a direct consequence of the different maps. Compared to the uniform map, empirical maps lead to more than ~1.5-times longer *CRF* segments in CEU, and higher *CRF* introgression rates. This enrichment in longer segments (which would be interpreted as evidence of even more recent Neanderthal admixture into *Hs*) is expected under variable recombination rates, and driven by the existence of large regions with lower probability of recombination along the genome. Surprisingly, although the empirical maps have the same effect in terms of match rate/introgression rate, they have an opposite effect in terms of segment lengths when inferred with *S'* (compared to *CRF*). For *S'*, empirical maps lead to ~2-times shorter *S'* segments. In the case of $T_{LD}$, the "HapMap" empirical map leads to slightly lower ages compared to the other maps, including the uniform one.

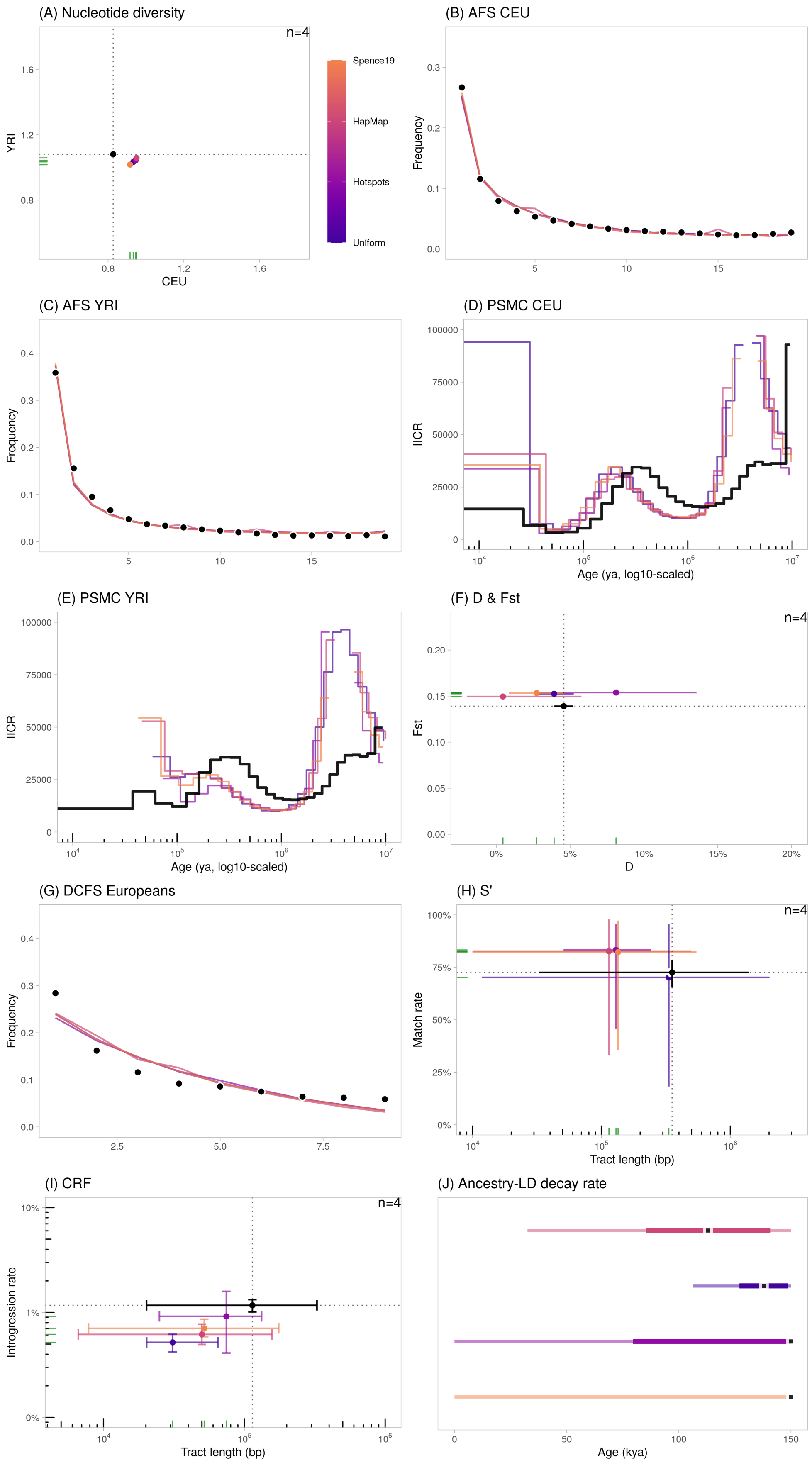

Figure S44: Summary statistics calculated on genetic data simulated with human empirical genetic maps ("Spence19", "HapMap"), simulated artificial heterogeneous map with hotspots ("Hotspots") and uniform map ("Uniform"). The error bars represent the confidence intervals at 95% ($= 1.96 \cdot SE$) except for $S'$ (H) and $CRF$ (I) where they represent the 2.5- and 97.5%-percentiles of the distributions.

#### S10.5 Neanderthal sampling times

We investigated the robustness of our results to the age of the Neanderthal sample that was simulated. The $^{14}C$ radiocarbon dating of observed Neanderthal genomes varies widely, with the Vindija33.19 genome dated ∼55 kya (Prüfer et al., 2017) and the Altai Neanderthal ∼115 kya (Prüfer et al., 2014). The observed statistics that we considered in this study have been produced using either one of the two Neanderthal samples in the original studies (Table S6). By default, in all our simulations, we sampled the Neanderthal genome 2,000 generations ago, corresponding to a sampling age of 50—58 kya if we assume a generation time of 25 or 29 years respectively (i.e. this would correspond to Vindija33.19 Neanderthal). In Fig. S45, we represented the statistics calculated for a sample living 2,000 generations BP, as well as for another one living 5,200 generations BP (130 kya for $g = 25$ years, i.e. corresponding to Altai Neanderthal).

**Results** The results obtained when sampling Neanderthal 2,000 generations ago vs. 5,200 generations ago are identical for all statistics (Fig. S45) which is not surprising given the very high level of genetic drift caused by the low Neanderthal population size in the accepted models (S7). As a conclusion, our results are robust to the sampling age of Neanderthal between ∼50 kya and ∼130 kya.

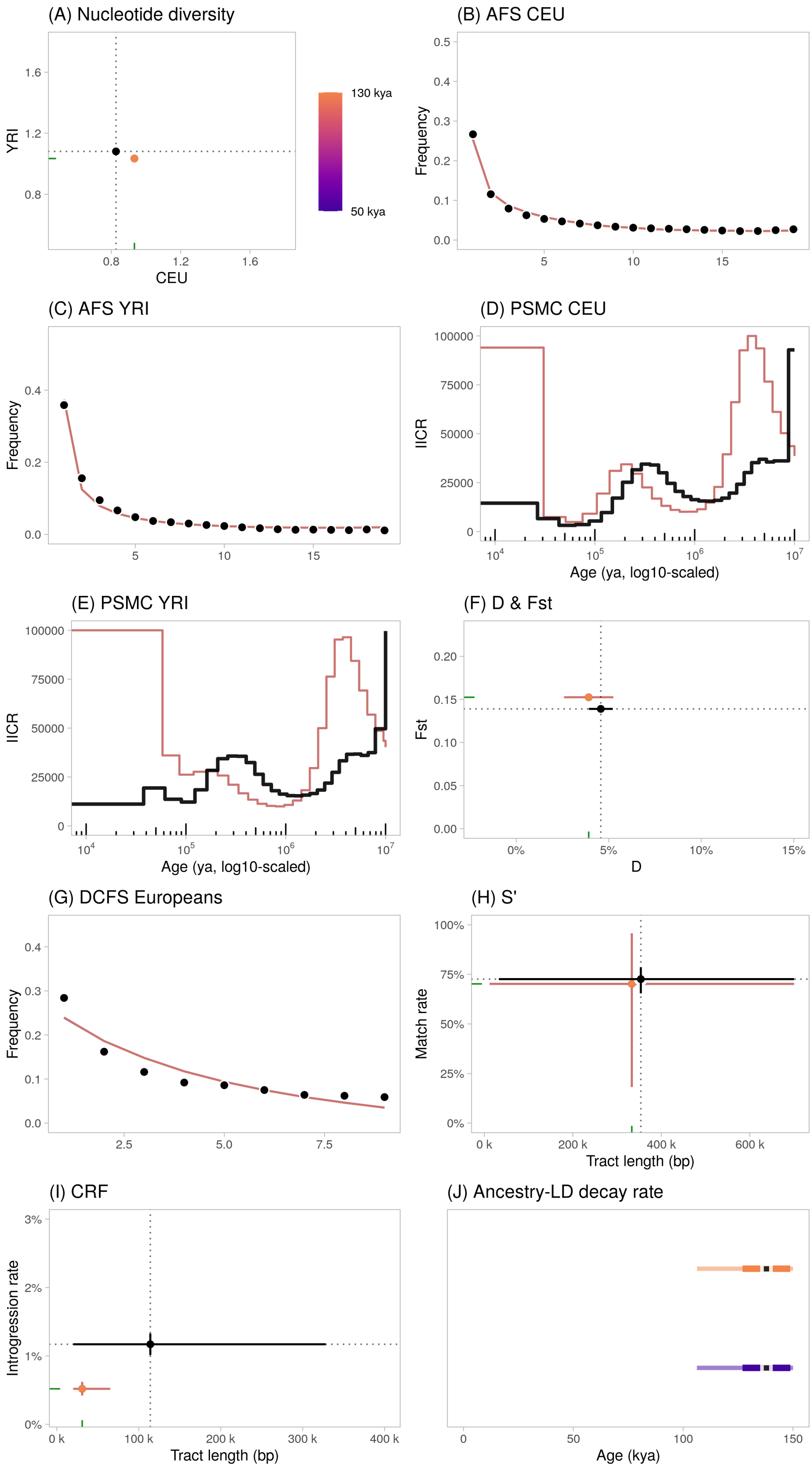

Figure S45: Summary statistics calculated on genetic data simulated with the Neanderthal genome sampled either at 50 kya (default, Vindija33.19) or 130 kya (Altai), assuming $g = 25$ years. The error bars represent the confidence intervals at 95% ($= 1.96 \cdot SE$) except for $S'$ (H) and $CRF$ (I) where they represent the 2.5- and 97.5%-percentiles of the distributions.

#### S10.6 Pseudodiploidization

To assess the robustness of some statistical estimates to genotype quality (i.e. diploid vs. pseudodiploid), we analyzed the genetic data simulated under the run 1014230 of our structured model (20×30 Mbp with uniform recombination rate) after pseudodiploidizing the heterozygous genotypes for every samples (modern humans as well as Neanderthals): i.e. we randomly sampled one of the two alleles for each genotype independently and set the sampled allele into two copies. We investigated the robustness on the two statistics—$D$-statistics (Green et al., 2010) and $T_{LD}$, the population-level ancestry-$LD$ decay constant (Sankararaman et al., 2012)—for which some empirical samples could have been analyzed as pseudodiploid in previous studies. We note that pseudodiploidization would not have been relevant or even possible for approaches which rely on heterozygous or phased genotypes (e.g. *AFS*, *DCFS*, *PSMC*, *CRF*).

**Results** The results for the $D$-statistics and the ancestry-$LD$ decay constant $T_{LD}$ were very similar (cf. Table S11), suggesting that the results for these two statistics are robust to pseudodiploidization.

Table S11: Comparison of the $D$ and $T_{LD}$ values when analyzing diploid vs. pseudodiploid genotypes.

| | Diploid | Pseudodiploid |
|---|---|---|
| $D$-statistics | 3.9% ± 6.8%<br>Z=5.77 | 3.9% ± 6.8%<br>Z=5.69 |
| $T_{LD}$<br>(generations BP) | 5,506 ± 649 | 5,795 ± 880 |

Durbin, R. M., Altshuler, D., Durbin, R. M., Abecasis, G. R., Bentley, D. R., Chakravarti, A., Clark, A. G., Collins, F. S., De La Vega, F. M., Donnelly, P., Egholm, M., Flicek, P., Gabriel, S. B., Gibbs, R. A., Knoppers, B. M., Lander, E. S., Lehrach, H., Mardis, E. R., McVean, G. A., Nickerson, D. A., Peltonen, L., Schafer, A. J., Sherry, S. T., Wang, J., Wilson, R. K., Gibbs, R. A., Deiros, D., Metzker, M., Muzny, D., Reid, J., Wheeler, D., Wang, J., Li, J., Jian, M., Li, G., Li, R., Liang, H., Tian, G., Wang, B., Wang, J., Wang, W., Yang, H., Zhang, X., Zheng, H., Lander, E. S., Altshuler, D., Ambrogio, L., Bloom, T., Cibulskis, K., Fennell, T. J., Gabriel, S. B., Jaffe, D. B., Shefler, E., Sougnez, C. L., Bentley, D. R., Gormley, N., Humphray, S., Kingsbury, Z., Kokko-Gonzales, P., Stone, J., McKernan, K. J., Costa, G. L., Ichikawa, J. K., Lee, C. C., Sudbrak, R., Lehrach, H., Borodina, T. A., Dahl, A., Davydov, A. N., Marquardt, P., Mertes, F., Nietfeld, W., Rosenstiel, P., Schreiber, S., Soldatov, A. V., Timmermann, B., Tolzmann, M., Egholm, M., Affourtit, J., Ashworth, D., Attiya, S., Bachorski, M., Buglione, E., Burke, A., Caprio, A., Celone, C., Clark, S., Conners, D., Desany, B., Gu, L., Guccione, L., Kao, K., Kebbel, A., Knowlton, J., Labrecque, M., McDade, L., Mealmaker, C., Minderman, M., Nawrocki, A., Niazi, F., Pareja, K., Ramenani, R., Riches, D., Song, W., Turcotte, C., Wang, S., Mardis, E. R., Wilson, R. K., Dooling, D., Fulton, L., Fulton, R., Weinstock, G., Durbin, R. M., Burton, J., Carter, D. M.,

Churcher, C., Coffey, A., Cox, A., Palotie, A., Quail, M., Skelly, T., Stalker, J., Swerdlow, H. P., Turner, D., De Witte, A., Giles, S., Gibbs, R. A., Wheeler, D., Bainbridge, M., Challis, D., Sabo, A., Yu, F., Yu, J., Wang, J., Fang, X., Guo, X., Li, R., Li, Y., Luo, R., Tai, S., Wu, H., Zheng, H., Zheng, X., Zhou, Y., Li, G., Wang, J., Yang, H., Marth, G. T., Garrison, E. P., Huang, W., Indap, A., Kural, D., Lee, W.-P., Fung Leong, W., Quinlan, A. R., Stewart, C., Stromberg, M. P., Ward, A. N., Wu, J., Lee, C., Mills, R. E., Shi, X., Daly, M. J., DePristo, M. A., Altshuler, D., Ball, A. D., Banks, E., Bloom, T., Browning, B. L., Cibulskis, K., Fennell, T. J., Garimella, K. V., Grossman, S. R., Handsaker, R. E., Hanna, M., Hartl, C., Jaffe, D. B., Kernytsky, A. M., Korn, J. M., Li, H., Maguire, J. R., McCarroll, S. A., McKenna, A., Nemesh, J. C., Philippakis, A. A., Poplin, R. E., Price, A., Rivas, M. A., Sabeti, P. C., Schaffner, S. F., Shefler, E., Shlyakhter, I. A., Cooper, D. N., Ball, E. V., Mort, M., Phillips, A. D., Stenson, P. D., Sebat, J., Makarov, V., Ye, K., Yoon, S. C., Bustamante, C. D., Clarke, L., Flicek, P., Cunningham, F., Herrero, J., Keenen, S., Kulesha, E., Leinonen, R., McLaren, W. M., Radhakrishnan, R., Smith, R. E., Zalunin, V., Zheng-Bradley, X., Korbel, J. O., Stütz, A. M., Humphray, S., Bauer, M., Keira Cheetham, R., Cox, T., Eberle, M., James, T., Kahn, S., Murray, L., Chakravarti, A., Ye, K., De La Vega, F. M., Fu, Y., Hyland, F. C. L., Manning, J. M., McLaughlin, S. F., Peckham, H. E., Sakarya, O., Sun, Y. A., Tsung, E. F., Batzer, M. A., Konkel, M. K., Walker, J. A., Sudbrak, R., Albrecht, M. W., Amstislavskiy, V. S., Herwig, R., Parkhomchuk, D. V., Sherry, S. T., Agarwala, R., Khouri, H. M., Morgulis, A. O., Paschall, J. E., Phan, L. D., Rotmistrovsky, K. E., Sanders, R. D., Shumway, M. F., Xiao, C., McVean, G. A., Auton, A., Iqbal, Z., Lunter, G., Marchini, J. L., Moutsianas, L., Myers, S., Tumian, A., Desany, B., Knight, J., Winer, R., Craig, D. W., Beckstrom-Sternberg, S. M., Christoforides, A., The 1000 Genomes Project Consortium, Corresponding author, Steering committee, Production group: Baylor College of Medicine, BGI-Shenzhen, Broad Institute of MIT and Harvard, Illumina, Life Technologies, Max Planck Institute for Molecular Genetics, Roche Applied Science, Washington University in St Louis, Wellcome Trust Sanger Institute, Analysis group: Agilent Technologies, Baylor College of Medicine, Boston College, Brigham and Women's Hospital, Cardiff University, T. H. G. M. D., Cold Spring Harbor Laboratory, Cornell and Stanford Universities, European Bioinformatics Institute, European Molecular Biology Laboratory, Johns Hopkins University, Leiden University Medical Center, Louisiana State University, US National Institutes of Health, Oxford University, and The Translational Genomics Research Institute (2010). A map of human genome variation from population-scale sequencing. *Nature*, 467(7319):1061–1073.

Durvasula, A. and Sankararaman, S. (2020). Recovering signals of ghost archaic introgression in African populations. *Science Advances*, 6(7):eaax5097.

Frazer, K. A., Ballinger, D. G., Cox, D. R., Hinds, D. A., Stuve, L. L., Gibbs, R. A., Belmont, J. W., Boudreau, A., Hardenbol, P., Leal, S. M., Pasternak, S., Wheeler, D. A., Willis, T. D., Yu, F., Yang, H., Zeng, C., Gao, Y., Hu, H., Hu, W., Li, C., Lin, W., Liu, S., Pan, H., Tang, X., Wang, J., Wang, W., Yu, J., Zhang, B., Zhang, Q., Zhao, H., Zhao, H., Zhou, J., Gabriel, S. B., Barry, R., Blumenstiel, B., Camargo, A., Defelice, M., Faggart, M., Goyette, M., Gupta, S., Moore, J., Nguyen, H., Onofrio, R. C., Parkin, M., Roy, J., Stahl, E., Winchester, E., Ziaugra, L., Altshuler, D., Shen, Y., Yao, Z., Huang, W., Chu, X., He, Y., Jin, L., Liu, Y., Shen, Y., Sun, W., Wang, H., Wang, Y., Wang, Y., Xiong, X., Xu, L., Waye, M. M. Y., Tsui, S. K. W., Xue, H., Wong, J. T.-F., Galver, L. M., Fan, J.-B., Gunderson, K., Murray, S. S., Oliphant, A. R., Chee, M. S., Montpetit, A., Chagnon, F., Ferretti, V., Leboeuf, M., Olivier, J.-F., Phillips, M. S., Roumy, S., Sallée, C., Verner, A., Hudson, T. J., Kwok, P.-Y., Cai, D., Koboldt, D. C., Miller, R. D., Pawlikowska, L., Taillon-Miller, P., Xiao, M., Tsui, L.-C., Mak, W., Qiang Song, Y., Tam, P. K. H., Nakamura, Y., Kawaguchi, T., Kitamoto, T., Morizono, T., Nagashima, A., Ohnishi, Y., Sekine, A., Tanaka, T., Tsunoda, T., Deloukas, P., Bird, C. P., Delgado, M., Dermitzakis, E. T., Gwilliam, R., Hunt, S., Morrison, J., Powell, D., Stranger, B. E., Whittaker, P., Bentley, D. R., Daly, M. J., de Bakker, P. I. W., Barrett, J., Chretien, Y. R., Maller, J., McCarroll, S., Patterson, N., Pe'er, I., Price, A., Purcell, S., Richter, D. J., Sabeti, P., Saxena, R., Schaffner, S. F., Sham, P. C., Varilly, P., Altshuler, D., Stein, L. D., Krishnan, L., Vernon Smith, A., Tello-Ruiz, M. K., Thorisson, G. A., Chakravarti, A., Chen, P. E., Cutler, D. J., Kashuk, C. S., Lin, S., Abecasis,

G. R., Guan, W., Li, Y., Munro, H. M., Steve Qin, Z., Thomas, D. J., McVean, G., Auton, A., Bottolo, L., Cardin, N., Eyheramendy, S., Freeman, C., Marchini, J., Myers, S., Spencer, C., Stephens, M., Donnelly, P., Cardon, L. R., Clarke, G., Evans, D. M., Morris, A. P., Weir, B. S., Tsunoda, T., Johnson, T., Mullikin, J. C., Sherry, S. T., Feolo, M., Skol, A., Zhang, H., Zeng, C., Zhao, H., Matsuda, I., Fukushima, Y., Macer, D. R., Suda, E., Rotimi, C. N., Adebamowo, C. A., Ajayi, I., Aniagwu, T., Marshall, P. A., Nkwodimmah, C., Royal, C. D. M., Leppert, M. F., Dixon, M., Peiffer, A., Qiu, R., Kent, A., Kato, K., Niikawa, N., Adewole, I. F., Knoppers, B. M., Foster, M. W., Wright Clayton, E., Watkin, J., Gibbs, R. A., Belmont, J. W., Muzny, D., Nazareth, L., Sodergren, E., Weinstock, G. M., Wheeler, D. A., Yakub, I., Gabriel, S. B., Onofrio, R. C., Richter, D. J., Ziaugra, L., Birren, B. W., Daly, M. J., Altshuler, D., Wilson, R. K., Fulton, L. L., Rogers, J., Burton, J., Carter, N. P., Clee, C. M., Griffiths, M., Jones, M. C., McLay, K., Plumb, R. W., Ross, M. T., Sims, S. K., Willey, D. L., Chen, Z., Han, H., Kang, L., Godbout, M., Wallenburg, J. C., L'Archevêque, P., Bellemare, G., Saeki, K., Wang, H., An, D., Fu, H., Li, Q., Wang, Z., Wang, R., Holden, A. L., Brooks, L. D., McEwen, J. E., Guyer, M. S., Ota Wang, V., Peterson, J. L., Shi, M., Spiegel, J., Sung, L. M., Zacharia, L. F., Collins, F. S., Kennedy, K., Jamieson, R., The International HapMap Consortium, Genotyping centres: Perlegen Sciences, Baylor College of Medicine and ParAllele BioScience, Beijing Genomics Institute, Broad Institute of Harvard and Massachusetts Institute of Technology, Chinese National Human Genome Center at Beijing, Chinese National Human Genome Center at Shanghai, Chinese University of Hong Kong, Hong Kong University of Science and Technology, Illumina, McGill University and Génome Québec Innovation Centre, University of California at San Francisco and Washington University, University of Hong Kong, University of Tokyo and RIKEN, Wellcome Trust Sanger Institute, Analysis groups: Broad Institute, Cold Spring Harbor Laboratory, Johns Hopkins University School of Medicine, University of Michigan, University of Oxford, University of Oxford, W. T. C. f. H. G., RIKEN, US National Institutes of Health, US National Institutes of Health National Center for Biotechnology Information, Community engagement/public consultation and sample collection groups: Beijing Normal University and Beijing Genomics Institute, Health Sciences University of Hokkaido, and Shinshu University, E. E. I., Howard University and University of Ibadan, University of Utah, Ethical, l. a. s. i. C. A. o. S. S., Genetic Interest Group, Kyoto University, Nagasaki University, University of Ibadan School of Medicine, University of Montréal, University of Oklahoma, Vanderbilt University, Wellcome Trust, SNP discovery: Baylor College of Medicine, Washington University, Scientific management: Chinese Academy of Sciences, Genome Canada, Génome Québec, Japanese Ministry of Education, Sports, S. a. T. C., Ministry of Science and Technology of the People's Republic of China, The Human Genetic Resource Administration of China, and The SNP Consortium (2007). A second generation human haplotype map of over 3.1 million SNPs. *Nature*, 449(7164):851–861.

Fu, Q., Hajdinjak, M., Moldovan, O. T., Constantin, S., Mallick, S., Skoglund, P., Patterson, N., Rohland, N., Lazaridis, I., Nickel, B., Viola, B., Prüfer, K., Meyer, M., Kelso, J., Reich, D., and Pääbo, S. (2015). An early modern human from Romania with a recent Neanderthal ancestor. *Nature*, 524(7564):216–219.

Fu, Q., Li, H., Moorjani, P., Jay, F., Slepchenko, S. M., Bondarev, A. A., Johnson, P. L. F., Aximu-Petri, A., Prüfer, K., de Filippo, C., Meyer, M., Zwyns, N., Salazar-García, D. C., Kuzmin, Y. V., Keates, S. G., Kosintsev, P. A., Razhev, D. I., Richards, M. P., Peristov, N. V., Lachmann, M., Douka, K., Higham, T. F. G., Slatkin, M., Hublin, J.-J., Reich, D., Kelso, J., Viola, T. B., and Pääbo, S. (2014). Genome sequence of a 45,000-year-old modern human from western Siberia. *Nature*, 514(7523):445–449.

Glémin, S., Arndt, P. F., Messer, P. W., Petrov, D., Galtier, N., and Duret, L. (2015). Quantification of GC-biased gene conversion in the human genome. *Genome Research*, 25(8):1215–1228.

Gower, G., Picazo, P. I., Fumagalli, M., and Racimo, F. (2021). Detecting adaptive introgression in human evolution using convolutional neural networks. *eLife*, 10:e64669.

Gravel, S., Henn, B. M., Gutenkunst, R. N., Indap, A. R., Marth, G. T., Clark, A. G., Yu, F.,

Gibbs, R. A., 1000 Genomes Project, and Bustamante, C. D. (2011). Demographic history and rare allele sharing among human populations. *Proceedings of the National Academy of Sciences of the United States of America*, 108(29):11983–11988.

Green, R. E., Krause, J., Briggs, A. W., Maricic, T., Stenzel, U., Kircher, M., Patterson, N., Li, H., Zhai, W., Fritz, M. H.-Y., Hansen, N. F., Durand, E. Y., Malaspinas, A.-S., Jensen, J. D., Marques-Bonet, T., Alkan, C., Prüfer, K., Meyer, M., Burbano, H. A., Good, J. M., Schultz, R., Aximu-Petri, A., Butthof, A., Höber, B., Höffner, B., Siegemund, M., Weihmann, A., Nusbaum, C., Lander, E. S., Russ, C., Novod, N., Affourtit, J., Egholm, M., Verna, C., Rudan, P., Brajkovic, D., Kucan, Ž., Gušic, I., Doronichev, V. B., Golovanova, L. V., Lalueza-Fox, C., de la Rasilla, M., Fortea, J., Rosas, A., Schmitz, R. W., Johnson, P. L. F., Eichler, E. E., Falush, D., Birney, E., Mullikin, J. C., Slatkin, M., Nielsen, R., Kelso, J., Lachmann, M., Reich, D., and Pääbo, S. (2010). A Draft Sequence of the Neandertal Genome. *Science*, 328(5979):710–722.

Gutenkunst, R. N., Hernandez, R. D., Williamson, S. H., and Bustamante, C. D. (2009). Inferring the Joint Demographic History of Multiple Populations from Multidimensional SNP Frequency Data. *PLOS Genetics*, 5(10):e1000695.

Halldorsson, B. V., Palsson, G., Stefansson, O. A., Jonsson, H., Hardarson, M. T., Eggertsson, H. P., Gunnarsson, B., Oddsson, A., Halldorsson, G. H., Zink, F., Gudjonsson, S. A., Frigge, M. L., Thorleifsson, G., Sigurdsson, A., Stacey, S. N., Sulem, P., Masson, G., Helgason, A., Gudbjartsson, D. F., Thorsteinsdottir, U., and Stefansson, K. (2019). Characterizing mutagenic effects of recombination through a sequence-level genetic map. *Science*, 363(6425):eaau1043.

Hellenthal, G. and Stephens, M. (2007). msHOT: Modifying Hudson's ms simulator to incorporate crossover and gene conversion hotspots. *Bioinformatics*, 23(4):520–521.

Hinch, A. G., Tandon, A., Patterson, N., Song, Y., Rohland, N., Palmer, C. D., Chen, G. K., Wang, K., Buxbaum, S. G., Akylbekova, E. L., Aldrich, M. C., Ambrosone, C. B., Amos, C., Bandera, E. V., Berndt, S. I., Bernstein, L., Blot, W. J., Bock, C. H., Boerwinkle, E., Cai, Q., Caporaso, N., Casey, G., Adrienne Cupples, L., Deming, S. L., Ryan Diver, W., Divers, J., Fornage, M., Gillanders, E. M., Glessner, J., Harris, C. C., Hu, J. J., Ingles, S. A., Isaacs, W., John, E. M., Linda Kao, W. H., Keating, B., Kittles, R. A., Kolonel, L. N., Larkin, E., Le Marchand, L., McNeill, L. H., Millikan, R. C., Murphy, Musani, S., Neslund-Dudas, C., Nyante, S., Papanicolaou, G. J., Press, M. F., Psaty, B. M., Reiner, A. P., Rich, S. S., Rodriguez-Gil, J. L., Rotter, J. I., Rybicki, B. A., Schwartz, A. G., Signorello, L. B., Spitz, M., Strom, S. S., Thun, M. J., Tucker, M. A., Wang, Z., Wiencke, J. K., Witte, J. S., Wrensch, M., Wu, X., Yamamura, Y., Zanetti, K. A., Zheng, W., Ziegler, R. G., Zhu, X., Redline, S., Hirschhorn, J. N., Henderson, B. E., Taylor Jr, H. A., Price, A. L., Hakonarson, H., Chanock, S. J., Haiman, C. A., Wilson, J. G., Reich, D., and Myers, S. R. (2011). The landscape of recombination in African Americans. *Nature*, 476(7359):170–175.

Hudson, R. R., Slatkin, M., and Maddison, W. P. (1992). Estimation of levels of gene flow from DNA sequence data. *Genetics*, 132(2):583–589.

Iasi, L. N. M., Ringbauer, H., and Peter, B. M. (2021). An Extended Admixture Pulse Model Reveals the Limitations to Human–Neandertal Introgression Dating. *Molecular Biology and Evolution*, 38(11):5156–5174.

Jacobs, G. S., Hudjashov, G., Saag, L., Kusuma, P., Darusallam, C. C., Lawson, D. J., Mondal, M., Pagani, L., Ricaut, F.-X., Stoneking, M., Metspalu, M., Sudoyo, H., Lansing, J. S., and Cox, M. P. (2019). Multiple Deeply Divergent Denisovan Ancestries in Papuans. *Cell*, 177(4):1010–1021.e32.

Jónsson, H., Sulem, P., Kehr, B., Kristmundsdottir, S., Zink, F., Hjartarson, E., Hardarson, M. T., Hjorleifsson, K. E., Eggertsson, H. P., Gudjonsson, S. A., Ward, L. D., Arnadottir, G. A., Helgason, E. A., Helgason, H., Gylfason, A., Jonasdottir, A., Jonasdottir, A., Rafnar, T., Frigge, M., Stacey, S. N., Th. Magnusson, O., Thorsteinsdottir, U., Masson, G., Kong, A., Halldorsson,

B. V., Helgason, A., Gudbjartsson, D. F., and Stefansson, K. (2017). Parental influence on human germline de novo mutations in 1,548 trios from Iceland. *Nature*, 549(7673):519–522.

Kamm, J., Terhorst, J., Durbin, R., and Song, Y. S. (2020). Efficiently Inferring the Demographic History of Many Populations With Allele Count Data. *Journal of the American Statistical Association*, 115(531):1472–1487.

Keightley, P. D. and Jackson, B. C. (2018). Inferring the Probability of the Derived vs. the Ancestral Allelic State at a Polymorphic Site. *Genetics*, 209(3):897–906.

Keinan, A., Mullikin, J. C., Patterson, N., and Reich, D. (2009). Accelerated genetic drift on chromosome X during the human dispersal out of Africa. *Nature Genetics*, 41(1):66–70.

Kelleher, J., Etheridge, A. M., and McVean, G. (2016). Efficient Coalescent Simulation and Genealogical Analysis for Large Sample Sizes. *PLOS Computational Biology*, 12(5):e1004842.

Kong, A., Thorleifsson, G., Gudbjartsson, D. F., Masson, G., Sigurdsson, A., Jonasdottir, A., Walters, G. B., Jonasdottir, A., Gylfason, A., Kristinsson, K. T., Gudjonsson, S. A., Frigge, M. L., Helgason, A., Thorsteinsdottir, U., and Stefansson, K. (2010). Fine-scale recombination rate differences between sexes, populations and individuals. *Nature*, 467(7319):1099–1103.

Kuhlwilm, M., Gronau, I., Hubisz, M. J., de Filippo, C., Prado-Martinez, J., Kircher, M., Fu, Q., Burbano, H. A., Lalueza-Fox, C., de la Rasilla, M., Rosas, A., Rudan, P., Brajkovic, D., Kucan, Ž., Gušic, I., Marques-Bonet, T., Andrés, A. M., Viola, B., Pääbo, S., Meyer, M., Siepel, A., and Castellano, S. (2016). Ancient gene flow from early modern humans into Eastern Neanderthals. *Nature*, 530(7591):429–433.

Li, H. and Durbin, R. (2011). Inference of human population history from individual whole-genome sequences. *Nature*, 475(7357):493–496.

Malaspinas, A.-S., Westaway, M. C., Muller, C., Sousa, V. C., Lao, O., Alves, I., Bergström, A., Athanasiadis, G., Cheng, J. Y., Crawford, J. E., Heupink, T. H., Macholdt, E., Peischl, S., Rasmussen, S., Schiffels, S., Subramanian, S., Wright, J. L., Albrechtsen, A., Barbieri, C., Dupanloup, I., Eriksson, A., Margaryan, A., Moltke, I., Pugach, I., Korneliussen, T. S., Levkivskyi, I. P., Moreno-Mayar, J. V., Ni, S., Racimo, F., Sikora, M., Xue, Y., Aghakhanian, F. A., Brucato, N., Brunak, S., Campos, P. F., Clark, W., Ellingvåg, S., Fourmile, G., Gerbault, P., Injie, D., Koki, G., Leavesley, M., Logan, B., Lynch, A., Matisoo-Smith, E. A., McAllister, P. J., Mentzer, A. J., Metspalu, M., Migliano, A. B., Murgha, L., Phipps, M. E., Pomat, W., Reynolds, D., Ricaut, F.-X., Siba, P., Thomas, M. G., Wales, T., Wall, C. M., Oppenheimer, S. J., Tyler-Smith, C., Durbin, R., Dortch, J., Manica, A., Schierup, M. H., Foley, R. A., Lahr, M. M., Bowern, C., Wall, J. D., Mailund, T., Stoneking, M., Nielsen, R., Sandhu, M. S., Excoffier, L., Lambert, D. M., and Willerslev, E. (2016). A genomic history of Aboriginal Australia. *Nature*, 538(7624):207–214.

Miles, A., io bot, p., R, M., Ralph, P., Harding, N., Pisupati, R., Rae, S., and Millar, T. (2021). Cggh/scikit-allel: V1.3.3. Zenodo.

Moorjani, P., Patterson, N., Hirschhorn, J. N., Keinan, A., Hao, L., Atzmon, G., Burns, E., Ostrer, H., Price, A. L., and Reich, D. (2011). The History of African Gene Flow into Southern Europeans, Levantines, and Jews. *PLOS Genetics*, 7(4):e1001373.

Moorjani, P., Sankararaman, S., Fu, Q., Przeworski, M., Patterson, N., and Reich, D. (2016). A genetic method for dating ancient genomes provides a direct estimate of human generation interval in the last 45,000 years. *Proceedings of the National Academy of Sciences*, 113(20):5652–5657.

Myers, S., Bottolo, L., Freeman, C., McVean, G., and Donnelly, P. (2005). A Fine-Scale Map of Recombination Rates and Hotspots Across the Human Genome. *Science*, 310(5746):321–324.

Ning, Z., Cox, A. J., and Mullikin, J. C. (2001). SSAHA: A Fast Search Method for Large DNA Databases. *Genome Research*, 11(10):1725–1729.

Patterson, N., Moorjani, P., Luo, Y., Mallick, S., Rohland, N., Zhan, Y., Genschoreck, T., Webster, T., and Reich, D. (2012). Ancient Admixture in Human History. *Genetics*, 192(3):1065–1093.

Prado-Martinez, J., Sudmant, P. H., Kidd, J. M., Li, H., Kelley, J. L., Lorente-Galdos, B., Veeramah, K. R., Woerner, A. E., O'Connor, T. D., Santpere, G., Cagan, A., Theunert, C., Casals, F., Laayouni, H., Munch, K., Hobolth, A., Halager, A. E., Malig, M., Hernandez-Rodriguez, J., Hernando-Herraez, I., Prüfer, K., Pybus, M., Johnstone, L., Lachmann, M., Alkan, C., Twigg, D., Petit, N., Baker, C., Hormozdiari, F., Fernandez-Callejo, M., Dabad, M., Wilson, M. L., Stevison, L., Camprubí, C., Carvalho, T., Ruiz-Herrera, A., Vives, L., Mele, M., Abello, T., Kondova, I., Bontrop, R. E., Pusey, A., Lankester, F., Kiyang, J. A., Bergl, R. A., Lonsdorf, E., Myers, S., Ventura, M., Gagneux, P., Comas, D., Siegismund, H., Blanc, J., Agueda-Calpena, L., Gut, M., Fulton, L., Tishkoff, S. A., Mullikin, J. C., Wilson, R. K., Gut, I. G., Gonder, M. K., Ryder, O. A., Hahn, B. H., Navarro, A., Akey, J. M., Bertranpetit, J., Reich, D., Mailund, T., Schierup, M. H., Hvilsom, C., Andrés, A. M., Wall, J. D., Bustamante, C. D., Hammer, M. F., Eichler, E. E., and Marques-Bonet, T. (2013). Great ape genetic diversity and population history. *Nature*, 499(7459):471–475.

Prüfer, K., de Filippo, C., Grote, S., Mafessoni, F., Korlević, P., Hajdinjak, M., Vernot, B., Skov, L., Hsieh, P., Peyrégne, S., Reher, D., Hopfe, C., Nagel, S., Maricic, T., Fu, Q., Theunert, C., Rogers, R., Skoglund, P., Chintalapati, M., Dannemann, M., Nelson, B. J., Key, F. M., Rudan, P., Kućan, Ž., Gušić, I., Golovanova, L. V., Doronichev, V. B., Patterson, N., Reich, D., Eichler, E. E., Slatkin, M., Schierup, M. H., Andrés, A. M., Kelso, J., Meyer, M., and Pääbo, S. (2017). A high-coverage Neandertal genome from Vindija Cave in Croatia. *Science*, 358(6363):655–658.

Prüfer, K., Racimo, F., Patterson, N., Jay, F., Sankararaman, S., Sawyer, S., Heinze, A., Renaud, G., Sudmant, P. H., de Filippo, C., Li, H., Mallick, S., Dannemann, M., Fu, Q., Kircher, M., Kuhlwilm, M., Lachmann, M., Meyer, M., Ongyerth, M., Siebauer, M., Theunert, C., Tandon, A., Moorjani, P., Pickrell, J., Mullikin, J. C., Vohr, S. H., Green, R. E., Hellmann, I., Johnson, P. L. F., Blanche, H., Cann, H., Kitzman, J. O., Shendure, J., Eichler, E. E., Lein, E. S., Bakken, T. E., Golovanova, L. V., Doronichev, V. B., Shunkov, M. V., Derevianko, A. P., Viola, B., Slatkin, M., Reich, D., Kelso, J., and Pääbo, S. (2014). The complete genome sequence of a Neanderthal from the Altai Mountains. *Nature*, 505(7481):43–49.

Ragsdale, A. P. and Gravel, S. (2019). Models of archaic admixture and recent history from two-locus statistics. *PLOS Genetics*, 15(6):e1008204.

Sankararaman, S., Mallick, S., Dannemann, M., Prüfer, K., Kelso, J., Pääbo, S., Patterson, N., and Reich, D. (2014). The genomic landscape of Neanderthal ancestry in present-day humans. *Nature*, 507(7492):354–357.

Sankararaman, S., Patterson, N., Li, H., Pääbo, S., and Reich, D. (2012). The Date of Interbreeding between Neandertals and Modern Humans. *PLOS Genetics*, 8(10):e1002947.

Scerri, E. M. L., Chikhi, L., and Thomas, M. G. (2019). Beyond multiregional and simple out-of-Africa models of human evolution. *Nature Ecology & Evolution*, 3(10):1370–1372.

Schaefer, N. K., Shapiro, B., and Green, R. E. (2021). An ancestral recombination graph of human, Neanderthal, and Denisovan genomes. *Science Advances*, 7(29):eabc0776.

Skoglund, P., Malmström, H., Raghavan, M., Storå, J., Hall, P., Willerslev, E., Gilbert, M. T. P., Götherström, A., and Jakobsson, M. (2012). Origins and Genetic Legacy of Neolithic Farmers and Hunter-Gatherers in Europe. *Science*, 336(6080):466–469.

Skov, L., Coll Macià, M., Sveinbjörnsson, G., Mafessoni, F., Lucotte, E. A., Einarsdóttir, M. S., Jonsson, H., Halldorsson, B., Gudbjartsson, D. F., Helgason, A., Schierup, M. H., and Stefansson, K. (2020). The nature of Neanderthal introgression revealed by 27,566 Icelandic genomes. *Nature*, 582(7810):78–83.

Spence, J. P. and Song, Y. S. (2019). Inference and analysis of population-specific fine-scale recombination maps across 26 diverse human populations. *Science Advances*, 5(10):eaaw9206.

Sudmant, P. H., Rausch, T., Gardner, E. J., Handsaker, R. E., Abyzov, A., Huddleston, J., Zhang, Y., Ye, K., Jun, G., Hsi-Yang Fritz, M., Konkel, M. K., Malhotra, A., Stütz, A. M., Shi, X., Paolo Casale, F., Chen, J., Hormozdiari, F., Dayama, G., Chen, K., Malig, M., Chaisson, M. J. P., Walter, K., Meiers, S., Kashin, S., Garrison, E., Auton, A., Lam, H. Y. K., Jasmine Mu, X., Alkan, C., Antaki, D., Bae, T., Cerveira, E., Chines, P., Chong, Z., Clarke, L., Dal, E., Ding, L., Emery, S., Fan, X., Gujral, M., Kahveci, F., Kidd, J. M., Kong, Y., Lameijer, E.-W., McCarthy, S., Flicek, P., Gibbs, R. A., Marth, G., Mason, C. E., Menelaou, A., Muzny, D. M., Nelson, B. J., Noor, A., Parrish, N. F., Pendleton, M., Quitadamo, A., Raeder, B., Schadt, E. E., Romanovitch, M., Schlattl, A., Sebra, R., Shabalin, A. A., Untergasser, A., Walker, J. A., Wang, M., Yu, F., Zhang, C., Zhang, J., Zheng-Bradley, X., Zhou, W., Zichner, T., Sebat, J., Batzer, M. A., McCarroll, S. A., Mills, R. E., Gerstein, M. B., Bashir, A., Stegle, O., Devine, S. E., Lee, C., Eichler, E. E., and Korbel, J. O. (2015). An integrated map of structural variation in 2,504 human genomes. *Nature*, 526(7571):75–81.

Williamson, S. H., Hernandez, R., Fledel-Alon, A., Zhu, L., Nielsen, R., and Bustamante, C. D. (2005). Simultaneous inference of selection and population growth from patterns of variation in the human genome. *Proceedings of the National Academy of Sciences of the United States of America*, 102(22):7882–7887.

Yang, M. A., Malaspinas, A.-S., Durand, E. Y., and Slatkin, M. (2012). Ancient Structure in Africa Unlikely to Explain Neanderthal and Non-African Genetic Similarity. *Molecular Biology and Evolution*, 29(10):2987–2995.

Zeng, K., Jackson, B. C., and Barton, H. J. (2019). Methods for Estimating Demography and Detecting Between-Locus Differences in the Effective Population Size and Mutation Rate. *Molecular Biology and Evolution*, 36(2):423–433.
